# The genomic diversity of Taiwanese Austronesian groups: implications for the ‘Into and Out of Taiwan’ models

**DOI:** 10.1101/2023.01.09.523210

**Authors:** Dang Liu, Albert Min-Shan Ko, Mark Stoneking

## Abstract

The origin and dispersal of the Austronesian language family, one of the largest and most widespread in the world, have long attracted the attention of linguists, archaeologists, and geneticists. Even though there is a growing consensus that Taiwan is the source of the spread of Austronesian languages, little is known about the migration patterns of the early Austronesians who settled in and left Taiwan, i.e., the “Into-Taiwan” and “Out-of-Taiwan” events. In particular, the genetic diversity/structure within Taiwan and how this relates to the Into/Out-of-Taiwan events is largely unexplored, because most genomic studies have largely utilized data from just two of the 16 recognized highland Austronesian groups in Taiwan. In this study, we generated the largest genome-wide dataset for Taiwanese Austronesians to date, including six highland groups and one lowland group from across the island, and two Taiwanese Han groups. We identified fine scale genomic structure in Taiwan, inferred the ancestry profile of the ancestors of Austronesians, and found that the southern Taiwanese Austronesians show excess genetic affinities toward the Austronesians outside of Taiwan. Our findings thus shed new light on the Into and Out-of-Taiwan dispersals.

## Introduction

Austronesian is one of the largest language families in the world, with more than 1200 languages spoken by almost 400 million people, spread from Madagascar in the west to Hawaii and Easter Island in the east (Eberhard, et al. 2021). Linguistic analyses strongly support a Taiwanese origin for Austronesian languages (Blust 1999; Gray, et al. 2009), and archaeological and genetic evidence further support an expansion of people from Taiwan associated with the spread of Austronesian languages (Diamond and Bellwood 2003; Bellwood 2011; Ko, et al. 2014; Lipson, et al. 2014). For this reason, the migration events “Into-Taiwan,” which detail the arrival of the ancestors of Taiwanese Austronesians on the island, and “Out-of-Taiwan,” which detail the departure of the same ancestors for other Austronesian groups, are of great importance. Regarding “Into-Taiwan”, based on the distribution of millet and rice in ancient sites, archeologists estimate that the ancestors of Taiwanese Austronesian arrived in Taiwan from the southeastern coast of mainland China ∼4.8 thousand years ago (kya) at the latest (Deng, et al. 2022). Linguistically, these people are considered “proto-Austronesians”, and their language shares features with the Tai-Kadai and Sino-Tibetan languages spoken in southeastern coast of China (Sagart 2008). Recent ancient DNA studies also find strong genetic links between the Taiwanese Austronesians and the ancient individuals from southern China (associated with the Neolithic agricultural culture) (Yang, et al. 2020; Wang, Yeh, et al. 2021). As for “Out-of-Taiwan”, archaeological evidence suggests that the agricultural complex associated with Austronesian ancestors began expanding from Taiwan into the Philippines ∼ 4.2 kya (Hung, et al. 2022) and then rapidly throughout Indonesia, west to Madagascar and east across the Pacific (Bellwood 2011; Kirch 2017). Among the ten divisions of the Austronesian language family recognized by linguists, nine (Formosan branches) are found only in Taiwan, while the remaining Austronesian languages outside Taiwan are grouped under the Malayo-Polynesian branch (Blust 2013). Phylogenetic analyses of Austronesian languages have also supported an origin in Taiwan and estimated that the Formosan and Malayo-Polynesian branches diverged ∼5 kya (Gray, et al. 2009). Genetic studies have found evidence for a “Out-of-Taiwan” migration, however with estimated dates ranging from 4 to 8 kya (Ko, et al. 2014; Soares, et al. 2016; Choin, et al. 2021; Larena, et al. 2021). Additionally, the expansion of Austronesian peoples are associated with the development of the Lapita culture (3.5-2.5 kya) in Near Oceania and the spread into Remote Oceania (Spriggs 1995; Kirch 2017); surprisingly, genomes from ancient individuals of Remote Oceania related to the Lapita have revealed a strong genetic link to the Taiwanese Austronesian groups (Skoglund, et al. 2016; Posth, et al. 2018; Lipson, et al. 2020), suggesting these individuals were among the earliest “Out-of-Taiwan” groups.

While there has been much investigation of the “Into-Taiwan” and “Out-of-Taiwan” migrations, the diversity and structure of Austronesian groups within Taiwan remains relatively unexplored. Most genome-wide studies have used data are from only two groups, the Amis and Atayal (Soares, et al. 2016; Lipson, et al. 2020; Larena, et al. 2021), when in fact there are over 20 recognized indigenous (Austronesian) groups in Taiwan according to the Council of Indigenous People (CIP) in Taiwan (https://www.cip.gov.tw; last accessed 12 December 2022). These are broadly classified as the “Highland” or “Lowland (called Pingpu in Mandarin)” groups based on where they live, although some of the “Highland” groups do not reside in the mountainous area. Presently, there are 16 officially defined indigenous “Highland” groups (the Atayal, Saysiyat, Truku, Sediq, Sakizaya, Thao, Tsou, Kavalan, Bunun, Hla’alua, Kanakanavu, Amis, Rukai, Puyuma, Paiwan, and the Tao); however, the Tao (or Yami) actually reside on Orchid Island, which is located off the southeastern coast of Taiwan. And while all the Highland groups speak Formosan languages, the Yami language is part of the Malayo-Polynesian branch (Blust 2013; Eberhard, et al. 2021). There are another 12 or so identified “Lowland” groups (the Ketagalan, Kavalan, Taokas, Kaxabu, Pazeh, Papora, Babuza, Lloa, Arikun, Siraya, Taivoan, and the Makatao); extensive contact with Han peoples (primarily Minnan and Hakka) who migrated from mainland China in the past 500 years has led to the extinction or endangerment of “Lowland” Austronesian languages (also described on the CIP website).

The “Highland” groups of Taiwan have also been the primary focus of genetic studies, while “Lowland” groups have received much less attention. There is evidence of genetic diversity among Taiwanese Austronesians, as shown by studies of “Highland” groups using uniparental genetic makers (Melton, et al. 1998; Tajima, et al. 2003; Ko, et al. 2014; Trejaut, et al. 2014; Trejaut, et al. 2019), though few of these studies included “Lowland” groups (Ko, et al. 2014; Trejaut, et al. 2014; Trejaut, et al. 2019). Furthermore, genome-wide studies, which can provide much more detailed insights into population histories, are very limited. To our knowledge, genome-wide data have only been generated for the Atayal, Amis, Paiwan, and Tao (Yami) (Patterson, et al. 2012; Mallick, et al. 2016; Choin, et al. 2021; Tatte, et al. 2021), and a systematic genomic assessment of the diversity and relationships among Taiwan groups, and how these relate to the “Into Taiwan” and “Out-of-Taiwan” migrations, is still lacking.

To remedy this, we generated new genome-wide data for 7 Taiwanese Austronesian groups from across the island (3 with no genome-wide data reported before, including one “Lowland” group) and 2 Taiwanese Han groups to investigate fine scale structure within Taiwan. We used the CIP’s definition of Taiwan Highland/Taiwan Orchid Island (THI/TOI) to categorize the Highlands’ various groups. Combing our new data with published comparative modern and ancient genomes, we leverage the diversity of Taiwanese Austronesians to gain new insights for the Into- and Out-of-Taiwan events.

## Results

### Taiwanese groups in the light of genetic variation in South Asia, East Asia and Oceania

We generated genome-wide single nucleotide polymorphism (SNP) array data on the Affymetrix Human Origins array for 55 individuals from 7 Austronesian (5 Highland, 1 Lowland, and 1 Orchid Island) and 2 Han groups from Taiwan (Fig. 1A). We merged the new data with comparative modern and ancient genomes from Asia and Oceania (Fig. S1).

**Fig. 1.**
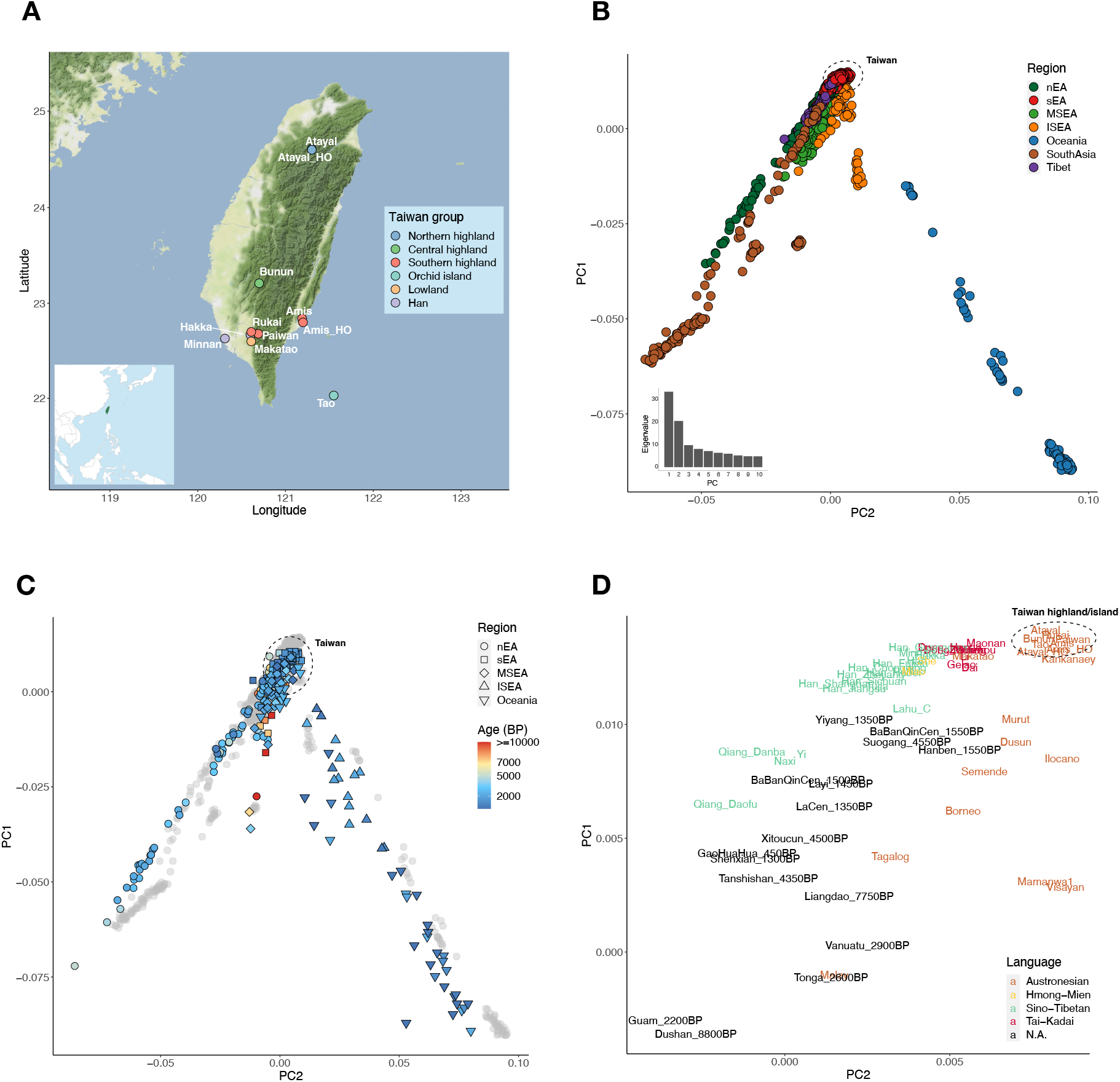
Map of sampled Taiwanese groups and PCA of the modern/ancient individuals from South Asia, East Asia, and Oceania. (A) Sampling locations of Taiwanese populations. Atayal_HO and Amis_HO are from published Human Origins data. The small map panel on the bottom-left indicates Taiwan (green) in the geographical context of East Asia. (B) PCA of modern individuals from South Asia, East Asia, and Oceania in the merged dataset, colored by regions. The dashed circle indicates where the Taiwanese individuals fall. The eigenvalues from PC1 to PC10 are shown on the bottom-left. (C) PCA with ancient individuals projected onto the PCA in (B) with modern individuals in grey. Symbol shapes indicate region and colors indicate age (D) Zoom-in of (C) with a focus on modern groups from sEA and ISEA and ancient groups from sEA and Lapita-related Oceania, colored by languages. The position of a group is the median of the positions of individuals from the group.

We first investigated overall patterns in the dataset using Principal Component Analysis (PCA) and ADMIXTURE (Alexander, et al. 2009) /DyStruct (Joseph and Pe’er 2019) analyses. In the PCA, East Asians (southern East Asians, sEA; northern East Asians, nEA; Mainland Southeast Asians, MSEA; and Island Southeast Asians, ISEA) are separated from South Asians and Oceanians. sEA/nEA is defined by the south/north of the Qinling-Huaihe (Yang, et al. 2020). Along PC1, sEA is at the extreme of the East Asian pole, ISEA are on the cline toward Oceanians, and the remaining East Asian groups are on the cline towards South Asians. The genetic profiles of all Taiwanese individuals fall within the genetic variation of sEA (Fig. 1B and 1C), with the Taiwan Austronesian groups on the extreme of the sEA pole together with the Filipino Austronesian group Kankanaey (Fig. 1D). Ancient individuals falling closest to Taiwanese groups are the sEA and early (∼2-3 kya) Oceanian individuals (Fig. 1D).

For the best fitting K value of the ADMIXTURE analysis (Fig. S2A), Atayal and Kankanaey have the highest frequencies of a purple component which is also enriched in Taiwanese Highland/Orchid Island (THI/TOI) groups as well as ancient sEA (e.g., ∼4.5 kya Suogang and ∼4.3 kya Tanshishan) and Oceanian (e.g., ∼2.9 kya Vanuatu and ∼2.6 kya Tonga) individuals (Fig. 2; Fig. S3A). In contrast, the Taiwanese Han groups are similar to the Han from Fujian in southern China, with two major components: light green and turquoise. In comparison to Han from Shandong of northern China, Taiwanese and Fujian Han groups show small amounts of the purple Austronesian-related component, suggesting potential interactions between the sEA Han and Austronesian groups. Similarly, the Taiwanese lowland Austronesian group, Makatao, has a similar profile to the Taiwanese Han groups but with more of the purple Austronesian-related component. The best fitting K value for the DyStruct analysis, an ADMIXTURE-like clustering algorithm that incorporates time transects and is hence more suitable for ancient DNA, indicates similar results for the THI/TOI being close to the Kankanaey and ancient sEA individuals. They share a relatively homogenous pattern of two components (Fig. 2; Fig. S3B): a brown one at highest frequency in ∼4.5 kya sEA Suogang (likely proto-Austronesian) and ∼2.9 kya Vanuatu individuals (likely early Austronesians), which is associated with all modern Austronesian groups in ISEA and Oceania, suggesting a link between all of them; and a grey one at highest frequency in the Tai-Kadai speaking Li and widespread among present day East Asians.

**Fig. 2.**
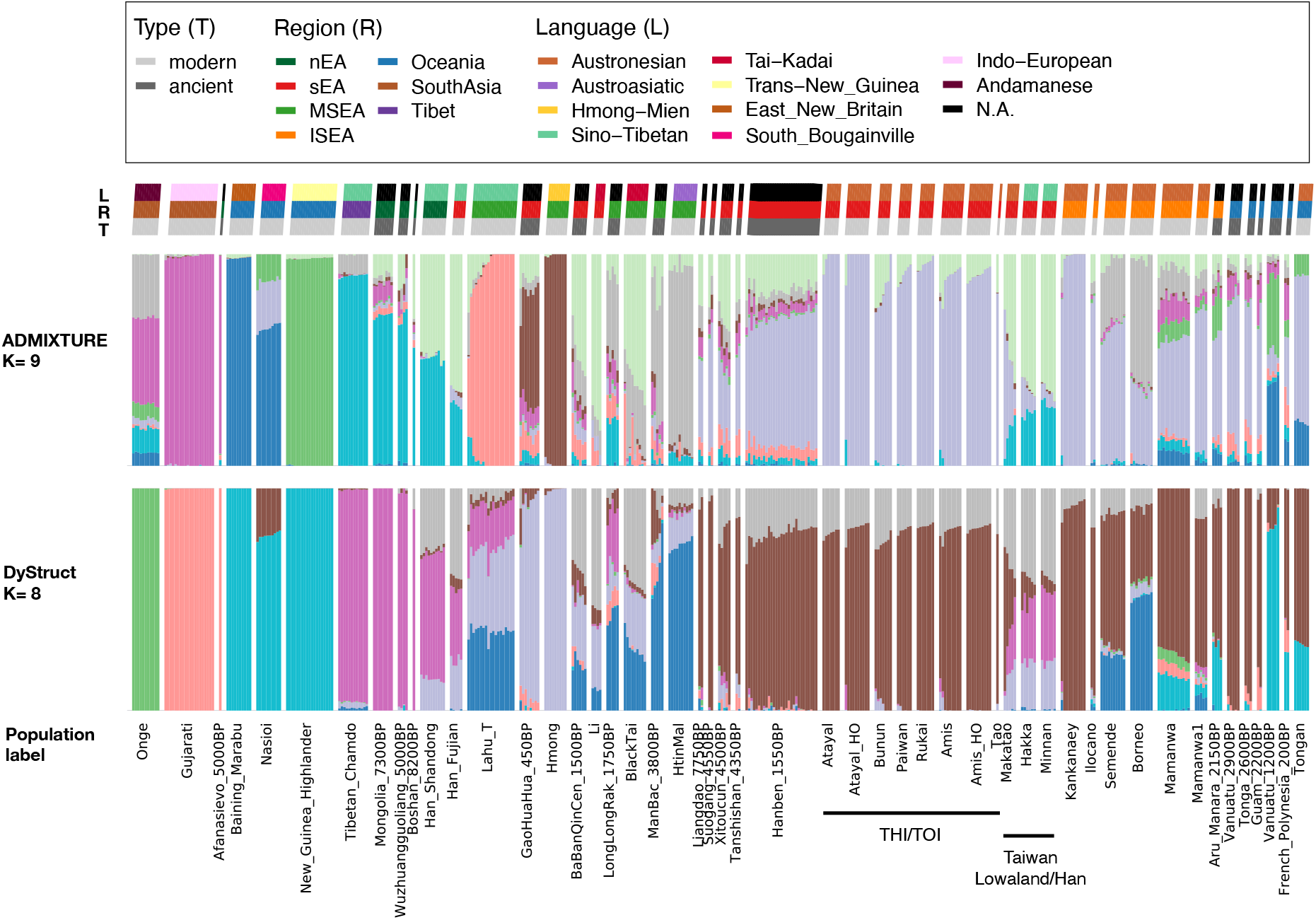
K = 9 of ADMIXTURE and K = 8 of DyStruct for selected representative groups. Both number of K are the best-fitting K for ADMIXTURE/DyStruct (Fig. S2). From the top to the bottom: there is a legend for the following three rows of keys indicating the language (L), the region (L), and the type of ancient/modern genomes (T) for the populations; below the keys, there are two rows of the results of ADMIXTURE and DyStruct, respectively; the last row indicates the specific population label. Each vertical thin bar represents an individuals, and different populations are separated with gaps. Representative groups are selected based on their enrichment of a source component, e.g., Andamanese, Indo-European, Papuan (Trans-New Guinea, East New Britain, South Bougainville), Sinto-Tibetan/nEA, Hmong-Mien, Austroasiatic; or their relevance to Into/Out-of-Taiwan events, e.g., Tai-Kadai, sEA, Taiwan, ISEA, Oceania. Results of the full dataset are presented in Fig. S3.

To investigate this grey component further, we noticed that the grey component is at higher frequency in the Taiwan Lowland/Han groups, and the light/dark purple components are also at a higher frequency in these groups, which is similar to the pattern of the light green and turquoise components in ADMIXTURE; furthermore, all of these are enriched in Han groups (Fig. 2). We therefore compared the proportions of these components in Taiwanese groups (ADMIXTURE light green vs. turquoise; DyStruct grey + light purple vs. dark purple) and found a correlation between them (r^2^=0.913 and 0.926, respectively; both p=0), suggesting they might be introduced via a single Han admixture event (Fig. S4). Moreover, the ratio of these components is correlated with the geographical distribution of Han groups (northern groups have higher turquoise/dark purple compared to southern groups), and the ratio in Taiwanese Han is close to that of the southern Chinese Han groups, while the cline of Taiwanese Austronesian groups points toward the Taiwanese Han (Fig. S4).

### Genetic structure within Taiwan

In the context of Asian and Oceanian genetic variation, the genetic profile of THI/TOI is relatively homogenous and there is little structure revealed, except that they are separated from the Han groups, with the Lowland group falling in between them (Fig. 3A). To further study the structure within Taiwan, we applied the haplotype-based method fineSTRUCTURE (Lawson, et al. 2012). To provide haplotype sources from neighbouring groups, we carried out chromosome painting on all Austronesian, Tai-Kadai, and Sino-Tibetan groups in our dataset (Fig. 3B), as these language families are proposed to be related (Sagart 2008). The fineSTRUCTURE results generally support clustering according to language family except that groups from MSEA show a heterogenous pattern, which is consistent with previous studies suggesting complex histories involving extensive admixture and probable cases of language shift (Liu, et al. 2020; Kutanan, et al. 2021). Within Taiwan, there is a clear separation between THI/TOI and Lowland/Han groups; the former clusters with other Austronesian groups while the latter cluster with Sino-Tibetan and Tai-Kadai groups (Fig. 3B). With respect to THI/TOI groups, the northern (Atayal) and central (Bunun) groups cluster in a clade; there is a division in the southern groups (Rukai, Paiwan, Amis): Rukai and Paiwan cluster together, while Amis cluster in another clade with the Orchid Island group (Tao) and the Filipino groups Kankanaey and Ilocano.

**Fig. 3.**
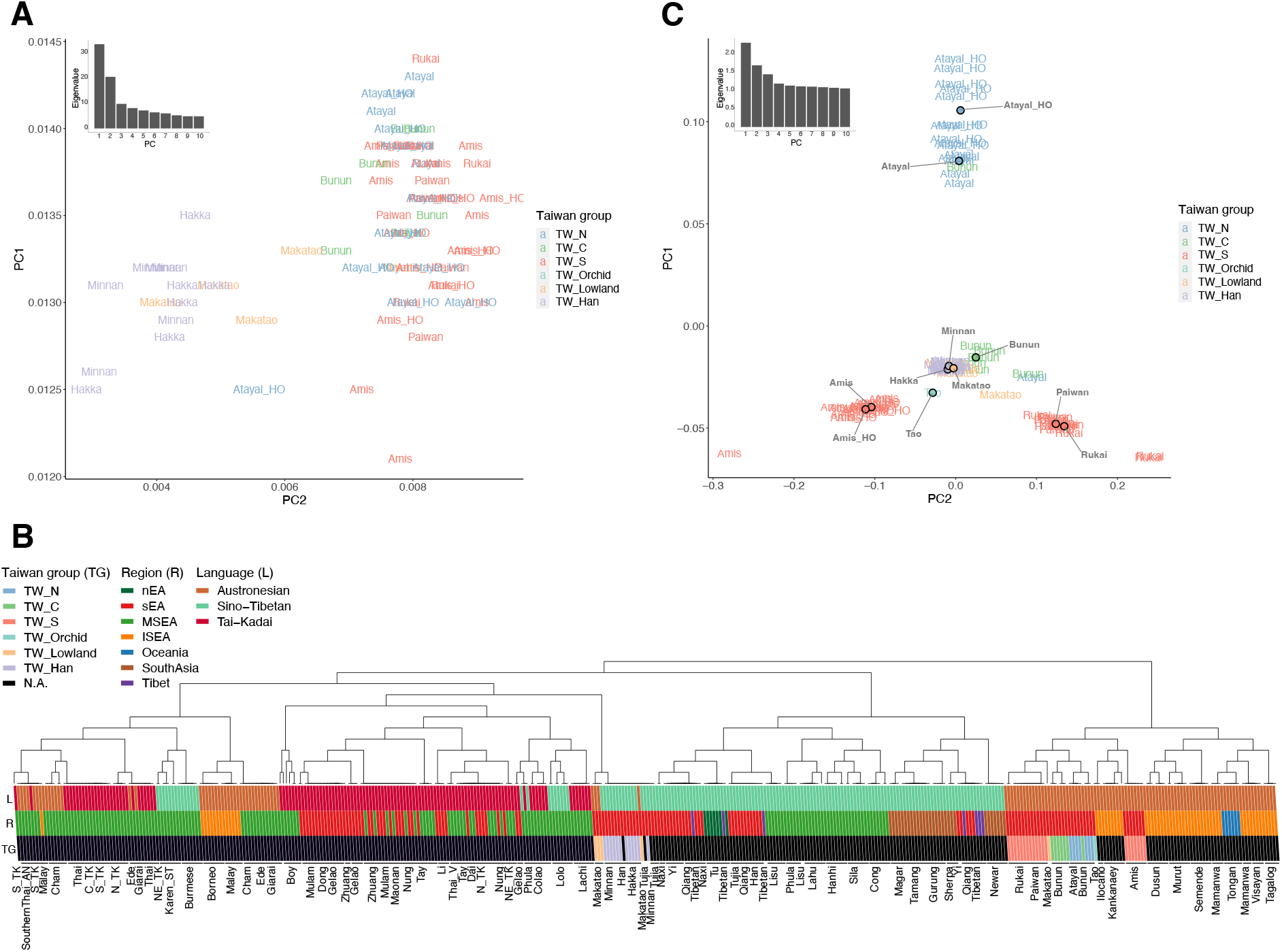
Structure within Taiwan. (A) Zoom-in of the PCA plot in Fig. 1B with a focus on Taiwanese individuals, labeled by populations and colored by groups. (B) FineStructure clustering of Sino-Tibetan, Tai-Kadai, and Austronesian groups based on haplotype painting profiles from ChromoPainter. Each vertical line represents an individual, colored according to regions (R), Taiwan groups (TG), and languages (L). Populations are labeled at the bottom. Note that the clustering does not necessarily imply phylogenetic structure. (C) PCA for which eigenvalues were computed using only THI/TOI individuals, with the Han and Lowland individuals projected. Individuals are labeled by groups and the median positions of individuals of each group are plotted with dots colored by groups. The eigenvalues from PC1 to PC10 are shown on the top-left. TW_N, TW_C, and TW_S denote northern, central, southern Taiwanese highland groups, respectively.

As there is structure revealed in THI/TOI groups, distinct from Lowland/Han groups, we performed a PCA with eigenvalues computed using only THI/TOI individuals and projected the Lowland/Han individuals (Fig. 3C). PC1 separates the northern group Atayal from southern groups and PC2 separates Amis from Rukai and Paiwan. The central group Bunun falls toward Atayal, the Orchid Island group Tao falls toward Amis, and the lowland group Makatao is heterogeneous, with most falling with Han groups in the middle, but some lying toward Rukai/Paiwan. Notably, the Atayal and Amis from the published dataset fall together with the Atayal and Amis genotyped in this study. The fact that Atayal, Amis, and Rukai drive the poles of this PCA probably reflects their isolation and drift, as they show high amounts of within-group IBD sharing (Fig. S5).

Both ADMIXTURE and PCA results show that the Lowland group Makatao has an intermediate genetic profile between the THI/TOI and Han groups (Figs. 2 and 3A), which suggests admixture. To test this, we computed the f3 admixture value f3(Minnan, Rukai; Makatao) and obtained Z= −7, indicating admixture in the Makatao involving these two proxy sources. Rukai is chosen as the Austronesian source proxy because the Makatao is projected toward the Rukai pole in the PCA of THI/TOI (Fig. 3C). This is further supported by a best-fitting admixture graph of Mbuti (outgroup), the Lowland Makatao, Han groups, and the three representative THI groups (Atayal, Amis, Rukai): the Makatao is modelled as an admixture of 42% ancestry from the ancestor of the THI group Rukai and 58% from the ancestor of the Han group Minnan (Fig. S6). Using GLOBETROTTER (Hellenthal, et al. 2014), we inferred the admixture as a single pulse event dating to ∼1.64 +/- 0.93 generations ago (∼50 years ago assuming 30 years per generation). Overall, we observe three distinct genetic clusters in the THI/TOI groups and an admixture event between the THI and Han groups in the Lowland group.

### Genetic structure of Taiwanese Austronesian groups: implications for Into-Taiwan

With a better understanding of the genetic structure within Taiwan, we can investigate the genetic profiles of the ancestors of Taiwanese Austronesians, or proto-Austronesians. We first focused on ancient genomes from or nearby Taiwan: Hanben from north-eastern Taiwan dated ∼1.5 kya; Suogang from Penghu Island offshore from south-western Taiwan, dated ∼4.5 kya; and Liangdao from Liangdao Island offshore from north-western Taiwan, dated ∼7.7 kya (Yang, et al. 2020; Wang, Yeh, et al. 2021). Among all projected ancient genomes on the PCA of THI/TOI, the Suogang and Liangdao individuals, who are thought to be related to proto-Austronesians (Yang, et al. 2020), position themselves toward the Rukai pole, while the Hanben individuals are closer to the Atayal pole (Fig. S7).

Previous studies used qpAdm with ancient genomes as sources and outgroups to model present day East Asians including Amis and Atayal (Yang, et al. 2020; Wang, Yeh, et al. 2021). However, Yang et al. modelled the Amis as a pure sEA source and the Atayal as an admixture of sEA and nEA sources while Wang et al. only included the Amis and modelled them as an admixture of sEA and nEA with different sources and outgroups. We investigated this and extended the analysis to the other Taiwanese groups in our study, using the sources and outgroups in Yang et al. as these were more clearly defined, with one modification (Fig. S8): we substituted the 7 kya Pha Faen genome (McColl, et al. 2018) (which overlaps with Onge in terms of Hoabinhian-related ancestry) with the 10 kya Longlin genome (Wang, Wang, et al. 2021) to provide more distinct sEA outgroups. For selected ancient groups, the ∼4.3 kya Tanshishan group from southern China, which is closely related to the ∼4.5 kya Suogang group (due to low coverage, we do not have enough power to model this group), is modelled as having mainly sEA (7.7 kya Liangdao) ancestry and a small proportion (∼8%) of nEA (8.2 kya Boshan) ancestry, while younger sEA and Taiwan groups have more (∼23-26%) nEA ancestry (Fig. 4A; Fig. S9). Likewise, we can model all the selected modern groups (including Taiwanese groups) as an admixture of ancient nEA and sEA ancestries, and the nEA ancestry has further increased (∼28-50%) in present day sEA samples (Fig. 4A; Fig. S9). The sEA ancestry is also present at low frequency in the ancient nEA groups and increases in present day samples, consistent with previous studies (Yang, et al. 2020; Wang, Yeh, et al. 2021). A best-fitting admixture graph supports that the ∼1.5 kya Hanben group is modelled as admixture of an nEA source (ancestor of 8.3 kya Boshan) and a sEA source (ancestor of 7.7 kya Liangdao); similarly, the THI Austronesians (Atayal, Amis, Rukai) and the Tai-Kadai speaking Li are modelled as an admixture of shared nEA (ancestor of Han from Shandong) and sEA (ancestor of 7.7 kya Liangdao) ancestries, while the Li receives additional sEA-related ancestry from an unsampled group (Fig. S10).

**Fig. 4.**
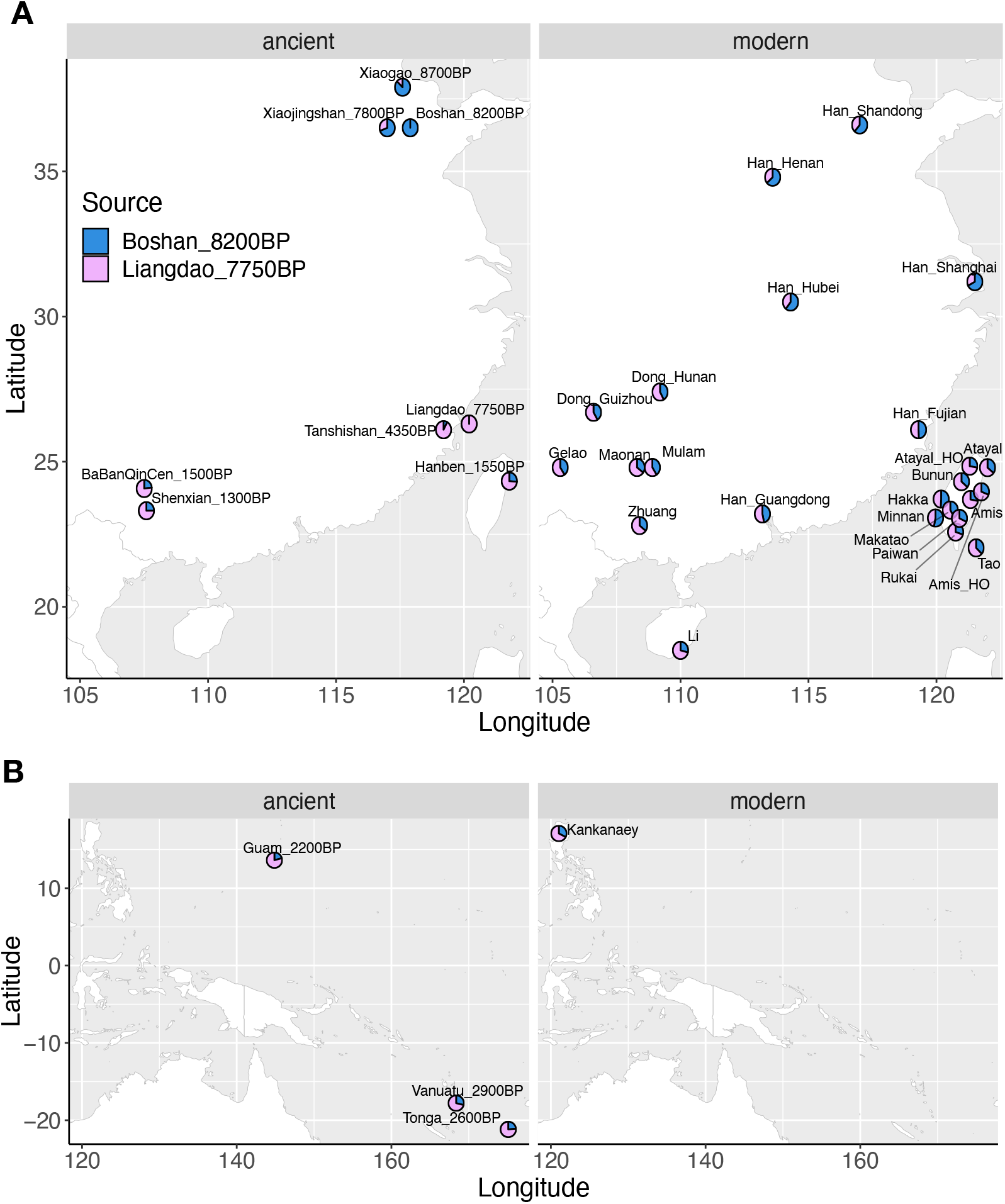
qpAdm modeling of the spatiotemporal dynamic of nEA vs. sEA ancestries in East Asia and Oceania. Using outgroups shown in Fig. S8, selected groups from (A) nEA and sEA as well as (B) ISEA and Oceania (with a focus on those related to early Out-of-Taiwan groups) are modelled with nEA (in blue, ∼8.2 kya Boshan as proxy) and sEA (in pink, ∼7.7 kya Liangdao as proxy) sources. The left panels show ancient groups while the right panels show modern groups. Bar plot visualization with the standard errors of the estimates is shown in Fig. S9.

### Genetic structure of Taiwanese Austronesian groups: implications for Out-of-Taiwan

We next investigated the relationships of Taiwanese Austronesian groups to groups in ISEA and Oceania, which would provide insights into the Out-of-Taiwan event(s). The Lapita-related ancient groups (∼2-3 kya Guam, Vanuatu, Tonga), who are thought to be related to the Out-of-Taiwan Austronesians, show a similar profile to the younger (∼1.5 kya) sEA ancient groups, while the present day Filipino Kankanaey, who are an isolated group often used as a modern proxy for the Out-of-Taiwan group, are more similar to the THI/TOI groups (Fig. 4B; Fig. S9). These together suggest that, intriguingly, early Out-of-Taiwan (Lapita) groups already possess increased nEA ancestry compared to early Into-Taiwan (sEA) groups (∼21-29% vs. ∼0-8%) and the nEA ancestry further slightly increased in present day THI/TOI (∼28-37%) and Kankanaey (∼33%) groups. Focussing on how these groups (as well as other ISEA and Oceanian groups) project on the THI/TOI PCA, we found that they are generally projected toward the southern groups Amis and Rukai and slightly more shifted to the Amis (Fig. S11). Results of f4 statistics of the form f4(Atayal, Amis/Rukai; modern/ancient groups, Mbuti) further support the observation that, compared to the northern group Atayal, the southern groups Amis and Rukai have excess sharing with modern/ancient ISEA and Oceanian groups (Fig. 5; Fig. S12). More precisely, the Atayal shows significant excess sharing with mainland sEA modern/ancient groups compared to the Amis (Fig. 5A and 5B) while the Rukai shows significant excess sharing with ISEA and Oceania groups compared to the Atayal (Fig. 5C and 5D). We finally tried to determine whether the ancestral out-of-Taiwan groups are closer to Amis or Rukai with an f4 statistic of the form f4(Amis, Rukai; modern/ancient groups, Mbuti). We found that the values are mostly negative (Fig. S13), suggesting that most populations share excess ancestry with Rukai. In keeping with these f4 results, a best-fitting admixture graph modeled the Atayal as the closest to the Into-Taiwan group (sEA Li) and the Rukai as the closest to the Out-of-Taiwan group (ISEA Kakanaey), while the Amis are placed in-between (Fig. S14).

**Fig. 5.**
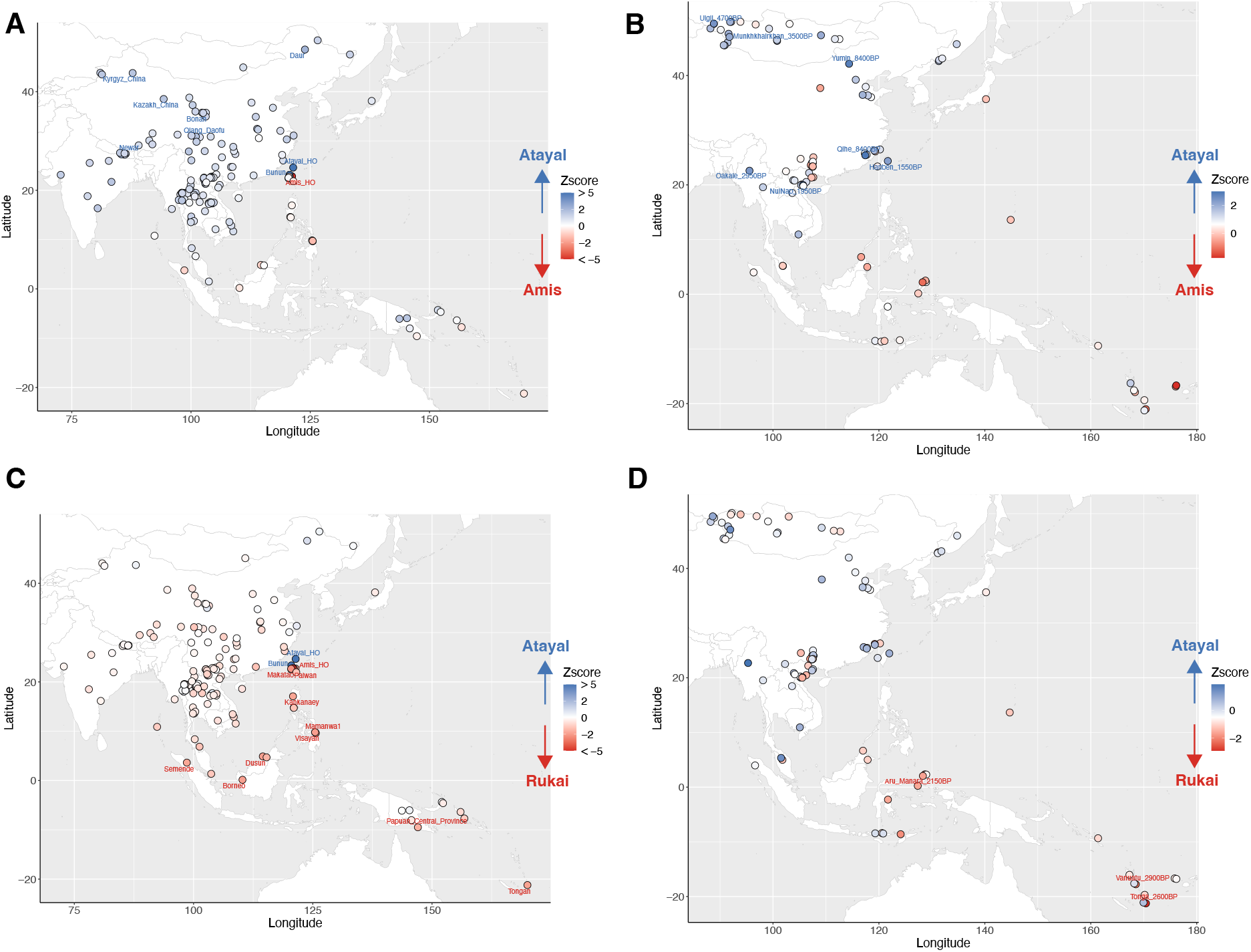
Differential allelic sharing to modern/ancient groups from East Asia and Oceania between Atayal and Rukai/Amis. Results of the form f4(Atayal, X; Y, Mbuti) where X is Amis (A and B) / Rukai (C and D) and Y are modern (A and C) / ancient (B and D) groups from East Asia and Oceania. The modern/ancient groups are plotted as dots on the map, colored in proportion to Z score. Positive values (in blue) indicate more sharing with Atayal while negative values (in red) indicate more sharing with Amis/Rukai. Significant values (absolute Z score value >= 2) are further labelled with population names. For the comparisons with ancient groups, additional tests using only transversions and French as outgroup are shown in Fig. S12, which reduce and the potential for false positives caused by DNA damage and/or attraction to deep outgroups but also the statistical power due to the decreased number of SNPs.

We further applied haplotype-based approaches, which enrich the signal of recent genetic sharing, to investigate the affinities between Taiwanese Austronesian and Out-of-Taiwan-related groups. Although results of frequency-based methods (i.e., f4 and qpgraph) indicate that the Rukai might be the closest to the Out-of-Taiwan groups, the fineSTRUCTURE (haplotype-based) results cluster Amis with Tao and Kankanaey, suggesting Amis might be the closest to the Out-of-Taiwan-related groups from a haplotype-based perspective. We therefore performed another ChromoPainter analysis using only the THI (Formosan branches) groups as the source to paint other Austronesians (Malayo-Polynesian branch) and indeed found that the Amis share most with other Austronesian groups, followed by the Rukai (Fig. 6). We also investigated the sharing of identity-by-descent (IBD) genomic segments in three main size ranges: 1 to 5 cM, 5 to 10 cM, and over 10 cM, roughly corresponding to ∼2.7 kya, ∼675 ya, and ∼225 ya, respectively (Al-Asadi, et al. 2019). Most of the signals are enriched in the size range of 1 to 5 cM (Fig. S15), while the THI groups show strong within group sharing and regional structure in the longer size ranges (Figs. S16 and S17), as found in the previous analyses of the genetic structure within Taiwan (Fig. 3; Fig. S5). Similarly, the Lowland Makatao also show an intermediate sharing profile between the Han and THI/TOI groups in the size range of 1 to 5 cM (Fig. S15). In this shortest size range, the THI/TOI groups only show notable sharing (the log value of average of the summed IBD length >= 1 cM) with Austronesian groups, except that they also share with Taiwanese Han (Hakka) and Papuan groups (Nasioi, Papuan Central Province, Papuan Gulf Province, Vella Lavella) with Austronesian admixture (Fig. S3). Finally, the IBD results indicate that the southern groups share more with Out-of-Taiwan groups than do the northern/central groups, e.g., the log value of the average summed IBD length for Rukai/Amis vs. Kankanaey is greater than 3 cM and for Rukai/Amis vs. Papuan Central Province is greater than 2 cM, while for Atayal/Bunun, these values are lower (Fig. S15).

**Fig. 6.**
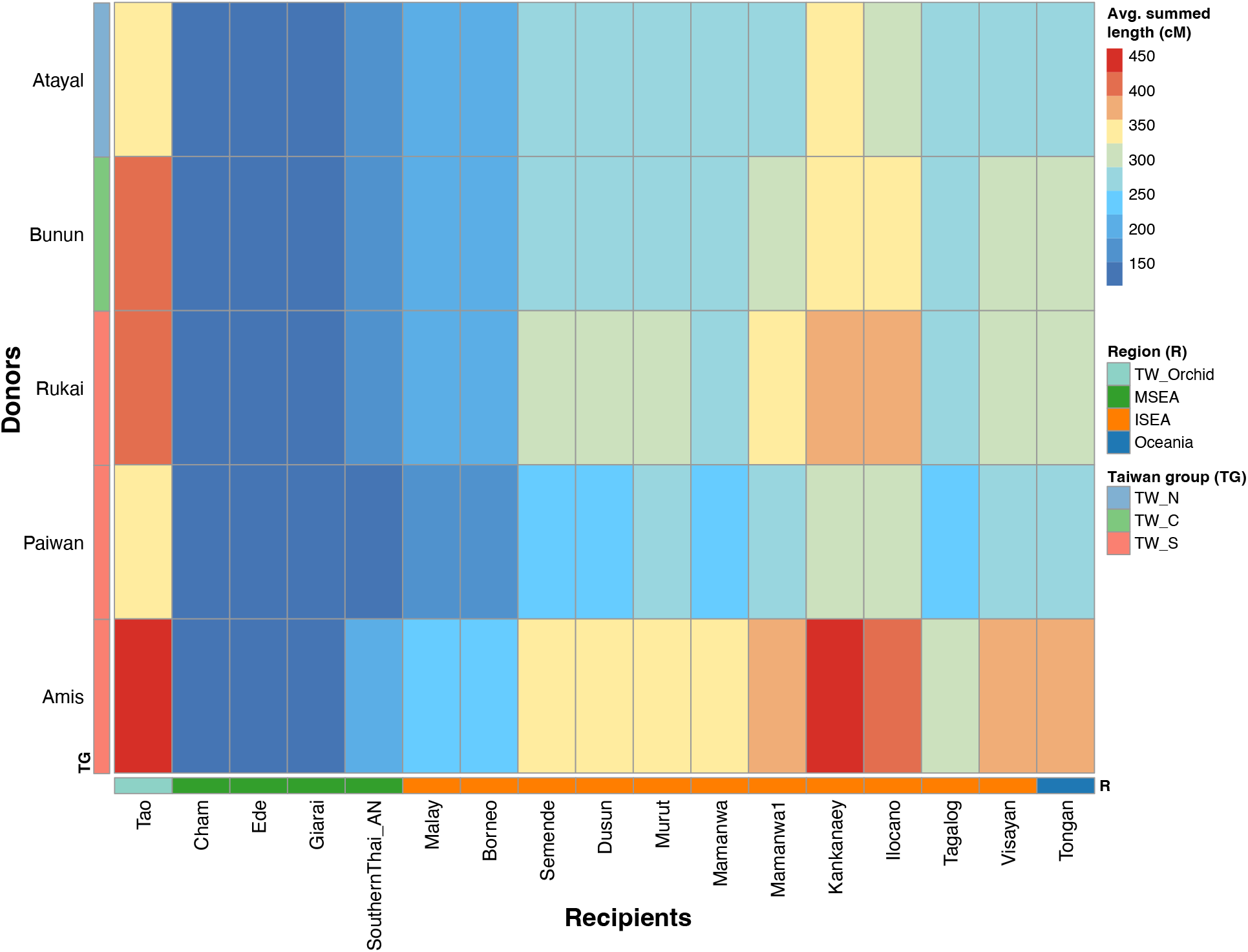
Chromosomal painting of the Malayo-Polynesian Austronesian recipients by the Formosan Austronesian donors. Results show the painting profile of recipients (in columns) by donors (in rows) colored in proportion to the average of the summed length (cM) of painted chromosomal regions, corresponding to the extent of haplotype sharing. The Taiwan group (TG) for the donors and the region (R) for the recipients are denoted by the colored keys. TW_N, TW_C, TW_S indicate northern, central, southern Taiwanese highland groups, respectively.

## Discussion

In this study, we generated new genome-wide data for 55 Taiwanese individuals (43 Austronesians) to characterize the genetic structure of Taiwan and address how this structure impacts questions regarding the Into- and Out-of-Taiwan migrations. Here, we highlight and discuss our most important findings concerning the genetic structure of THI/TOI groups, the genetic profile of lowland groups, and the Into-Taiwan and Out-of-Taiwan events.

### Genetic structure of Taiwan

The linguistic diversity of the Formosan branches of the Austronesian language family (Blust 1999; Li 2008) and the distinct cultures of the indigenous Taiwanese groups (CIP: https://www.cip.gov.tw) are suggestive of a population structure within Taiwan, particularly for the THI/TOI groups. Prior research into mitochondrial DNA (mtDNA) variation in Taiwanese groups revealed a cline of decreasing genetic diversity from north to south, with the groups further subdivided into northern, central, and southern clusters based on patterns of mtDNA sequence sharing. (Ko, et al. 2014). Furthermore, the central group is more closely related to the northern group than the southern groups are, and the southern groups can be further subdivided, as shown by our genome-wide data (Fig. 3B and 3C). In fact, the mtDNA study found that the northern group Atayal and the central group Bunun shared some haplotypes while the southern groups Rukai/Paiwan and Amis had distinct haplotype profiles (Ko, et al. 2014). In addition, our PCA results for the THI/TOI groups suggest a three-main-pole structure represented by Atayal, Rukai, and Amis (Fig. 3C), which also represents three distinct Formosan branches (Atayalic, Rukai, and East Formosan, respectively). The reason for the formation of such a structure is probably isolation between the groups, as shown by the high levels of IBD sharing within these groups (Fig. S5); the fact that Atayal exhibits the highest levels of within-group IBD sharing (Fig. S5) may explain why they are the most distinct of the THI/TOI groups in PCA and ADMIXTURE analyses (Figs. 2 and 3C). The Atayal have their own distinct haplotype network, according to previous Y-chromosomal DNA study (Trejaut, et al. 2014), suggesting they experienced founder/isolation events. The genetic structure observed within Taiwan could have formed (or strengthened) within the past 3 kya as the signal can be identified in all size ranges of IBD sharing between/within the THI/TOI group (Figs. S15-S17), which is in line with the ∼1-3 kya time estimate, based on mtDNA, for the formation of different groups (Ko, et al. 2014). However, not all THI/TOI groups are represented in our analyses; further studies that include these unsampled groups are needed to verify and extend our observations.

### Genetic profile of a lowland Taiwan group

There are around 12 identified lowland groups (belonging to the West Plains, Northwest Formosan, and East Formosan branches of the Austronesian language family) (Blust 1999), but only one is represented in our data. As the lowland groups are even less studied than the THI/TOI groups, our data provides the first genome-wide characterization of a lowland group. The lowland groups are known to have had extensive contact with the Han (Chiung 2001), and they share a similar demographic pattern of population size change with the Han that differs from the THI groups in the mtDNA study (Ko, et al. 2014) and larger population size and higher frequencies of sharing haplogroups in the Y-chromosomal DNA study (Trejaut, et al. 2014). Our data provide evidence of autosomal admixture between the THI and Han groups for the lowland group Makatao (Figs. 2 and 3A; Fig. S6), with the Rukai as the best proxy for the THI source (Fig. 3C; Fig. S6). Although the Makatao language belongs to the East Formosan branch and the Rukai language belongs to the Rukai branch, the two groups’ sampling locations of the two groups are in geographic proximity (Fig. 1A), which might explain for their genetic similarity and reflect their recent local contact through the sharing of IBD in the size range of 5 to 10 cM (Fig. S16). Evidence for ongoing gene-flow between the Han and Austronesian groups is suggested by the inferred admixture date of ∼50 years ago, which may be the most recent admixture date captured by our dating methods. Our sample size may not have been large enough to detect IBD in expanding populations (like Han), which may explain why we were unable to find evidence of IBD sharing between the Makatao and Han groups in the longer size range. Because we also find trace amounts of an Austronesian-related component in the Han groups, we speculate that this gene-flow is bi-directional (Fig. 2; Fig. S15). Since this signal is so weak, we cannot model this admixture in the Han groups using our admixture graph analyses; however, we do find that the Hakka cluster more closely with the THI groups than the Minnan do (Fig. S6). In addition, a recent study of Taiwanese Han from the Taiwan biobank found evidence of genetic sharing between the Han and the Austronesian groups (Lo, et al. 2021). Additional studies of Taiwan’s lowland groups are clearly needed.

### Implications for Into-Taiwan

The Into-Taiwan event describes the ancestors of Taiwanese Austronesians from wherever they originated. Linguistic studies indicate that “proto-Austronesian” likely formed in the southeast coast of mainland China (Bellwood 1984). Archeological studies of cultivated rice and millet show a link from southeastern China to Taiwan, and then to Southeast Asia (Deng, et al. 2022), which further supports a southeast Chinese origin of the ancestors of Austronesians under the “farming-language dispersal” model (Diamond and Bellwood 2003). Ancient DNA studies have confirmed a genetic link between ancient groups from the southeast Chinese coast and Taiwanese Austronesians (Yang, et al. 2020; Wang, Yeh, et al. 2021). Unfortunately, these studies only included Amis/Atayal, and the estimates of nEA vs. sEA ancestry were inconsistent (Yang, et al. 2020; Wang, Yeh, et al. 2021). Our results add context to the published ancient genomes from or nearby Taiwan by showing that the ∼1.5 kya Hanben individuals show additional affinities to the northern group Atayal while the ∼7.7 kya Liangdao and ∼4.5 kya Suogang individuals are slightly closer to the southern group Rukai (Fig. S7). The Atayal’s traditional territory overlaps with the Hanben archaeological site in Yilan County in northern Taiwan, suggesting a familial relationship between the two groups. In contrast, the Liangdao and Suogang archaeological sites, on islands off western Taiwan, are far from the active area of the Rukai; one potential explanation for their affinities to the Rukai might be that the Rukai are more closely related to the early Into-Taiwan groups that swiftly moved from the north to the south, as suggested by the mtDNA study (Ko, et al. 2014). In fact, linguistic studies indicate that the Rukai language is the earliest Formosan branches to split off (Starosta 1995; Ross 2009). We also revisited the modeling of nEA vs. sEA sources in THI/TOI groups and the surrounding ancient/modern groups, and confirmed that all of the groups tested can be modeled as having nEA (8.3 kya Boshan) and sEA (7.7 kya Liangdao) ancestries (Fig. 4; Figs. S9 and S10). This indicates that the “proto-Austronesian” and Austronesian-related genetic components are both a mixture of the nEA and sEA ancestries. Moreover, compared to the Into-Taiwan groups (e.g., 7.7 kya Liangdao and 4.3 kya Tanshishan), the 1.5 kya Hanben and present day THI/TOI groups show an increased amount of the nEA source, which suggests an additional influx of nEA ancestry into Taiwan after the Neolithic expansion, in line with ancient DNA studies showing post-Neolithic gene-flow between nEA and sEA (Yang, et al. 2020). Interestingly, the early Out-of-Taiwan groups (e.g., 2.2 kya Guam, 2.6 kya Tonga, and 2.9 kya Vanuatu) show more nEA ancestry than the early Into-Taiwan groups, but less than the THI/TOI groups and Kankanaey; the Atayal, Bunun, and Tao show the most nEA ancestry among these groups (Fig. 4; Fig. S9). This suggests that either more nEA gene-flow into Taiwan occurred prior to the Out-of-Taiwan expansion, or that there are un-sampled Into-Taiwan groups with more nEA ancestry than the early Out-of-Taiwan groups. Moreover, present day northern THI/TOI groups have had additional contact with nEA-related groups.

Previous research has indicated a close relationship between Austronesian and Tai-Kadai speakers (Thurgood 1994; Sagart 2008; Liu, et al. 2020; Wang, Yeh, et al. 2021). Linguistic studies raise two hypotheses: a shared ancestor for “proto-Austronesian” and “proto-Tai-Kadai”, or an Austronesian group from Taiwan returned to mainland China and became the ancestors of Tai-Kadai (Thurgood 1994; Sagart 2008). Our admixture graph result favors the former hypothesis, as the Tai-Kadai speaking Li, who has been shown to be an unadmixed proxy for Tai-Kadai ancestry (He, et al. 2020), shares the same source of nEA ancestry with the Austronesian groups rather than nesting within the Austronesian groups, and the Li even independently receive additional ancestry from the ancestral sEA branch (Fig. S10).

### Implications for Out-of-Taiwan

The Out-of-Taiwan event describes the migration of people from Taiwan to the ISEA and Oceania, coinciding with the spread of Austronesian languages and agriculture. While this migration and spread of culture have been confirmed by archeological, linguistic, and genetic evidence (Diamond and Bellwood 2003; Gray, et al. 2009; Ko, et al. 2014; Lipson, et al. 2014; Blust 2015), the mode/tempo of the migration remain debated (Bergstrom and Tyler-Smith 2018; Alam and Purugganan 2021) and how it relates to the structure within Taiwan is largely unexplored. In line with the mtDNA study (Ko, et al. 2014), our results support a closer relationship between the southern THI/TOI groups and the Out-of-Taiwan groups (Figs. 5 and 6; Figs S11 and S14). However, different results point to different southern groups as being the closest to the Out-of-Taiwan groups. The haplotype-based results favor the Amis being closer to the Out-of-Taiwan groups (Figs. 3B and 6), which is supported by a recent linguistic study suggesting that the East Formosan branch (which the Amis language belongs to) is the closest to the Out-of-Taiwan, Malayo-Polynesian branch (Chen, et al. 2022). However, the allele-based f4 comparisons show that the Rukai shares more ancestry with the Out-of-Taiwan groups than the Amis (Fig. 5), and this is also the conclusion of the allele-based admixture graph (Fig. S14). The haplotype-based analyses enrich the signal of recent contact while the allele-based analyses capture the average of overall sharing during population history, suggesting that, with the Out-of-Taiwan groups, the Amis received additional recent contact while the Rukai retained on average high genetic affinities throughout their history.

A further potential complication is that recent genetic studies have estimated an older divergence time between THI groups and Filipino groups than the linguistic Out-of-Taiwan time estimate, and suggested that the spread of Austronesian languages might not be associated with the migrations of people (Choin, et al. 2021; Larena, et al. 2021). Given that both southern groups Amis and Paiwan (close to Rukai) were used in their models, this older estimate is not due to using a more distant THI group to Out-of-Taiwan groups; however, even a small amount of additional interactions between the THI and nEA-related groups compared to the Filipino groups could inflate the estimate, as the inclusion of an nEA source in the model expands the confidence interval to overlap with the linguistic Out-of-Taiwan time (Choin, et al. 2021); in our study we do see nEA sources modeled for all the THI groups (Fig. 4; Fig. S9).

To sum up, one likely and concise synthesis of our results would be that 1) the early Into-Taiwan groups rapidly moved to the south and became Out-of-Taiwan groups, which is supported by our DyStruct results indicating that the early sEA groups share similar profiles with the early Oceania (Lapita-related) groups (Fig. 2) and by the previous mtDNA study (Ko, et al. 2014). 2) Given that there was little divergence between the Into-Taiwan and Out-of-Taiwan groups, our results of the Rukai retaining excess allelic sharing with both Into-Taiwan and Out-of-Taiwan groups (Fig. 4; Figs. S7 and S13) might reflect that their ancestor underwent less genetic drift than the ancestors of other present-day Taiwan groups; alternatively, the ancestors of Rukai might actually be the closest to the Into-Taiwan (which soon became Out-of-Taiwan) groups, which is in line with the linguistic evidence that they are the earliest diverged groups (Starosta 1995; Ross 2005b). 3) Populations further diverged within Taiwan after these early events, as suggested by IBD results (Fig. S15-S17) and the mtDNA study (Ko, et al. 2014), with the northern group Atayal experiencing the strongest bottleneck/isolation (Figs 2 and 3C; Fig. S5). 4) The closer haplotype sharing (and perhaps the linguistic relationship (Chen, et al. 2022)) between the Amis and Out-of-Taiwan groups might result from recent back-migration contact (Figs. 3B and 6); the East Formosan Amis is close to the Malayo-Polynesian Tao geographically and genetically, based on our results (Figs. 3 and 6) as well as previous findings (Tatte, et al. 2021), and it has been suggested that the Tao might be derived from a back-migration from ISEA (Ross 2005a). 5) Finally, during even more recent times, the lowland groups interacted extensively with Han groups, resulting in an admixed genetic profile (Figs. 2 and 3A; Fig. S6). In any case, more whole genome sequencing of THI/TOI groups would improve the power of more sophisticated analyses to test these models, along with more ancient genomes from Into-Taiwan, Taiwan, and Out-of-Taiwan groups.

## Materials and Methods

### Sample and data information

Sampling of Taiwanese individuals was done in Ko et al (Ko, et al. 2014); we selected a total of 43 Austronesian (37 highlanders from Atayal, Bunun, Rukai, Paiwan, Ami; 1 Tao; and 5 Makatao lowlanders) and 12 Taiwanese Han (Hakka; Minnan) individuals and generated their genome-wide data on the Affymetrix Human Origins array. The ethics committees of the China Medical University, the Taiwan National Health Research Institutes, and the University of Leipzig Medical Faculty have approved this study. Informed consent was obtained from all participants. We first merged the new data with published data on the same array from comparative modern populations (Patterson, et al. 2012; Lazaridis, et al. 2014; Qin and Stoneking 2015; Skoglund, et al. 2016; Lipson, Cheronet, et al. 2018; Liu, et al. 2020; Kutanan, et al. 2021; Wang, Yeh, et al. 2021) and we kept only autosomal markers for our analyses. For quality control of the merged data of modern individuals, we excluded sites with more than 5% missing data in the entire dataset and sites with more than 50% missing data and/or Hardy-Weinberg equilibrium p values below 0.00005 within a group (except for groups with only one individual). We removed individuals with more than 5% missing data and we excluded one individual from 1^st^ degree kinship pairs. Data missingness and Hardy-Weinberg equilibrium were calculated using PLINK v1.9 (Purcell, et al. 2007) while the individuals to be removed to avoid 1st degree relatedness (kinship coefficient >= 0.177) were inferred using KING (Manichaikul, et al. 2010), as implemented in PLINK v2 (Chang, et al. 2015). This gives a filtered dataset of modern individuals consisting of 540,697 SNPs and 1,664 individuals. Next, we further merged the filtered dataset with comparative ancient genomes (Skoglund, et al. 2016; Yang, et al. 2017; Lipson, Cheronet, et al. 2018; Lipson, Skoglund, et al. 2018; McColl, et al. 2018; Posth, et al. 2018; Lipson, et al. 2020; Yang, et al. 2020; Pugach, et al. 2021; Wang, Yeh, et al. 2021; Wang, Wang, et al. 2021; Oliveira, et al. 2022). Six positions with more than two variants or that were inconsistent between the array and sequencing data were excluded. Ancient individuals with <15,000 informative sites were also removed. This gives a filtered dataset of modern and ancient individuals consisting of 540,691 SNPs and 1,958 individuals. For Principal Component Analysis (PCA), ADMIXTURE, and DyStruct analyses (described in below sections), variants were pruned beforehand for linkage disequilibrium using PLINK v1.9, excluding one variant from pairs with r2 > 0.4 within windows of 200 variants and a step size of 25 variants. Metadata are in Table S1.

### PCA analyses

PCA was done with smartpca v16000 (Patterson, et al. 2006). Projection of groups on PCA was done with the “lsqproject” and “shrinkmode” options.

### ADMIXTURE and DyStruct analyses

The clustering algorithm was first run by ADMIXTURE v1.3.0 (Alexander, et al. 2009). From K = 2 to K = 15, we performed 100 replicates for each K with random seeds. ADMIXTURE outliers with >0.01 of the turquoise (Han-related) component were excluded for the Amis (TA199, TA81), Bunun (bun55, bun32), Atayal_HO (NA13600), and Kankanaey (Igorot11) groups in the F-statistics based analyses (e.g., f3, f4, qpAdm, admixture graph, described in the below sections). We also performed DyStruct v1.1.0 (Joseph and Pe’er 2019) incorporating time-series data from ancient genomes for the model. We transformed the age of ancient genomes into generations using a generation time of 30 years (Fenner 2005) and binned every 50 generations into a time point using the script bin_sample_times.py. From K = 2 to K = 15, we performed 25 replicates for each K with random seeds. Then, we used the script plot_Q.py to plot the highest likelihood results for the best K of ADMIXTURE (K=9, with the lowest cross-validation error) and DyStruct (K=8, with the highest likelihood). Scripts bin_sample_times.py and plot_Q.py are provided by the authors of DyStruct (available here: https://github.com/tyjo/dystruct/tree/master/supp/scripts).

### F3 and F4 statistics and qpAdm analyses

We used admixr v0.9.1 (Petr, et al. 2019) to compute f3- and f4-statistics from ADMIXTOOLS v7.0.2 (Patterson, et al. 2012), with significance assessed through block jackknife resampling across the genome. We also used admixr to compute qpAdm from ADMIXTOOLS with option inbreed = “YES” (required if using a single individual as a source or outgroup). We merged additional outgroups (Fu, et al. 2014; Seguin-Orlando, et al. 2014; Gallego Llorente, et al. 2015; Fu, et al. 2016; Lazaridis, et al. 2016; Damgaard, et al. 2018; Moreno-Mayar, et al. 2018; Narasimhan, et al. 2019; Sikora, et al. 2019) to follow Yang et al (Yang, et al. 2020), with metadata available in Table S1. The target groups are all Taiwanese groups and selected Tai-Kadai (Dong_Guizhou, Dong_Hunan, Gelao, Li, Maonan, Mulam, Zhuang), Han (Han_Fujian, Han_Guangdong, Han_Shandong, Han_Shanghai, Han_Henan, Han_Hubei), ancient nEA (Xiaojingshan_7800BP, Xiaogao_8700BP, Wuzhuangguoliang_5000BP) and ancient sEA (Suogang_4550BP, Xitoucun_4500BP, Tanshishan_4350BP, Hanben_1550BP, Shenxian_1300BP, BaBanQinCen_1500BP) groups; the source groups are nEA Boshan_8200BP and sEA Liangdao_7750BP; the outgroups are Mota_4470BP, Ust_Ishim_44350BP, Kostenki_38050BP, Iran_10000BP, Yana_31850BP, Karelia_8450BP, Okunevo_4300BP, Indus_Periphery_4500BP, New_Guinea_Highlander, Onge, Upward_Sun_River_11400BP, Tianyuan_40000BP, Longlin_10550BP, Kolyma_9750BP, Jomon_2800BP, Qihe_11550BP, Bianbian_9500BP, Mbuti, French, Australian, Tu, She, Ami, Kinh, and Baining. In contrast to the outgroups used in Yang et al, we substituted the 7 kya Pha Faen genome (McColl, et al. 2018) (which overlaps with Onge in terms of Hoabinhian-related ancestry) with the 10 kya Longlin genome (Wang, Wang, et al. 2021) to provide more distinct sEA outgroups. We excluded results with negative values inferred for either one of the sources, p-values > 0.05, or the absolute values of the standard error > 0.25.

### Data phasing

Phasing was done by SHAPEIT version 4.1.3 (Delaneau, et al. 2019), with East Asian (without the Kinh Vietnamese merged in our data set) and South Asian populations as a reference panel, and the recombination map from the 1000 Genomes Phase3 (Genomes Project, et al. 2015). To prepare the reference panel, we extracted the East and South Asian individuals as well as the overlapping sites with our data for each chromosome from the 1000 Genomes Phase3 data using bcftools version 1.4 (http://samtools.github.io/bcftools/; last accessed 10 July, 2020). The phasing accuracy of SHAPEIT4 can be enhanced by increasing the number of conditioning neighbors in the Positional Burrows–Wheeler Transform (PBWT) on which haplotype estimation is based (Delaneau, et al. 2019). We ran phasing with the options –pbwt-depth 8 for 8 conditioning neighbors and left other parameters as default.

### ChromoPainter, fineSTRUCTURE and GLOBETROTTER analyses

ChromoPainter v2 (Lawson, et al. 2012) was run on the phased dataset focusing only on Austronesian, Tai-Kadai, and Sino-Tibetan speaking groups, with sample sizes for each population group randomly down-sampled to 7 (all individuals were used for groups with sample sizes below 7). We began with 10 iterations of the EM (expectation maximization) process to estimate the switch rate and global mutation probability using chromosomes 1, 5, 10, 15, and 20. With the estimated switch and global mutation rates, we ran the chromosomal painting process for all chromosomes, which then gave the output for downstream analyses. We first attempted to paint the chromosomes of each individual, using all of the samples in the dataset as both donors and recipients, via the -a argument. The EM estimation of switch rate and global mutation probability were ∼409.33 and ∼0.00048, respectively, which were then used as the starting values for these parameters for all donors in the painting process. To specifically study the contribution of THI groups (Formosan branches) as sources for other Austronesians (Malayo-Polynesian branch), we also performed another run using all the samples except for non-THI Austronesians as both donors and recipients; non-THI Austronesians were used only as recipients. The EM estimation of switch rate and global mutation probability for this analysis were ∼ 430.08 and ∼ 0.00064, respectively.

fineSTRUCTURE v4.0.1 (Lawson, et al. 2012) was run on the first ChromoPainter output using all of the samples in the dataset as both donors and recipients. First, 1 million burn in steps were used (-x) and 1 million further iterations were sampled (-y) keeping every 10000th sample (-z). To infer a tree (-m T), we ran an additional 100,000 burn in steps (-x) and used the maximum number of 3000 tree comparisons for splitting/merging (-t). We processed and plotted the results in R with the assistance of the fineSTRUCTURE R Tools (https://people.maths.bris.ac.uk/~madjl/finestructure/finestructureR.html).

To investigate the admixture of Austronesian and Han ancestries in the Lowland group Makatao, GLOBETROTTER (Hellenthal, et al. 2014) was run on the second ChromoPainter output using THI/TOI Austronesian, Tai-Kadai, and Sino-Tibetan groups as surrogates. We first tested the certainty and potential waves of admixture events, and then estimated the major and minor sources as well as the dates of admixture. A single pulse of admixture was inferred for Makatao. The standard error was computed from the distribution of admixture dates accessed through 100 bootstraps. A generation time of 30 years was used for transforming generations to years (Fenner 2005).

### Identity by descent (IBD) analyses

We identified shared IBD blocks between each pair of individuals and homozygous-by-descent (HBD) blocks within each individual using RefinedIBD (Browning and Browning 2013). Both identified IBD and HBD blocks are considered as IBD blocks in our analyses, which is analogous to pairwise shared coalescence (PSC) segments in a previous study (Al-Asadi, et al. 2019). The IBD blocks within a 0.6 cM gap were merged using the program merge-ibd-segments from the RefinedIBD website (https://faculty.washington.edu/browning/refined-ibd.html), allowing only 1 inconsistent genotype between the gap and block regions. We used IBD blocks at least 2 cM in length shared by individuals within a population to investigate the demography of each population group. Then, we used IBD blocks in 1-5 cM, 5-10 cM, and over 10 cM ranges to investigate the sharing between individuals from different populations in different time periods (Al-Asadi, et al. 2019). For visualization of the sharing between populations, the pairs with an average of at least 0.5 shared IBD blocks (i.e., on average half of the pairs share IBD blocks) were kept to reduce noise and false positives. The average of the summed IBD length/number of blocks between two groups was calculated by dividing the summed IBD length/number of blocks between all pairs of individuals from the two groups by the product of the number of individuals from one group and the number of individuals from the other group.

### Admixture graph analyses

We selected groups to test our questions concerning the admixture profile of the Lowland group Makatao (Fig. S6), and the Into-Taiwan (Fig. S10) and Out-of-Taiwan (Fig. S14) events. For ancient groups, we removed individuals with more than 30% missing data except for individual TON002 from ∼2.6 kya Tonga, who is the highest quality ancient Lapita individual and is needed for the Out-of-Taiwan model. We used ADMIXTOOLS 2 (Maier, et al. 2022) to compute pairwise f2 statistics between the groups using extract_f2() with maxmiss=1 (no SNPs excluded) and then extracted the allele frequency products from the computed f2 blocks using f2_from_precomp() with afprod=TRUE (which will result in more precise f4-statistics when there are large amounts of missing data, as described on the tool website: https://uqrmaie1.github.io/admixtools/). Finally, for each of the scenarios, we searched for the best-fitting admixture graph by running ten independent runs of find_graphs() and selected the one with the lowest score (computed through the residuals between the expected and observed f-statistics given the data). We further confirmed a fitting graph by testing the graph with the lowest score using qpgraph() to check if the absolute value of the worst-fitting Z score is below 3. We started with no migrations (numadmix=0) and then added migrations till we found a fitting graph, which we denoted as the best-fitting graph for that scenario. We also added different admixture constraints for searching the graphs depending on our questions; for the admixture profile of Makatao, at least 1 admixture for Makatao, and no admixtures for Atayal, Rukai, and Amis; for Into-Taiwan events, no admixtures for ∼8.2 kya Boshan and ∼7.7 kya Liangdao; for Out-of-Taiwan events, no admixtures for New Guinea Highlander, Atayal, and Rukai.

## Data Availability

The genome-wide SNP array data generated in this study are available from the European Genome-Phenome Archive (EGA; https://ega-archive.org), under accession code EGASXXXXXXXXXXX.

## Acknowledgements

We thank all sample donors for making this work possible. We dedicate this paper to Prof. Ying-Chin Ko, a pioneer in the public health research of Taiwan aborigines and a tireless worker on behalf of Taiwan’s aboriginal communities. We thank Benjamin Peter, Stephan Schiffels, Sandra Oliveira, Irina Pugach, Hsiao-Chun Hung, Stacy Fang-Ching Teng and Etienne Patin for helpful discussion concerning the interpretation of results. This study was supported by the Max Planck Society.

