## Supplementary Materials for "The genomic diversity of Taiwanese Austronesian groups: implications for the ‘Into and Out of Taiwan’ models"

**This file includes:**

Figures S1 to S17
Table S1

### Supplementary Figures

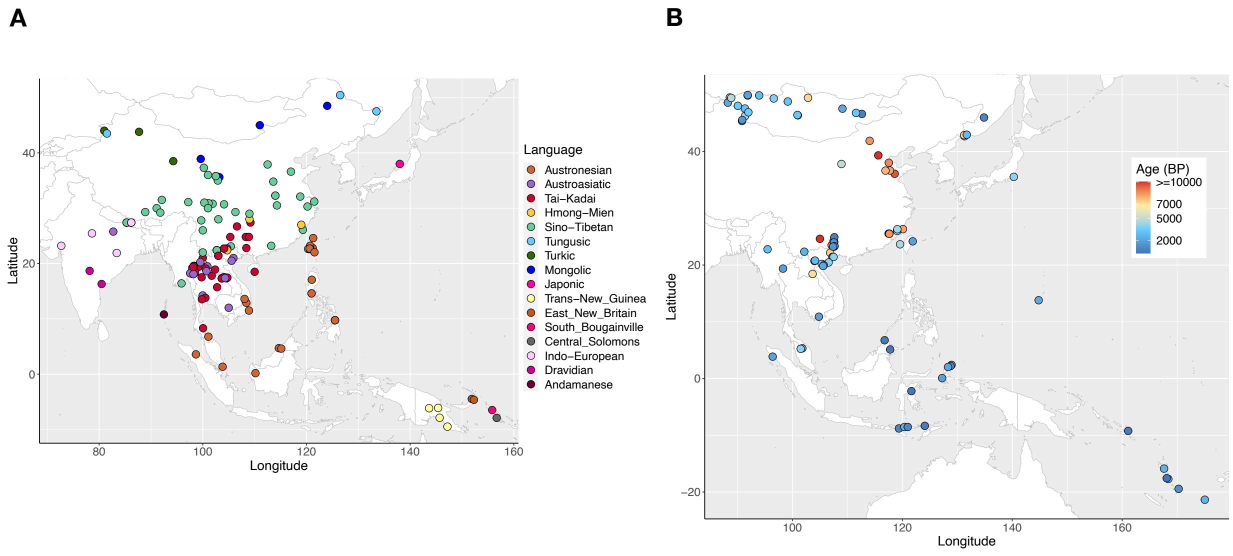


**Fig. S1. Map of comparative data.** Maps of comparative (A) modern and (B) ancient published data, colored by languages and sample ages, respectively. More details are in Table S1.

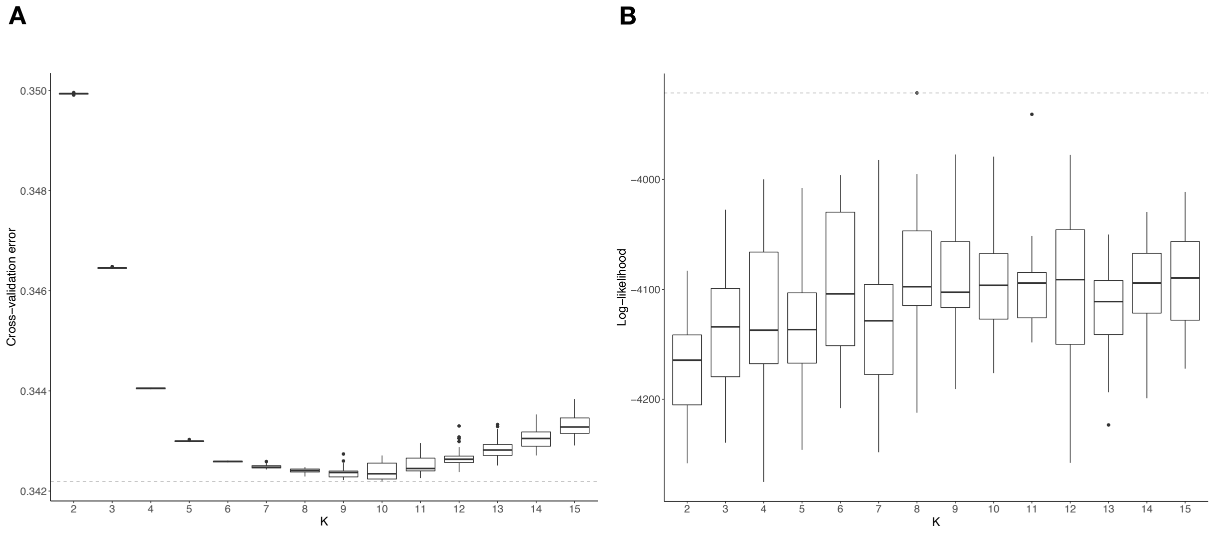


**Fig. S2. Identifying the best-fitting K for ADMIXTURE and DyStruct analyses.** (A) Cross-validation error plot for ADMIXTURE from K = 2 to K = 15. The best-fitting K (lowest cross-validation error) is K = 9. (B) Likelihood plot for DyStruct from K =2 to K = 15, indicating that the best-fitting K (highest likelihood) is K = 8. We ran 100 independent runs for ADMIXTURE (20 for DyStruct) for each K, and we show in Fig. 2 the runs with the highest likelihood for the best-fitting values of K.

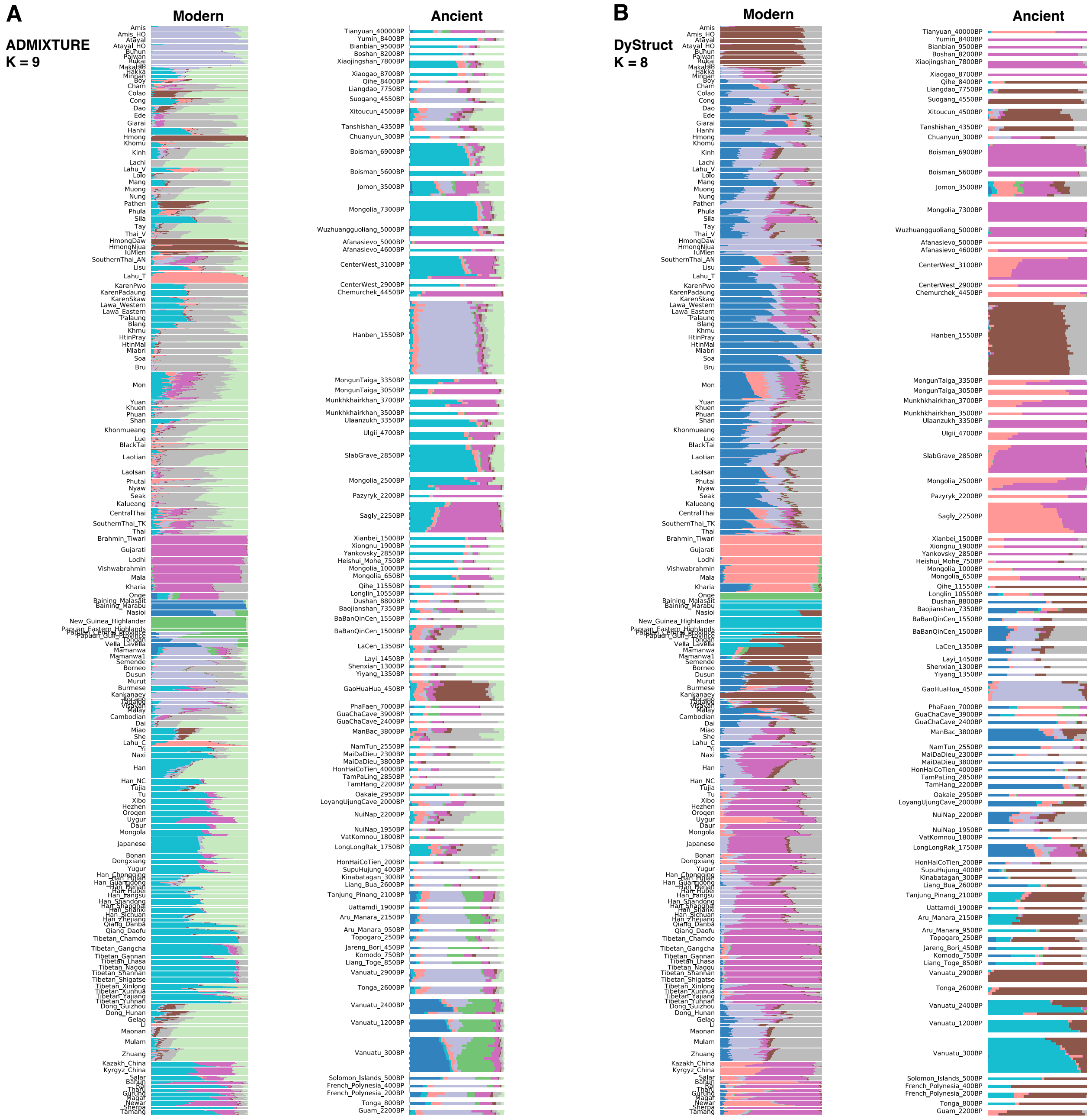

**Fig. S3. Full results of the best-fitting K for the ADMIXTURE and DyStruct analyses.** The highest likelihood run of (A) ADMIXTURE for K = 9 and (B) DyStruct for K = 8, with modern groups listed on the left and ancient samples on the right.

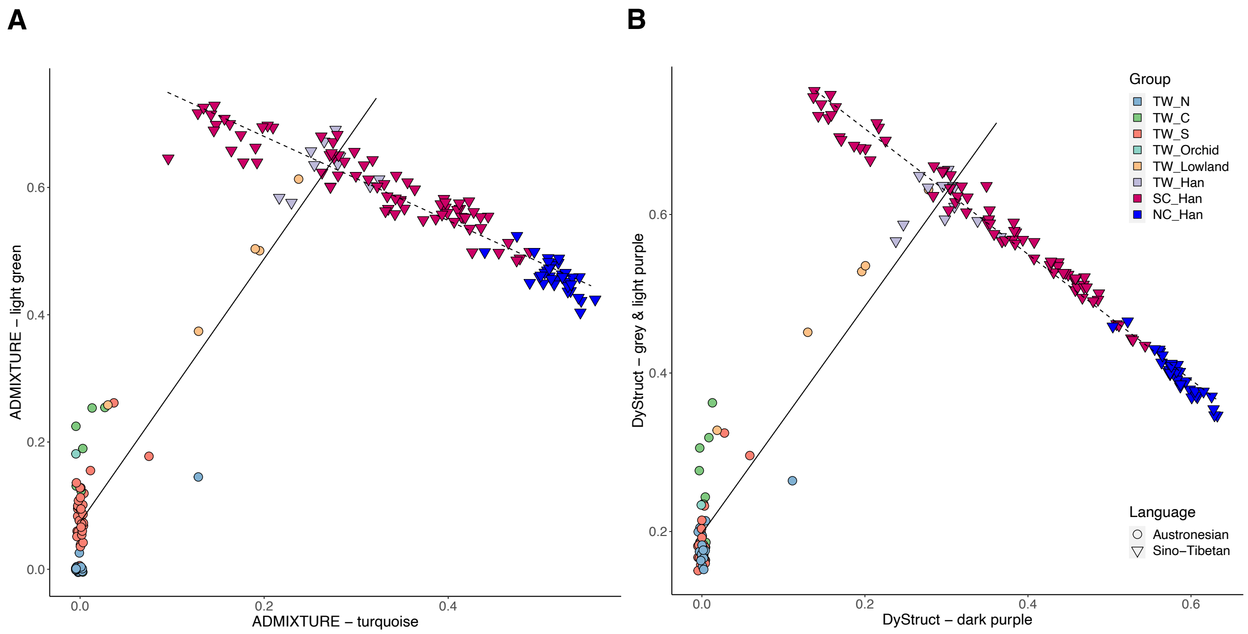


**Fig. S4. ADMIXTURE and DyStruct Han-related components in Taiwanese groups.** We plotted the components that show in some Taiwanese groups and largely shared with Han groups from China from the best-fitting K of (A) ADMIXTURE and (B) DyStruct results. Individuals were plotted as dots/triangles (according to language) and colored by groups. TW denotes Taiwan, and N, C, S denote northern, central, southern highland groups, respectively. SC_Han/NC_Han denote the southern/northern Han groups from China. The light green ADMIXTURE (grey and light purple DyStruct) component is negatively correlated with the turquoise ADMIXTURE (dark purple DyStruct) component for the Chinese Han groups ($r^2$=0.921, p=0 for ADMIXTURE; $r^2$=0.983, p=0 for DyStruct), but they are positively correlated in Taiwanese groups ($r^2$=0.913, p=0 for ADMIXTURE; $r^2$=0.926, p=0 for DyStruct).

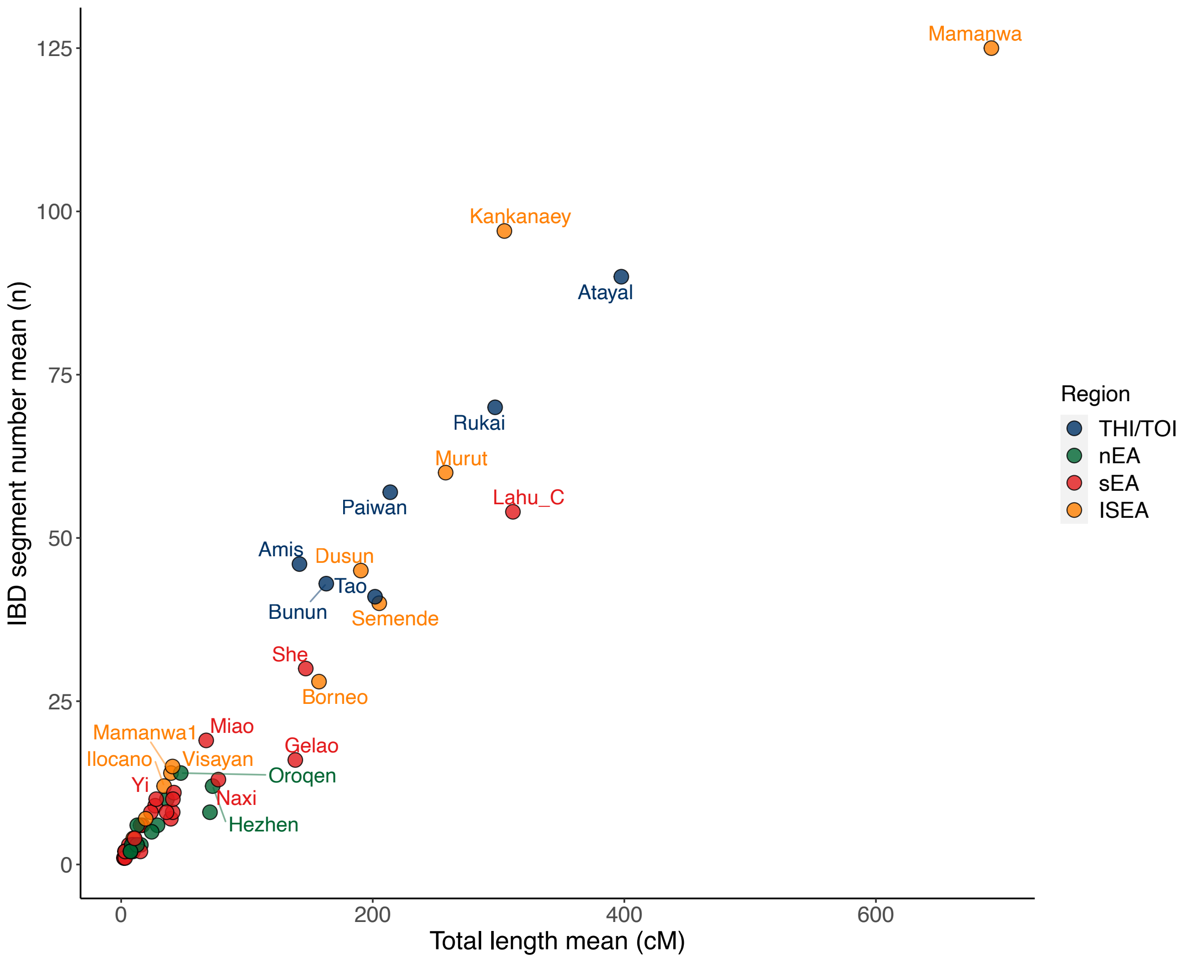


**Fig. S5. Within group IBD sharing.** We plotted on the x-axis the mean of the summed length of IBD segments, and on the y-axis the mean number of IBD segments for groups from nEA, sEA, and ISEA (colored accordingly). THI/TOI groups are highlighted in another color.

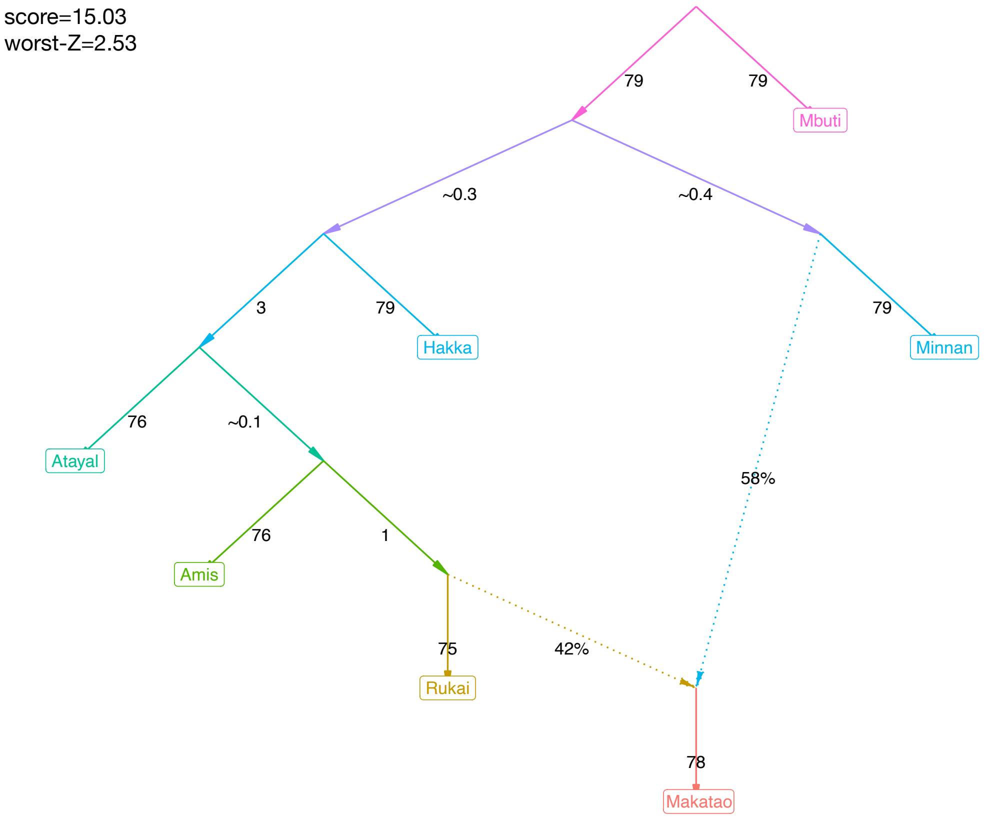


**Fig. S6. Best-fitting admixture graph for investigating admixture in the lowland group Makatao.** The numbers on the solid branch lines are in genetic drift units while those on the dashed lines indicate ancestry proportions. The ADMIXTOOLS 2 graph score and the worst-fitting Z score are shown on the top-left.

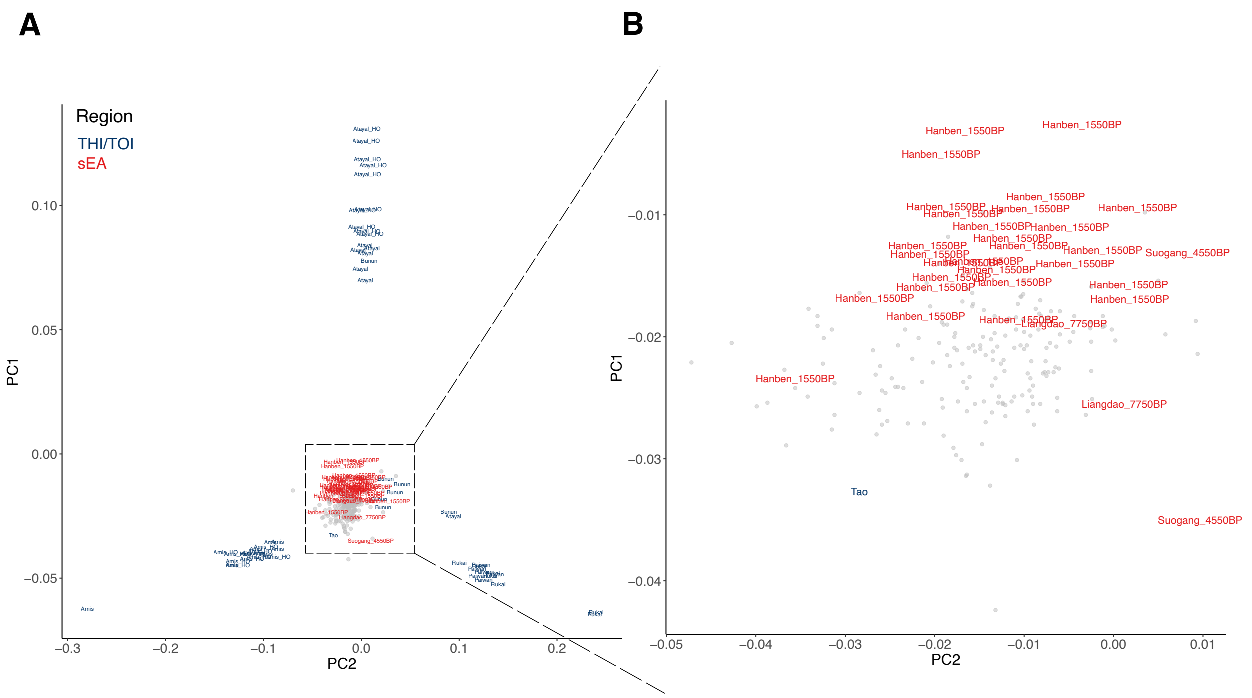


**Fig. S7. THI/TOI PCA with a focus on the projected ancient Taiwan-related individuals.** (A) the full results and (B) zoom-in of (A) onthe projected ancient sEA individuals. Our focusing groups are colored by regions while the other projected ancient individuals are colored in grey to provide an overall background variation.

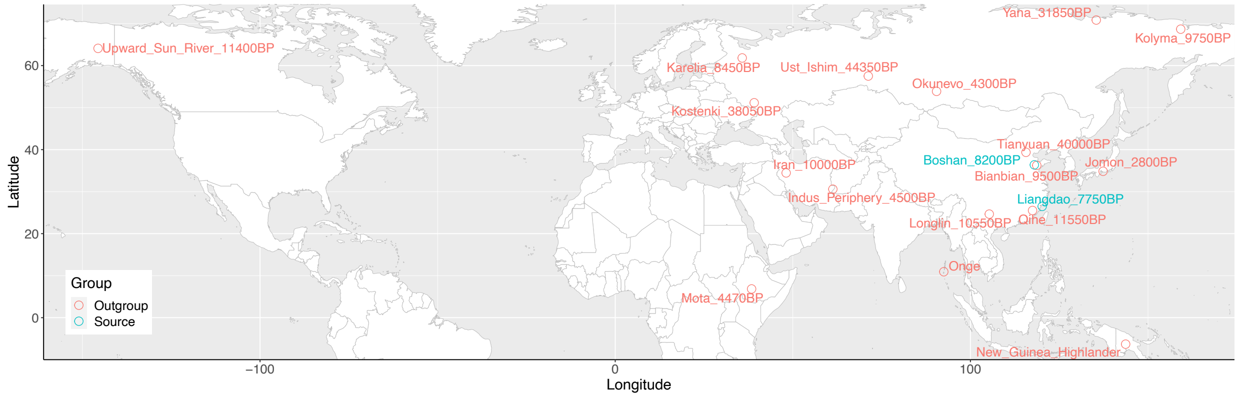


**Fig. S8. Map of source groups and outgroups for qpAdm.**

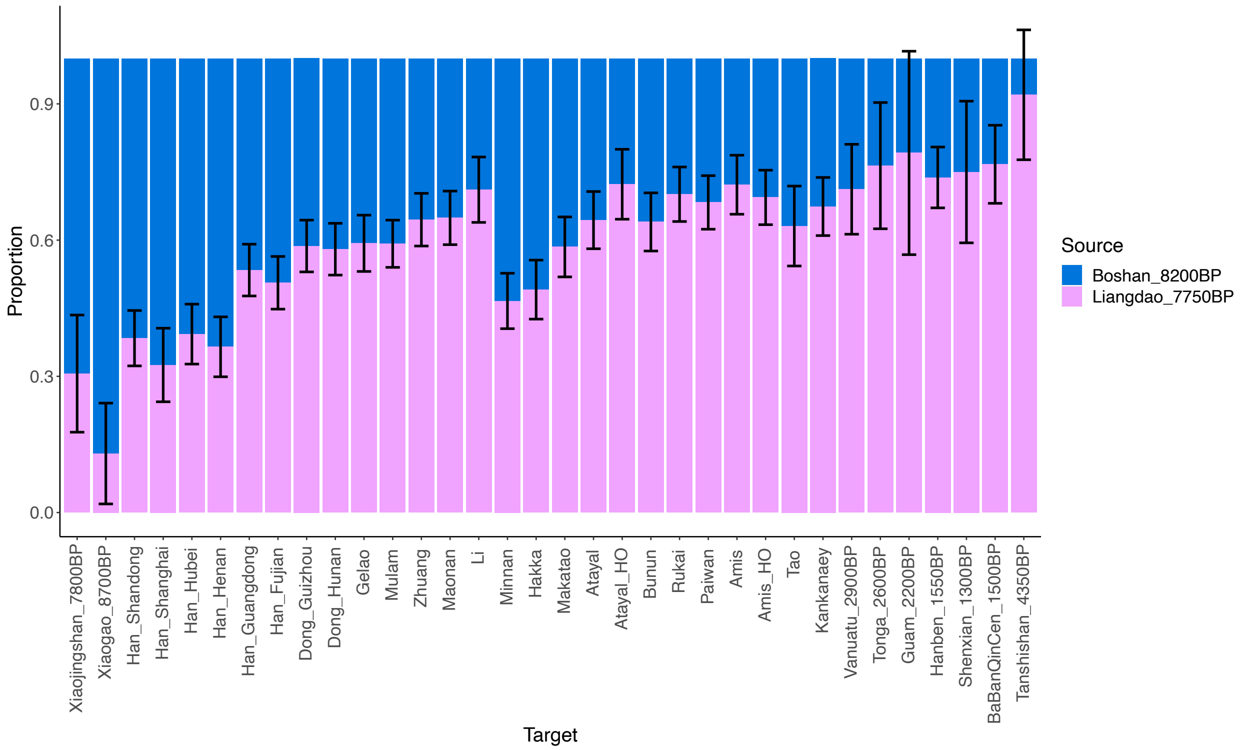


**Fig. S9. Bar plot visualization of the qpAdm results.** The error bars indicate the mean +/- one standard error.

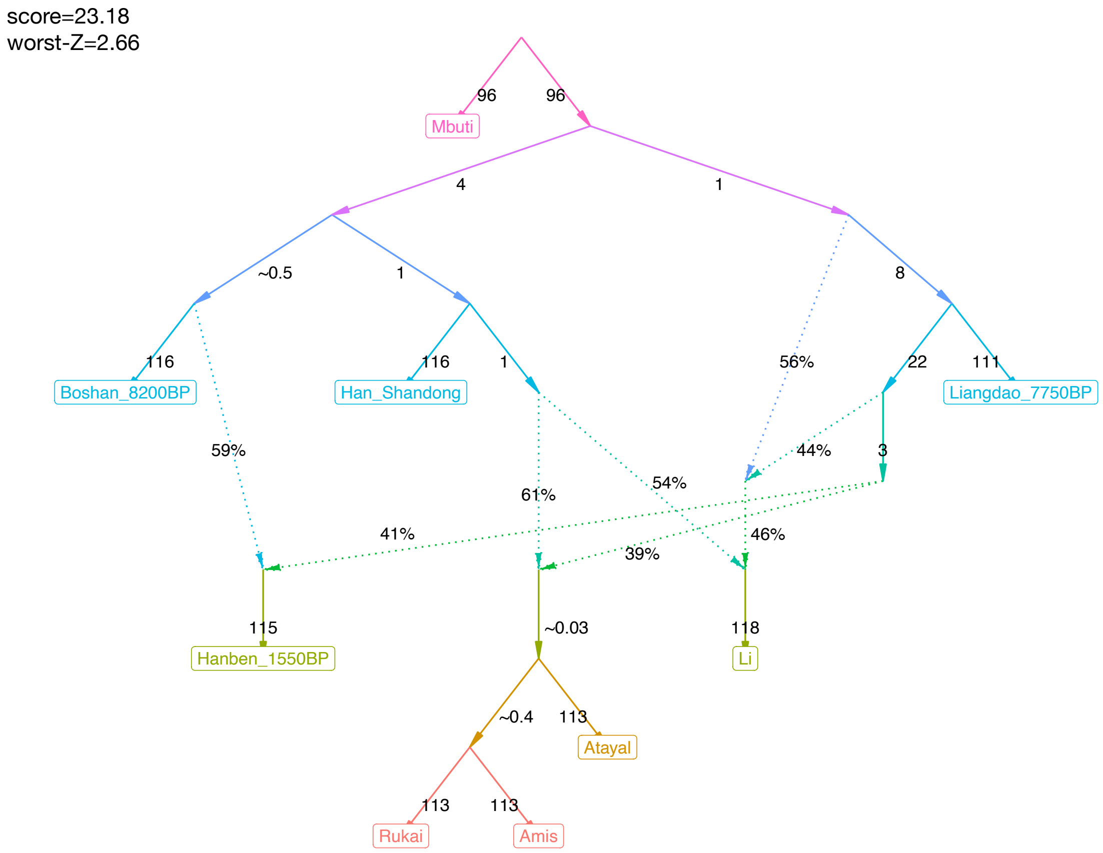


**Fig. S10. Best-fitting admixture graph for Into-Taiwan events.** The numbers on the solid lines denote are in genetic drift units while those on the dashed lines indicate ancestry proportions. The ADMIXTOOLS 2 graph score and the worst-fitting Z score are shown on the top-left.

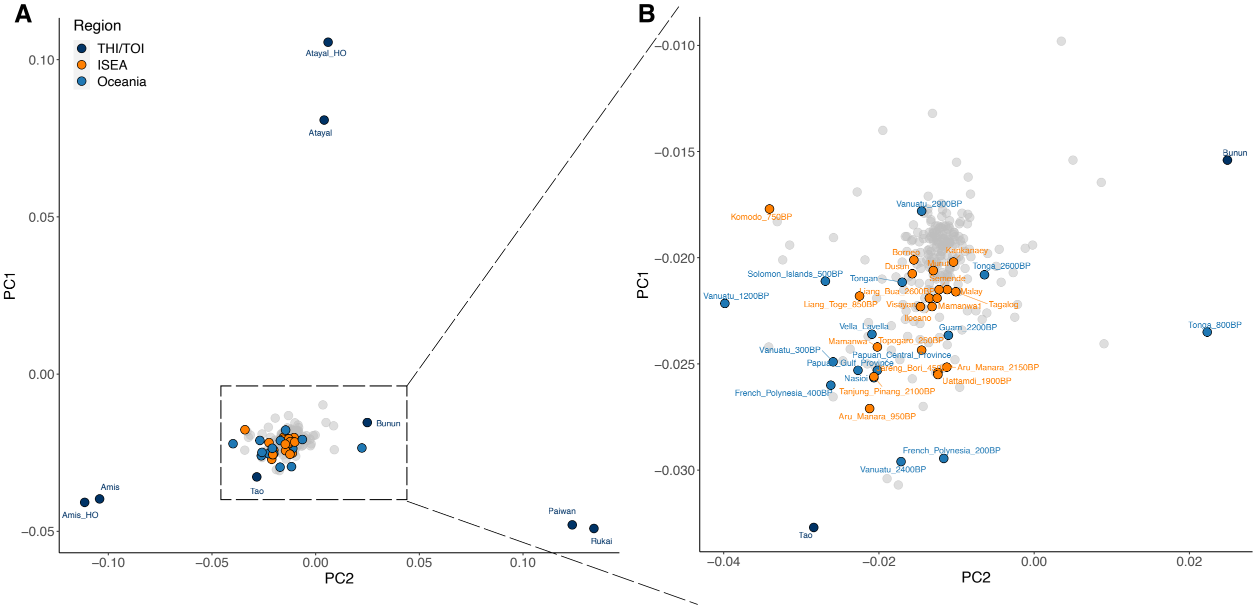


**Fig. S11. THI/TOI PCA with a focus on the projected Austronesian-related ISEA/Oceanian groups.** The PCA was computed using only THI/TOI groups. (A) the full results and (B) zoom-in of (A) show the projected modern and ancient groups with a focus on the ISEA/Oceanian groups whose individuals have at least 10% Austronesian-related (purple) component in ADMIXTURE K=9 (Fig. 2; Fig. S3). Dots represent the median position of individuals from a group. Our focusing groups are colored by regions while the other projected groups are colored in grey to provide an overall background variation.

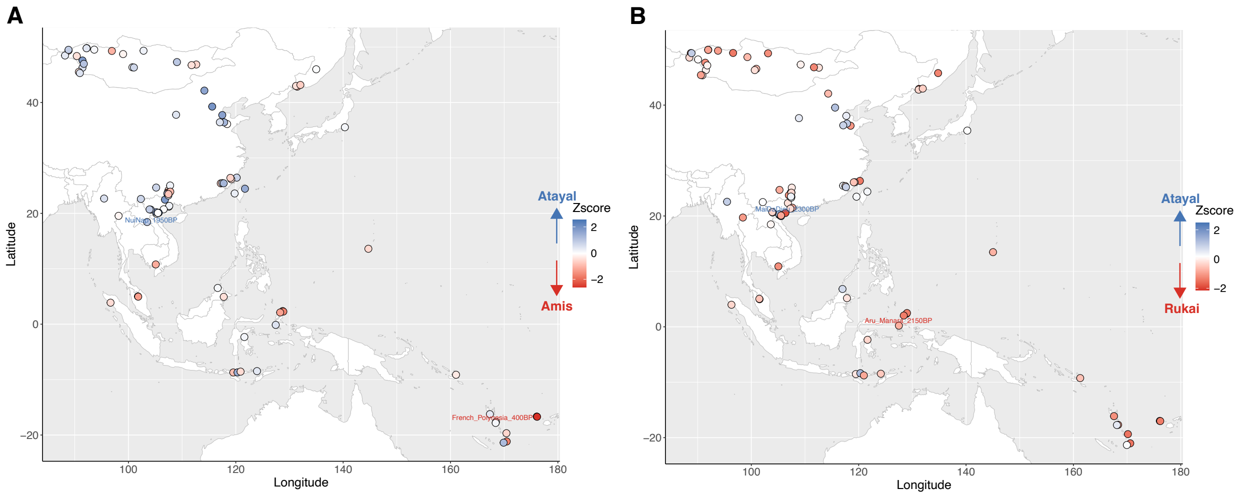


**Fig. S12. Differential allelic sharing to ancient groups from East Asia and Oceania between Atayal and Rukai/Amis using only transversions and French as outgroup.** Results of the form f4(Atayal, X; ancient groups from East Asia and Oceania, French) where X is Amis (A ) / Rukai (B). The ancient groups are plotted as dots on the map, colored in proportion to Z score. Positive values (in blue) indicate more sharing with Atayal while negative values (in red) indicate more sharing with Amis/Rukai. Significant values (absolute Z score value >= 2) are further labelled with the population name.

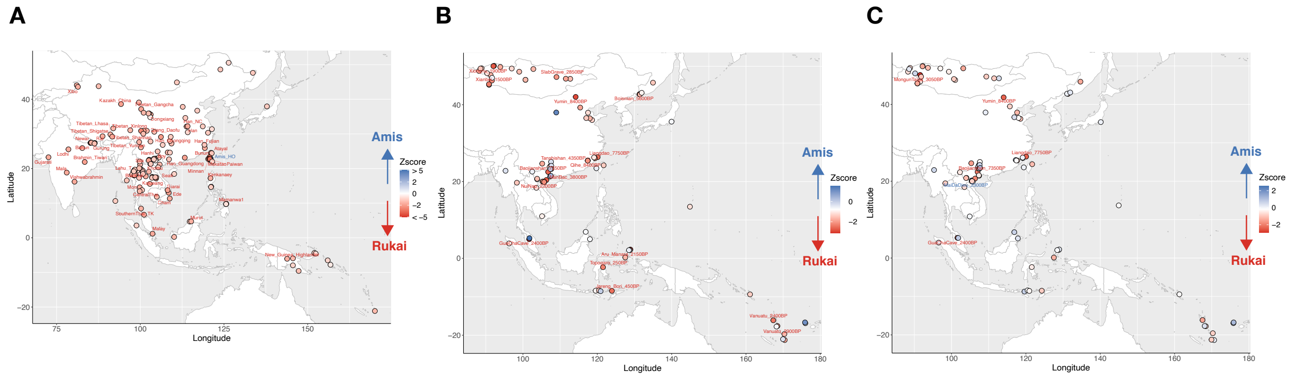


**Fig. S13. Differential allelic sharing to modern/ancient groups from East Asia and Oceania between Atayal and Rukai/Amis.** Results of the form f4(Amis, Rukai; Y, Mbuti) where Y are modern (A) / ancient (B) groups from East Asia and Oceania. Additional tests on (C) are results of the form f4(Amis, Rukai; ancient groups, French) using only transversions, which reduces the false positives caused by DNA damage and/or attraction to deep outgroups but also the statistical power due to the decreased number of SNPs. The modern/ancient groups are plotted as dots on the map, colored in proportion to Z score. Positive values (in blue) indicate more sharing with Amis while negative values (in red) indicate more sharing with Rukai. Significant values (absolute Z score value >= 2) are labelled with the population name.

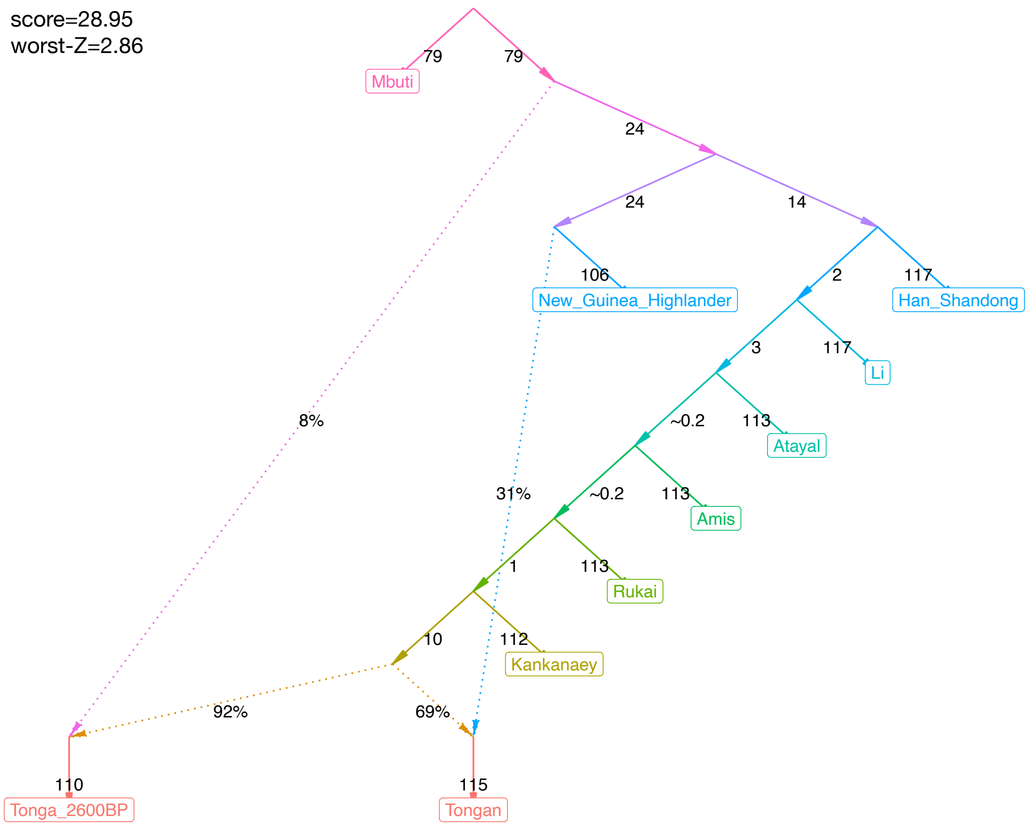


**Fig. S14. Best-fitting admixture graph for Out-of-Taiwan events.** The numbers on the solid lines denote are in genetic drift units while those on the dashed lines indicate the ancestry proportions. The ADMIXTOOLS 2 graph score and the worst-fitting Z score are shown on the top-left.

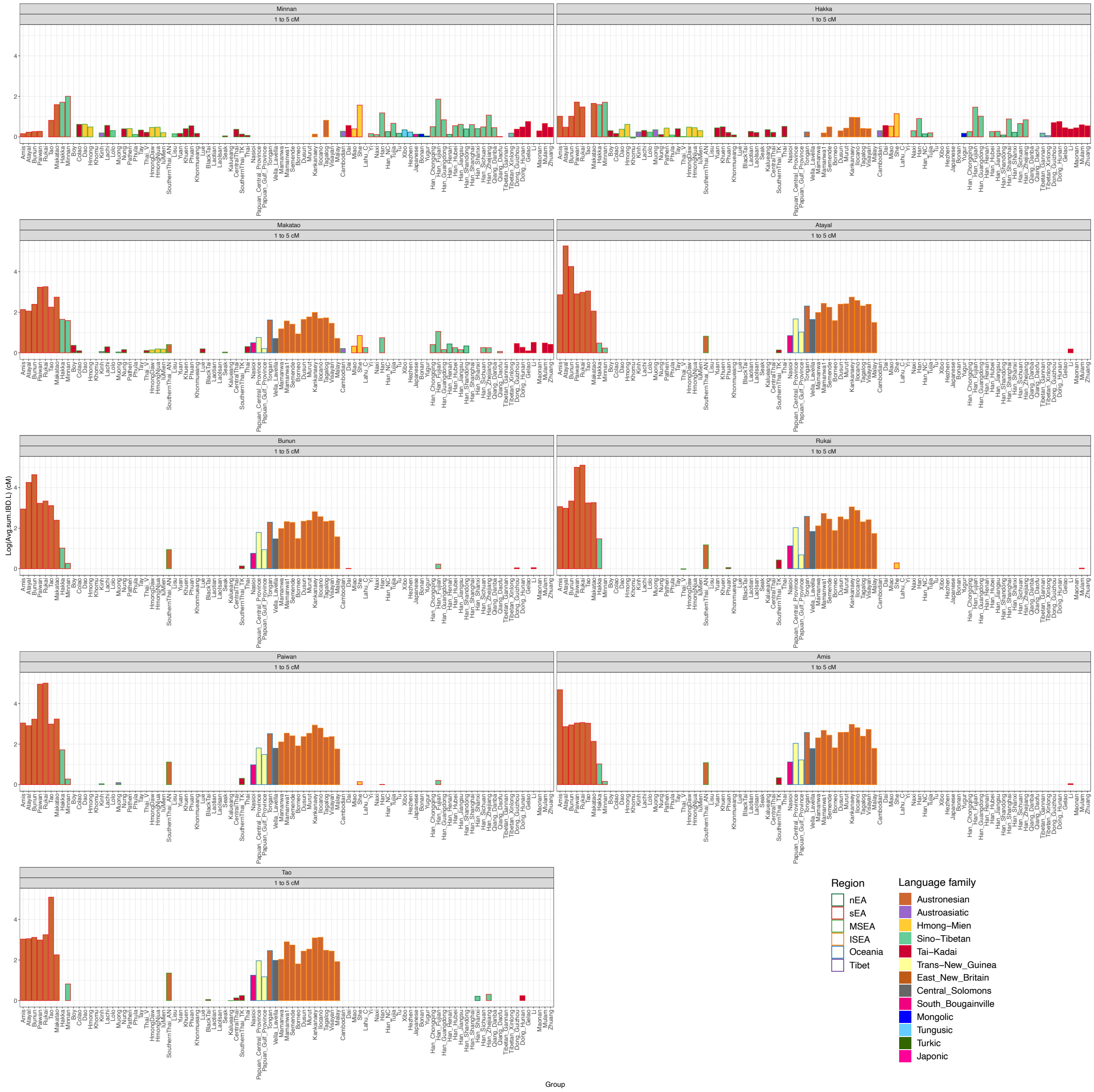


**Fig. S15. Quantification of all IBD sharing between each Taiwanese group and East Asian (including other Taiwanese) and Oceanian groups in segment size range of 1 to 5 cM.** The average summed IBD length between Taiwanese groups (in rows) and other East Asian and Oceanian groups is depicted in the bar plots; filled bars are colored according to the languages spoken by the compared East Asian/Oceanian groups shown on the x-axis while the outline color of the empty bars indicates their regions.

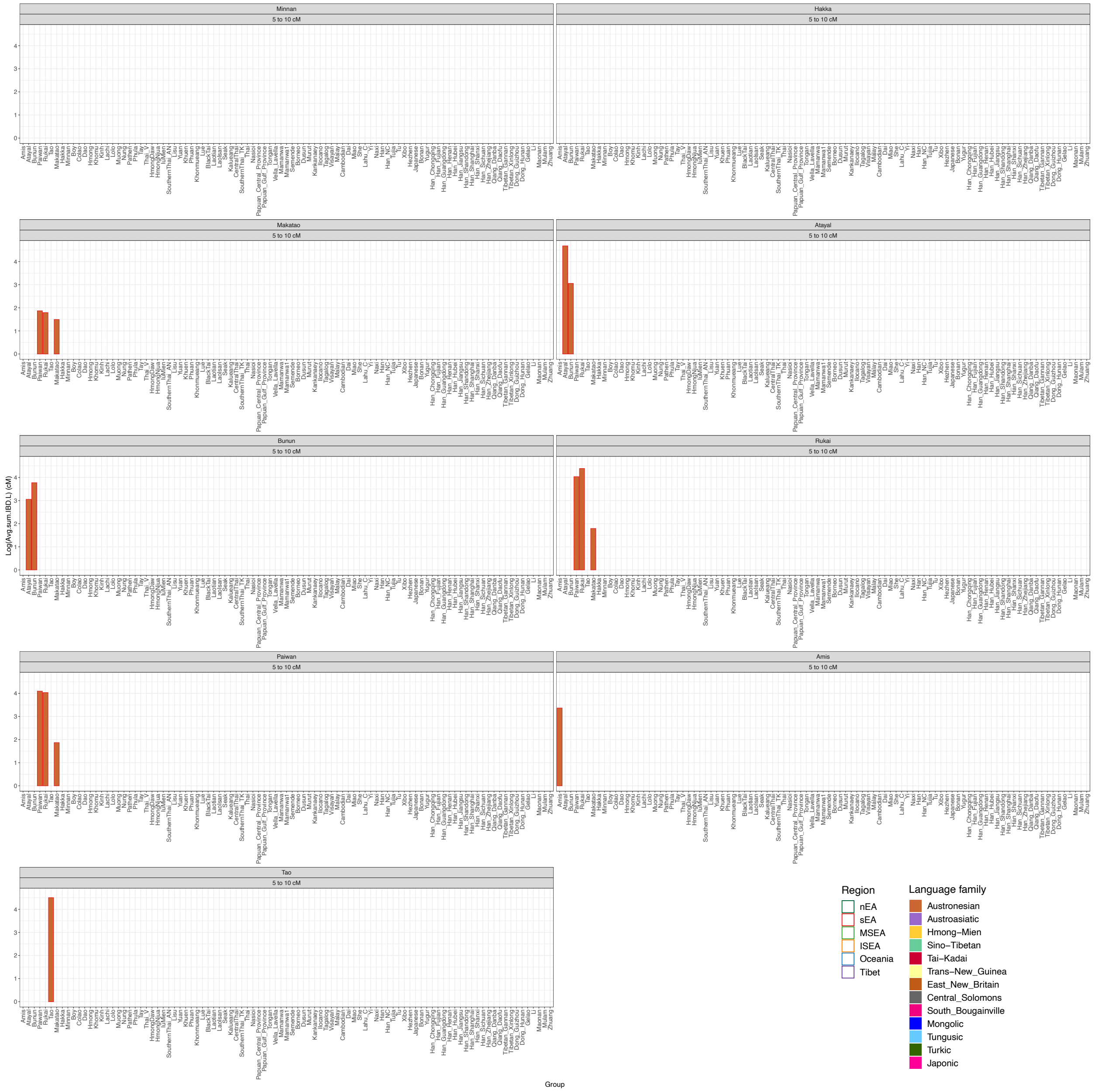


**Fig. S16. Quantification of all IBD sharing between each Taiwanese group and East Asian (including other Taiwanese) and Oceanian groups in segment size range of 5 to 10 cM.** The average summed IBD length between Taiwanese groups (in rows) and other East Asian and Oceanian groups is depicted in the bar plots; filled bars are colored according to the languages spoken by the compared East Asian/Oceanian groups shown on the x-axis while the outline color of the empty bars indicates their regions.

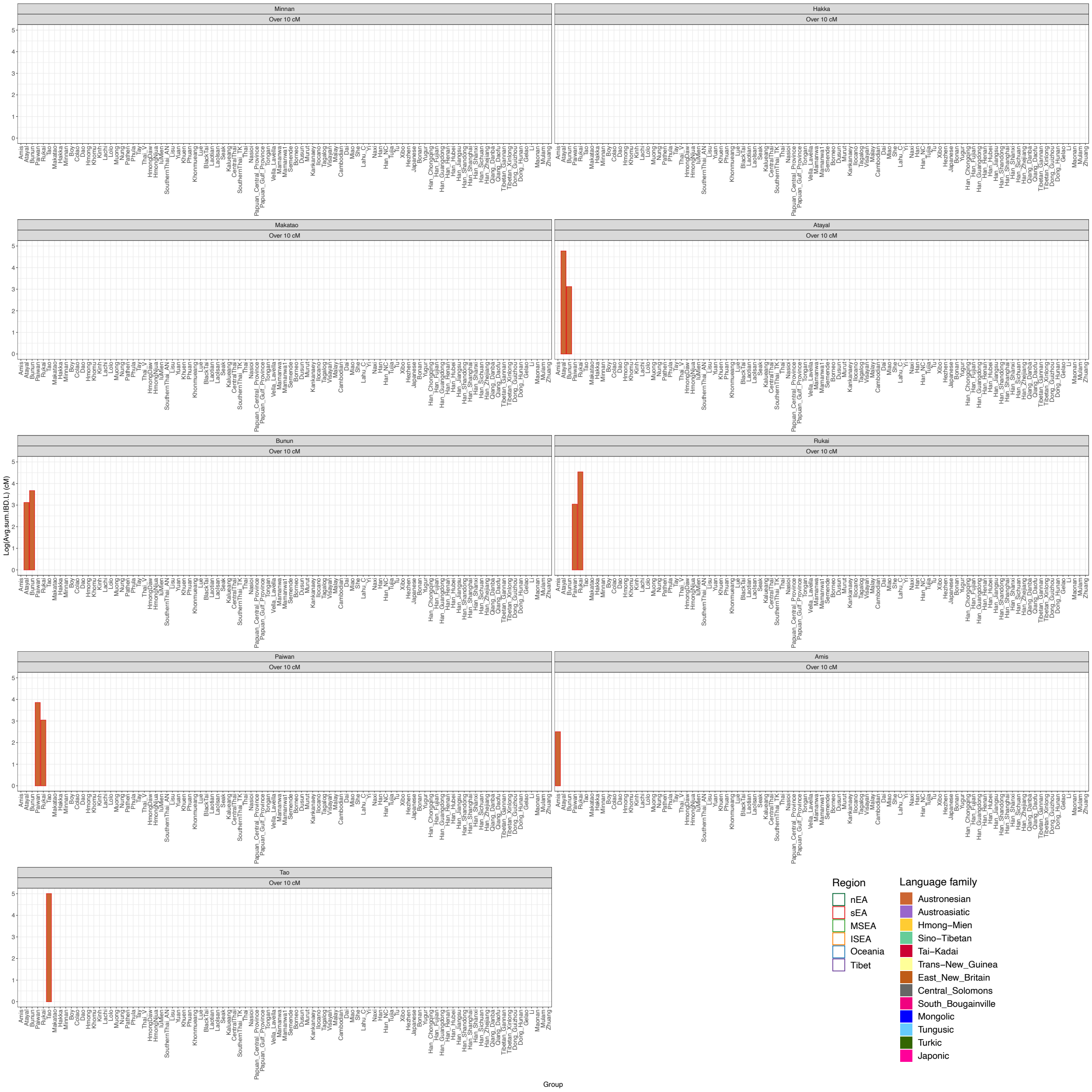


**Fig. S17. Quantification of all IBD sharing between each Taiwanese group and East Asian (including other Taiwanese) and Oceanian groups in segment size range of over 10 cM.** The average summed IBD length between Taiwanese groups (in rows) and other East Asian and Oceanian groups is depicted in the bar plots; filled bars are colored according to the languages spoken by the compared East Asian/Oceanian groups shown on the x-axis while the outline color of the empty bars indicates their regions.

### Supplementary Figures

**Table.S1. Metadata of the individuals in this study.** In Region column, sEA is southern East Asia, nEA is northern East Asia, MSEA is Mainland Southeast Asia, and ISEA is Island Southeast Asia. CP_Group is group for groupping in ChromoPainter analyses. In QC, imiss is missingness per individual, kin is 1st degree kinship, ancient indviduals only used in qpAdm as outgroups are noted.

| Individual_ID | Population | Region | Country | Type | Age | Language | Latitude | Longitude | CP_Group | TW_Group | Reference | QC |
|---|---|---|---|---|---|---|---|---|---|---|---|---|
| TA199 | Amis | sEA | Taiwan | modern | 0 | Austronesian | 22.84 | 121.19 | Amis | TW_S | This_study | PASS |
| TA29 | Amis | sEA | Taiwan | modern | 0 | Austronesian | 22.84 | 121.19 | Amis | TW_S | This_study | PASS |
| TA245 | Amis | sEA | Taiwan | modern | 0 | Austronesian | 22.84 | 121.19 | Amis | TW_S | This_study | PASS |
| TA81 | Amis | sEA | Taiwan | modern | 0 | Austronesian | 22.84 | 121.19 | Amis | TW_S | This_study | PASS |
| TA258 | Amis | sEA | Taiwan | modern | 0 | Austronesian | 22.84 | 121.19 | Amis | TW_S | This_study | PASS |
| TA95 | Amis | sEA | Taiwan | modern | 0 | Austronesian | 22.84 | 121.19 | Amis | TW_S | This_study | PASS |
| TA101 | Amis | sEA | Taiwan | modern | 0 | Austronesian | 22.84 | 121.19 | Amis | TW_S | This_study | PASS |
| TA131 | Amis | sEA | Taiwan | modern | 0 | Austronesian | 22.84 | 121.19 | Amis | TW_S | This_study | PASS |
| TA138 | Amis | sEA | Taiwan | modern | 0 | Austronesian | 22.84 | 121.19 | Amis | TW_S | This_study | PASS |
| ata34 | Atayal | sEA | Taiwan | modern | 0 | Austronesian | 24.61 | 121.3 | Atayal | TW_N | This_study | PASS |
| ata46 | Atayal | sEA | Taiwan | modern | 0 | Austronesian | 24.61 | 121.3 | Atayal | TW_N | This_study | PASS |
| ata30 | Atayal | sEA | Taiwan | modern | 0 | Austronesian | 24.61 | 121.3 | Atayal | TW_N | This_study | PASS |
| ata141 | Atayal | sEA | Taiwan | modern | 0 | Austronesian | 24.61 | 121.3 | Atayal | TW_N | This_study | PASS |
| ata10 | Atayal | sEA | Taiwan | modern | 0 | Austronesian | 24.61 | 121.3 | Atayal | TW_N | This_study | PASS |
| ata4 | Atayal | sEA | Taiwan | modern | 0 | Austronesian | 24.61 | 121.3 | Atayal | TW_N | This_study | PASS |
| ata1 | Atayal | sEA | Taiwan | modern | 0 | Austronesian | 24.61 | 121.3 | Atayal | TW_N | This_study | PASS |
| bun55 | Bunun | sEA | Taiwan | modern | 0 | Austronesian | 23.21 | 120.7 | Bunun | TW_C | This_study | PASS |
| bun34 | Bunun | sEA | Taiwan | modern | 0 | Austronesian | 23.21 | 120.7 | Bunun | TW_C | This_study | PASS |
| bun32 | Bunun | sEA | Taiwan | modern | 0 | Austronesian | 23.21 | 120.7 | Bunun | TW_C | This_study | PASS |
| bun38 | Bunun | sEA | Taiwan | modern | 0 | Austronesian | 23.21 | 120.7 | Bunun | TW_C | This_study | PASS |
| bun30 | Bunun | sEA | Taiwan | modern | 0 | Austronesian | 23.21 | 120.7 | Bunun | TW_C | This_study | PASS |
| bun39 | Bunun | sEA | Taiwan | modern | 0 | Austronesian | 23.21 | 120.7 | Bunun | TW_C | This_study | PASS |
| bun48 | Bunun | sEA | Taiwan | modern | 0 | Austronesian | 23.21 | 120.7 | Bunun | TW_C | This_study | PASS |
| pai10 | Paiwan | sEA | Taiwan | modern | 0 | Austronesian | 22.68 | 120.69 | Paiwan | TW_S | This_study | PASS |
| pai3 | Paiwan | sEA | Taiwan | modern | 0 | Austronesian | 22.68 | 120.69 | Paiwan | TW_S | This_study | PASS |
| pai58 | Paiwan | sEA | Taiwan | modern | 0 | Austronesian | 22.68 | 120.69 | Paiwan | TW_S | This_study | PASS |
| pai20 | Paiwan | sEA | Taiwan | modern | 0 | Austronesian | 22.68 | 120.69 | Paiwan | TW_S | This_study | PASS |
| pai103 | Paiwan | sEA | Taiwan | modern | 0 | Austronesian | 22.68 | 120.69 | Paiwan | TW_S | This_study | PASS |
| pai55 | Paiwan | sEA | Taiwan | modern | 0 | Austronesian | 22.68 | 120.69 | Paiwan | TW_S | This_study | PASS |
| pai25 | Paiwan | sEA | Taiwan | modern | 0 | Austronesian | 22.68 | 120.69 | Paiwan | TW_S | This_study | imiss |
| ruk487 | Rukai | sEA | Taiwan | modern | 0 | Austronesian | 22.7 | 120.61 | Rukai | TW_S | This_study | PASS |
| ruk540 | Rukai | sEA | Taiwan | modern | 0 | Austronesian | 22.7 | 120.61 | Rukai | TW_S | This_study | PASS |
| ruk111 | Rukai | sEA | Taiwan | modern | 0 | Austronesian | 22.7 | 120.61 | Rukai | TW_S | This_study | PASS |
| ruk502 | Rukai | sEA | Taiwan | modern | 0 | Austronesian | 22.7 | 120.61 | Rukai | TW_S | This_study | PASS |

| | | | | | | | | | | | | |
|---|---|---|---|---|---|---|---|---|---|---|---|---|
| ruk538 | Rukai | sEA | Taiwan | modern | 0 | Austronesian | 22.7 | 120.61 | Rukai | TW_S | This_study | PASS |
| ruk499 | Rukai | sEA | Taiwan | modern | 0 | Austronesian | 22.7 | 120.61 | Rukai | TW_S | This_study | PASS |
| ruk535 | Rukai | sEA | Taiwan | modern | 0 | Austronesian | 22.7 | 120.61 | Rukai | TW_S | This_study | PASS |
| tao45 | Tao | sEA | Taiwan | modern | 0 | Austronesian | 22.03 | 121.55 | Tao | TW_Orchid | This_study | PASS |
| pin33 | Makatao | sEA | Taiwan | modern | 0 | Austronesian | 22.6 | 120.61 | Makatao | TW_Lowland | This_study | PASS |
| pin29 | Makatao | sEA | Taiwan | modern | 0 | Austronesian | 22.6 | 120.61 | Makatao | TW_Lowland | This_study | PASS |
| pin74 | Makatao | sEA | Taiwan | modern | 0 | Austronesian | 22.6 | 120.61 | Makatao | TW_Lowland | This_study | PASS |
| pin126 | Makatao | sEA | Taiwan | modern | 0 | Austronesian | 22.6 | 120.61 | Makatao | TW_Lowland | This_study | PASS |
| pin81 | Makatao | sEA | Taiwan | modern | 0 | Austronesian | 22.6 | 120.61 | Makatao | TW_Lowland | This_study | PASS |
| hak95 | Hakka | sEA | Taiwan | modern | 0 | Sino-Tibetan | 22.67 | 120.6 | Hakka | TW_Han | This_study | PASS |
| hak19 | Hakka | sEA | Taiwan | modern | 0 | Sino-Tibetan | 22.67 | 120.6 | Hakka | TW_Han | This_study | PASS |
| hak13 | Hakka | sEA | Taiwan | modern | 0 | Sino-Tibetan | 22.67 | 120.6 | Hakka | TW_Han | This_study | PASS |
| hak23 | Hakka | sEA | Taiwan | modern | 0 | Sino-Tibetan | 22.67 | 120.6 | Hakka | TW_Han | This_study | PASS |
| hak33 | Hakka | sEA | Taiwan | modern | 0 | Sino-Tibetan | 22.67 | 120.6 | Hakka | TW_Han | This_study | PASS |
| hak36 | Hakka | sEA | Taiwan | modern | 0 | Sino-Tibetan | 22.67 | 120.6 | Hakka | TW_Han | This_study | PASS |
| min186 | Minnan | sEA | Taiwan | modern | 0 | Sino-Tibetan | 22.63 | 120.31 | Minnan | TW_Han | This_study | PASS |
| min235 | Minnan | sEA | Taiwan | modern | 0 | Sino-Tibetan | 22.63 | 120.31 | Minnan | TW_Han | This_study | PASS |
| min396 | Minnan | sEA | Taiwan | modern | 0 | Sino-Tibetan | 22.63 | 120.31 | Minnan | TW_Han | This_study | PASS |
| min305 | Minnan | sEA | Taiwan | modern | 0 | Sino-Tibetan | 22.63 | 120.31 | Minnan | TW_Han | This_study | PASS |
| min321 | Minnan | sEA | Taiwan | modern | 0 | Sino-Tibetan | 22.63 | 120.31 | Minnan | TW_Han | This_study | PASS |
| min76 | Minnan | sEA | Taiwan | modern | 0 | Sino-Tibetan | 22.63 | 120.31 | Minnan | TW_Han | This_study | PASS |
| BoY180 | Boy | MSEA | Vietnam | modern | 0 | Tai-Kadai | 22.7 | 104.1 | Boy | N.A. | Liu_et_al_2020 | PASS |
| BoY181 | Boy | MSEA | Vietnam | modern | 0 | Tai-Kadai | 22.7 | 104.1 | Boy | N.A. | Liu_et_al_2020 | kin |
| BoY182 | Boy | MSEA | Vietnam | modern | 0 | Tai-Kadai | 22.7 | 104.1 | Boy | N.A. | Liu_et_al_2020 | PASS |
| BoY184 | Boy | MSEA | Vietnam | modern | 0 | Tai-Kadai | 22.7 | 104.1 | Boy | N.A. | Liu_et_al_2020 | PASS |
| BoY185 | Boy | MSEA | Vietnam | modern | 0 | Tai-Kadai | 22.7 | 104.1 | Boy | N.A. | Liu_et_al_2020 | PASS |
| BoY186 | Boy | MSEA | Vietnam | modern | 0 | Tai-Kadai | 22.7 | 104.1 | Boy | N.A. | Liu_et_al_2020 | kin |
| BoY188 | Boy | MSEA | Vietnam | modern | 0 | Tai-Kadai | 22.7 | 104.1 | Boy | N.A. | Liu_et_al_2020 | kin |
| BoY189 | Boy | MSEA | Vietnam | modern | 0 | Tai-Kadai | 22.7 | 104.1 | Boy | N.A. | Liu_et_al_2020 | PASS |
| BoY190 | Boy | MSEA | Vietnam | modern | 0 | Tai-Kadai | 22.7 | 104.1 | Boy | N.A. | Liu_et_al_2020 | PASS |
| BoY191 | Boy | MSEA | Vietnam | modern | 0 | Tai-Kadai | 22.7 | 104.1 | Boy | N.A. | Liu_et_al_2020 | kin |
| BoY192_2 | Boy | MSEA | Vietnam | modern | 0 | Tai-Kadai | 22.7 | 104.1 | Boy | N.A. | Liu_et_al_2020 | kin |
| BoY193_2 | Boy | MSEA | Vietnam | modern | 0 | Tai-Kadai | 22.7 | 104.1 | Boy | N.A. | Liu_et_al_2020 | kin |
| Cham58 | Cham | MSEA | Vietnam | modern | 0 | Austronesian | 11.5 | 108.9 | Cham | N.A. | Liu_et_al_2020 | PASS |
| Cham64 | Cham | MSEA | Vietnam | modern | 0 | Austronesian | 11.5 | 108.9 | Cham | N.A. | Liu_et_al_2020 | PASS |
| Cham66 | Cham | MSEA | Vietnam | modern | 0 | Austronesian | 11.5 | 108.9 | Cham | N.A. | Liu_et_al_2020 | PASS |
| Cham75 | Cham | MSEA | Vietnam | modern | 0 | Austronesian | 11.5 | 108.9 | Cham | N.A. | Liu_et_al_2020 | PASS |
| Cham80 | Cham | MSEA | Vietnam | modern | 0 | Austronesian | 11.5 | 108.9 | Cham | N.A. | Liu_et_al_2020 | PASS |
| Cham83 | Cham | MSEA | Vietnam | modern | 0 | Austronesian | 11.5 | 108.9 | Cham | N.A. | Liu_et_al_2020 | PASS |
| Cham89 | Cham | MSEA | Vietnam | modern | 0 | Austronesian | 11.5 | 108.9 | Cham | N.A. | Liu_et_al_2020 | PASS |
| Cham127 | Cham | MSEA | Vietnam | modern | 0 | Austronesian | 11.5 | 108.9 | Cham | N.A. | Liu_et_al_2020 | PASS |

| | | | | | | | | | | | | |
|---|---|---|---|---|---|---|---|---|---|---|---|---|
| Cham133 | Cham | MSEA | Vietnam | modern | 0 | Austronesian | 11.5 | 108.9 | Cham | N.A. | Liu_et_al_2020 | PASS |
| Cham137 | Cham | MSEA | Vietnam | modern | 0 | Austronesian | 11.5 | 108.9 | Cham | N.A. | Liu_et_al_2020 | PASS |
| CoLao566 | Colao | MSEA | Vietnam | modern | 0 | Tai-Kadai | 22.7 | 104.7 | Colao | N.A. | Liu_et_al_2020 | PASS |
| CoLao567 | Colao | MSEA | Vietnam | modern | 0 | Tai-Kadai | 22.7 | 104.7 | Colao | N.A. | Liu_et_al_2020 | PASS |
| CoLao568 | Colao | MSEA | Vietnam | modern | 0 | Tai-Kadai | 22.7 | 104.7 | Colao | N.A. | Liu_et_al_2020 | PASS |
| CoLao569 | Colao | MSEA | Vietnam | modern | 0 | Tai-Kadai | 22.7 | 104.7 | Colao | N.A. | Liu_et_al_2020 | PASS |
| CoLao570 | Colao | MSEA | Vietnam | modern | 0 | Tai-Kadai | 22.7 | 104.7 | Colao | N.A. | Liu_et_al_2020 | PASS |
| CoLao571 | Colao | MSEA | Vietnam | modern | 0 | Tai-Kadai | 22.7 | 104.7 | Colao | N.A. | Liu_et_al_2020 | PASS |
| CoLao572 | Colao | MSEA | Vietnam | modern | 0 | Tai-Kadai | 22.7 | 104.7 | Colao | N.A. | Liu_et_al_2020 | PASS |
| CoLao573 | Colao | MSEA | Vietnam | modern | 0 | Tai-Kadai | 22.7 | 104.7 | Colao | N.A. | Liu_et_al_2020 | PASS |
| CoLao574 | Colao | MSEA | Vietnam | modern | 0 | Tai-Kadai | 22.7 | 104.7 | Colao | N.A. | Liu_et_al_2020 | PASS |
| CoLao575 | Colao | MSEA | Vietnam | modern | 0 | Tai-Kadai | 22.7 | 104.7 | Colao | N.A. | Liu_et_al_2020 | PASS |
| CoLao576 | Colao | MSEA | Vietnam | modern | 0 | Tai-Kadai | 22.7 | 104.7 | Colao | N.A. | Liu_et_al_2020 | PASS |
| CoLao577 | Colao | MSEA | Vietnam | modern | 0 | Tai-Kadai | 22.7 | 104.7 | Colao | N.A. | Liu_et_al_2020 | PASS |
| Cong410 | Cong | MSEA | Vietnam | modern | 0 | Sino-Tibetan | 22.4 | 102.7 | Cong | N.A. | Liu_et_al_2020 | PASS |
| Cong411 | Cong | MSEA | Vietnam | modern | 0 | Sino-Tibetan | 22.4 | 102.7 | Cong | N.A. | Liu_et_al_2020 | PASS |
| Cong412 | Cong | MSEA | Vietnam | modern | 0 | Sino-Tibetan | 22.4 | 102.7 | Cong | N.A. | Liu_et_al_2020 | kin |
| Cong413 | Cong | MSEA | Vietnam | modern | 0 | Sino-Tibetan | 22.4 | 102.7 | Cong | N.A. | Liu_et_al_2020 | PASS |
| Cong414 | Cong | MSEA | Vietnam | modern | 0 | Sino-Tibetan | 22.4 | 102.7 | Cong | N.A. | Liu_et_al_2020 | PASS |
| Cong415 | Cong | MSEA | Vietnam | modern | 0 | Sino-Tibetan | 22.4 | 102.7 | Cong | N.A. | Liu_et_al_2020 | PASS |
| Cong416 | Cong | MSEA | Vietnam | modern | 0 | Sino-Tibetan | 22.4 | 102.7 | Cong | N.A. | Liu_et_al_2020 | PASS |
| Cong420 | Cong | MSEA | Vietnam | modern | 0 | Sino-Tibetan | 22.4 | 102.7 | Cong | N.A. | Liu_et_al_2020 | PASS |
| Cong421 | Cong | MSEA | Vietnam | modern | 0 | Sino-Tibetan | 22.4 | 102.7 | Cong | N.A. | Liu_et_al_2020 | PASS |
| Cong423 | Cong | MSEA | Vietnam | modern | 0 | Sino-Tibetan | 22.4 | 102.7 | Cong | N.A. | Liu_et_al_2020 | PASS |
| Cong424 | Cong | MSEA | Vietnam | modern | 0 | Sino-Tibetan | 22.4 | 102.7 | Cong | N.A. | Liu_et_al_2020 | PASS |
| Cong425 | Cong | MSEA | Vietnam | modern | 0 | Sino-Tibetan | 22.4 | 102.7 | Cong | N.A. | Liu_et_al_2020 | PASS |
| Dao50 | Dao | MSEA | Vietnam | modern | 0 | Hmong-Mien | 22.7 | 104.7 | N.A. | N.A. | Liu_et_al_2020 | PASS |
| Dao51 | Dao | MSEA | Vietnam | modern | 0 | Hmong-Mien | 22.7 | 104.7 | N.A. | N.A. | Liu_et_al_2020 | PASS |
| Dao52 | Dao | MSEA | Vietnam | modern | 0 | Hmong-Mien | 22.7 | 104.7 | N.A. | N.A. | Liu_et_al_2020 | PASS |
| Dao623 | Dao | MSEA | Vietnam | modern | 0 | Hmong-Mien | 22.7 | 104.7 | N.A. | N.A. | Liu_et_al_2020 | PASS |
| Dao624_2 | Dao | MSEA | Vietnam | modern | 0 | Hmong-Mien | 22.7 | 104.7 | N.A. | N.A. | Liu_et_al_2020 | PASS |
| Dao625 | Dao | MSEA | Vietnam | modern | 0 | Hmong-Mien | 22.7 | 104.7 | N.A. | N.A. | Liu_et_al_2020 | PASS |
| Dao626 | Dao | MSEA | Vietnam | modern | 0 | Hmong-Mien | 22.7 | 104.7 | N.A. | N.A. | Liu_et_al_2020 | PASS |
| Dao627 | Dao | MSEA | Vietnam | modern | 0 | Hmong-Mien | 22.7 | 104.7 | N.A. | N.A. | Liu_et_al_2020 | PASS |
| Dao629 | Dao | MSEA | Vietnam | modern | 0 | Hmong-Mien | 22.7 | 104.7 | N.A. | N.A. | Liu_et_al_2020 | PASS |
| Dao630 | Dao | MSEA | Vietnam | modern | 0 | Hmong-Mien | 22.7 | 104.7 | N.A. | N.A. | Liu_et_al_2020 | PASS |
| Dao631 | Dao | MSEA | Vietnam | modern | 0 | Hmong-Mien | 22.7 | 104.7 | N.A. | N.A. | Liu_et_al_2020 | PASS |
| Dao632 | Dao | MSEA | Vietnam | modern | 0 | Hmong-Mien | 22.7 | 104.7 | N.A. | N.A. | Liu_et_al_2020 | PASS |
| Ede61 | Ede | MSEA | Vietnam | modern | 0 | Austronesian | 12.9 | 108.1 | Ede | N.A. | Liu_et_al_2020 | PASS |
| Ede74 | Ede | MSEA | Vietnam | modern | 0 | Austronesian | 12.9 | 108.2 | Ede | N.A. | Liu_et_al_2020 | PASS |
| Ede95 | Ede | MSEA | Vietnam | modern | 0 | Austronesian | 12.7 | 108.8 | Ede | N.A. | Liu_et_al_2020 | PASS |

| | | | | | | | | | | | | |
|---|---|---|---|---|---|---|---|---|---|---|---|---|
| Ede97 | Ede | MSEA | Vietnam | modern | 0 | Austronesian | 12.7 | 108 | Ede | N.A. | Liu_et_al_2020 | PASS |
| Ede102 | Ede | MSEA | Vietnam | modern | 0 | Austronesian | 13 | 108.4 | Ede | N.A. | Liu_et_al_2020 | PASS |
| Ede112 | Ede | MSEA | Vietnam | modern | 0 | Austronesian | 12.9 | 108.9 | Ede | N.A. | Liu_et_al_2020 | PASS |
| Ede123 | Ede | MSEA | Vietnam | modern | 0 | Austronesian | 12.9 | 108.2 | Ede | N.A. | Liu_et_al_2020 | PASS |
| Ede736 | Ede | MSEA | Vietnam | modern | 0 | Austronesian | 12.8 | 108.6 | Ede | N.A. | Liu_et_al_2020 | PASS |
| Ede737 | Ede | MSEA | Vietnam | modern | 0 | Austronesian | 13 | 108.4 | Ede | N.A. | Liu_et_al_2020 | PASS |
| Ede738 | Ede | MSEA | Vietnam | modern | 0 | Austronesian | 12.7 | 108.8 | Ede | N.A. | Liu_et_al_2020 | PASS |
| Ede740 | Ede | MSEA | Vietnam | modern | 0 | Austronesian | 12.7 | 108.8 | Ede | N.A. | Liu_et_al_2020 | PASS |
| Giarai60 | Giarai | MSEA | Vietnam | modern | 0 | Austronesian | 13.2 | 108.7 | Giarai | N.A. | Liu_et_al_2020 | PASS |
| Giarai67 | Giarai | MSEA | Vietnam | modern | 0 | Austronesian | 13.3 | 108.4 | Giarai | N.A. | Liu_et_al_2020 | PASS |
| Giarai91 | Giarai | MSEA | Vietnam | modern | 0 | Austronesian | 13.6 | 108.1 | Giarai | N.A. | Liu_et_al_2020 | PASS |
| Giarai100 | Giarai | MSEA | Vietnam | modern | 0 | Austronesian | 13.2 | 108.7 | Giarai | N.A. | Liu_et_al_2020 | PASS |
| Giarai107 | Giarai | MSEA | Vietnam | modern | 0 | Austronesian | 13.3 | 108.4 | Giarai | N.A. | Liu_et_al_2020 | PASS |
| Giarai124 | Giarai | MSEA | Vietnam | modern | 0 | Austronesian | 13.6 | 107.8 | Giarai | N.A. | Liu_et_al_2020 | PASS |
| Giarai134 | Giarai | MSEA | Vietnam | modern | 0 | Austronesian | 14.1 | 107.6 | Giarai | N.A. | Liu_et_al_2020 | PASS |
| Giarai716 | Giarai | MSEA | Vietnam | modern | 0 | Austronesian | 13.8 | 107.7 | Giarai | N.A. | Liu_et_al_2020 | PASS |
| Giarai718 | Giarai | MSEA | Vietnam | modern | 0 | Austronesian | 14.2 | 108 | Giarai | N.A. | Liu_et_al_2020 | PASS |
| Giarai719 | Giarai | MSEA | Vietnam | modern | 0 | Austronesian | 14 | 108 | Giarai | N.A. | Liu_et_al_2020 | PASS |
| Giarai722 | Giarai | MSEA | Vietnam | modern | 0 | Austronesian | 13.3 | 108.4 | Giarai | N.A. | Liu_et_al_2020 | PASS |
| Giarai723 | Giarai | MSEA | Vietnam | modern | 0 | Austronesian | 13.6 | 107.8 | Giarai | N.A. | Liu_et_al_2020 | PASS |
| HaNhi44 | Hanhi | MSEA | Vietnam | modern | 0 | Sino-Tibetan | 22.4 | 102.7 | Hanhi | N.A. | Liu_et_al_2020 | PASS |
| HaNhi45 | Hanhi | MSEA | Vietnam | modern | 0 | Sino-Tibetan | 22.4 | 102.7 | Hanhi | N.A. | Liu_et_al_2020 | PASS |
| HaNhi46 | Hanhi | MSEA | Vietnam | modern | 0 | Sino-Tibetan | 22.4 | 102.7 | Hanhi | N.A. | Liu_et_al_2020 | PASS |
| HaNhi315 | Hanhi | MSEA | Vietnam | modern | 0 | Sino-Tibetan | 22.4 | 102.7 | Hanhi | N.A. | Liu_et_al_2020 | PASS |
| HaNhi317 | Hanhi | MSEA | Vietnam | modern | 0 | Sino-Tibetan | 22.4 | 102.7 | Hanhi | N.A. | Liu_et_al_2020 | PASS |
| HaNhi319 | Hanhi | MSEA | Vietnam | modern | 0 | Sino-Tibetan | 22.4 | 102.7 | Hanhi | N.A. | Liu_et_al_2020 | PASS |
| HaNhi320 | Hanhi | MSEA | Vietnam | modern | 0 | Sino-Tibetan | 22.4 | 102.7 | Hanhi | N.A. | Liu_et_al_2020 | PASS |
| HaNhi321 | Hanhi | MSEA | Vietnam | modern | 0 | Sino-Tibetan | 22.4 | 102.7 | Hanhi | N.A. | Liu_et_al_2020 | PASS |
| HaNhi322 | Hanhi | MSEA | Vietnam | modern | 0 | Sino-Tibetan | 22.4 | 102.7 | Hanhi | N.A. | Liu_et_al_2020 | kin |
| HaNhi323_2 | Hanhi | MSEA | Vietnam | modern | 0 | Sino-Tibetan | 22.4 | 102.7 | Hanhi | N.A. | Liu_et_al_2020 | PASS |
| HaNhi328 | Hanhi | MSEA | Vietnam | modern | 0 | Sino-Tibetan | 22.4 | 102.7 | Hanhi | N.A. | Liu_et_al_2020 | PASS |
| HaNhi329 | Hanhi | MSEA | Vietnam | modern | 0 | Sino-Tibetan | 22.4 | 102.7 | Hanhi | N.A. | Liu_et_al_2020 | PASS |
| Hmong194 | Hmong | MSEA | Vietnam | modern | 0 | Hmong-Mien | 21.4 | 103 | N.A. | N.A. | Liu_et_al_2020 | PASS |
| Hmong195 | Hmong | MSEA | Vietnam | modern | 0 | Hmong-Mien | 21.4 | 103 | N.A. | N.A. | Liu_et_al_2020 | PASS |
| Hmong196_2 | Hmong | MSEA | Vietnam | modern | 0 | Hmong-Mien | 21.4 | 103 | N.A. | N.A. | Liu_et_al_2020 | kin |
| Hmong197_1 | Hmong | MSEA | Vietnam | modern | 0 | Hmong-Mien | 21.4 | 103 | N.A. | N.A. | Liu_et_al_2020 | kin |
| Hmong198 | Hmong | MSEA | Vietnam | modern | 0 | Hmong-Mien | 21.4 | 103 | N.A. | N.A. | Liu_et_al_2020 | PASS |
| Hmong199 | Hmong | MSEA | Vietnam | modern | 0 | Hmong-Mien | 21.4 | 103 | N.A. | N.A. | Liu_et_al_2020 | PASS |
| Hmong200 | Hmong | MSEA | Vietnam | modern | 0 | Hmong-Mien | 21.4 | 103 | N.A. | N.A. | Liu_et_al_2020 | PASS |
| Hmong201 | Hmong | MSEA | Vietnam | modern | 0 | Hmong-Mien | 21.4 | 103 | N.A. | N.A. | Liu_et_al_2020 | PASS |
| Hmong202 | Hmong | MSEA | Vietnam | modern | 0 | Hmong-Mien | 21.4 | 103 | N.A. | N.A. | Liu_et_al_2020 | kin |

| | | | | | | | | | | | | |
|---|---|---|---|---|---|---|---|---|---|---|---|---|
| Hmong203 | Hmong | MSEA | Vietnam | modern | 0 | Hmong-Mien | 21.4 | 103 | N.A. | N.A. | Liu_et_al_2020 | PASS |
| Hmong204 | Hmong | MSEA | Vietnam | modern | 0 | Hmong-Mien | 21.4 | 103 | N.A. | N.A. | Liu_et_al_2020 | PASS |
| Hmong205 | Hmong | MSEA | Vietnam | modern | 0 | Hmong-Mien | 21.4 | 103 | N.A. | N.A. | Liu_et_al_2020 | PASS |
| KhoMu235 | Khomu | MSEA | Vietnam | modern | 0 | Austroasiatic | 21.4 | 103 | N.A. | N.A. | Liu_et_al_2020 | PASS |
| KhoMu236 | Khomu | MSEA | Vietnam | modern | 0 | Austroasiatic | 21.4 | 103 | N.A. | N.A. | Liu_et_al_2020 | PASS |
| KhoMu237 | Khomu | MSEA | Vietnam | modern | 0 | Austroasiatic | 21.4 | 103 | N.A. | N.A. | Liu_et_al_2020 | PASS |
| KhoMu238 | Khomu | MSEA | Vietnam | modern | 0 | Austroasiatic | 21.4 | 103 | N.A. | N.A. | Liu_et_al_2020 | PASS |
| KhoMu239 | Khomu | MSEA | Vietnam | modern | 0 | Austroasiatic | 21.4 | 103 | N.A. | N.A. | Liu_et_al_2020 | PASS |
| KhoMu240 | Khomu | MSEA | Vietnam | modern | 0 | Austroasiatic | 21.4 | 103 | N.A. | N.A. | Liu_et_al_2020 | PASS |
| KhoMu241 | Khomu | MSEA | Vietnam | modern | 0 | Austroasiatic | 21.4 | 103 | N.A. | N.A. | Liu_et_al_2020 | kin |
| KhoMu242 | Khomu | MSEA | Vietnam | modern | 0 | Austroasiatic | 21.4 | 103 | N.A. | N.A. | Liu_et_al_2020 | PASS |
| KhoMu244 | Khomu | MSEA | Vietnam | modern | 0 | Austroasiatic | 21.4 | 103 | N.A. | N.A. | Liu_et_al_2020 | kin |
| KhoMu245_2 | Khomu | MSEA | Vietnam | modern | 0 | Austroasiatic | 21.4 | 103 | N.A. | N.A. | Liu_et_al_2020 | PASS |
| KhoMu246 | Khomu | MSEA | Vietnam | modern | 0 | Austroasiatic | 21.4 | 103 | N.A. | N.A. | Liu_et_al_2020 | kin |
| KhoMu247 | Khomu | MSEA | Vietnam | modern | 0 | Austroasiatic | 21.4 | 103 | N.A. | N.A. | Liu_et_al_2020 | PASS |
| Kinh01_1 | Kinh | MSEA | Vietnam | modern | 0 | Austroasiatic | 21.4 | 103 | N.A. | N.A. | Liu_et_al_2020 | PASS |
| Kinh03 | Kinh | MSEA | Vietnam | modern | 0 | Austroasiatic | 21.2 | 106 | N.A. | N.A. | Liu_et_al_2020 | PASS |
| Kinh05 | Kinh | MSEA | Vietnam | modern | 0 | Austroasiatic | 21 | 105.9 | N.A. | N.A. | Liu_et_al_2020 | PASS |
| Kinh06 | Kinh | MSEA | Vietnam | modern | 0 | Austroasiatic | 21 | 105.9 | N.A. | N.A. | Liu_et_al_2020 | PASS |
| Kinh07 | Kinh | MSEA | Vietnam | modern | 0 | Austroasiatic | 21 | 105.9 | N.A. | N.A. | Liu_et_al_2020 | PASS |
| Kinh08 | Kinh | MSEA | Vietnam | modern | 0 | Austroasiatic | 21 | 105.9 | N.A. | N.A. | Liu_et_al_2020 | PASS |
| Kinh09 | Kinh | MSEA | Vietnam | modern | 0 | Austroasiatic | 21 | 105.9 | N.A. | N.A. | Liu_et_al_2020 | PASS |
| Kinh10 | Kinh | MSEA | Vietnam | modern | 0 | Austroasiatic | 21 | 105.9 | N.A. | N.A. | Liu_et_al_2020 | PASS |
| Kinh11_1 | Kinh | MSEA | Vietnam | modern | 0 | Austroasiatic | 21 | 105.9 | N.A. | N.A. | Liu_et_al_2020 | PASS |
| Kinh12 | Kinh | MSEA | Vietnam | modern | 0 | Austroasiatic | 21 | 105.9 | N.A. | N.A. | Liu_et_al_2020 | PASS |
| Kinh14 | Kinh | MSEA | Vietnam | modern | 0 | Austroasiatic | 21 | 105.9 | N.A. | N.A. | Liu_et_al_2020 | PASS |
| Kinh15 | Kinh | MSEA | Vietnam | modern | 0 | Austroasiatic | 22 | 106.9 | N.A. | N.A. | Liu_et_al_2020 | PASS |
| LaChi530 | Lachi | MSEA | Vietnam | modern | 0 | Tai-Kadai | 22.7 | 104.7 | Lachi | N.A. | Liu_et_al_2020 | PASS |
| LaChi532 | Lachi | MSEA | Vietnam | modern | 0 | Tai-Kadai | 22.7 | 104.7 | Lachi | N.A. | Liu_et_al_2020 | PASS |
| LaChi533 | Lachi | MSEA | Vietnam | modern | 0 | Tai-Kadai | 22.7 | 104.7 | Lachi | N.A. | Liu_et_al_2020 | PASS |
| LaChi534 | Lachi | MSEA | Vietnam | modern | 0 | Tai-Kadai | 22.7 | 104.7 | Lachi | N.A. | Liu_et_al_2020 | PASS |
| LaChi535 | Lachi | MSEA | Vietnam | modern | 0 | Tai-Kadai | 22.7 | 104.7 | Lachi | N.A. | Liu_et_al_2020 | PASS |
| LaChi554 | Lachi | MSEA | Vietnam | modern | 0 | Tai-Kadai | 22.7 | 104.7 | Lachi | N.A. | Liu_et_al_2020 | PASS |
| LaChi555 | Lachi | MSEA | Vietnam | modern | 0 | Tai-Kadai | 22.7 | 104.7 | Lachi | N.A. | Liu_et_al_2020 | PASS |
| LaChi556 | Lachi | MSEA | Vietnam | modern | 0 | Tai-Kadai | 22.7 | 104.7 | Lachi | N.A. | Liu_et_al_2020 | PASS |
| LaChi557 | Lachi | MSEA | Vietnam | modern | 0 | Tai-Kadai | 22.7 | 104.7 | Lachi | N.A. | Liu_et_al_2020 | PASS |
| LaChi558 | Lachi | MSEA | Vietnam | modern | 0 | Tai-Kadai | 22.7 | 104.7 | Lachi | N.A. | Liu_et_al_2020 | PASS |
| LaChi559 | Lachi | MSEA | Vietnam | modern | 0 | Tai-Kadai | 22.7 | 104.7 | Lachi | N.A. | Liu_et_al_2020 | PASS |
| LaChi560 | Lachi | MSEA | Vietnam | modern | 0 | Tai-Kadai | 22.7 | 104.7 | Lachi | N.A. | Liu_et_al_2020 | PASS |
| LaHu342 | Lahu_V | MSEA | Vietnam | modern | 0 | Sino-Tibetan | 22.4 | 102.7 | Lahu | N.A. | Liu_et_al_2020 | PASS |
| LaHu344 | Lahu_V | MSEA | Vietnam | modern | 0 | Sino-Tibetan | 22.4 | 102.7 | Lahu | N.A. | Liu_et_al_2020 | PASS |

| | | | | | | | | | | | | |
|---|---|---|---|---|---|---|---|---|---|---|---|---|
| LaHu347 | Lahu_V | MSEA | Vietnam | modern | 0 | Sino-Tibetan | 22.4 | 102.7 | Lahu | N.A. | Liu_et_al_2020 | PASS |
| LaHu348 | Lahu_V | MSEA | Vietnam | modern | 0 | Sino-Tibetan | 22.4 | 102.7 | Lahu | N.A. | Liu_et_al_2020 | PASS |
| LaHu350 | Lahu_V | MSEA | Vietnam | modern | 0 | Sino-Tibetan | 22.4 | 102.7 | Lahu | N.A. | Liu_et_al_2020 | PASS |
| LaHu351 | Lahu_V | MSEA | Vietnam | modern | 0 | Sino-Tibetan | 22.4 | 102.7 | Lahu | N.A. | Liu_et_al_2020 | PASS |
| LaHu352 | Lahu_V | MSEA | Vietnam | modern | 0 | Sino-Tibetan | 22.4 | 102.7 | Lahu | N.A. | Liu_et_al_2020 | kin |
| LaHu353 | Lahu_V | MSEA | Vietnam | modern | 0 | Sino-Tibetan | 22.4 | 102.7 | Lahu | N.A. | Liu_et_al_2020 | kin |
| LaHu354 | Lahu_V | MSEA | Vietnam | modern | 0 | Sino-Tibetan | 22.4 | 102.7 | Lahu | N.A. | Liu_et_al_2020 | kin |
| LaHu355 | Lahu_V | MSEA | Vietnam | modern | 0 | Sino-Tibetan | 22.4 | 102.7 | Lahu | N.A. | Liu_et_al_2020 | PASS |
| LaHu356 | Lahu_V | MSEA | Vietnam | modern | 0 | Sino-Tibetan | 22.4 | 102.7 | Lahu | N.A. | Liu_et_al_2020 | PASS |
| LaHu360 | Lahu_V | MSEA | Vietnam | modern | 0 | Sino-Tibetan | 22.4 | 102.7 | Lahu | N.A. | Liu_et_al_2020 | kin |
| LoLo469 | Lolo | MSEA | Vietnam | modern | 0 | Sino-Tibetan | 23.1 | 105.4 | Lolo | N.A. | Liu_et_al_2020 | PASS |
| Lolo470 | Lolo | MSEA | Vietnam | modern | 0 | Sino-Tibetan | 23.1 | 105.4 | Lolo | N.A. | Liu_et_al_2020 | PASS |
| LoLo471 | Lolo | MSEA | Vietnam | modern | 0 | Sino-Tibetan | 23.1 | 105.4 | Lolo | N.A. | Liu_et_al_2020 | PASS |
| LoLo472 | Lolo | MSEA | Vietnam | modern | 0 | Sino-Tibetan | 23.1 | 105.4 | Lolo | N.A. | Liu_et_al_2020 | kin |
| LoLo473 | Lolo | MSEA | Vietnam | modern | 0 | Sino-Tibetan | 23.1 | 105.4 | Lolo | N.A. | Liu_et_al_2020 | PASS |
| LoLo474 | Lolo | MSEA | Vietnam | modern | 0 | Sino-Tibetan | 23.1 | 105.4 | Lolo | N.A. | Liu_et_al_2020 | PASS |
| LoLo475 | Lolo | MSEA | Vietnam | modern | 0 | Sino-Tibetan | 23.1 | 105.4 | Lolo | N.A. | Liu_et_al_2020 | PASS |
| LoLo476 | Lolo | MSEA | Vietnam | modern | 0 | Sino-Tibetan | 23.1 | 105.4 | Lolo | N.A. | Liu_et_al_2020 | PASS |
| LoLo477 | Lolo | MSEA | Vietnam | modern | 0 | Sino-Tibetan | 23.1 | 105.4 | Lolo | N.A. | Liu_et_al_2020 | kin |
| LoLo478 | Lolo | MSEA | Vietnam | modern | 0 | Sino-Tibetan | 23.1 | 105.4 | Lolo | N.A. | Liu_et_al_2020 | PASS |
| LoLo479 | Lolo | MSEA | Vietnam | modern | 0 | Sino-Tibetan | 23.1 | 105.4 | Lolo | N.A. | Liu_et_al_2020 | PASS |
| LoLo480 | Lolo | MSEA | Vietnam | modern | 0 | Sino-Tibetan | 23.1 | 105.4 | Lolo | N.A. | Liu_et_al_2020 | PASS |
| Mang273 | Mang | MSEA | Vietnam | modern | 0 | Austroasiatic | 22.4 | 102.7 | N.A. | N.A. | Liu_et_al_2020 | PASS |
| Mang274 | Mang | MSEA | Vietnam | modern | 0 | Austroasiatic | 22.4 | 102.7 | N.A. | N.A. | Liu_et_al_2020 | PASS |
| Mang276 | Mang | MSEA | Vietnam | modern | 0 | Austroasiatic | 22.4 | 102.7 | N.A. | N.A. | Liu_et_al_2020 | PASS |
| Mang278_2 | Mang | MSEA | Vietnam | modern | 0 | Austroasiatic | 22.4 | 102.7 | N.A. | N.A. | Liu_et_al_2020 | PASS |
| Mang284_1 | Mang | MSEA | Vietnam | modern | 0 | Austroasiatic | 22.4 | 102.7 | N.A. | N.A. | Liu_et_al_2020 | PASS |
| Mang289 | Mang | MSEA | Vietnam | modern | 0 | Austroasiatic | 22.4 | 102.7 | N.A. | N.A. | Liu_et_al_2020 | PASS |
| Mang290 | Mang | MSEA | Vietnam | modern | 0 | Austroasiatic | 22.4 | 102.7 | N.A. | N.A. | Liu_et_al_2020 | PASS |
| Mang291 | Mang | MSEA | Vietnam | modern | 0 | Austroasiatic | 22.4 | 102.7 | N.A. | N.A. | Liu_et_al_2020 | PASS |
| Mang298 | Mang | MSEA | Vietnam | modern | 0 | Austroasiatic | 22.4 | 102.7 | N.A. | N.A. | Liu_et_al_2020 | PASS |
| Mang307_1 | Mang | MSEA | Vietnam | modern | 0 | Austroasiatic | 22.4 | 102.7 | N.A. | N.A. | Liu_et_al_2020 | PASS |
| Mang308 | Mang | MSEA | Vietnam | modern | 0 | Austroasiatic | 22.4 | 102.7 | N.A. | N.A. | Liu_et_al_2020 | PASS |
| Mang310 | Mang | MSEA | Vietnam | modern | 0 | Austroasiatic | 22.4 | 102.7 | N.A. | N.A. | Liu_et_al_2020 | kin |
| Muong35 | Muong | MSEA | Vietnam | modern | 0 | Austroasiatic | 20.4 | 105.6 | N.A. | N.A. | Liu_et_al_2020 | PASS |
| Muong36 | Muong | MSEA | Vietnam | modern | 0 | Austroasiatic | 20.4 | 105.6 | N.A. | N.A. | Liu_et_al_2020 | PASS |
| Muong38 | Muong | MSEA | Vietnam | modern | 0 | Austroasiatic | 20.5 | 105.4 | N.A. | N.A. | Liu_et_al_2020 | PASS |
| Muong39 | Muong | MSEA | Vietnam | modern | 0 | Austroasiatic | 20.7 | 105.6 | N.A. | N.A. | Liu_et_al_2020 | PASS |
| Muong40 | Muong | MSEA | Vietnam | modern | 0 | Austroasiatic | 20.9 | 105.5 | N.A. | N.A. | Liu_et_al_2020 | PASS |
| Muong41 | Muong | MSEA | Vietnam | modern | 0 | Austroasiatic | 20.7 | 105.6 | N.A. | N.A. | Liu_et_al_2020 | PASS |
| Muong42 | Muong | MSEA | Vietnam | modern | 0 | Austroasiatic | 20.7 | 105.6 | N.A. | N.A. | Liu_et_al_2020 | PASS |

| | | | | | | | | | | | | |
|---|---|---|---|---|---|---|---|---|---|---|---|---|
| Muong43 | Muong | MSEA | Vietnam | modern | 0 | Austroasiatic | 20.5 | 105.4 | N.A. | N.A. | Liu_et_al_2020 | PASS |
| Muong121 | Muong | MSEA | Vietnam | modern | 0 | Austroasiatic | 20.5 | 105.4 | N.A. | N.A. | Liu_et_al_2020 | PASS |
| Muong132 | Muong | MSEA | Vietnam | modern | 0 | Austroasiatic | 20.5 | 105.4 | N.A. | N.A. | Liu_et_al_2020 | PASS |
| Nung47 | Nung | MSEA | Vietnam | modern | 0 | Tai-Kadai | 22.8 | 105 | Nung | N.A. | Liu_et_al_2020 | PASS |
| Nung48 | Nung | MSEA | Vietnam | modern | 0 | Tai-Kadai | 22.8 | 105 | Nung | N.A. | Liu_et_al_2020 | PASS |
| Nung49 | Nung | MSEA | Vietnam | modern | 0 | Tai-Kadai | 22.9 | 106.1 | Nung | N.A. | Liu_et_al_2020 | PASS |
| Nung664 | Nung | MSEA | Vietnam | modern | 0 | Tai-Kadai | 22.7 | 104.7 | Nung | N.A. | Liu_et_al_2020 | PASS |
| Nung672_1 | Nung | MSEA | Vietnam | modern | 0 | Tai-Kadai | 22.7 | 104.7 | Nung | N.A. | Liu_et_al_2020 | PASS |
| Nung674 | Nung | MSEA | Vietnam | modern | 0 | Tai-Kadai | 22.7 | 104.7 | Nung | N.A. | Liu_et_al_2020 | PASS |
| Nung677 | Nung | MSEA | Vietnam | modern | 0 | Tai-Kadai | 22.7 | 104.7 | Nung | N.A. | Liu_et_al_2020 | PASS |
| Nung678 | Nung | MSEA | Vietnam | modern | 0 | Tai-Kadai | 22.7 | 104.7 | Nung | N.A. | Liu_et_al_2020 | PASS |
| Nung680 | Nung | MSEA | Vietnam | modern | 0 | Tai-Kadai | 22.7 | 104.7 | Nung | N.A. | Liu_et_al_2020 | PASS |
| Nung681 | Nung | MSEA | Vietnam | modern | 0 | Tai-Kadai | 22.7 | 104.7 | Nung | N.A. | Liu_et_al_2020 | PASS |
| Nung687 | Nung | MSEA | Vietnam | modern | 0 | Tai-Kadai | 22.7 | 104.7 | Nung | N.A. | Liu_et_al_2020 | PASS |
| Nung691 | Nung | MSEA | Vietnam | modern | 0 | Tai-Kadai | 22.7 | 104.7 | Nung | N.A. | Liu_et_al_2020 | PASS |
| PaThen494 | Pathen | MSEA | Vietnam | modern | 0 | Hmong-Mien | 22.4 | 104.7 | N.A. | N.A. | Liu_et_al_2020 | PASS |
| PaThen495 | Pathen | MSEA | Vietnam | modern | 0 | Hmong-Mien | 22.4 | 104.7 | N.A. | N.A. | Liu_et_al_2020 | PASS |
| PaThen496 | Pathen | MSEA | Vietnam | modern | 0 | Hmong-Mien | 22.4 | 104.7 | N.A. | N.A. | Liu_et_al_2020 | PASS |
| PaThen497 | Pathen | MSEA | Vietnam | modern | 0 | Hmong-Mien | 22.4 | 104.7 | N.A. | N.A. | Liu_et_al_2020 | PASS |
| PaThen498 | Pathen | MSEA | Vietnam | modern | 0 | Hmong-Mien | 22.4 | 104.7 | N.A. | N.A. | Liu_et_al_2020 | PASS |
| PaThen499 | Pathen | MSEA | Vietnam | modern | 0 | Hmong-Mien | 22.4 | 104.7 | N.A. | N.A. | Liu_et_al_2020 | PASS |
| PaThen500 | Pathen | MSEA | Vietnam | modern | 0 | Hmong-Mien | 22.4 | 104.7 | N.A. | N.A. | Liu_et_al_2020 | PASS |
| PaThen501 | Pathen | MSEA | Vietnam | modern | 0 | Hmong-Mien | 22.4 | 104.7 | N.A. | N.A. | Liu_et_al_2020 | PASS |
| PaThen502 | Pathen | MSEA | Vietnam | modern | 0 | Hmong-Mien | 22.4 | 104.7 | N.A. | N.A. | Liu_et_al_2020 | kin |
| PaThen503 | Pathen | MSEA | Vietnam | modern | 0 | Hmong-Mien | 22.4 | 104.7 | N.A. | N.A. | Liu_et_al_2020 | PASS |
| PaThen504 | Pathen | MSEA | Vietnam | modern | 0 | Hmong-Mien | 22.4 | 104.7 | N.A. | N.A. | Liu_et_al_2020 | PASS |
| PaThen505 | Pathen | MSEA | Vietnam | modern | 0 | Hmong-Mien | 22.4 | 104.7 | N.A. | N.A. | Liu_et_al_2020 | PASS |
| Phula141 | Phula | MSEA | Vietnam | modern | 0 | Sino-Tibetan | 22.7 | 104.1 | Phula | N.A. | Liu_et_al_2020 | PASS |
| PhuLa142 | Phula | MSEA | Vietnam | modern | 0 | Sino-Tibetan | 22.7 | 104.1 | Phula | N.A. | Liu_et_al_2020 | PASS |
| PhuLa143 | Phula | MSEA | Vietnam | modern | 0 | Sino-Tibetan | 22.7 | 104.1 | Phula | N.A. | Liu_et_al_2020 | PASS |
| PhuLa144 | Phula | MSEA | Vietnam | modern | 0 | Sino-Tibetan | 22.7 | 104.1 | Phula | N.A. | Liu_et_al_2020 | PASS |
| PhuLa145 | Phula | MSEA | Vietnam | modern | 0 | Sino-Tibetan | 22.7 | 104.1 | Phula | N.A. | Liu_et_al_2020 | PASS |
| PhuLa146 | Phula | MSEA | Vietnam | modern | 0 | Sino-Tibetan | 22.7 | 104.1 | Phula | N.A. | Liu_et_al_2020 | PASS |
| PhuLa147 | Phula | MSEA | Vietnam | modern | 0 | Sino-Tibetan | 22.7 | 104.1 | Phula | N.A. | Liu_et_al_2020 | PASS |
| PhuLa148 | Phula | MSEA | Vietnam | modern | 0 | Sino-Tibetan | 22.7 | 104.1 | Phula | N.A. | Liu_et_al_2020 | PASS |
| PhuLa149 | Phula | MSEA | Vietnam | modern | 0 | Sino-Tibetan | 22.7 | 104.1 | Phula | N.A. | Liu_et_al_2020 | PASS |
| PhuLa152 | Phula | MSEA | Vietnam | modern | 0 | Sino-Tibetan | 22.7 | 104.1 | Phula | N.A. | Liu_et_al_2020 | PASS |
| PhuLa153 | Phula | MSEA | Vietnam | modern | 0 | Sino-Tibetan | 22.7 | 104.1 | Phula | N.A. | Liu_et_al_2020 | PASS |
| Phula1450 | Phula | MSEA | Vietnam | modern | 0 | Sino-Tibetan | 22.7 | 104.1 | Phula | N.A. | Liu_et_al_2020 | PASS |
| Sila427 | Sila | MSEA | Vietnam | modern | 0 | Sino-Tibetan | 22.4 | 102.7 | Sila | N.A. | Liu_et_al_2020 | PASS |
| Sila428 | Sila | MSEA | Vietnam | modern | 0 | Sino-Tibetan | 22.4 | 102.7 | Sila | N.A. | Liu_et_al_2020 | PASS |

| | | | | | | | | | | | | |
|---|---|---|---|---|---|---|---|---|---|---|---|---|
| Sila429 | Sila | MSEA | Vietnam | modern | 0 | Sino-Tibetan | 22.4 | 102.7 | Sila | N.A. | Liu_et_al_2020 | PASS |
| Sila433_1 | Sila | MSEA | Vietnam | modern | 0 | Sino-Tibetan | 22.4 | 102.7 | Sila | N.A. | Liu_et_al_2020 | PASS |
| Sila434 | Sila | MSEA | Vietnam | modern | 0 | Sino-Tibetan | 22.4 | 102.7 | Sila | N.A. | Liu_et_al_2020 | PASS |
| Sila435 | Sila | MSEA | Vietnam | modern | 0 | Sino-Tibetan | 22.4 | 102.7 | Sila | N.A. | Liu_et_al_2020 | kin |
| Sila436_2 | Sila | MSEA | Vietnam | modern | 0 | Sino-Tibetan | 22.4 | 102.7 | Sila | N.A. | Liu_et_al_2020 | PASS |
| Sila438 | Sila | MSEA | Vietnam | modern | 0 | Sino-Tibetan | 22.4 | 102.7 | Sila | N.A. | Liu_et_al_2020 | PASS |
| Sila439_2 | Sila | MSEA | Vietnam | modern | 0 | Sino-Tibetan | 22.4 | 102.7 | Sila | N.A. | Liu_et_al_2020 | PASS |
| Sila440 | Sila | MSEA | Vietnam | modern | 0 | Sino-Tibetan | 22.4 | 102.7 | Sila | N.A. | Liu_et_al_2020 | PASS |
| Sila441 | Sila | MSEA | Vietnam | modern | 0 | Sino-Tibetan | 22.4 | 102.7 | Sila | N.A. | Liu_et_al_2020 | PASS |
| Sila443 | Sila | MSEA | Vietnam | modern | 0 | Sino-Tibetan | 22.4 | 102.7 | Sila | N.A. | Liu_et_al_2020 | PASS |
| Tay84 | Tay | MSEA | Vietnam | modern | 0 | Tai-Kadai | 11.7 | 108.4 | Tay | N.A. | Liu_et_al_2020 | PASS |
| Tay87 | Tay | MSEA | Vietnam | modern | 0 | Tai-Kadai | 21.6 | 105.6 | Tay | N.A. | Liu_et_al_2020 | PASS |
| Tay138 | Tay | MSEA | Vietnam | modern | 0 | Tai-Kadai | 22.7 | 104.1 | Tay | N.A. | Liu_et_al_2020 | PASS |
| Tay155 | Tay | MSEA | Vietnam | modern | 0 | Tai-Kadai | 22.7 | 104.1 | Tay | N.A. | Liu_et_al_2020 | PASS |
| Tay156 | Tay | MSEA | Vietnam | modern | 0 | Tai-Kadai | 22.7 | 104.1 | Tay | N.A. | Liu_et_al_2020 | PASS |
| Tay157 | Tay | MSEA | Vietnam | modern | 0 | Tai-Kadai | 22.7 | 104.1 | Tay | N.A. | Liu_et_al_2020 | PASS |
| Tay158 | Tay | MSEA | Vietnam | modern | 0 | Tai-Kadai | 22.7 | 104.1 | Tay | N.A. | Liu_et_al_2020 | PASS |
| Tay160 | Tay | MSEA | Vietnam | modern | 0 | Tai-Kadai | 22.7 | 104.1 | Tay | N.A. | Liu_et_al_2020 | PASS |
| Tay161 | Tay | MSEA | Vietnam | modern | 0 | Tai-Kadai | 22.7 | 104.1 | Tay | N.A. | Liu_et_al_2020 | PASS |
| Tay163 | Tay | MSEA | Vietnam | modern | 0 | Tai-Kadai | 22.7 | 104.1 | Tay | N.A. | Liu_et_al_2020 | PASS |
| Tay164 | Tay | MSEA | Vietnam | modern | 0 | Tai-Kadai | 22.7 | 104.1 | Tay | N.A. | Liu_et_al_2020 | PASS |
| Tay162_1 | Tay | MSEA | Vietnam | modern | 0 | Tai-Kadai | 22.7 | 104.1 | Tay | N.A. | Liu_et_al_2020 | PASS |
| Thai55 | Thai_V | MSEA | Vietnam | modern | 0 | Tai-Kadai | 22 | 103.8 | Thai_V | N.A. | Liu_et_al_2020 | PASS |
| Thai56 | Thai_V | MSEA | Vietnam | modern | 0 | Tai-Kadai | 22 | 103.8 | Thai_V | N.A. | Liu_et_al_2020 | PASS |
| Thai57 | Thai_V | MSEA | Vietnam | modern | 0 | Tai-Kadai | 20.9 | 105.4 | Thai_V | N.A. | Liu_et_al_2020 | PASS |
| Thai248 | Thai_V | MSEA | Vietnam | modern | 0 | Tai-Kadai | 21.4 | 103 | Thai_V | N.A. | Liu_et_al_2020 | PASS |
| Thai249_2 | Thai_V | MSEA | Vietnam | modern | 0 | Tai-Kadai | 21.4 | 103 | Thai_V | N.A. | Liu_et_al_2020 | PASS |
| Thai250 | Thai_V | MSEA | Vietnam | modern | 0 | Tai-Kadai | 21.4 | 103 | Thai_V | N.A. | Liu_et_al_2020 | PASS |
| Thai251 | Thai_V | MSEA | Vietnam | modern | 0 | Tai-Kadai | 21.4 | 103 | Thai_V | N.A. | Liu_et_al_2020 | PASS |
| Thai252 | Thai_V | MSEA | Vietnam | modern | 0 | Tai-Kadai | 21.4 | 103 | Thai_V | N.A. | Liu_et_al_2020 | PASS |
| Thai253_2 | Thai_V | MSEA | Vietnam | modern | 0 | Tai-Kadai | 21.4 | 103 | Thai_V | N.A. | Liu_et_al_2020 | PASS |
| Thai254 | Thai_V | MSEA | Vietnam | modern | 0 | Tai-Kadai | 21.4 | 103 | Thai_V | N.A. | Liu_et_al_2020 | PASS |
| Thai255 | Thai_V | MSEA | Vietnam | modern | 0 | Tai-Kadai | 21.4 | 103 | Thai_V | N.A. | Liu_et_al_2020 | PASS |
| Thai257 | Thai_V | MSEA | Vietnam | modern | 0 | Tai-Kadai | 21.4 | 103 | Thai_V | N.A. | Liu_et_al_2020 | PASS |
| HM401 | HmongDaw | MSEA | Thailand | modern | 0 | Hmong-Mien | 19.9 | 100.11 | N.A. | N.A. | Kutanan_et_al_2021 | PASS |
| HM403 | HmongDaw | MSEA | Thailand | modern | 0 | Hmong-Mien | 19.9 | 100.11 | N.A. | N.A. | Kutanan_et_al_2021 | PASS |
| HM404 | HmongDaw | MSEA | Thailand | modern | 0 | Hmong-Mien | 19.9 | 100.11 | N.A. | N.A. | Kutanan_et_al_2021 | PASS |
| HM405 | HmongDaw | MSEA | Thailand | modern | 0 | Hmong-Mien | 19.9 | 100.11 | N.A. | N.A. | Kutanan_et_al_2021 | PASS |
| HM406 | HmongDaw | MSEA | Thailand | modern | 0 | Hmong-Mien | 19.9 | 100.11 | N.A. | N.A. | Kutanan_et_al_2021 | PASS |
| HM407 | HmongDaw | MSEA | Thailand | modern | 0 | Hmong-Mien | 19.9 | 100.11 | N.A. | N.A. | Kutanan_et_al_2021 | PASS |
| HM409 | HmongDaw | MSEA | Thailand | modern | 0 | Hmong-Mien | 19.9 | 100.11 | N.A. | N.A. | Kutanan_et_al_2021 | PASS |

| | | | | | | | | | | | | |
|---|---|---|---|---|---|---|---|---|---|---|---|---|
| HM413 | HmongDaw | MSEA | Thailand | modern | 0 | Hmong-Mien | 19.9 | 100.11 | N.A. | N.A. | Kutanan_et_al_2021 | PASS |
| HM415 | HmongDaw | MSEA | Thailand | modern | 0 | Hmong-Mien | 19.9 | 100.11 | N.A. | N.A. | Kutanan_et_al_2021 | PASS |
| HM101 | HmongNjua | MSEA | Thailand | modern | 0 | Hmong-Mien | 18.12 | 98.23 | N.A. | N.A. | Kutanan_et_al_2021 | PASS |
| HM102 | HmongNjua | MSEA | Thailand | modern | 0 | Hmong-Mien | 18.12 | 98.23 | N.A. | N.A. | Kutanan_et_al_2021 | PASS |
| HM103 | HmongNjua | MSEA | Thailand | modern | 0 | Hmong-Mien | 19.09 | 100.56 | N.A. | N.A. | Kutanan_et_al_2021 | PASS |
| HM104 | HmongNjua | MSEA | Thailand | modern | 0 | Hmong-Mien | 18.12 | 98.23 | N.A. | N.A. | Kutanan_et_al_2021 | PASS |
| HM105 | HmongNjua | MSEA | Thailand | modern | 0 | Hmong-Mien | 18.12 | 98.23 | N.A. | N.A. | Kutanan_et_al_2021 | kin |
| HM107 | HmongNjua | MSEA | Thailand | modern | 0 | Hmong-Mien | 18.12 | 98.23 | N.A. | N.A. | Kutanan_et_al_2021 | PASS |
| HM108 | HmongNjua | MSEA | Thailand | modern | 0 | Hmong-Mien | 18.12 | 98.23 | N.A. | N.A. | Kutanan_et_al_2021 | PASS |
| HM109 | HmongNjua | MSEA | Thailand | modern | 0 | Hmong-Mien | 18.12 | 98.23 | N.A. | N.A. | Kutanan_et_al_2021 | PASS |
| HM110 | HmongNjua | MSEA | Thailand | modern | 0 | Hmong-Mien | 18.12 | 98.23 | N.A. | N.A. | Kutanan_et_al_2021 | kin |
| HM111 | HmongNjua | MSEA | Thailand | modern | 0 | Hmong-Mien | 18.12 | 98.23 | N.A. | N.A. | Kutanan_et_al_2021 | PASS |
| Y101 | IuMien | MSEA | Thailand | modern | 0 | Hmong-Mien | 19.34 | 100.43 | N.A. | N.A. | Kutanan_et_al_2021 | PASS |
| Y102 | IuMien | MSEA | Thailand | modern | 0 | Hmong-Mien | 19.34 | 100.43 | N.A. | N.A. | Kutanan_et_al_2021 | PASS |
| Y104 | IuMien | MSEA | Thailand | modern | 0 | Hmong-Mien | 19.34 | 100.43 | N.A. | N.A. | Kutanan_et_al_2021 | PASS |
| Y107 | IuMien | MSEA | Thailand | modern | 0 | Hmong-Mien | 19.34 | 100.43 | N.A. | N.A. | Kutanan_et_al_2021 | PASS |
| Y109 | IuMien | MSEA | Thailand | modern | 0 | Hmong-Mien | 19.34 | 100.43 | N.A. | N.A. | Kutanan_et_al_2021 | PASS |
| Y215 | IuMien | MSEA | Thailand | modern | 0 | Hmong-Mien | 19.34 | 100.43 | N.A. | N.A. | Kutanan_et_al_2021 | PASS |
| Y273 | IuMien | MSEA | Thailand | modern | 0 | Hmong-Mien | 19.34 | 100.43 | N.A. | N.A. | Kutanan_et_al_2021 | PASS |
| Y275 | IuMien | MSEA | Thailand | modern | 0 | Hmong-Mien | 19.34 | 100.43 | N.A. | N.A. | Kutanan_et_al_2021 | PASS |
| MST101 | SouthernThai_AN | MSEA | Thailand | modern | 0 | Austronesian | 6.77 | 101.08 | SouthernThai_AN | N.A. | Kutanan_et_al_2021 | PASS |
| MST104 | SouthernThai_AN | MSEA | Thailand | modern | 0 | Austronesian | 6.77 | 101.08 | SouthernThai_AN | N.A. | Kutanan_et_al_2021 | PASS |
| MST106 | SouthernThai_AN | MSEA | Thailand | modern | 0 | Austronesian | 6.77 | 101.08 | SouthernThai_AN | N.A. | Kutanan_et_al_2021 | PASS |
| MST108 | SouthernThai_AN | MSEA | Thailand | modern | 0 | Austronesian | 6.77 | 101.08 | SouthernThai_AN | N.A. | Kutanan_et_al_2021 | PASS |
| MST202 | SouthernThai_AN | MSEA | Thailand | modern | 0 | Austronesian | 6.77 | 101.08 | SouthernThai_AN | N.A. | Kutanan_et_al_2021 | PASS |
| MST204 | SouthernThai_AN | MSEA | Thailand | modern | 0 | Austronesian | 6.77 | 101.08 | SouthernThai_AN | N.A. | Kutanan_et_al_2021 | PASS |
| MST206 | SouthernThai_AN | MSEA | Thailand | modern | 0 | Austronesian | 6.77 | 101.08 | SouthernThai_AN | N.A. | Kutanan_et_al_2021 | PASS |
| MST208 | SouthernThai_AN | MSEA | Thailand | modern | 0 | Austronesian | 6.77 | 101.08 | SouthernThai_AN | N.A. | Kutanan_et_al_2021 | PASS |
| MST302 | SouthernThai_AN | MSEA | Thailand | modern | 0 | Austronesian | 6.77 | 101.08 | SouthernThai_AN | N.A. | Kutanan_et_al_2021 | PASS |
| MST304 | SouthernThai_AN | MSEA | Thailand | modern | 0 | Austronesian | 6.77 | 101.08 | SouthernThai_AN | N.A. | Kutanan_et_al_2021 | PASS |
| MST306 | SouthernThai_AN | MSEA | Thailand | modern | 0 | Austronesian | 6.77 | 101.08 | SouthernThai_AN | N.A. | Kutanan_et_al_2021 | PASS |
| MST308 | SouthernThai_AN | MSEA | Thailand | modern | 0 | Austronesian | 6.77 | 101.08 | SouthernThai_AN | N.A. | Kutanan_et_al_2021 | PASS |
| MST401 | SouthernThai_AN | MSEA | Thailand | modern | 0 | Austronesian | 6.77 | 101.08 | SouthernThai_AN | N.A. | Kutanan_et_al_2021 | PASS |
| MST402 | SouthernThai_AN | MSEA | Thailand | modern | 0 | Austronesian | 6.77 | 101.08 | SouthernThai_AN | N.A. | Kutanan_et_al_2021 | PASS |
| MST403 | SouthernThai_AN | MSEA | Thailand | modern | 0 | Austronesian | 6.77 | 101.08 | SouthernThai_AN | N.A. | Kutanan_et_al_2021 | PASS |
| LS01 | Lisu | MSEA | Thailand | modern | 0 | Sino-Tibetan | 19.5 | 98.23 | Lisu | N.A. | Kutanan_et_al_2021 | PASS |
| LS02 | Lisu | MSEA | Thailand | modern | 0 | Sino-Tibetan | 19.5 | 98.23 | Lisu | N.A. | Kutanan_et_al_2021 | PASS |
| LS03 | Lisu | MSEA | Thailand | modern | 0 | Sino-Tibetan | 19.5 | 98.23 | Lisu | N.A. | Kutanan_et_al_2021 | PASS |
| LS04 | Lisu | MSEA | Thailand | modern | 0 | Sino-Tibetan | 19.5 | 98.23 | Lisu | N.A. | Kutanan_et_al_2021 | PASS |
| LS05 | Lisu | MSEA | Thailand | modern | 0 | Sino-Tibetan | 19.5 | 98.23 | Lisu | N.A. | Kutanan_et_al_2021 | PASS |
| LS06 | Lisu | MSEA | Thailand | modern | 0 | Sino-Tibetan | 19.5 | 98.23 | Lisu | N.A. | Kutanan_et_al_2021 | PASS |

| | | | | | | | | | | | | |
|---|---|---|---|---|---|---|---|---|---|---|---|---|
| LS07 | Lisu | MSEA | Thailand | modern | 0 | Sino-Tibetan | 19.5 | 98.23 | Lisu | N.A. | Kutanan_et_al_2021 | PASS |
| LS09 | Lisu | MSEA | Thailand | modern | 0 | Sino-Tibetan | 19.5 | 98.23 | Lisu | N.A. | Kutanan_et_al_2021 | PASS |
| LS10 | Lisu | MSEA | Thailand | modern | 0 | Sino-Tibetan | 19.5 | 98.23 | Lisu | N.A. | Kutanan_et_al_2021 | PASS |
| MB01 | Lahu_T | MSEA | Thailand | modern | 0 | Sino-Tibetan | 19.58 | 98.21 | Lahu | N.A. | Kutanan_et_al_2021 | PASS |
| MB02 | Lahu_T | MSEA | Thailand | modern | 0 | Sino-Tibetan | 19.58 | 98.21 | Lahu | N.A. | Kutanan_et_al_2021 | PASS |
| MB03 | Lahu_T | MSEA | Thailand | modern | 0 | Sino-Tibetan | 19.58 | 98.21 | Lahu | N.A. | Kutanan_et_al_2021 | PASS |
| MB04 | Lahu_T | MSEA | Thailand | modern | 0 | Sino-Tibetan | 19.58 | 98.21 | Lahu | N.A. | Kutanan_et_al_2021 | PASS |
| MB05 | Lahu_T | MSEA | Thailand | modern | 0 | Sino-Tibetan | 19.58 | 98.21 | Lahu | N.A. | Kutanan_et_al_2021 | PASS |
| MB06 | Lahu_T | MSEA | Thailand | modern | 0 | Sino-Tibetan | 19.58 | 98.21 | Lahu | N.A. | Kutanan_et_al_2021 | PASS |
| MB07 | Lahu_T | MSEA | Thailand | modern | 0 | Sino-Tibetan | 19.58 | 98.21 | Lahu | N.A. | Kutanan_et_al_2021 | PASS |
| MB08 | Lahu_T | MSEA | Thailand | modern | 0 | Sino-Tibetan | 19.58 | 98.21 | Lahu | N.A. | Kutanan_et_al_2021 | PASS |
| MB09 | Lahu_T | MSEA | Thailand | modern | 0 | Sino-Tibetan | 19.58 | 98.21 | Lahu | N.A. | Kutanan_et_al_2021 | kin |
| MB10 | Lahu_T | MSEA | Thailand | modern | 0 | Sino-Tibetan | 19.58 | 98.21 | Lahu | N.A. | Kutanan_et_al_2021 | PASS |
| MR01 | Lahu_T | MSEA | Thailand | modern | 0 | Sino-Tibetan | 19.53 | 98.27 | Lahu | N.A. | Kutanan_et_al_2021 | PASS |
| MR02 | Lahu_T | MSEA | Thailand | modern | 0 | Sino-Tibetan | 19.53 | 98.27 | Lahu | N.A. | Kutanan_et_al_2021 | PASS |
| MR03 | Lahu_T | MSEA | Thailand | modern | 0 | Sino-Tibetan | 19.53 | 98.27 | Lahu | N.A. | Kutanan_et_al_2021 | PASS |
| MR05 | Lahu_T | MSEA | Thailand | modern | 0 | Sino-Tibetan | 19.53 | 98.27 | Lahu | N.A. | Kutanan_et_al_2021 | PASS |
| MR06 | Lahu_T | MSEA | Thailand | modern | 0 | Sino-Tibetan | 19.53 | 98.27 | Lahu | N.A. | Kutanan_et_al_2021 | PASS |
| MR07 | Lahu_T | MSEA | Thailand | modern | 0 | Sino-Tibetan | 19.53 | 98.27 | Lahu | N.A. | Kutanan_et_al_2021 | PASS |
| MR08 | Lahu_T | MSEA | Thailand | modern | 0 | Sino-Tibetan | 19.53 | 98.27 | Lahu | N.A. | Kutanan_et_al_2021 | PASS |
| MR09 | Lahu_T | MSEA | Thailand | modern | 0 | Sino-Tibetan | 19.53 | 98.27 | Lahu | N.A. | Kutanan_et_al_2021 | PASS |
| MR10 | Lahu_T | MSEA | Thailand | modern | 0 | Sino-Tibetan | 19.53 | 98.27 | Lahu | N.A. | Kutanan_et_al_2021 | PASS |
| MR11 | Lahu_T | MSEA | Thailand | modern | 0 | Sino-Tibetan | 19.53 | 98.27 | Lahu | N.A. | Kutanan_et_al_2021 | PASS |
| EMS301 | KarenPwo | MSEA | Thailand | modern | 0 | Sino-Tibetan | 18.15 | 97.93 | Karen_ST | N.A. | Kutanan_et_al_2021 | PASS |
| EMS305 | KarenPwo | MSEA | Thailand | modern | 0 | Sino-Tibetan | 18.15 | 97.93 | Karen_ST | N.A. | Kutanan_et_al_2021 | PASS |
| EMS306 | KarenPwo | MSEA | Thailand | modern | 0 | Sino-Tibetan | 18.15 | 97.93 | Karen_ST | N.A. | Kutanan_et_al_2021 | PASS |
| EMS308 | KarenPwo | MSEA | Thailand | modern | 0 | Sino-Tibetan | 18.15 | 97.93 | Karen_ST | N.A. | Kutanan_et_al_2021 | PASS |
| EMS309 | KarenPwo | MSEA | Thailand | modern | 0 | Sino-Tibetan | 18.15 | 97.93 | Karen_ST | N.A. | Kutanan_et_al_2021 | PASS |
| EMS313 | KarenPwo | MSEA | Thailand | modern | 0 | Sino-Tibetan | 18.15 | 97.93 | Karen_ST | N.A. | Kutanan_et_al_2021 | PASS |
| EMS314 | KarenPwo | MSEA | Thailand | modern | 0 | Sino-Tibetan | 18.15 | 97.93 | Karen_ST | N.A. | Kutanan_et_al_2021 | PASS |
| EMS316 | KarenPwo | MSEA | Thailand | modern | 0 | Sino-Tibetan | 18.15 | 97.93 | Karen_ST | N.A. | Kutanan_et_al_2021 | PASS |
| EMS318 | KarenPwo | MSEA | Thailand | modern | 0 | Sino-Tibetan | 18.15 | 97.93 | Karen_ST | N.A. | Kutanan_et_al_2021 | PASS |
| EMS322 | KarenPwo | MSEA | Thailand | modern | 0 | Sino-Tibetan | 18.15 | 97.93 | Karen_ST | N.A. | Kutanan_et_al_2021 | PASS |
| EPB501 | KarenPadaung | MSEA | Thailand | modern | 0 | Sino-Tibetan | 19.14 | 97.93 | Karen_ST | N.A. | Kutanan_et_al_2021 | PASS |
| EPB504 | KarenPadaung | MSEA | Thailand | modern | 0 | Sino-Tibetan | 19.14 | 97.93 | Karen_ST | N.A. | Kutanan_et_al_2021 | PASS |
| EPB507 | KarenPadaung | MSEA | Thailand | modern | 0 | Sino-Tibetan | 19.14 | 97.93 | Karen_ST | N.A. | Kutanan_et_al_2021 | PASS |
| EPB508 | KarenPadaung | MSEA | Thailand | modern | 0 | Sino-Tibetan | 19.14 | 97.93 | Karen_ST | N.A. | Kutanan_et_al_2021 | PASS |
| EPB511 | KarenPadaung | MSEA | Thailand | modern | 0 | Sino-Tibetan | 19.14 | 97.93 | Karen_ST | N.A. | Kutanan_et_al_2021 | PASS |
| EPB516 | KarenPadaung | MSEA | Thailand | modern | 0 | Sino-Tibetan | 19.14 | 97.93 | Karen_ST | N.A. | Kutanan_et_al_2021 | PASS |
| EPB522 | KarenPadaung | MSEA | Thailand | modern | 0 | Sino-Tibetan | 19.14 | 97.93 | Karen_ST | N.A. | Kutanan_et_al_2021 | PASS |
| EPB523 | KarenPadaung | MSEA | Thailand | modern | 0 | Sino-Tibetan | 19.14 | 97.93 | Karen_ST | N.A. | Kutanan_et_al_2021 | PASS |

| | | | | | | | | | | | | |
|---|---|---|---|---|---|---|---|---|---|---|---|---|
| EPB525 | KarenPadaung | MSEA | Thailand | modern | 0 | Sino-Tibetan | 19.14 | 97.93 | Karen_ST | N.A. | Kutanan_et_al_2021 | PASS |
| EPB528 | KarenPadaung | MSEA | Thailand | modern | 0 | Sino-Tibetan | 19.14 | 97.93 | Karen_ST | N.A. | Kutanan_et_al_2021 | PASS |
| KR104 | KarenSkaw | MSEA | Thailand | modern | 0 | Sino-Tibetan | 19.58 | 98.31 | Karen_ST | N.A. | Kutanan_et_al_2021 | PASS |
| KR105 | KarenSkaw | MSEA | Thailand | modern | 0 | Sino-Tibetan | 19.58 | 98.31 | Karen_ST | N.A. | Kutanan_et_al_2021 | PASS |
| KR106 | KarenSkaw | MSEA | Thailand | modern | 0 | Sino-Tibetan | 19.58 | 98.31 | Karen_ST | N.A. | Kutanan_et_al_2021 | PASS |
| KR107 | KarenSkaw | MSEA | Thailand | modern | 0 | Sino-Tibetan | 19.58 | 98.31 | Karen_ST | N.A. | Kutanan_et_al_2021 | PASS |
| KR108 | KarenSkaw | MSEA | Thailand | modern | 0 | Sino-Tibetan | 19.58 | 98.31 | Karen_ST | N.A. | Kutanan_et_al_2021 | PASS |
| KR111 | KarenSkaw | MSEA | Thailand | modern | 0 | Sino-Tibetan | 19.58 | 98.31 | Karen_ST | N.A. | Kutanan_et_al_2021 | PASS |
| KR113 | KarenSkaw | MSEA | Thailand | modern | 0 | Sino-Tibetan | 19.58 | 98.31 | Karen_ST | N.A. | Kutanan_et_al_2021 | PASS |
| KR115 | KarenSkaw | MSEA | Thailand | modern | 0 | Sino-Tibetan | 19.58 | 98.31 | Karen_ST | N.A. | Kutanan_et_al_2021 | PASS |
| KR117 | KarenSkaw | MSEA | Thailand | modern | 0 | Sino-Tibetan | 19.58 | 98.31 | Karen_ST | N.A. | Kutanan_et_al_2021 | PASS |
| KR122 | KarenSkaw | MSEA | Thailand | modern | 0 | Sino-Tibetan | 19.58 | 98.31 | Karen_ST | N.A. | Kutanan_et_al_2021 | PASS |
| LW601 | Lawa_Western | MSEA | Thailand | modern | 0 | Austroasiatic | 18.08 | 98.2 | N.A. | N.A. | Kutanan_et_al_2021 | PASS |
| LW603 | Lawa_Western | MSEA | Thailand | modern | 0 | Austroasiatic | 18.08 | 98.2 | N.A. | N.A. | Kutanan_et_al_2021 | PASS |
| LW609 | Lawa_Western | MSEA | Thailand | modern | 0 | Austroasiatic | 18.08 | 98.2 | N.A. | N.A. | Kutanan_et_al_2021 | PASS |
| LW611 | Lawa_Western | MSEA | Thailand | modern | 0 | Austroasiatic | 18.08 | 98.2 | N.A. | N.A. | Kutanan_et_al_2021 | PASS |
| LW612 | Lawa_Western | MSEA | Thailand | modern | 0 | Austroasiatic | 18.08 | 98.2 | N.A. | N.A. | Kutanan_et_al_2021 | PASS |
| LW615 | Lawa_Western | MSEA | Thailand | modern | 0 | Austroasiatic | 18.08 | 98.2 | N.A. | N.A. | Kutanan_et_al_2021 | PASS |
| LW617 | Lawa_Western | MSEA | Thailand | modern | 0 | Austroasiatic | 18.08 | 98.2 | N.A. | N.A. | Kutanan_et_al_2021 | PASS |
| LW619 | Lawa_Western | MSEA | Thailand | modern | 0 | Austroasiatic | 18.08 | 98.2 | N.A. | N.A. | Kutanan_et_al_2021 | PASS |
| LW625 | Lawa_Western | MSEA | Thailand | modern | 0 | Austroasiatic | 18.08 | 98.2 | N.A. | N.A. | Kutanan_et_al_2021 | PASS |
| LW502 | Lawa_Eastern | MSEA | Thailand | modern | 0 | Austroasiatic | 18.23 | 97.56 | N.A. | N.A. | Kutanan_et_al_2021 | PASS |
| LW504 | Lawa_Eastern | MSEA | Thailand | modern | 0 | Austroasiatic | 18.23 | 97.56 | N.A. | N.A. | Kutanan_et_al_2021 | PASS |
| LW506 | Lawa_Eastern | MSEA | Thailand | modern | 0 | Austroasiatic | 18.23 | 97.56 | N.A. | N.A. | Kutanan_et_al_2021 | PASS |
| LW510 | Lawa_Eastern | MSEA | Thailand | modern | 0 | Austroasiatic | 18.23 | 97.56 | N.A. | N.A. | Kutanan_et_al_2021 | PASS |
| LW514 | Lawa_Eastern | MSEA | Thailand | modern | 0 | Austroasiatic | 18.23 | 97.56 | N.A. | N.A. | Kutanan_et_al_2021 | PASS |
| LW520 | Lawa_Eastern | MSEA | Thailand | modern | 0 | Austroasiatic | 18.23 | 97.56 | N.A. | N.A. | Kutanan_et_al_2021 | PASS |
| LW522 | Lawa_Eastern | MSEA | Thailand | modern | 0 | Austroasiatic | 18.23 | 97.56 | N.A. | N.A. | Kutanan_et_al_2021 | PASS |
| LW524 | Lawa_Eastern | MSEA | Thailand | modern | 0 | Austroasiatic | 18.23 | 97.56 | N.A. | N.A. | Kutanan_et_al_2021 | PASS |
| LW526 | Lawa_Eastern | MSEA | Thailand | modern | 0 | Austroasiatic | 18.23 | 97.56 | N.A. | N.A. | Kutanan_et_al_2021 | PASS |
| LW528 | Lawa_Eastern | MSEA | Thailand | modern | 0 | Austroasiatic | 18.23 | 97.56 | N.A. | N.A. | Kutanan_et_al_2021 | PASS |
| PL113 | Palaung | MSEA | Thailand | modern | 0 | Austroasiatic | 20.25 | 99.52 | N.A. | N.A. | Kutanan_et_al_2021 | PASS |
| PL117 | Palaung | MSEA | Thailand | modern | 0 | Austroasiatic | 20.25 | 99.52 | N.A. | N.A. | Kutanan_et_al_2021 | PASS |
| PL119 | Palaung | MSEA | Thailand | modern | 0 | Austroasiatic | 20.25 | 99.52 | N.A. | N.A. | Kutanan_et_al_2021 | PASS |
| PL121 | Palaung | MSEA | Thailand | modern | 0 | Austroasiatic | 20.25 | 99.52 | N.A. | N.A. | Kutanan_et_al_2021 | kin |
| PL123 | Palaung | MSEA | Thailand | modern | 0 | Austroasiatic | 20.25 | 99.52 | N.A. | N.A. | Kutanan_et_al_2021 | PASS |
| PL127 | Palaung | MSEA | Thailand | modern | 0 | Austroasiatic | 20.25 | 99.52 | N.A. | N.A. | Kutanan_et_al_2021 | PASS |
| PL129 | Palaung | MSEA | Thailand | modern | 0 | Austroasiatic | 20.25 | 99.52 | N.A. | N.A. | Kutanan_et_al_2021 | PASS |
| PL133 | Palaung | MSEA | Thailand | modern | 0 | Austroasiatic | 20.25 | 99.52 | N.A. | N.A. | Kutanan_et_al_2021 | PASS |
| PL134 | Palaung | MSEA | Thailand | modern | 0 | Austroasiatic | 20.25 | 99.52 | N.A. | N.A. | Kutanan_et_al_2021 | PASS |
| PL145 | Palaung | MSEA | Thailand | modern | 0 | Austroasiatic | 20.25 | 99.52 | N.A. | N.A. | Kutanan_et_al_2021 | PASS |

| | | | | | | | | | | | | |
|---|---|---|---|---|---|---|---|---|---|---|---|---|
| PP101 | Blang | MSEA | Thailand | modern | 0 | Austroasiatic | 19.56 | 99.09 | N.A. | N.A. | Kutanan_et_al_2021 | PASS |
| PP103 | Blang | MSEA | Thailand | modern | 0 | Austroasiatic | 19.56 | 99.09 | N.A. | N.A. | Kutanan_et_al_2021 | PASS |
| PP105 | Blang | MSEA | Thailand | modern | 0 | Austroasiatic | 19.56 | 99.09 | N.A. | N.A. | Kutanan_et_al_2021 | PASS |
| PP109 | Blang | MSEA | Thailand | modern | 0 | Austroasiatic | 19.56 | 99.09 | N.A. | N.A. | Kutanan_et_al_2021 | kin |
| PP119 | Blang | MSEA | Thailand | modern | 0 | Austroasiatic | 19.56 | 99.09 | N.A. | N.A. | Kutanan_et_al_2021 | PASS |
| PP211 | Blang | MSEA | Thailand | modern | 0 | Austroasiatic | 19.56 | 99.09 | N.A. | N.A. | Kutanan_et_al_2021 | PASS |
| PP257 | Blang | MSEA | Thailand | modern | 0 | Austroasiatic | 19.56 | 99.09 | N.A. | N.A. | Kutanan_et_al_2021 | PASS |
| PP259 | Blang | MSEA | Thailand | modern | 0 | Austroasiatic | 19.56 | 99.09 | N.A. | N.A. | Kutanan_et_al_2021 | PASS |
| PP263 | Blang | MSEA | Thailand | modern | 0 | Austroasiatic | 19.56 | 99.09 | N.A. | N.A. | Kutanan_et_al_2021 | PASS |
| PP265 | Blang | MSEA | Thailand | modern | 0 | Austroasiatic | 19.56 | 99.09 | N.A. | N.A. | Kutanan_et_al_2021 | PASS |
| KM102 | Khmu | MSEA | Thailand | modern | 0 | Austroasiatic | 19.49 | 100.89 | N.A. | N.A. | Kutanan_et_al_2021 | PASS |
| KM110 | Khmu | MSEA | Thailand | modern | 0 | Austroasiatic | 19.49 | 100.89 | N.A. | N.A. | Kutanan_et_al_2021 | PASS |
| KM115 | Khmu | MSEA | Thailand | modern | 0 | Austroasiatic | 19.49 | 100.89 | N.A. | N.A. | Kutanan_et_al_2021 | PASS |
| KM117 | Khmu | MSEA | Thailand | modern | 0 | Austroasiatic | 19.49 | 100.89 | N.A. | N.A. | Kutanan_et_al_2021 | PASS |
| KM119 | Khmu | MSEA | Thailand | modern | 0 | Austroasiatic | 19.49 | 100.89 | N.A. | N.A. | Kutanan_et_al_2021 | PASS |
| KM120 | Khmu | MSEA | Thailand | modern | 0 | Austroasiatic | 19.49 | 100.89 | N.A. | N.A. | Kutanan_et_al_2021 | PASS |
| KM121 | Khmu | MSEA | Thailand | modern | 0 | Austroasiatic | 19.49 | 100.89 | N.A. | N.A. | Kutanan_et_al_2021 | PASS |
| KM129 | Khmu | MSEA | Thailand | modern | 0 | Austroasiatic | 19.49 | 100.89 | N.A. | N.A. | Kutanan_et_al_2021 | PASS |
| KM133 | Khmu | MSEA | Thailand | modern | 0 | Austroasiatic | 19.49 | 100.89 | N.A. | N.A. | Kutanan_et_al_2021 | PASS |
| KM137 | Khmu | MSEA | Thailand | modern | 0 | Austroasiatic | 19.49 | 100.89 | N.A. | N.A. | Kutanan_et_al_2021 | PASS |
| TN201 | HtinPray | MSEA | Thailand | modern | 0 | Austroasiatic | 19.23 | 100.52 | N.A. | N.A. | Kutanan_et_al_2021 | PASS |
| TN207 | HtinPray | MSEA | Thailand | modern | 0 | Austroasiatic | 19.23 | 100.52 | N.A. | N.A. | Kutanan_et_al_2021 | PASS |
| TN209 | HtinPray | MSEA | Thailand | modern | 0 | Austroasiatic | 19.23 | 100.52 | N.A. | N.A. | Kutanan_et_al_2021 | PASS |
| TN213 | HtinPray | MSEA | Thailand | modern | 0 | Austroasiatic | 19.23 | 100.52 | N.A. | N.A. | Kutanan_et_al_2021 | PASS |
| TN219 | HtinPray | MSEA | Thailand | modern | 0 | Austroasiatic | 19.23 | 100.52 | N.A. | N.A. | Kutanan_et_al_2021 | kin |
| TN223 | HtinPray | MSEA | Thailand | modern | 0 | Austroasiatic | 19.23 | 100.52 | N.A. | N.A. | Kutanan_et_al_2021 | kin |
| TN227 | HtinPray | MSEA | Thailand | modern | 0 | Austroasiatic | 19.23 | 100.52 | N.A. | N.A. | Kutanan_et_al_2021 | PASS |
| TN229 | HtinPray | MSEA | Thailand | modern | 0 | Austroasiatic | 19.23 | 100.52 | N.A. | N.A. | Kutanan_et_al_2021 | PASS |
| TN231 | HtinPray | MSEA | Thailand | modern | 0 | Austroasiatic | 19.23 | 100.52 | N.A. | N.A. | Kutanan_et_al_2021 | PASS |
| TN234 | HtinPray | MSEA | Thailand | modern | 0 | Austroasiatic | 19.23 | 100.52 | N.A. | N.A. | Kutanan_et_al_2021 | kin |
| TN236 | HtinPray | MSEA | Thailand | modern | 0 | Austroasiatic | 19.23 | 100.52 | N.A. | N.A. | Kutanan_et_al_2021 | kin |
| TN301 | HtinPray | MSEA | Thailand | modern | 0 | Austroasiatic | 19.23 | 100.52 | N.A. | N.A. | Kutanan_et_al_2021 | PASS |
| TN303 | HtinPray | MSEA | Thailand | modern | 0 | Austroasiatic | 19.23 | 100.52 | N.A. | N.A. | Kutanan_et_al_2021 | PASS |
| TN307 | HtinPray | MSEA | Thailand | modern | 0 | Austroasiatic | 19.23 | 100.52 | N.A. | N.A. | Kutanan_et_al_2021 | PASS |
| TN313 | HtinPray | MSEA | Thailand | modern | 0 | Austroasiatic | 19.23 | 100.52 | N.A. | N.A. | Kutanan_et_al_2021 | PASS |
| TN102 | HtinMal | MSEA | Thailand | modern | 0 | Austroasiatic | 19.08 | 100.55 | N.A. | N.A. | Lipson_et_al_2018 | PASS |
| TN105 | HtinMal | MSEA | Thailand | modern | 0 | Austroasiatic | 19.08 | 100.55 | N.A. | N.A. | Lipson_et_al_2018 | PASS |
| TN106 | HtinMal | MSEA | Thailand | modern | 0 | Austroasiatic | 19.08 | 100.55 | N.A. | N.A. | Lipson_et_al_2018 | PASS |
| TN110 | HtinMal | MSEA | Thailand | modern | 0 | Austroasiatic | 19.08 | 100.55 | N.A. | N.A. | Lipson_et_al_2018 | PASS |
| TN112 | HtinMal | MSEA | Thailand | modern | 0 | Austroasiatic | 19.08 | 100.55 | N.A. | N.A. | Lipson_et_al_2018 | PASS |
| TN115 | HtinMal | MSEA | Thailand | modern | 0 | Austroasiatic | 19.08 | 100.55 | N.A. | N.A. | Lipson_et_al_2018 | PASS |

| | | | | | | | | | | | | |
|---|---|---|---|---|---|---|---|---|---|---|---|---|
| TN122 | HtinMal | MSEA | Thailand | modern | 0 | Austroasiatic | 19.08 | 100.55 | N.A. | N.A. | Lipson_et_al_2018 | PASS |
| TN123 | HtinMal | MSEA | Thailand | modern | 0 | Austroasiatic | 19.08 | 100.55 | N.A. | N.A. | Lipson_et_al_2018 | PASS |
| TN125 | HtinMal | MSEA | Thailand | modern | 0 | Austroasiatic | 19.08 | 100.55 | N.A. | N.A. | Lipson_et_al_2018 | PASS |
| TN147 | HtinMal | MSEA | Thailand | modern | 0 | Austroasiatic | 19.08 | 100.55 | N.A. | N.A. | Lipson_et_al_2018 | PASS |
| MA107 | Mlabri | MSEA | Thailand | modern | 0 | Austroasiatic | 18.61 | 100.71 | N.A. | N.A. | Lipson_et_al_2018 | PASS |
| MA112 | Mlabri | MSEA | Thailand | modern | 0 | Austroasiatic | 18.61 | 100.71 | N.A. | N.A. | Lipson_et_al_2018 | PASS |
| MA113 | Mlabri | MSEA | Thailand | modern | 0 | Austroasiatic | 18.61 | 100.71 | N.A. | N.A. | Lipson_et_al_2018 | PASS |
| MA115 | Mlabri | MSEA | Thailand | modern | 0 | Austroasiatic | 18.61 | 100.71 | N.A. | N.A. | Lipson_et_al_2018 | PASS |
| MA117 | Mlabri | MSEA | Thailand | modern | 0 | Austroasiatic | 18.61 | 100.71 | N.A. | N.A. | Lipson_et_al_2018 | PASS |
| MA119 | Mlabri | MSEA | Thailand | modern | 0 | Austroasiatic | 18.61 | 100.71 | N.A. | N.A. | Lipson_et_al_2018 | PASS |
| MA122 | Mlabri | MSEA | Thailand | modern | 0 | Austroasiatic | 18.61 | 100.71 | N.A. | N.A. | Lipson_et_al_2018 | PASS |
| MA124 | Mlabri | MSEA | Thailand | modern | 0 | Austroasiatic | 18.61 | 100.71 | N.A. | N.A. | Lipson_et_al_2018 | PASS |
| MA126 | Mlabri | MSEA | Thailand | modern | 0 | Austroasiatic | 18.61 | 100.71 | N.A. | N.A. | Lipson_et_al_2018 | PASS |
| MA128 | Mlabri | MSEA | Thailand | modern | 0 | Austroasiatic | 18.61 | 100.71 | N.A. | N.A. | Lipson_et_al_2018 | kin |
| SOA101 | Soa | MSEA | Thailand | modern | 0 | Austroasiatic | 17.37 | 104.3 | N.A. | N.A. | Kutanan_et_al_2021 | PASS |
| SOA105 | Soa | MSEA | Thailand | modern | 0 | Austroasiatic | 17.37 | 104.3 | N.A. | N.A. | Kutanan_et_al_2021 | PASS |
| SOA107 | Soa | MSEA | Thailand | modern | 0 | Austroasiatic | 17.37 | 104.3 | N.A. | N.A. | Kutanan_et_al_2021 | PASS |
| SOA109 | Soa | MSEA | Thailand | modern | 0 | Austroasiatic | 17.37 | 104.3 | N.A. | N.A. | Kutanan_et_al_2021 | PASS |
| SOA111 | Soa | MSEA | Thailand | modern | 0 | Austroasiatic | 17.37 | 104.3 | N.A. | N.A. | Kutanan_et_al_2021 | PASS |
| SOA113 | Soa | MSEA | Thailand | modern | 0 | Austroasiatic | 17.37 | 104.3 | N.A. | N.A. | Kutanan_et_al_2021 | PASS |
| SOA115 | Soa | MSEA | Thailand | modern | 0 | Austroasiatic | 17.37 | 104.3 | N.A. | N.A. | Kutanan_et_al_2021 | PASS |
| SOA117 | Soa | MSEA | Thailand | modern | 0 | Austroasiatic | 17.37 | 104.3 | N.A. | N.A. | Kutanan_et_al_2021 | PASS |
| SOA119 | Soa | MSEA | Thailand | modern | 0 | Austroasiatic | 17.37 | 104.3 | N.A. | N.A. | Kutanan_et_al_2021 | PASS |
| SOA121 | Soa | MSEA | Thailand | modern | 0 | Austroasiatic | 17.37 | 104.3 | N.A. | N.A. | Kutanan_et_al_2021 | PASS |
| SOA123 | Soa | MSEA | Thailand | modern | 0 | Austroasiatic | 17.37 | 104.3 | N.A. | N.A. | Kutanan_et_al_2021 | PASS |
| SOA125 | Soa | MSEA | Thailand | modern | 0 | Austroasiatic | 17.37 | 104.3 | N.A. | N.A. | Kutanan_et_al_2021 | PASS |
| SOA127 | Soa | MSEA | Thailand | modern | 0 | Austroasiatic | 17.37 | 104.3 | N.A. | N.A. | Kutanan_et_al_2021 | PASS |
| SOA129 | Soa | MSEA | Thailand | modern | 0 | Austroasiatic | 17.37 | 104.3 | N.A. | N.A. | Kutanan_et_al_2021 | PASS |
| SOA137 | Soa | MSEA | Thailand | modern | 0 | Austroasiatic | 17.37 | 104.3 | N.A. | N.A. | Kutanan_et_al_2021 | PASS |
| BRU101 | Bru | MSEA | Thailand | modern | 0 | Austroasiatic | 17.39 | 103.7 | N.A. | N.A. | Kutanan_et_al_2021 | PASS |
| BRU103 | Bru | MSEA | Thailand | modern | 0 | Austroasiatic | 17.39 | 103.7 | N.A. | N.A. | Kutanan_et_al_2021 | PASS |
| BRU105 | Bru | MSEA | Thailand | modern | 0 | Austroasiatic | 17.39 | 103.7 | N.A. | N.A. | Kutanan_et_al_2021 | kin |
| BRU107 | Bru | MSEA | Thailand | modern | 0 | Austroasiatic | 17.39 | 103.7 | N.A. | N.A. | Kutanan_et_al_2021 | PASS |
| BRU109 | Bru | MSEA | Thailand | modern | 0 | Austroasiatic | 17.39 | 103.7 | N.A. | N.A. | Kutanan_et_al_2021 | kin |
| BRU111 | Bru | MSEA | Thailand | modern | 0 | Austroasiatic | 17.39 | 103.7 | N.A. | N.A. | Kutanan_et_al_2021 | PASS |
| BRU113 | Bru | MSEA | Thailand | modern | 0 | Austroasiatic | 17.39 | 103.7 | N.A. | N.A. | Kutanan_et_al_2021 | kin |
| BRU115 | Bru | MSEA | Thailand | modern | 0 | Austroasiatic | 17.39 | 103.7 | N.A. | N.A. | Kutanan_et_al_2021 | PASS |
| BRU117 | Bru | MSEA | Thailand | modern | 0 | Austroasiatic | 17.39 | 103.7 | N.A. | N.A. | Kutanan_et_al_2021 | PASS |
| BRU119 | Bru | MSEA | Thailand | modern | 0 | Austroasiatic | 17.39 | 103.7 | N.A. | N.A. | Kutanan_et_al_2021 | PASS |
| BRU121 | Bru | MSEA | Thailand | modern | 0 | Austroasiatic | 17.39 | 103.7 | N.A. | N.A. | Kutanan_et_al_2021 | PASS |
| BRU123 | Bru | MSEA | Thailand | modern | 0 | Austroasiatic | 17.39 | 103.7 | N.A. | N.A. | Kutanan_et_al_2021 | PASS |

| | | | | | | | | | | | | |
|---|---|---|---|---|---|---|---|---|---|---|---|---|
| BRU125 | Bru | MSEA | Thailand | modern | 0 | Austroasiatic | 17.39 | 103.7 | N.A. | N.A. | Kutanan_et_al_2021 | PASS |
| BRU127 | Bru | MSEA | Thailand | modern | 0 | Austroasiatic | 17.39 | 103.7 | N.A. | N.A. | Kutanan_et_al_2021 | PASS |
| BRU129 | Bru | MSEA | Thailand | modern | 0 | Austroasiatic | 17.39 | 103.7 | N.A. | N.A. | Kutanan_et_al_2021 | PASS |
| MO103 | Mon | MSEA | Thailand | modern | 0 | Austroasiatic | 14.69 | 102.06 | N.A. | N.A. | Kutanan_et_al_2021 | PASS |
| MO105 | Mon | MSEA | Thailand | modern | 0 | Austroasiatic | 14.69 | 102.06 | N.A. | N.A. | Kutanan_et_al_2021 | PASS |
| MO109 | Mon | MSEA | Thailand | modern | 0 | Austroasiatic | 14.69 | 102.06 | N.A. | N.A. | Kutanan_et_al_2021 | PASS |
| MO113 | Mon | MSEA | Thailand | modern | 0 | Austroasiatic | 14.69 | 102.06 | N.A. | N.A. | Kutanan_et_al_2021 | PASS |
| MO115 | Mon | MSEA | Thailand | modern | 0 | Austroasiatic | 14.69 | 102.06 | N.A. | N.A. | Kutanan_et_al_2021 | PASS |
| MO117 | Mon | MSEA | Thailand | modern | 0 | Austroasiatic | 14.69 | 102.06 | N.A. | N.A. | Kutanan_et_al_2021 | PASS |
| MO119 | Mon | MSEA | Thailand | modern | 0 | Austroasiatic | 14.69 | 102.06 | N.A. | N.A. | Kutanan_et_al_2021 | PASS |
| MO121 | Mon | MSEA | Thailand | modern | 0 | Austroasiatic | 14.69 | 102.06 | N.A. | N.A. | Kutanan_et_al_2021 | PASS |
| MO125 | Mon | MSEA | Thailand | modern | 0 | Austroasiatic | 14.69 | 102.06 | N.A. | N.A. | Kutanan_et_al_2021 | PASS |
| MO127 | Mon | MSEA | Thailand | modern | 0 | Austroasiatic | 14.69 | 102.06 | N.A. | N.A. | Kutanan_et_al_2021 | PASS |
| MO129 | Mon | MSEA | Thailand | modern | 0 | Austroasiatic | 14.69 | 102.06 | N.A. | N.A. | Kutanan_et_al_2021 | PASS |
| MO135 | Mon | MSEA | Thailand | modern | 0 | Austroasiatic | 14.69 | 102.06 | N.A. | N.A. | Kutanan_et_al_2021 | PASS |
| MO139 | Mon | MSEA | Thailand | modern | 0 | Austroasiatic | 14.69 | 102.06 | N.A. | N.A. | Kutanan_et_al_2021 | PASS |
| MO141 | Mon | MSEA | Thailand | modern | 0 | Austroasiatic | 14.69 | 102.06 | N.A. | N.A. | Kutanan_et_al_2021 | PASS |
| MO143 | Mon | MSEA | Thailand | modern | 0 | Austroasiatic | 14.69 | 102.06 | N.A. | N.A. | Kutanan_et_al_2021 | PASS |
| MO301 | Mon | MSEA | Thailand | modern | 0 | Austroasiatic | 15.3 | 98.38 | N.A. | N.A. | Kutanan_et_al_2021 | PASS |
| MO303 | Mon | MSEA | Thailand | modern | 0 | Austroasiatic | 15.3 | 98.38 | N.A. | N.A. | Kutanan_et_al_2021 | PASS |
| MO305 | Mon | MSEA | Thailand | modern | 0 | Austroasiatic | 15.3 | 98.38 | N.A. | N.A. | Kutanan_et_al_2021 | PASS |
| MO307 | Mon | MSEA | Thailand | modern | 0 | Austroasiatic | 15.3 | 98.38 | N.A. | N.A. | Kutanan_et_al_2021 | PASS |
| MO309 | Mon | MSEA | Thailand | modern | 0 | Austroasiatic | 15.3 | 98.38 | N.A. | N.A. | Kutanan_et_al_2021 | PASS |
| MO311 | Mon | MSEA | Thailand | modern | 0 | Austroasiatic | 15.3 | 98.38 | N.A. | N.A. | Kutanan_et_al_2021 | PASS |
| MO313 | Mon | MSEA | Thailand | modern | 0 | Austroasiatic | 15.3 | 98.38 | N.A. | N.A. | Kutanan_et_al_2021 | PASS |
| MO315 | Mon | MSEA | Thailand | modern | 0 | Austroasiatic | 15.3 | 98.38 | N.A. | N.A. | Kutanan_et_al_2021 | PASS |
| MO317 | Mon | MSEA | Thailand | modern | 0 | Austroasiatic | 15.3 | 98.38 | N.A. | N.A. | Kutanan_et_al_2021 | PASS |
| MO319 | Mon | MSEA | Thailand | modern | 0 | Austroasiatic | 15.3 | 98.38 | N.A. | N.A. | Kutanan_et_al_2021 | PASS |
| MO701 | Mon | MSEA | Thailand | modern | 0 | Austroasiatic | 13.58 | 100.1 | N.A. | N.A. | Kutanan_et_al_2021 | PASS |
| MO703 | Mon | MSEA | Thailand | modern | 0 | Austroasiatic | 13.58 | 100.1 | N.A. | N.A. | Kutanan_et_al_2021 | PASS |
| MO705 | Mon | MSEA | Thailand | modern | 0 | Austroasiatic | 13.58 | 100.1 | N.A. | N.A. | Kutanan_et_al_2021 | PASS |
| MO707 | Mon | MSEA | Thailand | modern | 0 | Austroasiatic | 13.58 | 100.1 | N.A. | N.A. | Kutanan_et_al_2021 | PASS |
| MO709 | Mon | MSEA | Thailand | modern | 0 | Austroasiatic | 13.58 | 100.1 | N.A. | N.A. | Kutanan_et_al_2021 | PASS |
| MO711 | Mon | MSEA | Thailand | modern | 0 | Austroasiatic | 13.58 | 100.1 | N.A. | N.A. | Kutanan_et_al_2021 | PASS |
| MO713 | Mon | MSEA | Thailand | modern | 0 | Austroasiatic | 13.58 | 100.1 | N.A. | N.A. | Kutanan_et_al_2021 | PASS |
| MO715 | Mon | MSEA | Thailand | modern | 0 | Austroasiatic | 13.58 | 100.1 | N.A. | N.A. | Kutanan_et_al_2021 | PASS |
| MO717 | Mon | MSEA | Thailand | modern | 0 | Austroasiatic | 13.58 | 100.1 | N.A. | N.A. | Kutanan_et_al_2021 | kin |
| MO719 | Mon | MSEA | Thailand | modern | 0 | Austroasiatic | 13.58 | 100.1 | N.A. | N.A. | Kutanan_et_al_2021 | PASS |
| RM101 | Mon | MSEA | Thailand | modern | 0 | Austroasiatic | 13.77 | 99.85 | N.A. | N.A. | Kutanan_et_al_2021 | PASS |
| RM103 | Mon | MSEA | Thailand | modern | 0 | Austroasiatic | 13.77 | 99.85 | N.A. | N.A. | Kutanan_et_al_2021 | PASS |
| RM105 | Mon | MSEA | Thailand | modern | 0 | Austroasiatic | 13.77 | 99.85 | N.A. | N.A. | Kutanan_et_al_2021 | PASS |

| | | | | | | | | | | | | |
|---|---|---|---|---|---|---|---|---|---|---|---|---|
| RM107 | Mon | MSEA | Thailand | modern | 0 | Austroasiatic | 13.77 | 99.85 | N.A. | N.A. | Kutanan_et_al_2021 | PASS |
| RM109 | Mon | MSEA | Thailand | modern | 0 | Austroasiatic | 13.77 | 99.85 | N.A. | N.A. | Kutanan_et_al_2021 | PASS |
| RM110 | Mon | MSEA | Thailand | modern | 0 | Austroasiatic | 13.77 | 99.85 | N.A. | N.A. | Kutanan_et_al_2021 | PASS |
| RM111 | Mon | MSEA | Thailand | modern | 0 | Austroasiatic | 13.77 | 99.85 | N.A. | N.A. | Kutanan_et_al_2021 | PASS |
| RM112 | Mon | MSEA | Thailand | modern | 0 | Austroasiatic | 13.77 | 99.85 | N.A. | N.A. | Kutanan_et_al_2021 | PASS |
| RM115 | Mon | MSEA | Thailand | modern | 0 | Austroasiatic | 13.77 | 99.85 | N.A. | N.A. | Kutanan_et_al_2021 | PASS |
| RM117 | Mon | MSEA | Thailand | modern | 0 | Austroasiatic | 13.77 | 99.85 | N.A. | N.A. | Kutanan_et_al_2021 | PASS |
| RM119 | Mon | MSEA | Thailand | modern | 0 | Austroasiatic | 13.77 | 99.85 | N.A. | N.A. | Kutanan_et_al_2021 | PASS |
| RM121 | Mon | MSEA | Thailand | modern | 0 | Austroasiatic | 13.77 | 99.85 | N.A. | N.A. | Kutanan_et_al_2021 | PASS |
| RM123 | Mon | MSEA | Thailand | modern | 0 | Austroasiatic | 13.77 | 99.85 | N.A. | N.A. | Kutanan_et_al_2021 | PASS |
| RM127 | Mon | MSEA | Thailand | modern | 0 | Austroasiatic | 13.77 | 99.85 | N.A. | N.A. | Kutanan_et_al_2021 | PASS |
| RM129 | Mon | MSEA | Thailand | modern | 0 | Austroasiatic | 13.77 | 99.85 | N.A. | N.A. | Kutanan_et_al_2021 | PASS |
| RY112 | Yuan | MSEA | Thailand | modern | 0 | Tai-Kadai | 13.54 | 99.83 | N_TK | N.A. | Kutanan_et_al_2021 | PASS |
| RY130 | Yuan | MSEA | Thailand | modern | 0 | Tai-Kadai | 13.54 | 99.83 | N_TK | N.A. | Kutanan_et_al_2021 | PASS |
| RY132 | Yuan | MSEA | Thailand | modern | 0 | Tai-Kadai | 13.54 | 99.83 | N_TK | N.A. | Kutanan_et_al_2021 | PASS |
| RY136 | Yuan | MSEA | Thailand | modern | 0 | Tai-Kadai | 13.54 | 99.83 | N_TK | N.A. | Kutanan_et_al_2021 | PASS |
| RY137 | Yuan | MSEA | Thailand | modern | 0 | Tai-Kadai | 13.54 | 99.83 | N_TK | N.A. | Kutanan_et_al_2021 | PASS |
| RY139 | Yuan | MSEA | Thailand | modern | 0 | Tai-Kadai | 13.54 | 99.83 | N_TK | N.A. | Kutanan_et_al_2021 | PASS |
| RY141 | Yuan | MSEA | Thailand | modern | 0 | Tai-Kadai | 13.54 | 99.83 | N_TK | N.A. | Kutanan_et_al_2021 | PASS |
| RY143 | Yuan | MSEA | Thailand | modern | 0 | Tai-Kadai | 13.54 | 99.83 | N_TK | N.A. | Kutanan_et_al_2021 | PASS |
| TK102 | Khuen | MSEA | Thailand | modern | 0 | Tai-Kadai | 18.38 | 98.51 | N_TK | N.A. | Kutanan_et_al_2021 | PASS |
| TK104 | Khuen | MSEA | Thailand | modern | 0 | Tai-Kadai | 18.38 | 98.51 | N_TK | N.A. | Kutanan_et_al_2021 | PASS |
| TK112 | Khuen | MSEA | Thailand | modern | 0 | Tai-Kadai | 18.38 | 98.51 | N_TK | N.A. | Kutanan_et_al_2021 | PASS |
| TK114 | Khuen | MSEA | Thailand | modern | 0 | Tai-Kadai | 18.38 | 98.51 | N_TK | N.A. | Kutanan_et_al_2021 | kin |
| TK118 | Khuen | MSEA | Thailand | modern | 0 | Tai-Kadai | 18.38 | 98.51 | N_TK | N.A. | Kutanan_et_al_2021 | PASS |
| TK119 | Khuen | MSEA | Thailand | modern | 0 | Tai-Kadai | 18.38 | 98.51 | N_TK | N.A. | Kutanan_et_al_2021 | PASS |
| TK120 | Khuen | MSEA | Thailand | modern | 0 | Tai-Kadai | 18.38 | 98.51 | N_TK | N.A. | Kutanan_et_al_2021 | PASS |
| TK121 | Khuen | MSEA | Thailand | modern | 0 | Tai-Kadai | 18.38 | 98.51 | N_TK | N.A. | Kutanan_et_al_2021 | PASS |
| TK127 | Khuen | MSEA | Thailand | modern | 0 | Tai-Kadai | 18.38 | 98.51 | N_TK | N.A. | Kutanan_et_al_2021 | PASS |
| TK139 | Khuen | MSEA | Thailand | modern | 0 | Tai-Kadai | 18.38 | 98.51 | N_TK | N.A. | Kutanan_et_al_2021 | PASS |
| STP101 | Phuan | MSEA | Thailand | modern | 0 | Tai-Kadai | 17.51 | 99.77 | N_TK | N.A. | Kutanan_et_al_2021 | PASS |
| STP103 | Phuan | MSEA | Thailand | modern | 0 | Tai-Kadai | 17.51 | 99.77 | N_TK | N.A. | Kutanan_et_al_2021 | PASS |
| STP106 | Phuan | MSEA | Thailand | modern | 0 | Tai-Kadai | 17.51 | 99.77 | N_TK | N.A. | Kutanan_et_al_2021 | PASS |
| STP107 | Phuan | MSEA | Thailand | modern | 0 | Tai-Kadai | 17.51 | 99.77 | N_TK | N.A. | Kutanan_et_al_2021 | PASS |
| STP111 | Phuan | MSEA | Thailand | modern | 0 | Tai-Kadai | 17.51 | 99.77 | N_TK | N.A. | Kutanan_et_al_2021 | PASS |
| STP113 | Phuan | MSEA | Thailand | modern | 0 | Tai-Kadai | 17.51 | 99.77 | N_TK | N.A. | Kutanan_et_al_2021 | PASS |
| STP115 | Phuan | MSEA | Thailand | modern | 0 | Tai-Kadai | 17.51 | 99.77 | N_TK | N.A. | Kutanan_et_al_2021 | PASS |
| STP117 | Phuan | MSEA | Thailand | modern | 0 | Tai-Kadai | 17.51 | 99.77 | N_TK | N.A. | Kutanan_et_al_2021 | PASS |
| STP121 | Phuan | MSEA | Thailand | modern | 0 | Tai-Kadai | 17.51 | 99.77 | N_TK | N.A. | Kutanan_et_al_2021 | PASS |
| STP123 | Phuan | MSEA | Thailand | modern | 0 | Tai-Kadai | 17.51 | 99.77 | N_TK | N.A. | Kutanan_et_al_2021 | PASS |
| YA102 | Shan | MSEA | Thailand | modern | 0 | Tai-Kadai | 19.37 | 98.14 | N_TK | N.A. | Kutanan_et_al_2021 | PASS |

| | | | | | | | | | | | | |
|---|---|---|---|---|---|---|---|---|---|---|---|---|
| YA107 | Shan | MSEA | Thailand | modern | 0 | Tai-Kadai | 19.37 | 98.14 | N_TK | N.A. | Kutanan_et_al_2021 | PASS |
| YA108 | Shan | MSEA | Thailand | modern | 0 | Tai-Kadai | 19.37 | 98.14 | N_TK | N.A. | Kutanan_et_al_2021 | PASS |
| YA111 | Shan | MSEA | Thailand | modern | 0 | Tai-Kadai | 19.37 | 98.14 | N_TK | N.A. | Kutanan_et_al_2021 | PASS |
| YA113 | Shan | MSEA | Thailand | modern | 0 | Tai-Kadai | 19.37 | 98.14 | N_TK | N.A. | Kutanan_et_al_2021 | PASS |
| YA115 | Shan | MSEA | Thailand | modern | 0 | Tai-Kadai | 19.37 | 98.14 | N_TK | N.A. | Kutanan_et_al_2021 | PASS |
| YA117 | Shan | MSEA | Thailand | modern | 0 | Tai-Kadai | 19.37 | 98.14 | N_TK | N.A. | Kutanan_et_al_2021 | PASS |
| YA118 | Shan | MSEA | Thailand | modern | 0 | Tai-Kadai | 19.37 | 98.14 | N_TK | N.A. | Kutanan_et_al_2021 | kin |
| YA121 | Shan | MSEA | Thailand | modern | 0 | Tai-Kadai | 19.37 | 98.14 | N_TK | N.A. | Kutanan_et_al_2021 | PASS |
| YA122 | Shan | MSEA | Thailand | modern | 0 | Tai-Kadai | 19.37 | 98.14 | N_TK | N.A. | Kutanan_et_al_2021 | PASS |
| A1YU101 | Khonmueang | MSEA | Thailand | modern | 0 | Tai-Kadai | 19.58 | 99.51 | N_TK | N.A. | Kutanan_et_al_2021 | PASS |
| A1YU105 | Khonmueang | MSEA | Thailand | modern | 0 | Tai-Kadai | 19.58 | 99.51 | N_TK | N.A. | Kutanan_et_al_2021 | PASS |
| A1YU107 | Khonmueang | MSEA | Thailand | modern | 0 | Tai-Kadai | 19.58 | 99.51 | N_TK | N.A. | Kutanan_et_al_2021 | PASS |
| A1YU111 | Khonmueang | MSEA | Thailand | modern | 0 | Tai-Kadai | 19.58 | 99.51 | N_TK | N.A. | Kutanan_et_al_2021 | PASS |
| A1YU113 | Khonmueang | MSEA | Thailand | modern | 0 | Tai-Kadai | 19.58 | 99.51 | N_TK | N.A. | Kutanan_et_al_2021 | PASS |
| A1YU115 | Khonmueang | MSEA | Thailand | modern | 0 | Tai-Kadai | 19.58 | 99.51 | N_TK | N.A. | Kutanan_et_al_2021 | PASS |
| A1YU125 | Khonmueang | MSEA | Thailand | modern | 0 | Tai-Kadai | 19.58 | 99.51 | N_TK | N.A. | Kutanan_et_al_2021 | PASS |
| A1YU129 | Khonmueang | MSEA | Thailand | modern | 0 | Tai-Kadai | 19.58 | 99.51 | N_TK | N.A. | Kutanan_et_al_2021 | PASS |
| A1YU131 | Khonmueang | MSEA | Thailand | modern | 0 | Tai-Kadai | 19.58 | 99.51 | N_TK | N.A. | Kutanan_et_al_2021 | PASS |
| A1YU132 | Khonmueang | MSEA | Thailand | modern | 0 | Tai-Kadai | 19.58 | 99.51 | N_TK | N.A. | Kutanan_et_al_2021 | PASS |
| A2YU301 | Khonmueang | MSEA | Thailand | modern | 0 | Tai-Kadai | 19.02 | 98.58 | N_TK | N.A. | Kutanan_et_al_2021 | PASS |
| A2YU303 | Khonmueang | MSEA | Thailand | modern | 0 | Tai-Kadai | 19.02 | 98.58 | N_TK | N.A. | Kutanan_et_al_2021 | PASS |
| A2YU305 | Khonmueang | MSEA | Thailand | modern | 0 | Tai-Kadai | 19.02 | 98.58 | N_TK | N.A. | Kutanan_et_al_2021 | PASS |
| A2YU307 | Khonmueang | MSEA | Thailand | modern | 0 | Tai-Kadai | 19.02 | 98.58 | N_TK | N.A. | Kutanan_et_al_2021 | PASS |
| A2YU313 | Khonmueang | MSEA | Thailand | modern | 0 | Tai-Kadai | 19.02 | 98.58 | N_TK | N.A. | Kutanan_et_al_2021 | PASS |
| A2YU321 | Khonmueang | MSEA | Thailand | modern | 0 | Tai-Kadai | 19.02 | 98.58 | N_TK | N.A. | Kutanan_et_al_2021 | PASS |
| A2YU329 | Khonmueang | MSEA | Thailand | modern | 0 | Tai-Kadai | 19.02 | 98.58 | N_TK | N.A. | Kutanan_et_al_2021 | PASS |
| A2YU331 | Khonmueang | MSEA | Thailand | modern | 0 | Tai-Kadai | 19.02 | 98.58 | N_TK | N.A. | Kutanan_et_al_2021 | PASS |
| A2YU337 | Khonmueang | MSEA | Thailand | modern | 0 | Tai-Kadai | 19.02 | 98.58 | N_TK | N.A. | Kutanan_et_al_2021 | PASS |
| A2YU353 | Khonmueang | MSEA | Thailand | modern | 0 | Tai-Kadai | 19.02 | 98.58 | N_TK | N.A. | Kutanan_et_al_2021 | PASS |
| TL114 | Lue | MSEA | Thailand | modern | 0 | Tai-Kadai | 19.09 | 100.56 | N_TK | N.A. | Kutanan_et_al_2021 | PASS |
| TL116 | Lue | MSEA | Thailand | modern | 0 | Tai-Kadai | 19.09 | 100.56 | N_TK | N.A. | Kutanan_et_al_2021 | PASS |
| TL118 | Lue | MSEA | Thailand | modern | 0 | Tai-Kadai | 19.09 | 100.56 | N_TK | N.A. | Kutanan_et_al_2021 | PASS |
| TL120 | Lue | MSEA | Thailand | modern | 0 | Tai-Kadai | 19.09 | 100.56 | N_TK | N.A. | Kutanan_et_al_2021 | PASS |
| TL128 | Lue | MSEA | Thailand | modern | 0 | Tai-Kadai | 19.09 | 100.56 | N_TK | N.A. | Kutanan_et_al_2021 | PASS |
| TL148 | Lue | MSEA | Thailand | modern | 0 | Tai-Kadai | 19.09 | 100.56 | N_TK | N.A. | Kutanan_et_al_2021 | PASS |
| TL152 | Lue | MSEA | Thailand | modern | 0 | Tai-Kadai | 19.09 | 100.56 | N_TK | N.A. | Kutanan_et_al_2021 | PASS |
| TL160 | Lue | MSEA | Thailand | modern | 0 | Tai-Kadai | 19.09 | 100.56 | N_TK | N.A. | Kutanan_et_al_2021 | PASS |
| TL166 | Lue | MSEA | Thailand | modern | 0 | Tai-Kadai | 19.09 | 100.56 | N_TK | N.A. | Kutanan_et_al_2021 | PASS |
| TL170 | Lue | MSEA | Thailand | modern | 0 | Tai-Kadai | 19.09 | 100.56 | N_TK | N.A. | Kutanan_et_al_2021 | kin |
| BTA101 | BlackTai | MSEA | Thailand | modern | 0 | Tai-Kadai | 17.79 | 101.73 | NE_TK | N.A. | Kutanan_et_al_2021 | PASS |
| BTA103 | BlackTai | MSEA | Thailand | modern | 0 | Tai-Kadai | 17.79 | 101.73 | NE_TK | N.A. | Kutanan_et_al_2021 | PASS |

| | | | | | | | | | | | | |
|---|---|---|---|---|---|---|---|---|---|---|---|---|
| BTA105 | BlackTai | MSEA | Thailand | modern | 0 | Tai-Kadai | 17.79 | 101.73 | NE_TK | N.A. | Kutanan_et_al_2021 | PASS |
| BTA107 | BlackTai | MSEA | Thailand | modern | 0 | Tai-Kadai | 17.79 | 101.73 | NE_TK | N.A. | Kutanan_et_al_2021 | PASS |
| BTA111 | BlackTai | MSEA | Thailand | modern | 0 | Tai-Kadai | 17.79 | 101.73 | NE_TK | N.A. | Kutanan_et_al_2021 | PASS |
| BTA113 | BlackTai | MSEA | Thailand | modern | 0 | Tai-Kadai | 17.79 | 101.73 | NE_TK | N.A. | Kutanan_et_al_2021 | PASS |
| BTA115 | BlackTai | MSEA | Thailand | modern | 0 | Tai-Kadai | 17.79 | 101.73 | NE_TK | N.A. | Kutanan_et_al_2021 | PASS |
| BTA117 | BlackTai | MSEA | Thailand | modern | 0 | Tai-Kadai | 17.79 | 101.73 | NE_TK | N.A. | Kutanan_et_al_2021 | PASS |
| BTA119 | BlackTai | MSEA | Thailand | modern | 0 | Tai-Kadai | 17.79 | 101.73 | NE_TK | N.A. | Kutanan_et_al_2021 | PASS |
| LUA101 | Laotian | MSEA | Laos | modern | 0 | Tai-Kadai | 19.84 | 102.16 | NE_TK | N.A. | Kutanan_et_al_2021 | PASS |
| LUA105 | Laotian | MSEA | Laos | modern | 0 | Tai-Kadai | 19.84 | 102.16 | NE_TK | N.A. | Kutanan_et_al_2021 | PASS |
| LUA107 | Laotian | MSEA | Laos | modern | 0 | Tai-Kadai | 19.84 | 102.16 | NE_TK | N.A. | Kutanan_et_al_2021 | PASS |
| LUA109 | Laotian | MSEA | Laos | modern | 0 | Tai-Kadai | 19.84 | 102.16 | NE_TK | N.A. | Kutanan_et_al_2021 | PASS |
| LUA111 | Laotian | MSEA | Laos | modern | 0 | Tai-Kadai | 19.84 | 102.16 | NE_TK | N.A. | Kutanan_et_al_2021 | PASS |
| LUA113 | Laotian | MSEA | Laos | modern | 0 | Tai-Kadai | 19.84 | 102.16 | NE_TK | N.A. | Kutanan_et_al_2021 | PASS |
| LUA115 | Laotian | MSEA | Laos | modern | 0 | Tai-Kadai | 19.84 | 102.16 | NE_TK | N.A. | Kutanan_et_al_2021 | PASS |
| LUA117 | Laotian | MSEA | Laos | modern | 0 | Tai-Kadai | 19.84 | 102.16 | NE_TK | N.A. | Kutanan_et_al_2021 | PASS |
| LUA119 | Laotian | MSEA | Laos | modern | 0 | Tai-Kadai | 19.84 | 102.16 | NE_TK | N.A. | Kutanan_et_al_2021 | PASS |
| LUA121 | Laotian | MSEA | Laos | modern | 0 | Tai-Kadai | 19.84 | 102.16 | NE_TK | N.A. | Kutanan_et_al_2021 | PASS |
| LUA123 | Laotian | MSEA | Laos | modern | 0 | Tai-Kadai | 19.84 | 102.16 | NE_TK | N.A. | Kutanan_et_al_2021 | PASS |
| LUA125 | Laotian | MSEA | Laos | modern | 0 | Tai-Kadai | 19.84 | 102.16 | NE_TK | N.A. | Kutanan_et_al_2021 | PASS |
| LUA127 | Laotian | MSEA | Laos | modern | 0 | Tai-Kadai | 19.84 | 102.16 | NE_TK | N.A. | Kutanan_et_al_2021 | PASS |
| LUA129 | Laotian | MSEA | Laos | modern | 0 | Tai-Kadai | 19.84 | 102.16 | NE_TK | N.A. | Kutanan_et_al_2021 | PASS |
| LUA131 | Laotian | MSEA | Laos | modern | 0 | Tai-Kadai | 19.84 | 102.16 | NE_TK | N.A. | Kutanan_et_al_2021 | PASS |
| VIE126 | Laotian | MSEA | Laos | modern | 0 | Tai-Kadai | 17.97 | 102.59 | NE_TK | N.A. | Kutanan_et_al_2021 | PASS |
| VIE128 | Laotian | MSEA | Laos | modern | 0 | Tai-Kadai | 17.97 | 102.59 | NE_TK | N.A. | Kutanan_et_al_2021 | PASS |
| VIE136 | Laotian | MSEA | Laos | modern | 0 | Tai-Kadai | 17.97 | 102.59 | NE_TK | N.A. | Kutanan_et_al_2021 | PASS |
| VIE138 | Laotian | MSEA | Laos | modern | 0 | Tai-Kadai | 17.97 | 102.59 | NE_TK | N.A. | Kutanan_et_al_2021 | PASS |
| VIE140 | Laotian | MSEA | Laos | modern | 0 | Tai-Kadai | 17.97 | 102.59 | NE_TK | N.A. | Kutanan_et_al_2021 | PASS |
| VIE142 | Laotian | MSEA | Laos | modern | 0 | Tai-Kadai | 17.97 | 102.59 | NE_TK | N.A. | Kutanan_et_al_2021 | PASS |
| VIE150 | Laotian | MSEA | Laos | modern | 0 | Tai-Kadai | 17.97 | 102.59 | NE_TK | N.A. | Kutanan_et_al_2021 | PASS |
| VIE152 | Laotian | MSEA | Laos | modern | 0 | Tai-Kadai | 17.97 | 102.59 | NE_TK | N.A. | Kutanan_et_al_2021 | PASS |
| VIE154 | Laotian | MSEA | Laos | modern | 0 | Tai-Kadai | 17.97 | 102.59 | NE_TK | N.A. | Kutanan_et_al_2021 | PASS |
| VIE156 | Laotian | MSEA | Laos | modern | 0 | Tai-Kadai | 17.97 | 102.59 | NE_TK | N.A. | Kutanan_et_al_2021 | PASS |
| VIE158 | Laotian | MSEA | Laos | modern | 0 | Tai-Kadai | 17.97 | 102.59 | NE_TK | N.A. | Kutanan_et_al_2021 | PASS |
| VIE162 | Laotian | MSEA | Laos | modern | 0 | Tai-Kadai | 17.97 | 102.59 | NE_TK | N.A. | Kutanan_et_al_2021 | PASS |
| VIE164 | Laotian | MSEA | Laos | modern | 0 | Tai-Kadai | 17.97 | 102.59 | NE_TK | N.A. | Kutanan_et_al_2021 | PASS |
| VIE168 | Laotian | MSEA | Laos | modern | 0 | Tai-Kadai | 17.97 | 102.59 | NE_TK | N.A. | Kutanan_et_al_2021 | PASS |
| VIE170 | Laotian | MSEA | Laos | modern | 0 | Tai-Kadai | 17.97 | 102.59 | NE_TK | N.A. | Kutanan_et_al_2021 | PASS |
| LAO101 | LaoIsan | MSEA | Thailand | modern | 0 | Tai-Kadai | 15.62 | 103.5 | NE_TK | N.A. | Kutanan_et_al_2021 | PASS |
| LAO103 | LaoIsan | MSEA | Thailand | modern | 0 | Tai-Kadai | 15.62 | 103.5 | NE_TK | N.A. | Kutanan_et_al_2021 | PASS |
| LAO105 | LaoIsan | MSEA | Thailand | modern | 0 | Tai-Kadai | 15.62 | 103.5 | NE_TK | N.A. | Kutanan_et_al_2021 | PASS |
| LAO107 | LaoIsan | MSEA | Thailand | modern | 0 | Tai-Kadai | 15.62 | 103.5 | NE_TK | N.A. | Kutanan_et_al_2021 | PASS |

| | | | | | | | | | | | | |
|---|---|---|---|---|---|---|---|---|---|---|---|---|
| LAO109 | LaoIsan | MSEA | Thailand | modern | 0 | Tai-Kadai | 15.62 | 103.5 | NE_TK | N.A. | Kutanan_et_al_2021 | PASS |
| LAO111 | LaoIsan | MSEA | Thailand | modern | 0 | Tai-Kadai | 15.62 | 103.5 | NE_TK | N.A. | Kutanan_et_al_2021 | PASS |
| LAO113 | LaoIsan | MSEA | Thailand | modern | 0 | Tai-Kadai | 15.62 | 103.5 | NE_TK | N.A. | Kutanan_et_al_2021 | PASS |
| LAO115 | LaoIsan | MSEA | Thailand | modern | 0 | Tai-Kadai | 15.62 | 103.5 | NE_TK | N.A. | Kutanan_et_al_2021 | PASS |
| LAO117 | LaoIsan | MSEA | Thailand | modern | 0 | Tai-Kadai | 15.62 | 103.5 | NE_TK | N.A. | Kutanan_et_al_2021 | PASS |
| LAO119 | LaoIsan | MSEA | Thailand | modern | 0 | Tai-Kadai | 15.62 | 103.5 | NE_TK | N.A. | Kutanan_et_al_2021 | PASS |
| LAO401 | LaoIsan | MSEA | Thailand | modern | 0 | Tai-Kadai | 15.81 | 102.03 | NE_TK | N.A. | Kutanan_et_al_2021 | PASS |
| LAO403 | LaoIsan | MSEA | Thailand | modern | 0 | Tai-Kadai | 15.81 | 102.03 | NE_TK | N.A. | Kutanan_et_al_2021 | PASS |
| LAO405 | LaoIsan | MSEA | Thailand | modern | 0 | Tai-Kadai | 15.81 | 102.03 | NE_TK | N.A. | Kutanan_et_al_2021 | PASS |
| LAO407 | LaoIsan | MSEA | Thailand | modern | 0 | Tai-Kadai | 15.81 | 102.03 | NE_TK | N.A. | Kutanan_et_al_2021 | PASS |
| LAO409 | LaoIsan | MSEA | Thailand | modern | 0 | Tai-Kadai | 15.81 | 102.03 | NE_TK | N.A. | Kutanan_et_al_2021 | PASS |
| LAO411 | LaoIsan | MSEA | Thailand | modern | 0 | Tai-Kadai | 15.81 | 102.03 | NE_TK | N.A. | Kutanan_et_al_2021 | PASS |
| LAO413 | LaoIsan | MSEA | Thailand | modern | 0 | Tai-Kadai | 15.81 | 102.03 | NE_TK | N.A. | Kutanan_et_al_2021 | PASS |
| LAO417 | LaoIsan | MSEA | Thailand | modern | 0 | Tai-Kadai | 15.81 | 102.03 | NE_TK | N.A. | Kutanan_et_al_2021 | PASS |
| LAO421 | LaoIsan | MSEA | Thailand | modern | 0 | Tai-Kadai | 15.81 | 102.03 | NE_TK | N.A. | Kutanan_et_al_2021 | PASS |
| LAO423 | LaoIsan | MSEA | Thailand | modern | 0 | Tai-Kadai | 15.81 | 102.03 | NE_TK | N.A. | Kutanan_et_al_2021 | PASS |
| PUT101 | Phutai | MSEA | Thailand | modern | 0 | Tai-Kadai | 17.28 | 103.65 | NE_TK | N.A. | Kutanan_et_al_2021 | PASS |
| PUT102 | Phutai | MSEA | Thailand | modern | 0 | Tai-Kadai | 17.28 | 103.65 | NE_TK | N.A. | Kutanan_et_al_2021 | PASS |
| PUT103 | Phutai | MSEA | Thailand | modern | 0 | Tai-Kadai | 17.28 | 103.65 | NE_TK | N.A. | Kutanan_et_al_2021 | PASS |
| PUT105 | Phutai | MSEA | Thailand | modern | 0 | Tai-Kadai | 17.28 | 103.65 | NE_TK | N.A. | Kutanan_et_al_2021 | PASS |
| PUT109 | Phutai | MSEA | Thailand | modern | 0 | Tai-Kadai | 17.28 | 103.65 | NE_TK | N.A. | Kutanan_et_al_2021 | PASS |
| PUT110 | Phutai | MSEA | Thailand | modern | 0 | Tai-Kadai | 17.28 | 103.65 | NE_TK | N.A. | Kutanan_et_al_2021 | PASS |
| PUT111 | Phutai | MSEA | Thailand | modern | 0 | Tai-Kadai | 17.28 | 103.65 | NE_TK | N.A. | Kutanan_et_al_2021 | PASS |
| PUT112 | Phutai | MSEA | Thailand | modern | 0 | Tai-Kadai | 17.28 | 103.65 | NE_TK | N.A. | Kutanan_et_al_2021 | PASS |
| PUT113 | Phutai | MSEA | Thailand | modern | 0 | Tai-Kadai | 17.28 | 103.65 | NE_TK | N.A. | Kutanan_et_al_2021 | PASS |
| PUT114 | Phutai | MSEA | Thailand | modern | 0 | Tai-Kadai | 17.28 | 103.65 | NE_TK | N.A. | Kutanan_et_al_2021 | PASS |
| YOH103 | Nyaw | MSEA | Thailand | modern | 0 | Tai-Kadai | 17.55 | 104.09 | NE_TK | N.A. | Kutanan_et_al_2021 | PASS |
| YOH105 | Nyaw | MSEA | Thailand | modern | 0 | Tai-Kadai | 17.55 | 104.09 | NE_TK | N.A. | Kutanan_et_al_2021 | PASS |
| YOH107 | Nyaw | MSEA | Thailand | modern | 0 | Tai-Kadai | 17.55 | 104.09 | NE_TK | N.A. | Kutanan_et_al_2021 | PASS |
| YOH109 | Nyaw | MSEA | Thailand | modern | 0 | Tai-Kadai | 17.55 | 104.09 | NE_TK | N.A. | Kutanan_et_al_2021 | PASS |
| YOH111 | Nyaw | MSEA | Thailand | modern | 0 | Tai-Kadai | 17.55 | 104.09 | NE_TK | N.A. | Kutanan_et_al_2021 | PASS |
| YOH113 | Nyaw | MSEA | Thailand | modern | 0 | Tai-Kadai | 17.55 | 104.09 | NE_TK | N.A. | Kutanan_et_al_2021 | PASS |
| YOH115 | Nyaw | MSEA | Thailand | modern | 0 | Tai-Kadai | 17.55 | 104.09 | NE_TK | N.A. | Kutanan_et_al_2021 | PASS |
| YOH117 | Nyaw | MSEA | Thailand | modern | 0 | Tai-Kadai | 17.55 | 104.09 | NE_TK | N.A. | Kutanan_et_al_2021 | PASS |
| YOH121 | Nyaw | MSEA | Thailand | modern | 0 | Tai-Kadai | 17.55 | 104.09 | NE_TK | N.A. | Kutanan_et_al_2021 | PASS |
| YOH123 | Nyaw | MSEA | Thailand | modern | 0 | Tai-Kadai | 17.55 | 104.09 | NE_TK | N.A. | Kutanan_et_al_2021 | PASS |
| SAK104 | Seak | MSEA | Thailand | modern | 0 | Tai-Kadai | 17.45 | 104.74 | NE_TK | N.A. | Kutanan_et_al_2021 | PASS |
| SAK105 | Seak | MSEA | Thailand | modern | 0 | Tai-Kadai | 17.45 | 104.74 | NE_TK | N.A. | Kutanan_et_al_2021 | kin |
| SAK108 | Seak | MSEA | Thailand | modern | 0 | Tai-Kadai | 17.45 | 104.74 | NE_TK | N.A. | Kutanan_et_al_2021 | PASS |
| SAK110 | Seak | MSEA | Thailand | modern | 0 | Tai-Kadai | 17.45 | 104.74 | NE_TK | N.A. | Kutanan_et_al_2021 | PASS |
| SAK111 | Seak | MSEA | Thailand | modern | 0 | Tai-Kadai | 17.45 | 104.74 | NE_TK | N.A. | Kutanan_et_al_2021 | PASS |

| | | | | | | | | | | | | |
|---|---|---|---|---|---|---|---|---|---|---|---|---|
| SAK112 | Seak | MSEA | Thailand | modern | 0 | Tai-Kadai | 17.45 | 104.74 | NE_TK | N.A. | Kutanan_et_al_2021 | PASS |
| SAK114 | Seak | MSEA | Thailand | modern | 0 | Tai-Kadai | 17.45 | 104.74 | NE_TK | N.A. | Kutanan_et_al_2021 | PASS |
| SAK116 | Seak | MSEA | Thailand | modern | 0 | Tai-Kadai | 17.45 | 104.74 | NE_TK | N.A. | Kutanan_et_al_2021 | PASS |
| SAK118 | Seak | MSEA | Thailand | modern | 0 | Tai-Kadai | 17.45 | 104.74 | NE_TK | N.A. | Kutanan_et_al_2021 | PASS |
| SAK120 | Seak | MSEA | Thailand | modern | 0 | Tai-Kadai | 17.45 | 104.74 | NE_TK | N.A. | Kutanan_et_al_2021 | PASS |
| SAK126 | Seak | MSEA | Thailand | modern | 0 | Tai-Kadai | 17.45 | 104.74 | NE_TK | N.A. | Kutanan_et_al_2021 | PASS |
| SAK128 | Seak | MSEA | Thailand | modern | 0 | Tai-Kadai | 17.45 | 104.74 | NE_TK | N.A. | Kutanan_et_al_2021 | PASS |
| SAK130 | Seak | MSEA | Thailand | modern | 0 | Tai-Kadai | 17.45 | 104.74 | NE_TK | N.A. | Kutanan_et_al_2021 | PASS |
| SAK136 | Seak | MSEA | Thailand | modern | 0 | Tai-Kadai | 17.45 | 104.74 | NE_TK | N.A. | Kutanan_et_al_2021 | PASS |
| SAK138 | Seak | MSEA | Thailand | modern | 0 | Tai-Kadai | 17.45 | 104.74 | NE_TK | N.A. | Kutanan_et_al_2021 | PASS |
| KAL101 | Kalueang | MSEA | Thailand | modern | 0 | Tai-Kadai | 17.33 | 104.59 | NE_TK | N.A. | Kutanan_et_al_2021 | PASS |
| KAL102 | Kalueang | MSEA | Thailand | modern | 0 | Tai-Kadai | 17.33 | 104.59 | NE_TK | N.A. | Kutanan_et_al_2021 | PASS |
| KAL105 | Kalueang | MSEA | Thailand | modern | 0 | Tai-Kadai | 17.33 | 104.59 | NE_TK | N.A. | Kutanan_et_al_2021 | PASS |
| KAL107 | Kalueang | MSEA | Thailand | modern | 0 | Tai-Kadai | 17.33 | 104.59 | NE_TK | N.A. | Kutanan_et_al_2021 | PASS |
| KAL110 | Kalueang | MSEA | Thailand | modern | 0 | Tai-Kadai | 17.33 | 104.59 | NE_TK | N.A. | Kutanan_et_al_2021 | PASS |
| KAL112 | Kalueang | MSEA | Thailand | modern | 0 | Tai-Kadai | 17.33 | 104.59 | NE_TK | N.A. | Kutanan_et_al_2021 | PASS |
| KAL113 | Kalueang | MSEA | Thailand | modern | 0 | Tai-Kadai | 17.33 | 104.59 | NE_TK | N.A. | Kutanan_et_al_2021 | PASS |
| KAL115 | Kalueang | MSEA | Thailand | modern | 0 | Tai-Kadai | 17.33 | 104.59 | NE_TK | N.A. | Kutanan_et_al_2021 | PASS |
| KAL116 | Kalueang | MSEA | Thailand | modern | 0 | Tai-Kadai | 17.33 | 104.59 | NE_TK | N.A. | Kutanan_et_al_2021 | PASS |
| KAL118 | Kalueang | MSEA | Thailand | modern | 0 | Tai-Kadai | 17.33 | 104.59 | NE_TK | N.A. | Kutanan_et_al_2021 | PASS |
| CT201 | CentralThai | MSEA | Thailand | modern | 0 | Tai-Kadai | 13.71 | 99.77 | C_TK | N.A. | Kutanan_et_al_2021 | PASS |
| CT203 | CentralThai | MSEA | Thailand | modern | 0 | Tai-Kadai | 13.71 | 99.77 | C_TK | N.A. | Kutanan_et_al_2021 | PASS |
| CT205 | CentralThai | MSEA | Thailand | modern | 0 | Tai-Kadai | 13.71 | 99.77 | C_TK | N.A. | Kutanan_et_al_2021 | PASS |
| CT207 | CentralThai | MSEA | Thailand | modern | 0 | Tai-Kadai | 18.12 | 98.23 | C_TK | N.A. | Kutanan_et_al_2021 | PASS |
| CT209 | CentralThai | MSEA | Thailand | modern | 0 | Tai-Kadai | 13.71 | 99.77 | C_TK | N.A. | Kutanan_et_al_2021 | PASS |
| CT211 | CentralThai | MSEA | Thailand | modern | 0 | Tai-Kadai | 13.71 | 99.77 | C_TK | N.A. | Kutanan_et_al_2021 | PASS |
| CT213 | CentralThai | MSEA | Thailand | modern | 0 | Tai-Kadai | 13.71 | 99.77 | C_TK | N.A. | Kutanan_et_al_2021 | PASS |
| CT215 | CentralThai | MSEA | Thailand | modern | 0 | Tai-Kadai | 13.71 | 99.77 | C_TK | N.A. | Kutanan_et_al_2021 | PASS |
| CT217 | CentralThai | MSEA | Thailand | modern | 0 | Tai-Kadai | 13.71 | 99.77 | C_TK | N.A. | Kutanan_et_al_2021 | PASS |
| CT219 | CentralThai | MSEA | Thailand | modern | 0 | Tai-Kadai | 13.71 | 99.77 | C_TK | N.A. | Kutanan_et_al_2021 | PASS |
| CT301 | CentralThai | MSEA | Thailand | modern | 0 | Tai-Kadai | 13.74 | 100.07 | C_TK | N.A. | Kutanan_et_al_2021 | PASS |
| CT303 | CentralThai | MSEA | Thailand | modern | 0 | Tai-Kadai | 13.71 | 100.07 | C_TK | N.A. | Kutanan_et_al_2021 | PASS |
| CT305 | CentralThai | MSEA | Thailand | modern | 0 | Tai-Kadai | 13.74 | 100.07 | C_TK | N.A. | Kutanan_et_al_2021 | PASS |
| CT307 | CentralThai | MSEA | Thailand | modern | 0 | Tai-Kadai | 13.74 | 100.07 | C_TK | N.A. | Kutanan_et_al_2021 | PASS |
| CT309 | CentralThai | MSEA | Thailand | modern | 0 | Tai-Kadai | 13.74 | 100.07 | C_TK | N.A. | Kutanan_et_al_2021 | PASS |
| CT311 | CentralThai | MSEA | Thailand | modern | 0 | Tai-Kadai | 13.74 | 100.07 | C_TK | N.A. | Kutanan_et_al_2021 | PASS |
| CT313 | CentralThai | MSEA | Thailand | modern | 0 | Tai-Kadai | 13.74 | 100.07 | C_TK | N.A. | Kutanan_et_al_2021 | PASS |
| CT315 | CentralThai | MSEA | Thailand | modern | 0 | Tai-Kadai | 13.74 | 100.07 | C_TK | N.A. | Kutanan_et_al_2021 | PASS |
| CT317 | CentralThai | MSEA | Thailand | modern | 0 | Tai-Kadai | 13.74 | 100.07 | C_TK | N.A. | Kutanan_et_al_2021 | PASS |
| CT319 | CentralThai | MSEA | Thailand | modern | 0 | Tai-Kadai | 13.74 | 100.07 | C_TK | N.A. | Kutanan_et_al_2021 | PASS |
| BST101 | SouthernThai_TK | MSEA | Thailand | modern | 0 | Tai-Kadai | 8.31 | 100.05 | S_TK | N.A. | Kutanan_et_al_2021 | PASS |

| | | | | | | | | | | | | |
|---|---|---|---|---|---|---|---|---|---|---|---|---|
| BST103 | SouthernThai_TK | MSEA | Thailand | modern | 0 | Tai-Kadai | 8.31 | 100.05 | S_TK | N.A. | Kutanan_et_al_2021 | PASS |
| BST105 | SouthernThai_TK | MSEA | Thailand | modern | 0 | Tai-Kadai | 8.31 | 100.05 | S_TK | N.A. | Kutanan_et_al_2021 | PASS |
| BST107 | SouthernThai_TK | MSEA | Thailand | modern | 0 | Tai-Kadai | 8.31 | 100.05 | S_TK | N.A. | Kutanan_et_al_2021 | PASS |
| BST109 | SouthernThai_TK | MSEA | Thailand | modern | 0 | Tai-Kadai | 8.31 | 100.05 | S_TK | N.A. | Kutanan_et_al_2021 | PASS |
| BST111 | SouthernThai_TK | MSEA | Thailand | modern | 0 | Tai-Kadai | 8.31 | 100.05 | S_TK | N.A. | Kutanan_et_al_2021 | PASS |
| BST113 | SouthernThai_TK | MSEA | Thailand | modern | 0 | Tai-Kadai | 8.31 | 100.05 | S_TK | N.A. | Kutanan_et_al_2021 | PASS |
| BST115 | SouthernThai_TK | MSEA | Thailand | modern | 0 | Tai-Kadai | 8.31 | 100.05 | S_TK | N.A. | Kutanan_et_al_2021 | PASS |
| BST117 | SouthernThai_TK | MSEA | Thailand | modern | 0 | Tai-Kadai | 8.31 | 100.05 | S_TK | N.A. | Kutanan_et_al_2021 | PASS |
| BST119 | SouthernThai_TK | MSEA | Thailand | modern | 0 | Tai-Kadai | 8.31 | 100.05 | S_TK | N.A. | Kutanan_et_al_2021 | PASS |
| BST121 | SouthernThai_TK | MSEA | Thailand | modern | 0 | Tai-Kadai | 8.31 | 100.05 | S_TK | N.A. | Kutanan_et_al_2021 | PASS |
| BST123 | SouthernThai_TK | MSEA | Thailand | modern | 0 | Tai-Kadai | 8.31 | 100.05 | S_TK | N.A. | Kutanan_et_al_2021 | PASS |
| BST125 | SouthernThai_TK | MSEA | Thailand | modern | 0 | Tai-Kadai | 8.31 | 100.05 | S_TK | N.A. | Kutanan_et_al_2021 | PASS |
| BST129 | SouthernThai_TK | MSEA | Thailand | modern | 0 | Tai-Kadai | 8.31 | 100.05 | S_TK | N.A. | Kutanan_et_al_2021 | PASS |
| BST131 | SouthernThai_TK | MSEA | Thailand | modern | 0 | Tai-Kadai | 8.31 | 100.05 | S_TK | N.A. | Kutanan_et_al_2021 | PASS |
| CHI007 | Thai | MSEA | Thailand | modern | 0 | Tai-Kadai | 13.8 | 100.5 | Thai | N.A. | Lazaridis_et_al_2014 | PASS |
| CHI034 | Thai | MSEA | Thailand | modern | 0 | Tai-Kadai | 13.8 | 100.5 | Thai | N.A. | Lazaridis_et_al_2014 | PASS |
| DCH002 | Thai | MSEA | Thailand | modern | 0 | Tai-Kadai | 13.8 | 100.5 | Thai | N.A. | Lazaridis_et_al_2014 | kin |
| DCH006 | Thai | MSEA | Thailand | modern | 0 | Tai-Kadai | 13.8 | 100.5 | Thai | N.A. | Lazaridis_et_al_2014 | PASS |
| DCH007 | Thai | MSEA | Thailand | modern | 0 | Tai-Kadai | 13.8 | 100.5 | Thai | N.A. | Lazaridis_et_al_2014 | PASS |
| DCH008 | Thai | MSEA | Thailand | modern | 0 | Tai-Kadai | 13.8 | 100.5 | Thai | N.A. | Lazaridis_et_al_2014 | PASS |
| DCH009 | Thai | MSEA | Thailand | modern | 0 | Tai-Kadai | 13.8 | 100.5 | Thai | N.A. | Lazaridis_et_al_2014 | PASS |
| DCH010 | Thai | MSEA | Thailand | modern | 0 | Tai-Kadai | 13.8 | 100.5 | Thai | N.A. | Lazaridis_et_al_2014 | PASS |
| DCH011 | Thai | MSEA | Thailand | modern | 0 | Tai-Kadai | 13.8 | 100.5 | Thai | N.A. | Lazaridis_et_al_2014 | kin |
| DCH012 | Thai | MSEA | Thailand | modern | 0 | Tai-Kadai | 13.8 | 100.5 | Thai | N.A. | Lazaridis_et_al_2014 | PASS |
| HGDP00449 | Mbuti | Africa | Congo | modern | 0 | Central_Sudanic | 1 | 29 | N.A. | N.A. | Patterson_et_al_2012 | PASS |
| HGDP00462 | Mbuti | Africa | Congo | modern | 0 | Central_Sudanic | 1 | 29 | N.A. | N.A. | Patterson_et_al_2012 | PASS |
| HGDP00463 | Mbuti | Africa | Congo | modern | 0 | Central_Sudanic | 1 | 29 | N.A. | N.A. | Patterson_et_al_2012 | PASS |
| HGDP00467 | Mbuti | Africa | Congo | modern | 0 | Central_Sudanic | 1 | 29 | N.A. | N.A. | Patterson_et_al_2012 | PASS |
| HGDP00474 | Mbuti | Africa | Congo | modern | 0 | Central_Sudanic | 1 | 29 | N.A. | N.A. | Patterson_et_al_2012 | PASS |
| HGDP00476 | Mbuti | Africa | Congo | modern | 0 | Central_Sudanic | 1 | 29 | N.A. | N.A. | Patterson_et_al_2012 | PASS |
| HGDP00478 | Mbuti | Africa | Congo | modern | 0 | Central_Sudanic | 1 | 29 | N.A. | N.A. | Patterson_et_al_2012 | PASS |
| HGDP00982 | Mbuti | Africa | Congo | modern | 0 | Central_Sudanic | 1 | 29 | N.A. | N.A. | Patterson_et_al_2012 | PASS |
| HGDP00984 | Mbuti | Africa | Congo | modern | 0 | Central_Sudanic | 1 | 29 | N.A. | N.A. | Patterson_et_al_2012 | PASS |
| HGDP01081 | Mbuti | Africa | Congo | modern | 0 | Central_Sudanic | 1 | 29 | N.A. | N.A. | Patterson_et_al_2012 | PASS |
| HGDP00511 | French | Europe | France | modern | 0 | Indo-European | 46 | 2 | N.A. | N.A. | Patterson_et_al_2012 | PASS |
| HGDP00512 | French | Europe | France | modern | 0 | Indo-European | 46 | 2 | N.A. | N.A. | Patterson_et_al_2012 | PASS |
| HGDP00513 | French | Europe | France | modern | 0 | Indo-European | 46 | 2 | N.A. | N.A. | Patterson_et_al_2012 | PASS |
| HGDP00514 | French | Europe | France | modern | 0 | Indo-European | 46 | 2 | N.A. | N.A. | Patterson_et_al_2012 | PASS |
| HGDP00515 | French | Europe | France | modern | 0 | Indo-European | 46 | 2 | N.A. | N.A. | Patterson_et_al_2012 | PASS |
| HGDP00516 | French | Europe | France | modern | 0 | Indo-European | 46 | 2 | N.A. | N.A. | Patterson_et_al_2012 | PASS |
| HGDP00517 | French | Europe | France | modern | 0 | Indo-European | 46 | 2 | N.A. | N.A. | Patterson_et_al_2012 | PASS |

| | | | | | | | | | | | | |
|---|---|---|---|---|---|---|---|---|---|---|---|---|
| HGDP00518 | French | Europe | France | modern | 0 | Indo-European | 46 | 2 | N.A. | N.A. | Patterson_et_al_2012 | PASS |
| HGDP00519 | French | Europe | France | modern | 0 | Indo-European | 46 | 2 | N.A. | N.A. | Patterson_et_al_2012 | PASS |
| HGDP00522 | French | Europe | France | modern | 0 | Indo-European | 46 | 2 | N.A. | N.A. | Patterson_et_al_2012 | PASS |
| Tiwari1 | Brahmin_Tiwari | SouthAsia | India | modern | 0 | Indo-European | 21.9 | 83.4 | N.A. | N.A. | Lazaridis_et_al_2014 | PASS |
| Tiwari2 | Brahmin_Tiwari | SouthAsia | India | modern | 0 | Indo-European | 21.9 | 83.4 | N.A. | N.A. | Lazaridis_et_al_2014 | PASS |
| Tiwari3 | Brahmin_Tiwari | SouthAsia | India | modern | 0 | Indo-European | 21.9 | 83.4 | N.A. | N.A. | Lazaridis_et_al_2014 | PASS |
| Tiwari4 | Brahmin_Tiwari | SouthAsia | India | modern | 0 | Indo-European | 21.9 | 83.4 | N.A. | N.A. | Lazaridis_et_al_2014 | PASS |
| Tiwari5 | Brahmin_Tiwari | SouthAsia | India | modern | 0 | Indo-European | 21.9 | 83.4 | N.A. | N.A. | Lazaridis_et_al_2014 | PASS |
| Tiwari6 | Brahmin_Tiwari | SouthAsia | India | modern | 0 | Indo-European | 21.9 | 83.4 | N.A. | N.A. | Lazaridis_et_al_2014 | PASS |
| Tiwari7 | Brahmin_Tiwari | SouthAsia | India | modern | 0 | Indo-European | 21.9 | 83.4 | N.A. | N.A. | Lazaridis_et_al_2014 | PASS |
| Tiwari8 | Brahmin_Tiwari | SouthAsia | India | modern | 0 | Indo-European | 21.9 | 83.4 | N.A. | N.A. | Lazaridis_et_al_2014 | PASS |
| Tiwari9 | Brahmin_Tiwari | SouthAsia | India | modern | 0 | Indo-European | 21.9 | 83.4 | N.A. | N.A. | Lazaridis_et_al_2014 | PASS |
| Tiwari10 | Brahmin_Tiwari | SouthAsia | India | modern | 0 | Indo-European | 21.9 | 83.4 | N.A. | N.A. | Lazaridis_et_al_2014 | PASS |
| Tiwari11 | Brahmin_Tiwari | SouthAsia | India | modern | 0 | Indo-European | 21.9 | 83.4 | N.A. | N.A. | Lazaridis_et_al_2014 | PASS |
| Tiwari12 | Brahmin_Tiwari | SouthAsia | India | modern | 0 | Indo-European | 21.9 | 83.4 | N.A. | N.A. | Lazaridis_et_al_2014 | PASS |
| Tiwari13 | Brahmin_Tiwari | SouthAsia | India | modern | 0 | Indo-European | 21.9 | 83.4 | N.A. | N.A. | Lazaridis_et_al_2014 | PASS |
| Tiwari14 | Brahmin_Tiwari | SouthAsia | India | modern | 0 | Indo-European | 21.9 | 83.4 | N.A. | N.A. | Lazaridis_et_al_2014 | PASS |
| Tiwari15 | Brahmin_Tiwari | SouthAsia | India | modern | 0 | Indo-European | 21.9 | 83.4 | N.A. | N.A. | Lazaridis_et_al_2014 | PASS |
| NA20847 | Gujarati | SouthAsia | India | modern | 0 | Indo-European | 23.2 | 72.7 | N.A. | N.A. | Lazaridis_et_al_2014 | PASS |
| NA20854 | Gujarati | SouthAsia | India | modern | 0 | Indo-European | 23.2 | 72.7 | N.A. | N.A. | Lazaridis_et_al_2014 | PASS |
| NA20881 | Gujarati | SouthAsia | India | modern | 0 | Indo-European | 23.2 | 72.7 | N.A. | N.A. | Lazaridis_et_al_2014 | PASS |
| NA20882 | Gujarati | SouthAsia | India | modern | 0 | Indo-European | 23.2 | 72.7 | N.A. | N.A. | Lazaridis_et_al_2014 | PASS |
| NA20899 | Gujarati | SouthAsia | India | modern | 0 | Indo-European | 23.2 | 72.7 | N.A. | N.A. | Lazaridis_et_al_2014 | PASS |
| NA20910 | Gujarati | SouthAsia | India | modern | 0 | Indo-European | 23.2 | 72.7 | N.A. | N.A. | Lazaridis_et_al_2014 | PASS |
| NA21125 | Gujarati | SouthAsia | India | modern | 0 | Indo-European | 23.2 | 72.7 | N.A. | N.A. | Lazaridis_et_al_2014 | PASS |
| NA20858 | Gujarati | SouthAsia | India | modern | 0 | Indo-European | 23.2 | 72.7 | N.A. | N.A. | Lazaridis_et_al_2014 | PASS |
| NA20866 | Gujarati | SouthAsia | India | modern | 0 | Indo-European | 23.2 | 72.7 | N.A. | N.A. | Lazaridis_et_al_2014 | PASS |
| NA20873 | Gujarati | SouthAsia | India | modern | 0 | Indo-European | 23.2 | 72.7 | N.A. | N.A. | Lazaridis_et_al_2014 | PASS |
| NA20883 | Gujarati | SouthAsia | India | modern | 0 | Indo-European | 23.2 | 72.7 | N.A. | N.A. | Lazaridis_et_al_2014 | PASS |
| NA20885 | Gujarati | SouthAsia | India | modern | 0 | Indo-European | 23.2 | 72.7 | N.A. | N.A. | Lazaridis_et_al_2014 | PASS |
| NA20898 | Gujarati | SouthAsia | India | modern | 0 | Indo-European | 23.2 | 72.7 | N.A. | N.A. | Lazaridis_et_al_2014 | PASS |
| NA20903 | Gujarati | SouthAsia | India | modern | 0 | Indo-European | 23.2 | 72.7 | N.A. | N.A. | Lazaridis_et_al_2014 | PASS |
| NA20904 | Gujarati | SouthAsia | India | modern | 0 | Indo-European | 23.2 | 72.7 | N.A. | N.A. | Lazaridis_et_al_2014 | PASS |
| NA20911 | Gujarati | SouthAsia | India | modern | 0 | Indo-European | 23.2 | 72.7 | N.A. | N.A. | Lazaridis_et_al_2014 | PASS |
| NA21116 | Gujarati | SouthAsia | India | modern | 0 | Indo-European | 23.2 | 72.7 | N.A. | N.A. | Lazaridis_et_al_2014 | PASS |
| NA21123 | Gujarati | SouthAsia | India | modern | 0 | Indo-European | 23.2 | 72.7 | N.A. | N.A. | Lazaridis_et_al_2014 | PASS |
| NA21097 | Gujarati | SouthAsia | India | modern | 0 | Indo-European | 23.2 | 72.7 | N.A. | N.A. | Lazaridis_et_al_2014 | PASS |
| NA20862 | Gujarati | SouthAsia | India | modern | 0 | Indo-European | 23.2 | 72.7 | N.A. | N.A. | Lazaridis_et_al_2014 | PASS |
| Lodhi2 | Lodhi | SouthAsia | India | modern | 0 | Indo-European | 25.45 | 78.62 | N.A. | N.A. | Lazaridis_et_al_2014 | PASS |
| Lodhi3 | Lodhi | SouthAsia | India | modern | 0 | Indo-European | 25.45 | 78.62 | N.A. | N.A. | Lazaridis_et_al_2014 | PASS |
| Lodhi4 | Lodhi | SouthAsia | India | modern | 0 | Indo-European | 25.45 | 78.62 | N.A. | N.A. | Lazaridis_et_al_2014 | PASS |

| | | | | | | | | | | | | |
|---|---|---|---|---|---|---|---|---|---|---|---|---|
| Lodhi5 | Lodhi | SouthAsia | India | modern | 0 | Indo-European | 25.45 | 78.62 | N.A. | N.A. | Lazaridis_et_al_2014 | PASS |
| Lodhi6 | Lodhi | SouthAsia | India | modern | 0 | Indo-European | 25.45 | 78.62 | N.A. | N.A. | Lazaridis_et_al_2014 | PASS |
| Lodhi7 | Lodhi | SouthAsia | India | modern | 0 | Indo-European | 25.45 | 78.62 | N.A. | N.A. | Lazaridis_et_al_2014 | PASS |
| Lodhi8 | Lodhi | SouthAsia | India | modern | 0 | Indo-European | 25.45 | 78.62 | N.A. | N.A. | Lazaridis_et_al_2014 | PASS |
| Lodhi9 | Lodhi | SouthAsia | India | modern | 0 | Indo-European | 25.45 | 78.62 | N.A. | N.A. | Lazaridis_et_al_2014 | PASS |
| Lodhi10 | Lodhi | SouthAsia | India | modern | 0 | Indo-European | 25.45 | 78.62 | N.A. | N.A. | Lazaridis_et_al_2014 | PASS |
| Lodhi11 | Lodhi | SouthAsia | India | modern | 0 | Indo-European | 25.45 | 78.62 | N.A. | N.A. | Lazaridis_et_al_2014 | PASS |
| Lodhi13 | Lodhi | SouthAsia | India | modern | 0 | Indo-European | 25.45 | 78.62 | N.A. | N.A. | Lazaridis_et_al_2014 | PASS |
| Lodhi14 | Lodhi | SouthAsia | India | modern | 0 | Indo-European | 25.45 | 78.62 | N.A. | N.A. | Lazaridis_et_al_2014 | PASS |
| Lodhi15 | Lodhi | SouthAsia | India | modern | 0 | Indo-European | 25.45 | 78.62 | N.A. | N.A. | Lazaridis_et_al_2014 | PASS |
| Vishwabrahmin1 | Vishwabrahmin | SouthAsia | India | modern | 0 | Dravidian | 16.3 | 80.48 | N.A. | N.A. | Lazaridis_et_al_2014 | PASS |
| Vishwabrahmin3 | Vishwabrahmin | SouthAsia | India | modern | 0 | Dravidian | 16.3 | 80.48 | N.A. | N.A. | Lazaridis_et_al_2014 | PASS |
| Vishwabrahmin4 | Vishwabrahmin | SouthAsia | India | modern | 0 | Dravidian | 16.3 | 80.48 | N.A. | N.A. | Lazaridis_et_al_2014 | PASS |
| Vishwabrahmin5 | Vishwabrahmin | SouthAsia | India | modern | 0 | Dravidian | 16.3 | 80.48 | N.A. | N.A. | Lazaridis_et_al_2014 | PASS |
| Vishwabrahmin6 | Vishwabrahmin | SouthAsia | India | modern | 0 | Dravidian | 16.3 | 80.48 | N.A. | N.A. | Lazaridis_et_al_2014 | PASS |
| Vishwabrahmin7 | Vishwabrahmin | SouthAsia | India | modern | 0 | Dravidian | 16.3 | 80.48 | N.A. | N.A. | Lazaridis_et_al_2014 | PASS |
| Vishwabrahmin8 | Vishwabrahmin | SouthAsia | India | modern | 0 | Dravidian | 16.3 | 80.48 | N.A. | N.A. | Lazaridis_et_al_2014 | PASS |
| Vishwabrahmin9 | Vishwabrahmin | SouthAsia | India | modern | 0 | Dravidian | 16.3 | 80.48 | N.A. | N.A. | Lazaridis_et_al_2014 | PASS |
| Vishwabrahmin10 | Vishwabrahmin | SouthAsia | India | modern | 0 | Dravidian | 16.3 | 80.48 | N.A. | N.A. | Lazaridis_et_al_2014 | PASS |
| Vishwabrahmin11 | Vishwabrahmin | SouthAsia | India | modern | 0 | Dravidian | 16.3 | 80.48 | N.A. | N.A. | Lazaridis_et_al_2014 | PASS |
| Vishwabrahmin12 | Vishwabrahmin | SouthAsia | India | modern | 0 | Dravidian | 16.3 | 80.48 | N.A. | N.A. | Lazaridis_et_al_2014 | PASS |
| Vishwabrahmin13 | Vishwabrahmin | SouthAsia | India | modern | 0 | Dravidian | 16.3 | 80.48 | N.A. | N.A. | Lazaridis_et_al_2014 | PASS |
| Vishwabrahmin14 | Vishwabrahmin | SouthAsia | India | modern | 0 | Dravidian | 16.3 | 80.48 | N.A. | N.A. | Lazaridis_et_al_2014 | PASS |
| Vishwabrahmin15 | Vishwabrahmin | SouthAsia | India | modern | 0 | Dravidian | 16.3 | 80.48 | N.A. | N.A. | Lazaridis_et_al_2014 | PASS |
| Mala1 | Mala | SouthAsia | India | modern | 0 | Dravidian | 18.67 | 78.17 | N.A. | N.A. | Lazaridis_et_al_2014 | PASS |
| Mala2 | Mala | SouthAsia | India | modern | 0 | Dravidian | 18.67 | 78.17 | N.A. | N.A. | Lazaridis_et_al_2014 | PASS |
| Mala3 | Mala | SouthAsia | India | modern | 0 | Dravidian | 18.67 | 78.17 | N.A. | N.A. | Lazaridis_et_al_2014 | PASS |
| Mala4 | Mala | SouthAsia | India | modern | 0 | Dravidian | 18.67 | 78.17 | N.A. | N.A. | Lazaridis_et_al_2014 | PASS |
| Mala5 | Mala | SouthAsia | India | modern | 0 | Dravidian | 18.67 | 78.17 | N.A. | N.A. | Lazaridis_et_al_2014 | PASS |
| Mala6 | Mala | SouthAsia | India | modern | 0 | Dravidian | 18.67 | 78.17 | N.A. | N.A. | Lazaridis_et_al_2014 | PASS |
| Mala7 | Mala | SouthAsia | India | modern | 0 | Dravidian | 18.67 | 78.17 | N.A. | N.A. | Lazaridis_et_al_2014 | PASS |
| Mala8 | Mala | SouthAsia | India | modern | 0 | Dravidian | 18.67 | 78.17 | N.A. | N.A. | Lazaridis_et_al_2014 | PASS |
| Mala9 | Mala | SouthAsia | India | modern | 0 | Dravidian | 18.67 | 78.17 | N.A. | N.A. | Lazaridis_et_al_2014 | PASS |
| Mala10 | Mala | SouthAsia | India | modern | 0 | Dravidian | 18.67 | 78.17 | N.A. | N.A. | Lazaridis_et_al_2014 | PASS |
| Mala11 | Mala | SouthAsia | India | modern | 0 | Dravidian | 18.67 | 78.17 | N.A. | N.A. | Lazaridis_et_al_2014 | PASS |
| Mala12 | Mala | SouthAsia | India | modern | 0 | Dravidian | 18.67 | 78.17 | N.A. | N.A. | Lazaridis_et_al_2014 | PASS |
| Mala13 | Mala | SouthAsia | India | modern | 0 | Dravidian | 18.67 | 78.17 | N.A. | N.A. | Lazaridis_et_al_2014 | PASS |
| Mala14 | Mala | SouthAsia | India | modern | 0 | Dravidian | 18.67 | 78.17 | N.A. | N.A. | Lazaridis_et_al_2014 | PASS |
| Mala15 | Mala | SouthAsia | India | modern | 0 | Dravidian | 18.67 | 78.17 | N.A. | N.A. | Lazaridis_et_al_2014 | PASS |
| Kharia1 | Kharia | SouthAsia | India | modern | 0 | Austroasiatic | 25.77 | 82.73 | N.A. | N.A. | Lazaridis_et_al_2014 | PASS |
| Kharia2 | Kharia | SouthAsia | India | modern | 0 | Austroasiatic | 25.77 | 82.73 | N.A. | N.A. | Lazaridis_et_al_2014 | PASS |

| | | | | | | | | | | | | |
|---|---|---|---|---|---|---|---|---|---|---|---|---|
| Kharia3 | Kharia | SouthAsia | India | modern | 0 | Austroasiatic | 25.77 | 82.73 | N.A. | N.A. | Lazaridis_et_al_2014 | PASS |
| Kharia4 | Kharia | SouthAsia | India | modern | 0 | Austroasiatic | 25.77 | 82.73 | N.A. | N.A. | Lazaridis_et_al_2014 | PASS |
| Kharia5 | Kharia | SouthAsia | India | modern | 0 | Austroasiatic | 25.77 | 82.73 | N.A. | N.A. | Lazaridis_et_al_2014 | PASS |
| Kharia6 | Kharia | SouthAsia | India | modern | 0 | Austroasiatic | 25.77 | 82.73 | N.A. | N.A. | Lazaridis_et_al_2014 | PASS |
| Kharia7 | Kharia | SouthAsia | India | modern | 0 | Austroasiatic | 25.77 | 82.73 | N.A. | N.A. | Lazaridis_et_al_2014 | PASS |
| Kharia8 | Kharia | SouthAsia | India | modern | 0 | Austroasiatic | 25.77 | 82.73 | N.A. | N.A. | Lazaridis_et_al_2014 | PASS |
| Kharia9 | Kharia | SouthAsia | India | modern | 0 | Austroasiatic | 25.77 | 82.73 | N.A. | N.A. | Lazaridis_et_al_2014 | PASS |
| Kharia10 | Kharia | SouthAsia | India | modern | 0 | Austroasiatic | 25.77 | 82.73 | N.A. | N.A. | Lazaridis_et_al_2014 | PASS |
| Kharia11 | Kharia | SouthAsia | India | modern | 0 | Austroasiatic | 25.77 | 82.73 | N.A. | N.A. | Lazaridis_et_al_2014 | PASS |
| Kharia12 | Kharia | SouthAsia | India | modern | 0 | Austroasiatic | 25.77 | 82.73 | N.A. | N.A. | Lazaridis_et_al_2014 | PASS |
| Kharia13 | Kharia | SouthAsia | India | modern | 0 | Austroasiatic | 25.77 | 82.73 | N.A. | N.A. | Lazaridis_et_al_2014 | PASS |
| Kharia14 | Kharia | SouthAsia | India | modern | 0 | Austroasiatic | 25.77 | 82.73 | N.A. | N.A. | Lazaridis_et_al_2014 | PASS |
| Kharia15 | Kharia | SouthAsia | India | modern | 0 | Austroasiatic | 25.77 | 82.73 | N.A. | N.A. | Lazaridis_et_al_2014 | PASS |
| Onge1 | Onge | SouthAsia | India | modern | 0 | Andamanese | 10.8 | 92.5 | N.A. | N.A. | Lazaridis_et_al_2014 | PASS |
| Onge2 | Onge | SouthAsia | India | modern | 0 | Andamanese | 10.8 | 92.5 | N.A. | N.A. | Lazaridis_et_al_2014 | PASS |
| Onge3 | Onge | SouthAsia | India | modern | 0 | Andamanese | 10.8 | 92.5 | N.A. | N.A. | Lazaridis_et_al_2014 | PASS |
| Onge7 | Onge | SouthAsia | India | modern | 0 | Andamanese | 10.8 | 92.5 | N.A. | N.A. | Lazaridis_et_al_2014 | PASS |
| Onge9 | Onge | SouthAsia | India | modern | 0 | Andamanese | 10.8 | 92.5 | N.A. | N.A. | Lazaridis_et_al_2014 | PASS |
| Onge12 | Onge | SouthAsia | India | modern | 0 | Andamanese | 10.8 | 92.5 | N.A. | N.A. | Lazaridis_et_al_2014 | PASS |
| Onge13 | Onge | SouthAsia | India | modern | 0 | Andamanese | 10.8 | 92.5 | N.A. | N.A. | Lazaridis_et_al_2014 | PASS |
| Onge14 | Onge | SouthAsia | India | modern | 0 | Andamanese | 10.8 | 92.5 | N.A. | N.A. | Lazaridis_et_al_2014 | PASS |
| Onge15 | Onge | SouthAsia | India | modern | 0 | Andamanese | 10.8 | 92.5 | N.A. | N.A. | Lazaridis_et_al_2014 | PASS |
| Onge16 | Onge | SouthAsia | India | modern | 0 | Andamanese | 10.8 | 92.5 | N.A. | N.A. | Lazaridis_et_al_2014 | PASS |
| Onge17 | Onge | SouthAsia | India | modern | 0 | Andamanese | 10.8 | 92.5 | N.A. | N.A. | Lazaridis_et_al_2014 | PASS |
| UV345 | Baining_Malasait | Oceania | Papua_New_Guinea | modern | 0 | East_New_Britain | -4.47 | 151.9 | N.A. | N.A. | Skoglund_et_al_2016 | PASS |
| UV346 | Baining_Malasait | Oceania | Papua_New_Guinea | modern | 0 | East_New_Britain | -4.47 | 151.9 | N.A. | N.A. | Skoglund_et_al_2016 | PASS |
| UV347 | Baining_Malasait | Oceania | Papua_New_Guinea | modern | 0 | East_New_Britain | -4.47 | 151.9 | N.A. | N.A. | Skoglund_et_al_2016 | PASS |
| UV368 | Baining_Malasait | Oceania | Papua_New_Guinea | modern | 0 | East_New_Britain | -4.47 | 151.9 | N.A. | N.A. | Skoglund_et_al_2016 | kin |
| UV370 | Baining_Malasait | Oceania | Papua_New_Guinea | modern | 0 | East_New_Britain | -4.47 | 151.9 | N.A. | N.A. | Skoglund_et_al_2016 | PASS |
| UV001 | Baining_Marabu | Oceania | Papua_New_Guinea | modern | 0 | East_New_Britain | -4.63 | 152.3 | N.A. | N.A. | Skoglund_et_al_2016 | PASS |
| UV076 | Baining_Marabu | Oceania | Papua_New_Guinea | modern | 0 | East_New_Britain | -4.63 | 152.3 | N.A. | N.A. | Skoglund_et_al_2016 | PASS |
| UV005 | Baining_Marabu | Oceania | Papua_New_Guinea | modern | 0 | East_New_Britain | -4.63 | 152.3 | N.A. | N.A. | Skoglund_et_al_2016 | PASS |
| UV02 | Baining_Marabu | Oceania | Papua_New_Guinea | modern | 0 | East_New_Britain | -4.63 | 152.3 | N.A. | N.A. | Skoglund_et_al_2016 | PASS |
| UV08 | Baining_Marabu | Oceania | Papua_New_Guinea | modern | 0 | East_New_Britain | -4.63 | 152.3 | N.A. | N.A. | Skoglund_et_al_2016 | PASS |
| UV11 | Baining_Marabu | Oceania | Papua_New_Guinea | modern | 0 | East_New_Britain | -4.63 | 152.3 | N.A. | N.A. | Skoglund_et_al_2016 | PASS |
| UV23 | Baining_Marabu | Oceania | Papua_New_Guinea | modern | 0 | East_New_Britain | -4.63 | 152.3 | N.A. | N.A. | Skoglund_et_al_2016 | PASS |
| UV24 | Baining_Marabu | Oceania | Papua_New_Guinea | modern | 0 | East_New_Britain | -4.63 | 152.3 | N.A. | N.A. | Skoglund_et_al_2016 | PASS |
| UV48 | Baining_Marabu | Oceania | Papua_New_Guinea | modern | 0 | East_New_Britain | -4.63 | 152.3 | N.A. | N.A. | Skoglund_et_al_2016 | PASS |
| UV73 | Baining_Marabu | Oceania | Papua_New_Guinea | modern | 0 | East_New_Britain | -4.63 | 152.3 | N.A. | N.A. | Skoglund_et_al_2016 | PASS |
| HGDP00655 | Nasioi | Oceania | Papua_New_Guinea | modern | 0 | South_Bougainville | -6.48 | 155.83 | N.A. | N.A. | Patterson_et_al_2012 | PASS |
| HGDP00656 | Nasioi | Oceania | Papua_New_Guinea | modern | 0 | South_Bougainville | -6.48 | 155.83 | N.A. | N.A. | Patterson_et_al_2012 | PASS |

| | | | | | | | | | | | | |
|---|---|---|---|---|---|---|---|---|---|---|---|---|
| HGDP00660 | Nasioi | Oceania | Papua_New_Guinea | modern | 0 | South_Bougainville | -6.48 | 155.83 | N.A. | N.A. | Patterson_et_al_2012 | PASS |
| HGDP00661 | Nasioi | Oceania | Papua_New_Guinea | modern | 0 | South_Bougainville | -6.48 | 155.83 | N.A. | N.A. | Patterson_et_al_2012 | PASS |
| HGDP00662 | Nasioi | Oceania | Papua_New_Guinea | modern | 0 | South_Bougainville | -6.48 | 155.83 | N.A. | N.A. | Patterson_et_al_2012 | PASS |
| HGDP00663 | Nasioi | Oceania | Papua_New_Guinea | modern | 0 | South_Bougainville | -6.48 | 155.83 | N.A. | N.A. | Patterson_et_al_2012 | PASS |
| HGDP00664 | Nasioi | Oceania | Papua_New_Guinea | modern | 0 | South_Bougainville | -6.48 | 155.83 | N.A. | N.A. | Patterson_et_al_2012 | PASS |
| HGDP00787 | Nasioi | Oceania | Papua_New_Guinea | modern | 0 | South_Bougainville | -6.48 | 155.83 | N.A. | N.A. | Patterson_et_al_2012 | PASS |
| HGDP00788 | Nasioi | Oceania | Papua_New_Guinea | modern | 0 | South_Bougainville | -6.48 | 155.83 | N.A. | N.A. | Patterson_et_al_2012 | PASS |
| HGDP01027 | Nasioi | Oceania | Papua_New_Guinea | modern | 0 | South_Bougainville | -6.48 | 155.83 | N.A. | N.A. | Patterson_et_al_2012 | PASS |
| NG18 | New_Guinea_Highlander | Oceania | Papua_New_Guinea | modern | 0 | Trans-New_Guinea | -6.15 | 143.66 | N.A. | N.A. | Qin_and_Stoneking_2015 | PASS |
| SH09 | New_Guinea_Highlander | Oceania | Papua_New_Guinea | modern | 0 | Trans-New_Guinea | -6.15 | 143.66 | N.A. | N.A. | Qin_and_Stoneking_2015 | PASS |
| NG20 | New_Guinea_Highlander | Oceania | Papua_New_Guinea | modern | 0 | Trans-New_Guinea | -6.15 | 143.66 | N.A. | N.A. | Qin_and_Stoneking_2015 | PASS |
| SH10 | New_Guinea_Highlander | Oceania | Papua_New_Guinea | modern | 0 | Trans-New_Guinea | -6.15 | 143.66 | N.A. | N.A. | Qin_and_Stoneking_2015 | PASS |
| NG4 | New_Guinea_Highlander | Oceania | Papua_New_Guinea | modern | 0 | Trans-New_Guinea | -6.15 | 143.66 | N.A. | N.A. | Qin_and_Stoneking_2015 | PASS |
| NG22 | New_Guinea_Highlander | Oceania | Papua_New_Guinea | modern | 0 | Trans-New_Guinea | -6.15 | 143.66 | N.A. | N.A. | Qin_and_Stoneking_2015 | PASS |
| SH18 | New_Guinea_Highlander | Oceania | Papua_New_Guinea | modern | 0 | Trans-New_Guinea | -6.15 | 143.66 | N.A. | N.A. | Qin_and_Stoneking_2015 | PASS |
| SH33 | New_Guinea_Highlander | Oceania | Papua_New_Guinea | modern | 0 | Trans-New_Guinea | -6.15 | 143.66 | N.A. | N.A. | Qin_and_Stoneking_2015 | PASS |
| NG25 | New_Guinea_Highlander | Oceania | Papua_New_Guinea | modern | 0 | Trans-New_Guinea | -6.15 | 143.66 | N.A. | N.A. | Qin_and_Stoneking_2015 | PASS |
| SH19 | New_Guinea_Highlander | Oceania | Papua_New_Guinea | modern | 0 | Trans-New_Guinea | -6.15 | 143.66 | N.A. | N.A. | Qin_and_Stoneking_2015 | PASS |
| NG7 | New_Guinea_Highlander | Oceania | Papua_New_Guinea | modern | 0 | Trans-New_Guinea | -6.15 | 143.66 | N.A. | N.A. | Qin_and_Stoneking_2015 | PASS |
| NG26 | New_Guinea_Highlander | Oceania | Papua_New_Guinea | modern | 0 | Trans-New_Guinea | -6.15 | 143.66 | N.A. | N.A. | Qin_and_Stoneking_2015 | PASS |
| SH20 | New_Guinea_Highlander | Oceania | Papua_New_Guinea | modern | 0 | Trans-New_Guinea | -6.15 | 143.66 | N.A. | N.A. | Qin_and_Stoneking_2015 | PASS |
| NG13 | New_Guinea_Highlander | Oceania | Papua_New_Guinea | modern | 0 | Trans-New_Guinea | -6.15 | 143.66 | N.A. | N.A. | Qin_and_Stoneking_2015 | PASS |
| SH05 | New_Guinea_Highlander | Oceania | Papua_New_Guinea | modern | 0 | Trans-New_Guinea | -6.15 | 143.66 | N.A. | N.A. | Qin_and_Stoneking_2015 | PASS |
| SH31 | New_Guinea_Highlander | Oceania | Papua_New_Guinea | modern | 0 | Trans-New_Guinea | -6.15 | 143.66 | N.A. | N.A. | Qin_and_Stoneking_2015 | PASS |
| SH07 | New_Guinea_Highlander | Oceania | Papua_New_Guinea | modern | 0 | Trans-New_Guinea | -6.15 | 143.66 | N.A. | N.A. | Qin_and_Stoneking_2015 | PASS |
| NG17 | New_Guinea_Highlander | Oceania | Papua_New_Guinea | modern | 0 | Trans-New_Guinea | -6.15 | 143.66 | N.A. | N.A. | Qin_and_Stoneking_2015 | PASS |
| SH08 | New_Guinea_Highlander | Oceania | Papua_New_Guinea | modern | 0 | Trans-New_Guinea | -6.15 | 143.66 | N.A. | N.A. | Qin_and_Stoneking_2015 | PASS |
| NG24 | Papuan_Eastern_Highlands | Oceania | Papua_New_Guinea | modern | 0 | Trans-New_Guinea | -6.1 | 145.4 | N.A. | N.A. | Qin_and_Stoneking_2015 | PASS |
| NG10 | Papuan_Eastern_Highlands | Oceania | Papua_New_Guinea | modern | 0 | Trans-New_Guinea | -6.1 | 145.4 | N.A. | N.A. | Qin_and_Stoneking_2015 | PASS |
| NG11 | Papuan_Eastern_Highlands | Oceania | Papua_New_Guinea | modern | 0 | Trans-New_Guinea | -6.1 | 145.4 | N.A. | N.A. | Qin_and_Stoneking_2015 | PASS |
| NG28 | Papuan_Eastern_Highlands | Oceania | Papua_New_Guinea | modern | 0 | Trans-New_Guinea | -6.1 | 145.4 | N.A. | N.A. | Qin_and_Stoneking_2015 | PASS |
| CP3 | Papuan_Central_Province | Oceania | Papua_New_Guinea | modern | 0 | Trans-New_Guinea | -9.48 | 147.2 | N.A. | N.A. | Qin_and_Stoneking_2015 | PASS |
| CP7 | Papuan_Central_Province | Oceania | Papua_New_Guinea | modern | 0 | Trans-New_Guinea | -9.48 | 147.2 | N.A. | N.A. | Qin_and_Stoneking_2015 | PASS |
| CP10 | Papuan_Central_Province | Oceania | Papua_New_Guinea | modern | 0 | Trans-New_Guinea | -9.48 | 147.2 | N.A. | N.A. | Qin_and_Stoneking_2015 | PASS |

| | | | | | | | | | | | | |
|---|---|---|---|---|---|---|---|---|---|---|---|---|
| CP13 | Papuan_Central_Province | Oceania | Papua_New_Guinea | modern | 0 | Trans-New_Guinea | -9.48 | 147.2 | N.A. | N.A. | Qin_and_Stoneking_2015 | PASS |
| CP14 | Papuan_Central_Province | Oceania | Papua_New_Guinea | modern | 0 | Trans-New_Guinea | -9.48 | 147.2 | N.A. | N.A. | Qin_and_Stoneking_2015 | PASS |
| GP3 | Papuan_Gulf_Province | Oceania | Papua_New_Guinea | modern | 0 | Trans-New_Guinea | -7.88 | 145.69 | N.A. | N.A. | Qin_and_Stoneking_2015 | PASS |
| GP4 | Papuan_Gulf_Province | Oceania | Papua_New_Guinea | modern | 0 | Trans-New_Guinea | -7.88 | 145.69 | N.A. | N.A. | Qin_and_Stoneking_2015 | PASS |
| GP6 | Papuan_Gulf_Province | Oceania | Papua_New_Guinea | modern | 0 | Trans-New_Guinea | -7.88 | 145.69 | N.A. | N.A. | Qin_and_Stoneking_2015 | PASS |
| Y4349 | Tongan | Oceania | Kingdom_of_Tonga | modern | 0 | Austronesian | -21.19 | -175.18 | Tongan | N.A. | Qin_and_Stoneking_2015 | PASS |
| Y4383 | Tongan | Oceania | Kingdom_of_Tonga | modern | 0 | Austronesian | -21.19 | -175.18 | Tongan | N.A. | Qin_and_Stoneking_2015 | PASS |
| Y4914 | Tongan | Oceania | Kingdom_of_Tonga | modern | 0 | Austronesian | -21.19 | -175.18 | Tongan | N.A. | Qin_and_Stoneking_2015 | PASS |
| Y6254 | Tongan | Oceania | Kingdom_of_Tonga | modern | 0 | Austronesian | -21.19 | -175.18 | Tongan | N.A. | Qin_and_Stoneking_2015 | PASS |
| Y5707 | Tongan | Oceania | Kingdom_of_Tonga | modern | 0 | Austronesian | -21.19 | -175.18 | Tongan | N.A. | Qin_and_Stoneking_2015 | PASS |
| Y5718 | Tongan | Oceania | Kingdom_of_Tonga | modern | 0 | Austronesian | -21.19 | -175.18 | Tongan | N.A. | Qin_and_Stoneking_2015 | PASS |
| VL43 | Vella_Lavella | Oceania | Solomon_Islands | modern | 0 | Central_Solomons | -7.9 | 156.7 | N.A. | N.A. | Qin_and_Stoneking_2015 | PASS |
| KO39 | Vella_Lavella | Oceania | Solomon_Islands | modern | 0 | Central_Solomons | -7.9 | 156.7 | N.A. | N.A. | Qin_and_Stoneking_2015 | PASS |
| VL02 | Vella_Lavella | Oceania | Solomon_Islands | modern | 0 | Central_Solomons | -7.9 | 156.7 | N.A. | N.A. | Qin_and_Stoneking_2015 | PASS |
| VL08 | Vella_Lavella | Oceania | Solomon_Islands | modern | 0 | Central_Solomons | -7.9 | 156.7 | N.A. | N.A. | Qin_and_Stoneking_2015 | PASS |
| VL09 | Vella_Lavella | Oceania | Solomon_Islands | modern | 0 | Central_Solomons | -7.9 | 156.7 | N.A. | N.A. | Qin_and_Stoneking_2015 | PASS |
| VL11 | Vella_Lavella | Oceania | Solomon_Islands | modern | 0 | Central_Solomons | -7.9 | 156.7 | N.A. | N.A. | Qin_and_Stoneking_2015 | PASS |
| PH10_1 | Mamanwa | ISEA | Philippines | modern | 0 | Austronesian | 9.76 | 125.51 | Mamanwa | N.A. | Qin_and_Stoneking_2015 | PASS |
| PH65_1 | Mamanwa | ISEA | Philippines | modern | 0 | Austronesian | 9.76 | 125.51 | Mamanwa | N.A. | Qin_and_Stoneking_2015 | PASS |
| PH2 | Mamanwa | ISEA | Philippines | modern | 0 | Austronesian | 9.76 | 125.51 | Mamanwa | N.A. | Qin_and_Stoneking_2015 | PASS |
| PH35_1 | Mamanwa | ISEA | Philippines | modern | 0 | Austronesian | 9.76 | 125.51 | Mamanwa | N.A. | Qin_and_Stoneking_2015 | PASS |
| PH38_1 | Mamanwa | ISEA | Philippines | modern | 0 | Austronesian | 9.76 | 125.51 | Mamanwa | N.A. | Qin_and_Stoneking_2015 | PASS |
| PH4 | Mamanwa | ISEA | Philippines | modern | 0 | Austronesian | 9.76 | 125.51 | Mamanwa | N.A. | Qin_and_Stoneking_2015 | PASS |
| PH45_1 | Mamanwa | ISEA | Philippines | modern | 0 | Austronesian | 9.76 | 125.51 | Mamanwa | N.A. | Qin_and_Stoneking_2015 | PASS |
| PH66_2 | Mamanwa | ISEA | Philippines | modern | 0 | Austronesian | 9.76 | 125.51 | Mamanwa | N.A. | Qin_and_Stoneking_2015 | PASS |
| PH67_2 | Mamanwa | ISEA | Philippines | modern | 0 | Austronesian | 9.76 | 125.51 | Mamanwa | N.A. | Qin_and_Stoneking_2015 | PASS |
| PH72_3 | Mamanwa | ISEA | Philippines | modern | 0 | Austronesian | 9.76 | 125.51 | Mamanwa | N.A. | Qin_and_Stoneking_2015 | PASS |
| PH77_1 | Mamanwa | ISEA | Philippines | modern | 0 | Austronesian | 9.76 | 125.51 | Mamanwa | N.A. | Qin_and_Stoneking_2015 | PASS |
| PH8 | Mamanwa | ISEA | Philippines | modern | 0 | Austronesian | 9.76 | 125.51 | Mamanwa | N.A. | Qin_and_Stoneking_2015 | PASS |
| PH9 | Mamanwa | ISEA | Philippines | modern | 0 | Austronesian | 9.76 | 125.51 | Mamanwa | N.A. | Qin_and_Stoneking_2015 | PASS |
| PH50_1 | Mamanwa1 | ISEA | Philippines | modern | 0 | Austronesian | 9.76 | 125.51 | Mamanwa | N.A. | Qin_and_Stoneking_2015 | PASS |
| PH52_3 | Mamanwa1 | ISEA | Philippines | modern | 0 | Austronesian | 9.76 | 125.51 | Mamanwa | N.A. | Qin_and_Stoneking_2015 | PASS |
| PH53_1 | Mamanwa1 | ISEA | Philippines | modern | 0 | Austronesian | 9.76 | 125.51 | Mamanwa | N.A. | Qin_and_Stoneking_2015 | PASS |
| PH55_2 | Mamanwa1 | ISEA | Philippines | modern | 0 | Austronesian | 9.76 | 125.51 | Mamanwa | N.A. | Qin_and_Stoneking_2015 | PASS |
| PH56_1 | Mamanwa1 | ISEA | Philippines | modern | 0 | Austronesian | 9.76 | 125.51 | Mamanwa | N.A. | Qin_and_Stoneking_2015 | PASS |
| Smd1_1 | Semende | ISEA | Indonesia | modern | 0 | Austronesian | 3.58 | 98.67 | Semende | N.A. | Qin_and_Stoneking_2015 | PASS |
| Smd10_1 | Semende | ISEA | Indonesia | modern | 0 | Austronesian | 3.58 | 98.67 | Semende | N.A. | Qin_and_Stoneking_2015 | PASS |
| Smd14_1 | Semende | ISEA | Indonesia | modern | 0 | Austronesian | 3.58 | 98.67 | Semende | N.A. | Qin_and_Stoneking_2015 | PASS |
| Smd15_1 | Semende | ISEA | Indonesia | modern | 0 | Austronesian | 3.58 | 98.67 | Semende | N.A. | Qin_and_Stoneking_2015 | PASS |
| Smd18_1 | Semende | ISEA | Indonesia | modern | 0 | Austronesian | 3.58 | 98.67 | Semende | N.A. | Qin_and_Stoneking_2015 | PASS |

| | | | | | | | | | | | | |
|---|---|---|---|---|---|---|---|---|---|---|---|---|
| Smd19_1 | Semende | ISEA | Indonesia | modern | 0 | Austronesian | 3.58 | 98.67 | Semende | N.A. | Qin_and_Stoneking_2015 | PASS |
| Smd3_1 | Semende | ISEA | Indonesia | modern | 0 | Austronesian | 3.58 | 98.67 | Semende | N.A. | Qin_and_Stoneking_2015 | PASS |
| Smd5_1 | Semende | ISEA | Indonesia | modern | 0 | Austronesian | 3.58 | 98.67 | Semende | N.A. | Qin_and_Stoneking_2015 | PASS |
| Smd6_1 | Semende | ISEA | Indonesia | modern | 0 | Austronesian | 3.58 | 98.67 | Semende | N.A. | Qin_and_Stoneking_2015 | PASS |
| Smd9_1 | Semende | ISEA | Indonesia | modern | 0 | Austronesian | 3.58 | 98.67 | Semende | N.A. | Qin_and_Stoneking_2015 | PASS |
| Bor1_5 | Borneo | ISEA | Indonesia | modern | 0 | Austronesian | 0.18 | 110.16 | Borneo | N.A. | Qin_and_Stoneking_2015 | PASS |
| Bor16_1 | Borneo | ISEA | Indonesia | modern | 0 | Austronesian | 0.18 | 110.16 | Borneo | N.A. | Qin_and_Stoneking_2015 | PASS |
| Bor2_3 | Borneo | ISEA | Indonesia | modern | 0 | Austronesian | 0.18 | 110.16 | Borneo | N.A. | Qin_and_Stoneking_2015 | PASS |
| Bor20_1 | Borneo | ISEA | Indonesia | modern | 0 | Austronesian | 0.18 | 110.16 | Borneo | N.A. | Qin_and_Stoneking_2015 | PASS |
| Bor3_3 | Borneo | ISEA | Indonesia | modern | 0 | Austronesian | 0.18 | 110.16 | Borneo | N.A. | Qin_and_Stoneking_2015 | PASS |
| Bor4_5 | Borneo | ISEA | Indonesia | modern | 0 | Austronesian | 0.18 | 110.16 | Borneo | N.A. | Qin_and_Stoneking_2015 | PASS |
| Bor6_1 | Borneo | ISEA | Indonesia | modern | 0 | Austronesian | 0.18 | 110.16 | Borneo | N.A. | Qin_and_Stoneking_2015 | PASS |
| Bor7_2 | Borneo | ISEA | Indonesia | modern | 0 | Austronesian | 0.18 | 110.16 | Borneo | N.A. | Qin_and_Stoneking_2015 | PASS |
| Bor9_3 | Borneo | ISEA | Indonesia | modern | 0 | Austronesian | 0.18 | 110.16 | Borneo | N.A. | Qin_and_Stoneking_2015 | PASS |
| Dusun4 | Dusun | ISEA | Brunei | modern | 0 | Austronesian | 4.71 | 114.67 | Dusun | N.A. | Skoglund_et_al_2016 | PASS |
| Dusun7 | Dusun | ISEA | Brunei | modern | 0 | Austronesian | 4.71 | 114.67 | Dusun | N.A. | Skoglund_et_al_2016 | PASS |
| Dusun8 | Dusun | ISEA | Brunei | modern | 0 | Austronesian | 4.71 | 114.67 | Dusun | N.A. | Skoglund_et_al_2016 | PASS |
| Dusun10 | Dusun | ISEA | Brunei | modern | 0 | Austronesian | 4.71 | 114.67 | Dusun | N.A. | Skoglund_et_al_2016 | PASS |
| Dusun11 | Dusun | ISEA | Brunei | modern | 0 | Austronesian | 4.71 | 114.67 | Dusun | N.A. | Skoglund_et_al_2016 | PASS |
| Dusun12 | Dusun | ISEA | Brunei | modern | 0 | Austronesian | 4.71 | 114.67 | Dusun | N.A. | Skoglund_et_al_2016 | PASS |
| Dusun14 | Dusun | ISEA | Brunei | modern | 0 | Austronesian | 4.71 | 114.67 | Dusun | N.A. | Skoglund_et_al_2016 | PASS |
| Dusun21 | Dusun | ISEA | Brunei | modern | 0 | Austronesian | 4.71 | 114.67 | Dusun | N.A. | Skoglund_et_al_2016 | PASS |
| Dusun22 | Dusun | ISEA | Brunei | modern | 0 | Austronesian | 4.71 | 114.67 | Dusun | N.A. | Skoglund_et_al_2016 | PASS |
| Dusun23 | Dusun | ISEA | Brunei | modern | 0 | Austronesian | 4.71 | 114.67 | Dusun | N.A. | Skoglund_et_al_2016 | PASS |
| Murut2 | Murut | ISEA | Brunei | modern | 0 | Austronesian | 4.62 | 115.14 | Murut | N.A. | Skoglund_et_al_2016 | PASS |
| Murut3 | Murut | ISEA | Brunei | modern | 0 | Austronesian | 4.62 | 115.14 | Murut | N.A. | Skoglund_et_al_2016 | PASS |
| Murut4 | Murut | ISEA | Brunei | modern | 0 | Austronesian | 4.62 | 115.14 | Murut | N.A. | Skoglund_et_al_2016 | PASS |
| Murut5 | Murut | ISEA | Brunei | modern | 0 | Austronesian | 4.62 | 115.14 | Murut | N.A. | Skoglund_et_al_2016 | PASS |
| Murut6 | Murut | ISEA | Brunei | modern | 0 | Austronesian | 4.62 | 115.14 | Murut | N.A. | Skoglund_et_al_2016 | PASS |
| Murut9 | Murut | ISEA | Brunei | modern | 0 | Austronesian | 4.62 | 115.14 | Murut | N.A. | Skoglund_et_al_2016 | PASS |
| Murut11 | Murut | ISEA | Brunei | modern | 0 | Austronesian | 4.62 | 115.14 | Murut | N.A. | Skoglund_et_al_2016 | PASS |
| Murut13 | Murut | ISEA | Brunei | modern | 0 | Austronesian | 4.62 | 115.14 | Murut | N.A. | Skoglund_et_al_2016 | PASS |
| Murut15 | Murut | ISEA | Brunei | modern | 0 | Austronesian | 4.62 | 115.14 | Murut | N.A. | Skoglund_et_al_2016 | PASS |
| Murut17 | Murut | ISEA | Brunei | modern | 0 | Austronesian | 4.62 | 115.14 | Murut | N.A. | Skoglund_et_al_2016 | PASS |
| Burm4 | Burmese | MSEA | Myanmar | modern | 0 | Sino-Tibetan | 16.41 | 95.89 | Burmese | N.A. | Skoglund_et_al_2016 | PASS |
| Burm7 | Burmese | MSEA | Myanmar | modern | 0 | Sino-Tibetan | 16.41 | 95.89 | Burmese | N.A. | Skoglund_et_al_2016 | PASS |
| Burm8 | Burmese | MSEA | Myanmar | modern | 0 | Sino-Tibetan | 16.41 | 95.89 | Burmese | N.A. | Skoglund_et_al_2016 | PASS |
| Burm9 | Burmese | MSEA | Myanmar | modern | 0 | Sino-Tibetan | 16.41 | 95.89 | Burmese | N.A. | Skoglund_et_al_2016 | PASS |
| Burm10 | Burmese | MSEA | Myanmar | modern | 0 | Sino-Tibetan | 16.41 | 95.89 | Burmese | N.A. | Skoglund_et_al_2016 | PASS |
| Burm12 | Burmese | MSEA | Myanmar | modern | 0 | Sino-Tibetan | 16.41 | 95.89 | Burmese | N.A. | Skoglund_et_al_2016 | PASS |
| Burm13 | Burmese | MSEA | Myanmar | modern | 0 | Sino-Tibetan | 16.41 | 95.89 | Burmese | N.A. | Skoglund_et_al_2016 | PASS |

| | | | | | | | | | | | | |
|---|---|---|---|---|---|---|---|---|---|---|---|---|
| Burm14 | Burmese | MSEA | Myanmar | modern | 0 | Sino-Tibetan | 16.41 | 95.89 | Burmese | N.A. | Skoglund_et_al_2016 | PASS |
| Burm15 | Burmese | MSEA | Myanmar | modern | 0 | Sino-Tibetan | 16.41 | 95.89 | Burmese | N.A. | Skoglund_et_al_2016 | PASS |
| Burm16 | Burmese | MSEA | Myanmar | modern | 0 | Sino-Tibetan | 16.41 | 95.89 | Burmese | N.A. | Skoglund_et_al_2016 | PASS |
| Igorot3 | Kankanaey | ISEA | Philippines | modern | 0 | Austronesian | 17.07 | 121.03 | Kankanaey | N.A. | Skoglund_et_al_2016 | PASS |
| Igorot4 | Kankanaey | ISEA | Philippines | modern | 0 | Austronesian | 17.07 | 121.03 | Kankanaey | N.A. | Skoglund_et_al_2016 | PASS |
| Igorot5 | Kankanaey | ISEA | Philippines | modern | 0 | Austronesian | 17.07 | 121.03 | Kankanaey | N.A. | Skoglund_et_al_2016 | PASS |
| Igorot6 | Kankanaey | ISEA | Philippines | modern | 0 | Austronesian | 17.07 | 121.03 | Kankanaey | N.A. | Skoglund_et_al_2016 | PASS |
| Igorot10 | Kankanaey | ISEA | Philippines | modern | 0 | Austronesian | 17.07 | 121.03 | Kankanaey | N.A. | Skoglund_et_al_2016 | PASS |
| Igorot11 | Kankanaey | ISEA | Philippines | modern | 0 | Austronesian | 17.07 | 121.03 | Kankanaey | N.A. | Skoglund_et_al_2016 | PASS |
| Igorot13 | Kankanaey | ISEA | Philippines | modern | 0 | Austronesian | 17.07 | 121.03 | Kankanaey | N.A. | Skoglund_et_al_2016 | PASS |
| Igorot17 | Kankanaey | ISEA | Philippines | modern | 0 | Austronesian | 17.07 | 121.03 | Kankanaey | N.A. | Skoglund_et_al_2016 | PASS |
| Igorot21 | Kankanaey | ISEA | Philippines | modern | 0 | Austronesian | 17.07 | 121.03 | Kankanaey | N.A. | Skoglund_et_al_2016 | PASS |
| Igorot23 | Kankanaey | ISEA | Philippines | modern | 0 | Austronesian | 17.07 | 121.03 | Kankanaey | N.A. | Skoglund_et_al_2016 | PASS |
| Luz2 | Ilocano | ISEA | Philippines | modern | 0 | Austronesian | 14.6 | 120.98 | Ilocano | N.A. | Skoglund_et_al_2016 | PASS |
| Luz6 | Ilocano | ISEA | Philippines | modern | 0 | Austronesian | 14.6 | 120.98 | Ilocano | N.A. | Skoglund_et_al_2016 | PASS |
| Luz5 | Tagalog | ISEA | Philippines | modern | 0 | Austronesian | 14.6 | 120.98 | Tagalog | N.A. | Skoglund_et_al_2016 | PASS |
| Luz8 | Tagalog | ISEA | Philippines | modern | 0 | Austronesian | 14.6 | 120.98 | Tagalog | N.A. | Skoglund_et_al_2016 | PASS |
| Luz9 | Tagalog | ISEA | Philippines | modern | 0 | Austronesian | 14.6 | 120.98 | Tagalog | N.A. | Skoglund_et_al_2016 | PASS |
| Luz10 | Tagalog | ISEA | Philippines | modern | 0 | Austronesian | 14.6 | 120.98 | Tagalog | N.A. | Skoglund_et_al_2016 | PASS |
| Luz11 | Tagalog | ISEA | Philippines | modern | 0 | Austronesian | 14.6 | 120.98 | Tagalog | N.A. | Skoglund_et_al_2016 | PASS |
| Vizaya1 | Visayan | ISEA | Philippines | modern | 0 | Austronesian | 9.76 | 125.51 | Visayan | N.A. | Skoglund_et_al_2016 | PASS |
| Vizaya2 | Visayan | ISEA | Philippines | modern | 0 | Austronesian | 9.76 | 125.51 | Visayan | N.A. | Skoglund_et_al_2016 | PASS |
| Vizaya3 | Visayan | ISEA | Philippines | modern | 0 | Austronesian | 9.76 | 125.51 | Visayan | N.A. | Skoglund_et_al_2016 | PASS |
| Vizaya4 | Visayan | ISEA | Philippines | modern | 0 | Austronesian | 9.76 | 125.51 | Visayan | N.A. | Skoglund_et_al_2016 | PASS |
| Malay2 | Malay | ISEA | Singapore | modern | 0 | Austronesian | 1.35 | 103.82 | Malay | N.A. | Skoglund_et_al_2016 | PASS |
| Malay5 | Malay | ISEA | Singapore | modern | 0 | Austronesian | 1.35 | 103.82 | Malay | N.A. | Skoglund_et_al_2016 | PASS |
| Malay6 | Malay | ISEA | Singapore | modern | 0 | Austronesian | 1.35 | 103.82 | Malay | N.A. | Skoglund_et_al_2016 | PASS |
| Malay10 | Malay | ISEA | Singapore | modern | 0 | Austronesian | 1.35 | 103.82 | Malay | N.A. | Skoglund_et_al_2016 | PASS |
| Malay11 | Malay | ISEA | Singapore | modern | 0 | Austronesian | 1.35 | 103.82 | Malay | N.A. | Skoglund_et_al_2016 | PASS |
| Malay13 | Malay | ISEA | Singapore | modern | 0 | Austronesian | 1.35 | 103.82 | Malay | N.A. | Skoglund_et_al_2016 | PASS |
| Malay14 | Malay | ISEA | Singapore | modern | 0 | Austronesian | 1.35 | 103.82 | Malay | N.A. | Skoglund_et_al_2016 | PASS |
| Malay20 | Malay | ISEA | Singapore | modern | 0 | Austronesian | 1.35 | 103.82 | Malay | N.A. | Skoglund_et_al_2016 | PASS |
| Malay21 | Malay | ISEA | Singapore | modern | 0 | Austronesian | 1.35 | 103.82 | Malay | N.A. | Skoglund_et_al_2016 | PASS |
| NA13607 | Amis_HO | sEA | Taiwan | modern | 0 | Austronesian | 22.8 | 121.2 | Amis | TW_S | Lazaridis_et_al_2014 | PASS |
| NA13608 | Amis_HO | sEA | Taiwan | modern | 0 | Austronesian | 22.8 | 121.2 | Amis | TW_S | Lazaridis_et_al_2014 | PASS |
| NA13609 | Amis_HO | sEA | Taiwan | modern | 0 | Austronesian | 22.8 | 121.2 | Amis | TW_S | Lazaridis_et_al_2014 | PASS |
| NA13610 | Amis_HO | sEA | Taiwan | modern | 0 | Austronesian | 22.8 | 121.2 | Amis | TW_S | Lazaridis_et_al_2014 | PASS |
| NA13611 | Amis_HO | sEA | Taiwan | modern | 0 | Austronesian | 22.8 | 121.2 | Amis | TW_S | Lazaridis_et_al_2014 | PASS |
| NA13612 | Amis_HO | sEA | Taiwan | modern | 0 | Austronesian | 22.8 | 121.2 | Amis | TW_S | Lazaridis_et_al_2014 | PASS |
| NA13613 | Amis_HO | sEA | Taiwan | modern | 0 | Austronesian | 22.8 | 121.2 | Amis | TW_S | Lazaridis_et_al_2014 | PASS |
| NA13614 | Amis_HO | sEA | Taiwan | modern | 0 | Austronesian | 22.8 | 121.2 | Amis | TW_S | Lazaridis_et_al_2014 | PASS |

| | | | | | | | | | | | | |
|---|---|---|---|---|---|---|---|---|---|---|---|---|
| NA13615 | Amis_HO | sEA | Taiwan | modern | 0 | Austronesian | 22.8 | 121.2 | Amis | TW_S | Lazaridis_et_al_2014 | PASS |
| NA13616 | Amis_HO | sEA | Taiwan | modern | 0 | Austronesian | 22.8 | 121.2 | Amis | TW_S | Lazaridis_et_al_2014 | PASS |
| NA13597 | Atayal_HO | sEA | Taiwan | modern | 0 | Austronesian | 24.6 | 121.3 | Atayal | TW_N | Lazaridis_et_al_2014 | PASS |
| NA13598 | Atayal_HO | sEA | Taiwan | modern | 0 | Austronesian | 24.6 | 121.3 | Atayal | TW_N | Lazaridis_et_al_2014 | PASS |
| NA13599 | Atayal_HO | sEA | Taiwan | modern | 0 | Austronesian | 24.6 | 121.3 | Atayal | TW_N | Lazaridis_et_al_2014 | PASS |
| NA13600 | Atayal_HO | sEA | Taiwan | modern | 0 | Austronesian | 24.6 | 121.3 | Atayal | TW_N | Lazaridis_et_al_2014 | PASS |
| NA13601 | Atayal_HO | sEA | Taiwan | modern | 0 | Austronesian | 24.6 | 121.3 | Atayal | TW_N | Lazaridis_et_al_2014 | PASS |
| NA13602 | Atayal_HO | sEA | Taiwan | modern | 0 | Austronesian | 24.6 | 121.3 | Atayal | TW_N | Lazaridis_et_al_2014 | PASS |
| NA13603 | Atayal_HO | sEA | Taiwan | modern | 0 | Austronesian | 24.6 | 121.3 | Atayal | TW_N | Lazaridis_et_al_2014 | PASS |
| NA13604 | Atayal_HO | sEA | Taiwan | modern | 0 | Austronesian | 24.6 | 121.3 | Atayal | TW_N | Lazaridis_et_al_2014 | PASS |
| NA13605 | Atayal_HO | sEA | Taiwan | modern | 0 | Austronesian | 24.6 | 121.3 | Atayal | TW_N | Lazaridis_et_al_2014 | PASS |
| NA13606 | Atayal_HO | sEA | Taiwan | modern | 0 | Austronesian | 24.6 | 121.3 | Atayal | TW_N | Lazaridis_et_al_2014 | PASS |
| HGDP00711 | Cambodian | MSEA | Cambodia | modern | 0 | Austroasiatic | 12 | 105 | N.A. | N.A. | Patterson_et_al_2012 | PASS |
| HGDP00712 | Cambodian | MSEA | Cambodia | modern | 0 | Austroasiatic | 12 | 105 | N.A. | N.A. | Patterson_et_al_2012 | PASS |
| HGDP00713 | Cambodian | MSEA | Cambodia | modern | 0 | Austroasiatic | 12 | 105 | N.A. | N.A. | Patterson_et_al_2012 | PASS |
| HGDP00714 | Cambodian | MSEA | Cambodia | modern | 0 | Austroasiatic | 12 | 105 | N.A. | N.A. | Patterson_et_al_2012 | PASS |
| HGDP00715 | Cambodian | MSEA | Cambodia | modern | 0 | Austroasiatic | 12 | 105 | N.A. | N.A. | Patterson_et_al_2012 | PASS |
| HGDP00716 | Cambodian | MSEA | Cambodia | modern | 0 | Austroasiatic | 12 | 105 | N.A. | N.A. | Patterson_et_al_2012 | PASS |
| HGDP00717 | Cambodian | MSEA | Cambodia | modern | 0 | Austroasiatic | 12 | 105 | N.A. | N.A. | Patterson_et_al_2012 | PASS |
| HGDP00719 | Cambodian | MSEA | Cambodia | modern | 0 | Austroasiatic | 12 | 105 | N.A. | N.A. | Patterson_et_al_2012 | PASS |
| HGDP00720 | Cambodian | MSEA | Cambodia | modern | 0 | Austroasiatic | 12 | 105 | N.A. | N.A. | Patterson_et_al_2012 | PASS |
| HGDP00721 | Cambodian | MSEA | Cambodia | modern | 0 | Austroasiatic | 12 | 105 | N.A. | N.A. | Patterson_et_al_2012 | PASS |
| HG01842 | Kinh | MSEA | Vietnam | modern | 0 | Austroasiatic | 21 | 105.9 | N.A. | N.A. | Patterson_et_al_2012 | PASS |
| HG01844 | Kinh | MSEA | Vietnam | modern | 0 | Austroasiatic | 21 | 105.9 | N.A. | N.A. | Patterson_et_al_2012 | PASS |
| HG01846 | Kinh | MSEA | Vietnam | modern | 0 | Austroasiatic | 21 | 105.9 | N.A. | N.A. | Patterson_et_al_2012 | PASS |
| HG01600 | Kinh | MSEA | Vietnam | modern | 0 | Austroasiatic | 21 | 105.9 | N.A. | N.A. | Patterson_et_al_2012 | PASS |
| HG01595 | Kinh | MSEA | Vietnam | modern | 0 | Austroasiatic | 21 | 105.9 | N.A. | N.A. | Patterson_et_al_2012 | PASS |
| HG01596 | Kinh | MSEA | Vietnam | modern | 0 | Austroasiatic | 21 | 105.9 | N.A. | N.A. | Patterson_et_al_2012 | PASS |
| HG01597 | Kinh | MSEA | Vietnam | modern | 0 | Austroasiatic | 21 | 105.9 | N.A. | N.A. | Patterson_et_al_2012 | PASS |
| HG01598 | Kinh | MSEA | Vietnam | modern | 0 | Austroasiatic | 21 | 105.9 | N.A. | N.A. | Patterson_et_al_2012 | PASS |
| HGDP01307 | Dai | sEA | China | modern | 0 | Tai-Kadai | 21 | 100 | Dai | N.A. | Patterson_et_al_2012 | PASS |
| HGDP01308 | Dai | sEA | China | modern | 0 | Tai-Kadai | 21 | 100 | Dai | N.A. | Patterson_et_al_2012 | PASS |
| HGDP01309 | Dai | sEA | China | modern | 0 | Tai-Kadai | 21 | 100 | Dai | N.A. | Patterson_et_al_2012 | PASS |
| HGDP01310 | Dai | sEA | China | modern | 0 | Tai-Kadai | 21 | 100 | Dai | N.A. | Patterson_et_al_2012 | PASS |
| HGDP01311 | Dai | sEA | China | modern | 0 | Tai-Kadai | 21 | 100 | Dai | N.A. | Patterson_et_al_2012 | PASS |
| HGDP01312 | Dai | sEA | China | modern | 0 | Tai-Kadai | 21 | 100 | Dai | N.A. | Patterson_et_al_2012 | PASS |
| HGDP01313 | Dai | sEA | China | modern | 0 | Tai-Kadai | 21 | 100 | Dai | N.A. | Patterson_et_al_2012 | PASS |
| HGDP01314 | Dai | sEA | China | modern | 0 | Tai-Kadai | 21 | 100 | Dai | N.A. | Patterson_et_al_2012 | PASS |
| HGDP01315 | Dai | sEA | China | modern | 0 | Tai-Kadai | 21 | 100 | Dai | N.A. | Patterson_et_al_2012 | PASS |
| HGDP01316 | Dai | sEA | China | modern | 0 | Tai-Kadai | 21 | 100 | Dai | N.A. | Patterson_et_al_2012 | PASS |
| HGDP01189 | Miao | sEA | China | modern | 0 | Hmong-Mien | 28 | 109 | N.A. | N.A. | Patterson_et_al_2012 | PASS |

| | | | | | | | | | | | | |
|---|---|---|---|---|---|---|---|---|---|---|---|---|
| HGDP01190 | Miao | sEA | China | modern | 0 | Hmong-Mien | 28 | 109 | N.A. | N.A. | Patterson_et_al_2012 | PASS |
| HGDP01191 | Miao | sEA | China | modern | 0 | Hmong-Mien | 28 | 109 | N.A. | N.A. | Patterson_et_al_2012 | PASS |
| HGDP01192 | Miao | sEA | China | modern | 0 | Hmong-Mien | 28 | 109 | N.A. | N.A. | Patterson_et_al_2012 | PASS |
| HGDP01193 | Miao | sEA | China | modern | 0 | Hmong-Mien | 28 | 109 | N.A. | N.A. | Patterson_et_al_2012 | PASS |
| HGDP01194 | Miao | sEA | China | modern | 0 | Hmong-Mien | 28 | 109 | N.A. | N.A. | Patterson_et_al_2012 | PASS |
| HGDP01195 | Miao | sEA | China | modern | 0 | Hmong-Mien | 28 | 109 | N.A. | N.A. | Patterson_et_al_2012 | PASS |
| HGDP01196 | Miao | sEA | China | modern | 0 | Hmong-Mien | 28 | 109 | N.A. | N.A. | Patterson_et_al_2012 | PASS |
| HGDP01197 | Miao | sEA | China | modern | 0 | Hmong-Mien | 28 | 109 | N.A. | N.A. | Patterson_et_al_2012 | PASS |
| HGDP01198 | Miao | sEA | China | modern | 0 | Hmong-Mien | 28 | 109 | N.A. | N.A. | Patterson_et_al_2012 | PASS |
| HGDP01327 | She | sEA | China | modern | 0 | Hmong-Mien | 27 | 119 | N.A. | N.A. | Patterson_et_al_2012 | PASS |
| HGDP01328 | She | sEA | China | modern | 0 | Hmong-Mien | 27 | 119 | N.A. | N.A. | Patterson_et_al_2012 | PASS |
| HGDP01329 | She | sEA | China | modern | 0 | Hmong-Mien | 27 | 119 | N.A. | N.A. | Patterson_et_al_2012 | PASS |
| HGDP01330 | She | sEA | China | modern | 0 | Hmong-Mien | 27 | 119 | N.A. | N.A. | Patterson_et_al_2012 | PASS |
| HGDP01331 | She | sEA | China | modern | 0 | Hmong-Mien | 27 | 119 | N.A. | N.A. | Patterson_et_al_2012 | PASS |
| HGDP01332 | She | sEA | China | modern | 0 | Hmong-Mien | 27 | 119 | N.A. | N.A. | Patterson_et_al_2012 | PASS |
| HGDP01333 | She | sEA | China | modern | 0 | Hmong-Mien | 27 | 119 | N.A. | N.A. | Patterson_et_al_2012 | PASS |
| HGDP01334 | She | sEA | China | modern | 0 | Hmong-Mien | 27 | 119 | N.A. | N.A. | Patterson_et_al_2012 | PASS |
| HGDP01335 | She | sEA | China | modern | 0 | Hmong-Mien | 27 | 119 | N.A. | N.A. | Patterson_et_al_2012 | PASS |
| HGDP01336 | She | sEA | China | modern | 0 | Hmong-Mien | 27 | 119 | N.A. | N.A. | Patterson_et_al_2012 | PASS |
| HGDP01317 | Lahu_C | sEA | China | modern | 0 | Sino-Tibetan | 22 | 100 | Lahu | N.A. | Patterson_et_al_2012 | PASS |
| HGDP01318 | Lahu_C | sEA | China | modern | 0 | Sino-Tibetan | 22 | 100 | Lahu | N.A. | Patterson_et_al_2012 | PASS |
| HGDP01319 | Lahu_C | sEA | China | modern | 0 | Sino-Tibetan | 22 | 100 | Lahu | N.A. | Patterson_et_al_2012 | PASS |
| HGDP01320 | Lahu_C | sEA | China | modern | 0 | Sino-Tibetan | 22 | 100 | Lahu | N.A. | Patterson_et_al_2012 | PASS |
| HGDP01321 | Lahu_C | sEA | China | modern | 0 | Sino-Tibetan | 22 | 100 | Lahu | N.A. | Patterson_et_al_2012 | PASS |
| HGDP01322 | Lahu_C | sEA | China | modern | 0 | Sino-Tibetan | 22 | 100 | Lahu | N.A. | Patterson_et_al_2012 | PASS |
| HGDP01323 | Lahu_C | sEA | China | modern | 0 | Sino-Tibetan | 22 | 100 | Lahu | N.A. | Patterson_et_al_2012 | PASS |
| HGDP01326 | Lahu_C | sEA | China | modern | 0 | Sino-Tibetan | 22 | 100 | Lahu | N.A. | Patterson_et_al_2012 | PASS |
| HGDP01179 | Yi | sEA | China | modern | 0 | Sino-Tibetan | 28 | 103 | Yi | N.A. | Patterson_et_al_2012 | PASS |
| HGDP01180 | Yi | sEA | China | modern | 0 | Sino-Tibetan | 28 | 103 | Yi | N.A. | Patterson_et_al_2012 | PASS |
| HGDP01181 | Yi | sEA | China | modern | 0 | Sino-Tibetan | 28 | 103 | Yi | N.A. | Patterson_et_al_2012 | PASS |
| HGDP01182 | Yi | sEA | China | modern | 0 | Sino-Tibetan | 28 | 103 | Yi | N.A. | Patterson_et_al_2012 | PASS |
| HGDP01183 | Yi | sEA | China | modern | 0 | Sino-Tibetan | 28 | 103 | Yi | N.A. | Patterson_et_al_2012 | PASS |
| HGDP01184 | Yi | sEA | China | modern | 0 | Sino-Tibetan | 28 | 103 | Yi | N.A. | Patterson_et_al_2012 | PASS |
| HGDP01185 | Yi | sEA | China | modern | 0 | Sino-Tibetan | 28 | 103 | Yi | N.A. | Patterson_et_al_2012 | PASS |
| HGDP01186 | Yi | sEA | China | modern | 0 | Sino-Tibetan | 28 | 103 | Yi | N.A. | Patterson_et_al_2012 | PASS |
| HGDP01187 | Yi | sEA | China | modern | 0 | Sino-Tibetan | 28 | 103 | Yi | N.A. | Patterson_et_al_2012 | PASS |
| HGDP01188 | Yi | sEA | China | modern | 0 | Sino-Tibetan | 28 | 103 | Yi | N.A. | Patterson_et_al_2012 | PASS |
| HGDP01337 | Naxi | sEA | China | modern | 0 | Sino-Tibetan | 26 | 100 | Naxi | N.A. | Patterson_et_al_2012 | PASS |
| HGDP01338 | Naxi | sEA | China | modern | 0 | Sino-Tibetan | 26 | 100 | Naxi | N.A. | Patterson_et_al_2012 | PASS |
| HGDP01339 | Naxi | sEA | China | modern | 0 | Sino-Tibetan | 26 | 100 | Naxi | N.A. | Patterson_et_al_2012 | PASS |
| HGDP01340 | Naxi | sEA | China | modern | 0 | Sino-Tibetan | 26 | 100 | Naxi | N.A. | Patterson_et_al_2012 | PASS |

| | | | | | | | | | | | | |
|---|---|---|---|---|---|---|---|---|---|---|---|---|
| HGDP01341 | Naxi | sEA | China | modern | 0 | Sino-Tibetan | 26 | 100 | Naxi | N.A. | Patterson_et_al_2012 | PASS |
| HGDP01342 | Naxi | sEA | China | modern | 0 | Sino-Tibetan | 26 | 100 | Naxi | N.A. | Patterson_et_al_2012 | PASS |
| HGDP01344 | Naxi | sEA | China | modern | 0 | Sino-Tibetan | 26 | 100 | Naxi | N.A. | Patterson_et_al_2012 | PASS |
| HGDP01345 | Naxi | sEA | China | modern | 0 | Sino-Tibetan | 26 | 100 | Naxi | N.A. | Patterson_et_al_2012 | PASS |
| HGDP01346 | Naxi | sEA | China | modern | 0 | Sino-Tibetan | 26 | 100 | Naxi | N.A. | Patterson_et_al_2012 | PASS |
| HGDP00774 | Han | sEA | China | modern | 0 | Sino-Tibetan | 32.27 | 114 | Han | N.A. | Patterson_et_al_2012 | PASS |
| HGDP00775 | Han | sEA | China | modern | 0 | Sino-Tibetan | 32.27 | 114 | Han | N.A. | Patterson_et_al_2012 | PASS |
| HGDP00776 | Han | sEA | China | modern | 0 | Sino-Tibetan | 32.27 | 114 | Han | N.A. | Patterson_et_al_2012 | PASS |
| HGDP00777 | Han | sEA | China | modern | 0 | Sino-Tibetan | 32.27 | 114 | Han | N.A. | Patterson_et_al_2012 | PASS |
| HGDP00779 | Han | sEA | China | modern | 0 | Sino-Tibetan | 32.27 | 114 | Han | N.A. | Patterson_et_al_2012 | PASS |
| HGDP00780 | Han | sEA | China | modern | 0 | Sino-Tibetan | 32.27 | 114 | Han | N.A. | Patterson_et_al_2012 | PASS |
| HGDP00781 | Han | sEA | China | modern | 0 | Sino-Tibetan | 32.27 | 114 | Han | N.A. | Patterson_et_al_2012 | PASS |
| HGDP00782 | Han | sEA | China | modern | 0 | Sino-Tibetan | 32.27 | 114 | Han | N.A. | Patterson_et_al_2012 | PASS |
| HGDP00783 | Han | sEA | China | modern | 0 | Sino-Tibetan | 32.27 | 114 | Han | N.A. | Patterson_et_al_2012 | PASS |
| HGDP00784 | Han | sEA | China | modern | 0 | Sino-Tibetan | 32.27 | 114 | Han | N.A. | Patterson_et_al_2012 | PASS |
| HGDP00785 | Han | sEA | China | modern | 0 | Sino-Tibetan | 32.27 | 114 | Han | N.A. | Patterson_et_al_2012 | PASS |
| HGDP00786 | Han | sEA | China | modern | 0 | Sino-Tibetan | 32.27 | 114 | Han | N.A. | Patterson_et_al_2012 | PASS |
| HGDP00811 | Han | sEA | China | modern | 0 | Sino-Tibetan | 32.27 | 114 | Han | N.A. | Patterson_et_al_2012 | PASS |
| HGDP00812 | Han | sEA | China | modern | 0 | Sino-Tibetan | 32.27 | 114 | Han | N.A. | Patterson_et_al_2012 | PASS |
| HGDP00813 | Han | sEA | China | modern | 0 | Sino-Tibetan | 32.27 | 114 | Han | N.A. | Patterson_et_al_2012 | PASS |
| HGDP00814 | Han | sEA | China | modern | 0 | Sino-Tibetan | 32.27 | 114 | Han | N.A. | Patterson_et_al_2012 | PASS |
| HGDP00815 | Han | sEA | China | modern | 0 | Sino-Tibetan | 32.27 | 114 | Han | N.A. | Patterson_et_al_2012 | PASS |
| HGDP00817 | Han | sEA | China | modern | 0 | Sino-Tibetan | 32.27 | 114 | Han | N.A. | Patterson_et_al_2012 | PASS |
| HGDP00818 | Han | sEA | China | modern | 0 | Sino-Tibetan | 32.27 | 114 | Han | N.A. | Patterson_et_al_2012 | PASS |
| HGDP00819 | Han | sEA | China | modern | 0 | Sino-Tibetan | 32.27 | 114 | Han | N.A. | Patterson_et_al_2012 | PASS |
| HGDP00820 | Han | sEA | China | modern | 0 | Sino-Tibetan | 32.27 | 114 | Han | N.A. | Patterson_et_al_2012 | PASS |
| HGDP00821 | Han | sEA | China | modern | 0 | Sino-Tibetan | 32.27 | 114 | Han | N.A. | Patterson_et_al_2012 | PASS |
| HGDP00822 | Han | sEA | China | modern | 0 | Sino-Tibetan | 32.27 | 114 | Han | N.A. | Patterson_et_al_2012 | PASS |
| HGDP00971 | Han | sEA | China | modern | 0 | Sino-Tibetan | 32.27 | 114 | Han | N.A. | Patterson_et_al_2012 | PASS |
| HGDP00972 | Han | sEA | China | modern | 0 | Sino-Tibetan | 32.27 | 114 | Han | N.A. | Patterson_et_al_2012 | PASS |
| HGDP00973 | Han | sEA | China | modern | 0 | Sino-Tibetan | 32.27 | 114 | Han | N.A. | Patterson_et_al_2012 | PASS |
| HGDP00974 | Han | sEA | China | modern | 0 | Sino-Tibetan | 32.27 | 114 | Han | N.A. | Patterson_et_al_2012 | PASS |
| HGDP00975 | Han | sEA | China | modern | 0 | Sino-Tibetan | 32.27 | 114 | Han | N.A. | Patterson_et_al_2012 | PASS |
| HGDP00976 | Han | sEA | China | modern | 0 | Sino-Tibetan | 32.27 | 114 | Han | N.A. | Patterson_et_al_2012 | PASS |
| HGDP00977 | Han | sEA | China | modern | 0 | Sino-Tibetan | 32.27 | 114 | Han | N.A. | Patterson_et_al_2012 | PASS |
| HGDP01021 | Han | sEA | China | modern | 0 | Sino-Tibetan | 32.27 | 114 | Han | N.A. | Patterson_et_al_2012 | PASS |
| HGDP01023 | Han | sEA | China | modern | 0 | Sino-Tibetan | 32.27 | 114 | Han | N.A. | Patterson_et_al_2012 | PASS |
| HGDP01024 | Han | sEA | China | modern | 0 | Sino-Tibetan | 32.27 | 114 | Han | N.A. | Patterson_et_al_2012 | PASS |
| HGDP01287 | Han_NC | nEA | China | modern | 0 | Sino-Tibetan | 32.27 | 114 | Han | N.A. | Patterson_et_al_2012 | PASS |
| HGDP01288 | Han_NC | nEA | China | modern | 0 | Sino-Tibetan | 32.27 | 114 | Han | N.A. | Patterson_et_al_2012 | PASS |
| HGDP01289 | Han_NC | nEA | China | modern | 0 | Sino-Tibetan | 32.27 | 114 | Han | N.A. | Patterson_et_al_2012 | PASS |

| | | | | | | | | | | | | |
|---|---|---|---|---|---|---|---|---|---|---|---|---|
| HGDP01290 | Han_NC | nEA | China | modern | 0 | Sino-Tibetan | 32.27 | 114 | Han | N.A. | Patterson_et_al_2012 | PASS |
| HGDP01291 | Han_NC | nEA | China | modern | 0 | Sino-Tibetan | 32.27 | 114 | Han | N.A. | Patterson_et_al_2012 | PASS |
| HGDP01292 | Han_NC | nEA | China | modern | 0 | Sino-Tibetan | 32.27 | 114 | Han | N.A. | Patterson_et_al_2012 | PASS |
| HGDP01293 | Han_NC | nEA | China | modern | 0 | Sino-Tibetan | 32.27 | 114 | Han | N.A. | Patterson_et_al_2012 | PASS |
| HGDP01294 | Han_NC | nEA | China | modern | 0 | Sino-Tibetan | 32.27 | 114 | Han | N.A. | Patterson_et_al_2012 | PASS |
| HGDP01295 | Han_NC | nEA | China | modern | 0 | Sino-Tibetan | 32.27 | 114 | Han | N.A. | Patterson_et_al_2012 | PASS |
| HGDP01296 | Han_NC | nEA | China | modern | 0 | Sino-Tibetan | 32.27 | 114 | Han | N.A. | Patterson_et_al_2012 | PASS |
| HGDP01095 | Tujia | sEA | China | modern | 0 | Sino-Tibetan | 29 | 109 | Tujia | N.A. | Patterson_et_al_2012 | PASS |
| HGDP01096 | Tujia | sEA | China | modern | 0 | Sino-Tibetan | 29 | 109 | Tujia | N.A. | Patterson_et_al_2012 | PASS |
| HGDP01097 | Tujia | sEA | China | modern | 0 | Sino-Tibetan | 29 | 109 | Tujia | N.A. | Patterson_et_al_2012 | PASS |
| HGDP01098 | Tujia | sEA | China | modern | 0 | Sino-Tibetan | 29 | 109 | Tujia | N.A. | Patterson_et_al_2012 | PASS |
| HGDP01099 | Tujia | sEA | China | modern | 0 | Sino-Tibetan | 29 | 109 | Tujia | N.A. | Patterson_et_al_2012 | PASS |
| HGDP01100 | Tujia | sEA | China | modern | 0 | Sino-Tibetan | 29 | 109 | Tujia | N.A. | Patterson_et_al_2012 | PASS |
| HGDP01101 | Tujia | sEA | China | modern | 0 | Sino-Tibetan | 29 | 109 | Tujia | N.A. | Patterson_et_al_2012 | PASS |
| HGDP01102 | Tujia | sEA | China | modern | 0 | Sino-Tibetan | 29 | 109 | Tujia | N.A. | Patterson_et_al_2012 | PASS |
| HGDP01103 | Tujia | sEA | China | modern | 0 | Sino-Tibetan | 29 | 109 | Tujia | N.A. | Patterson_et_al_2012 | PASS |
| HGDP01104 | Tujia | sEA | China | modern | 0 | Sino-Tibetan | 29 | 109 | Tujia | N.A. | Patterson_et_al_2012 | PASS |
| HGDP01347 | Tu | nEA | China | modern | 0 | Sino-Tibetan | 36 | 101 | Tu | N.A. | Patterson_et_al_2012 | PASS |
| HGDP01348 | Tu | nEA | China | modern | 0 | Sino-Tibetan | 36 | 101 | Tu | N.A. | Patterson_et_al_2012 | PASS |
| HGDP01349 | Tu | nEA | China | modern | 0 | Sino-Tibetan | 36 | 101 | Tu | N.A. | Patterson_et_al_2012 | PASS |
| HGDP01350 | Tu | nEA | China | modern | 0 | Sino-Tibetan | 36 | 101 | Tu | N.A. | Patterson_et_al_2012 | PASS |
| HGDP01351 | Tu | nEA | China | modern | 0 | Sino-Tibetan | 36 | 101 | Tu | N.A. | Patterson_et_al_2012 | PASS |
| HGDP01352 | Tu | nEA | China | modern | 0 | Sino-Tibetan | 36 | 101 | Tu | N.A. | Patterson_et_al_2012 | PASS |
| HGDP01353 | Tu | nEA | China | modern | 0 | Sino-Tibetan | 36 | 101 | Tu | N.A. | Patterson_et_al_2012 | PASS |
| HGDP01354 | Tu | nEA | China | modern | 0 | Sino-Tibetan | 36 | 101 | Tu | N.A. | Patterson_et_al_2012 | PASS |
| HGDP01355 | Tu | nEA | China | modern | 0 | Sino-Tibetan | 36 | 101 | Tu | N.A. | Patterson_et_al_2012 | PASS |
| HGDP01356 | Tu | nEA | China | modern | 0 | Sino-Tibetan | 36 | 101 | Tu | N.A. | Patterson_et_al_2012 | PASS |
| HGDP01243 | Xibo | nEA | China | modern | 0 | Tungusic | 43.5 | 81.5 | N.A. | N.A. | Patterson_et_al_2012 | PASS |
| HGDP01244 | Xibo | nEA | China | modern | 0 | Tungusic | 43.5 | 81.5 | N.A. | N.A. | Patterson_et_al_2012 | PASS |
| HGDP01245 | Xibo | nEA | China | modern | 0 | Tungusic | 43.5 | 81.5 | N.A. | N.A. | Patterson_et_al_2012 | PASS |
| HGDP01246 | Xibo | nEA | China | modern | 0 | Tungusic | 43.5 | 81.5 | N.A. | N.A. | Patterson_et_al_2012 | PASS |
| HGDP01247 | Xibo | nEA | China | modern | 0 | Tungusic | 43.5 | 81.5 | N.A. | N.A. | Patterson_et_al_2012 | PASS |
| HGDP01248 | Xibo | nEA | China | modern | 0 | Tungusic | 43.5 | 81.5 | N.A. | N.A. | Patterson_et_al_2012 | PASS |
| HGDP01249 | Xibo | nEA | China | modern | 0 | Tungusic | 43.5 | 81.5 | N.A. | N.A. | Patterson_et_al_2012 | PASS |
| HGDP01250 | Xibo | nEA | China | modern | 0 | Tungusic | 43.5 | 81.5 | N.A. | N.A. | Patterson_et_al_2012 | PASS |
| HGDP01251 | Xibo | nEA | China | modern | 0 | Tungusic | 43.5 | 81.5 | N.A. | N.A. | Patterson_et_al_2012 | PASS |
| HGDP01233 | Hezhen | nEA | China | modern | 0 | Tungusic | 47.5 | 133.5 | N.A. | N.A. | Patterson_et_al_2012 | PASS |
| HGDP01234 | Hezhen | nEA | China | modern | 0 | Tungusic | 47.5 | 133.5 | N.A. | N.A. | Patterson_et_al_2012 | PASS |
| HGDP01236 | Hezhen | nEA | China | modern | 0 | Tungusic | 47.5 | 133.5 | N.A. | N.A. | Patterson_et_al_2012 | PASS |
| HGDP01237 | Hezhen | nEA | China | modern | 0 | Tungusic | 47.5 | 133.5 | N.A. | N.A. | Patterson_et_al_2012 | PASS |
| HGDP01238 | Hezhen | nEA | China | modern | 0 | Tungusic | 47.5 | 133.5 | N.A. | N.A. | Patterson_et_al_2012 | PASS |

| | | | | | | | | | | | | |
|---|---|---|---|---|---|---|---|---|---|---|---|---|
| HGDP01239 | Hezhen | nEA | China | modern | 0 | Tungusic | 47.5 | 133.5 | N.A. | N.A. | Patterson_et_al_2012 | PASS |
| HGDP01240 | Hezhen | nEA | China | modern | 0 | Tungusic | 47.5 | 133.5 | N.A. | N.A. | Patterson_et_al_2012 | PASS |
| HGDP01241 | Hezhen | nEA | China | modern | 0 | Tungusic | 47.5 | 133.5 | N.A. | N.A. | Patterson_et_al_2012 | PASS |
| HGDP01242 | Hezhen | nEA | China | modern | 0 | Tungusic | 47.5 | 133.5 | N.A. | N.A. | Patterson_et_al_2012 | PASS |
| HGDP01203 | Oroqen | nEA | China | modern | 0 | Tungusic | 50.43 | 126.5 | N.A. | N.A. | Patterson_et_al_2012 | PASS |
| HGDP01204 | Oroqen | nEA | China | modern | 0 | Tungusic | 50.43 | 126.5 | N.A. | N.A. | Patterson_et_al_2012 | PASS |
| HGDP01205 | Oroqen | nEA | China | modern | 0 | Tungusic | 50.43 | 126.5 | N.A. | N.A. | Patterson_et_al_2012 | PASS |
| HGDP01206 | Oroqen | nEA | China | modern | 0 | Tungusic | 50.43 | 126.5 | N.A. | N.A. | Patterson_et_al_2012 | PASS |
| HGDP01207 | Oroqen | nEA | China | modern | 0 | Tungusic | 50.43 | 126.5 | N.A. | N.A. | Patterson_et_al_2012 | PASS |
| HGDP01208 | Oroqen | nEA | China | modern | 0 | Tungusic | 50.43 | 126.5 | N.A. | N.A. | Patterson_et_al_2012 | PASS |
| HGDP01209 | Oroqen | nEA | China | modern | 0 | Tungusic | 50.43 | 126.5 | N.A. | N.A. | Patterson_et_al_2012 | PASS |
| HGDP01211 | Oroqen | nEA | China | modern | 0 | Tungusic | 50.43 | 126.5 | N.A. | N.A. | Patterson_et_al_2012 | PASS |
| HGDP01212 | Oroqen | nEA | China | modern | 0 | Tungusic | 50.43 | 126.5 | N.A. | N.A. | Patterson_et_al_2012 | PASS |
| HGDP01297 | Uygur | nEA | China | modern | 0 | Turkic | 44 | 81 | N.A. | N.A. | Patterson_et_al_2012 | PASS |
| HGDP01298 | Uygur | nEA | China | modern | 0 | Turkic | 44 | 81 | N.A. | N.A. | Patterson_et_al_2012 | PASS |
| HGDP01299 | Uygur | nEA | China | modern | 0 | Turkic | 44 | 81 | N.A. | N.A. | Patterson_et_al_2012 | PASS |
| HGDP01300 | Uygur | nEA | China | modern | 0 | Turkic | 44 | 81 | N.A. | N.A. | Patterson_et_al_2012 | PASS |
| HGDP01301 | Uygur | nEA | China | modern | 0 | Turkic | 44 | 81 | N.A. | N.A. | Patterson_et_al_2012 | PASS |
| HGDP01302 | Uygur | nEA | China | modern | 0 | Turkic | 44 | 81 | N.A. | N.A. | Patterson_et_al_2012 | PASS |
| HGDP01303 | Uygur | nEA | China | modern | 0 | Turkic | 44 | 81 | N.A. | N.A. | Patterson_et_al_2012 | PASS |
| HGDP01304 | Uygur | nEA | China | modern | 0 | Turkic | 44 | 81 | N.A. | N.A. | Patterson_et_al_2012 | PASS |
| HGDP01305 | Uygur | nEA | China | modern | 0 | Turkic | 44 | 81 | N.A. | N.A. | Patterson_et_al_2012 | PASS |
| HGDP01306 | Uygur | nEA | China | modern | 0 | Turkic | 44 | 81 | N.A. | N.A. | Patterson_et_al_2012 | PASS |
| HGDP01213 | Daur | nEA | China | modern | 0 | Mongolic | 48.5 | 124 | N.A. | N.A. | Patterson_et_al_2012 | PASS |
| HGDP01214 | Daur | nEA | China | modern | 0 | Mongolic | 48.5 | 124 | N.A. | N.A. | Patterson_et_al_2012 | PASS |
| HGDP01215 | Daur | nEA | China | modern | 0 | Mongolic | 48.5 | 124 | N.A. | N.A. | Patterson_et_al_2012 | PASS |
| HGDP01216 | Daur | nEA | China | modern | 0 | Mongolic | 48.5 | 124 | N.A. | N.A. | Patterson_et_al_2012 | PASS |
| HGDP01217 | Daur | nEA | China | modern | 0 | Mongolic | 48.5 | 124 | N.A. | N.A. | Patterson_et_al_2012 | PASS |
| HGDP01218 | Daur | nEA | China | modern | 0 | Mongolic | 48.5 | 124 | N.A. | N.A. | Patterson_et_al_2012 | PASS |
| HGDP01220 | Daur | nEA | China | modern | 0 | Mongolic | 48.5 | 124 | N.A. | N.A. | Patterson_et_al_2012 | PASS |
| HGDP01221 | Daur | nEA | China | modern | 0 | Mongolic | 48.5 | 124 | N.A. | N.A. | Patterson_et_al_2012 | PASS |
| HGDP01222 | Daur | nEA | China | modern | 0 | Mongolic | 48.5 | 124 | N.A. | N.A. | Patterson_et_al_2012 | PASS |
| HGDP01223 | Mongola | nEA | Mongolia | modern | 0 | Mongolic | 45 | 111 | N.A. | N.A. | Patterson_et_al_2012 | PASS |
| HGDP01224 | Mongola | nEA | Mongolia | modern | 0 | Mongolic | 45 | 111 | N.A. | N.A. | Patterson_et_al_2012 | PASS |
| HGDP01225 | Mongola | nEA | Mongolia | modern | 0 | Mongolic | 45 | 111 | N.A. | N.A. | Patterson_et_al_2012 | PASS |
| HGDP01226 | Mongola | nEA | Mongolia | modern | 0 | Mongolic | 45 | 111 | N.A. | N.A. | Patterson_et_al_2012 | PASS |
| HGDP01227 | Mongola | nEA | Mongolia | modern | 0 | Mongolic | 45 | 111 | N.A. | N.A. | Patterson_et_al_2012 | PASS |
| HGDP01228 | Mongola | nEA | Mongolia | modern | 0 | Mongolic | 45 | 111 | N.A. | N.A. | Patterson_et_al_2012 | PASS |
| HGDP01229 | Mongola | nEA | Mongolia | modern | 0 | Mongolic | 45 | 111 | N.A. | N.A. | Patterson_et_al_2012 | PASS |
| HGDP01230 | Mongola | nEA | Mongolia | modern | 0 | Mongolic | 45 | 111 | N.A. | N.A. | Patterson_et_al_2012 | PASS |
| HGDP01231 | Mongola | nEA | Mongolia | modern | 0 | Mongolic | 45 | 111 | N.A. | N.A. | Patterson_et_al_2012 | PASS |

| | | | | | | | | | | | | |
|---|---|---|---|---|---|---|---|---|---|---|---|---|
| HGDP01232 | Mongola | nEA | Mongolia | modern | 0 | Mongolic | 45 | 111 | N.A. | N.A. | Patterson_et_al_2012 | PASS |
| HGDP00747 | Japanese | nEA | Japan | modern | 0 | Japonic | 38 | 138 | N.A. | N.A. | Patterson_et_al_2012 | PASS |
| HGDP00748 | Japanese | nEA | Japan | modern | 0 | Japonic | 38 | 138 | N.A. | N.A. | Patterson_et_al_2012 | PASS |
| HGDP00749 | Japanese | nEA | Japan | modern | 0 | Japonic | 38 | 138 | N.A. | N.A. | Patterson_et_al_2012 | PASS |
| HGDP00750 | Japanese | nEA | Japan | modern | 0 | Japonic | 38 | 138 | N.A. | N.A. | Patterson_et_al_2012 | PASS |
| HGDP00751 | Japanese | nEA | Japan | modern | 0 | Japonic | 38 | 138 | N.A. | N.A. | Patterson_et_al_2012 | PASS |
| HGDP00752 | Japanese | nEA | Japan | modern | 0 | Japonic | 38 | 138 | N.A. | N.A. | Patterson_et_al_2012 | PASS |
| HGDP00753 | Japanese | nEA | Japan | modern | 0 | Japonic | 38 | 138 | N.A. | N.A. | Patterson_et_al_2012 | PASS |
| HGDP00754 | Japanese | nEA | Japan | modern | 0 | Japonic | 38 | 138 | N.A. | N.A. | Patterson_et_al_2012 | PASS |
| HGDP00755 | Japanese | nEA | Japan | modern | 0 | Japonic | 38 | 138 | N.A. | N.A. | Patterson_et_al_2012 | PASS |
| HGDP00756 | Japanese | nEA | Japan | modern | 0 | Japonic | 38 | 138 | N.A. | N.A. | Patterson_et_al_2012 | PASS |
| HGDP00757 | Japanese | nEA | Japan | modern | 0 | Japonic | 38 | 138 | N.A. | N.A. | Patterson_et_al_2012 | PASS |
| HGDP00758 | Japanese | nEA | Japan | modern | 0 | Japonic | 38 | 138 | N.A. | N.A. | Patterson_et_al_2012 | PASS |
| HGDP00759 | Japanese | nEA | Japan | modern | 0 | Japonic | 38 | 138 | N.A. | N.A. | Patterson_et_al_2012 | PASS |
| HGDP00760 | Japanese | nEA | Japan | modern | 0 | Japonic | 38 | 138 | N.A. | N.A. | Patterson_et_al_2012 | PASS |
| HGDP00761 | Japanese | nEA | Japan | modern | 0 | Japonic | 38 | 138 | N.A. | N.A. | Patterson_et_al_2012 | PASS |
| HGDP00762 | Japanese | nEA | Japan | modern | 0 | Japonic | 38 | 138 | N.A. | N.A. | Patterson_et_al_2012 | PASS |
| HGDP00763 | Japanese | nEA | Japan | modern | 0 | Japonic | 38 | 138 | N.A. | N.A. | Patterson_et_al_2012 | PASS |
| HGDP00764 | Japanese | nEA | Japan | modern | 0 | Japonic | 38 | 138 | N.A. | N.A. | Patterson_et_al_2012 | PASS |
| HGDP00765 | Japanese | nEA | Japan | modern | 0 | Japonic | 38 | 138 | N.A. | N.A. | Patterson_et_al_2012 | PASS |
| HGDP00766 | Japanese | nEA | Japan | modern | 0 | Japonic | 38 | 138 | N.A. | N.A. | Patterson_et_al_2012 | PASS |
| HGDP00767 | Japanese | nEA | Japan | modern | 0 | Japonic | 38 | 138 | N.A. | N.A. | Patterson_et_al_2012 | PASS |
| HGDP00768 | Japanese | nEA | Japan | modern | 0 | Japonic | 38 | 138 | N.A. | N.A. | Patterson_et_al_2012 | PASS |
| HGDP00769 | Japanese | nEA | Japan | modern | 0 | Japonic | 38 | 138 | N.A. | N.A. | Patterson_et_al_2012 | PASS |
| HGDP00771 | Japanese | nEA | Japan | modern | 0 | Japonic | 38 | 138 | N.A. | N.A. | Patterson_et_al_2012 | PASS |
| HGDP00772 | Japanese | nEA | Japan | modern | 0 | Japonic | 38 | 138 | N.A. | N.A. | Patterson_et_al_2012 | PASS |
| HGDP00773 | Japanese | nEA | Japan | modern | 0 | Japonic | 38 | 138 | N.A. | N.A. | Patterson_et_al_2012 | PASS |
| HGDP00790 | Japanese | nEA | Japan | modern | 0 | Japonic | 38 | 138 | N.A. | N.A. | Patterson_et_al_2012 | PASS |
| HGDP00791 | Japanese | nEA | Japan | modern | 0 | Japonic | 38 | 138 | N.A. | N.A. | Patterson_et_al_2012 | PASS |
| HGDP00828 | Japanese | nEA | Japan | modern | 0 | Japonic | 38 | 138 | N.A. | N.A. | Patterson_et_al_2012 | PASS |
| BAO60 | Bonan | nEA | China | modern | 0 | Mongolic | 35.7 | 102.8 | N.A. | N.A. | Wang_C_et_al_2021 | PASS |
| BAO53 | Bonan | nEA | China | modern | 0 | Mongolic | 35.7 | 102.8 | N.A. | N.A. | Wang_C_et_al_2021 | PASS |
| BAO15 | Bonan | nEA | China | modern | 0 | Mongolic | 35.7 | 102.8 | N.A. | N.A. | Wang_C_et_al_2021 | PASS |
| BAO12 | Bonan | nEA | China | modern | 0 | Mongolic | 35.7 | 102.8 | N.A. | N.A. | Wang_C_et_al_2021 | PASS |
| BAO26 | Bonan | nEA | China | modern | 0 | Mongolic | 35.7 | 102.8 | N.A. | N.A. | Wang_C_et_al_2021 | PASS |
| BAO47 | Bonan | nEA | China | modern | 0 | Mongolic | 35.7 | 102.8 | N.A. | N.A. | Wang_C_et_al_2021 | PASS |
| BAO54 | Bonan | nEA | China | modern | 0 | Mongolic | 35.7 | 102.8 | N.A. | N.A. | Wang_C_et_al_2021 | PASS |
| BAO22 | Bonan | nEA | China | modern | 0 | Mongolic | 35.7 | 102.8 | N.A. | N.A. | Wang_C_et_al_2021 | PASS |
| BAO11 | Bonan | nEA | China | modern | 0 | Mongolic | 35.7 | 102.8 | N.A. | N.A. | Wang_C_et_al_2021 | PASS |
| BAO03 | Bonan | nEA | China | modern | 0 | Mongolic | 35.7 | 102.8 | N.A. | N.A. | Wang_C_et_al_2021 | kin |
| DX1157 | Dongxiang | nEA | China | modern | 0 | Mongolic | 35.6 | 103.2 | N.A. | N.A. | Wang_C_et_al_2021 | PASS |

| | | | | | | | | | | | | |
|---|---|---|---|---|---|---|---|---|---|---|---|---|
| DX1175 | Dongxiang | nEA | China | modern | 0 | Mongolic | 35.6 | 103.2 | N.A. | N.A. | Wang_C_et_al_2021 | PASS |
| DX1170 | Dongxiang | nEA | China | modern | 0 | Mongolic | 35.6 | 103.2 | N.A. | N.A. | Wang_C_et_al_2021 | PASS |
| DX1047 | Dongxiang | nEA | China | modern | 0 | Mongolic | 35.6 | 103.2 | N.A. | N.A. | Wang_C_et_al_2021 | PASS |
| DX1146 | Dongxiang | nEA | China | modern | 0 | Mongolic | 35.6 | 103.2 | N.A. | N.A. | Wang_C_et_al_2021 | PASS |
| DX1058 | Dongxiang | nEA | China | modern | 0 | Mongolic | 35.6 | 103.2 | N.A. | N.A. | Wang_C_et_al_2021 | PASS |
| DX1131 | Dongxiang | nEA | China | modern | 0 | Mongolic | 35.6 | 103.2 | N.A. | N.A. | Wang_C_et_al_2021 | PASS |
| EYG48 | Yugur | nEA | China | modern | 0 | Mongolic | 38.9 | 99.6 | N.A. | N.A. | Wang_C_et_al_2021 | PASS |
| EYG28 | Yugur | nEA | China | modern | 0 | Mongolic | 38.9 | 99.6 | N.A. | N.A. | Wang_C_et_al_2021 | PASS |
| EYG13 | Yugur | nEA | China | modern | 0 | Mongolic | 38.9 | 99.6 | N.A. | N.A. | Wang_C_et_al_2021 | PASS |
| EYG18 | Yugur | nEA | China | modern | 0 | Mongolic | 38.9 | 99.6 | N.A. | N.A. | Wang_C_et_al_2021 | PASS |
| EYG31 | Yugur | nEA | China | modern | 0 | Mongolic | 38.9 | 99.6 | N.A. | N.A. | Wang_C_et_al_2021 | PASS |
| XYG19 | Yugur | nEA | China | modern | 0 | Mongolic | 38.9 | 99.6 | N.A. | N.A. | Wang_C_et_al_2021 | PASS |
| EYG8 | Yugur | nEA | China | modern | 0 | Mongolic | 38.9 | 99.6 | N.A. | N.A. | Wang_C_et_al_2021 | PASS |
| EYG04 | Yugur | nEA | China | modern | 0 | Mongolic | 38.9 | 99.6 | N.A. | N.A. | Wang_C_et_al_2021 | PASS |
| EYG45 | Yugur | nEA | China | modern | 0 | Mongolic | 38.9 | 99.6 | N.A. | N.A. | Wang_C_et_al_2021 | PASS |
| EYG44 | Yugur | nEA | China | modern | 0 | Mongolic | 38.9 | 99.6 | N.A. | N.A. | Wang_C_et_al_2021 | PASS |
| EYG40 | Yugur | nEA | China | modern | 0 | Mongolic | 38.9 | 99.6 | N.A. | N.A. | Wang_C_et_al_2021 | PASS |
| EYG06 | Yugur | nEA | China | modern | 0 | Mongolic | 38.9 | 99.6 | N.A. | N.A. | Wang_C_et_al_2021 | PASS |
| EYG25 | Yugur | nEA | China | modern | 0 | Mongolic | 38.9 | 99.6 | N.A. | N.A. | Wang_C_et_al_2021 | PASS |
| EYG42 | Yugur | nEA | China | modern | 0 | Mongolic | 38.9 | 99.6 | N.A. | N.A. | Wang_C_et_al_2021 | PASS |
| EYG46 | Yugur | nEA | China | modern | 0 | Mongolic | 38.9 | 99.6 | N.A. | N.A. | Wang_C_et_al_2021 | PASS |
| EYG24 | Yugur | nEA | China | modern | 0 | Mongolic | 38.9 | 99.6 | N.A. | N.A. | Wang_C_et_al_2021 | PASS |
| Han516 | Han_Chongqing | sEA | China | modern | 0 | Sino-Tibetan | 29.3 | 106.3 | Han | N.A. | Wang_C_et_al_2021 | PASS |
| Han1968 | Han_Chongqing | sEA | China | modern | 0 | Sino-Tibetan | 29.3 | 106.3 | Han | N.A. | Wang_C_et_al_2021 | PASS |
| Han2150 | Han_Chongqing | sEA | China | modern | 0 | Sino-Tibetan | 29.3 | 106.3 | Han | N.A. | Wang_C_et_al_2021 | PASS |
| Han1994 | Han_Fujian | sEA | China | modern | 0 | Sino-Tibetan | 26.1 | 119.3 | Han | N.A. | Wang_C_et_al_2021 | PASS |
| Han1467 | Han_Fujian | sEA | China | modern | 0 | Sino-Tibetan | 26.1 | 119.3 | Han | N.A. | Wang_C_et_al_2021 | PASS |
| Han1619 | Han_Fujian | sEA | China | modern | 0 | Sino-Tibetan | 26.1 | 119.3 | Han | N.A. | Wang_C_et_al_2021 | PASS |
| Han1900 | Han_Fujian | sEA | China | modern | 0 | Sino-Tibetan | 26.1 | 119.3 | Han | N.A. | Wang_C_et_al_2021 | PASS |
| Han1934 | Han_Fujian | sEA | China | modern | 0 | Sino-Tibetan | 26.1 | 119.3 | Han | N.A. | Wang_C_et_al_2021 | PASS |
| Han1207 | Han_Guangdong | sEA | China | modern | 0 | Sino-Tibetan | 23.2 | 113.2 | Han | N.A. | Wang_C_et_al_2021 | PASS |
| Han1235 | Han_Guangdong | sEA | China | modern | 0 | Sino-Tibetan | 23.2 | 113.2 | Han | N.A. | Wang_C_et_al_2021 | PASS |
| Han1415 | Han_Guangdong | sEA | China | modern | 0 | Sino-Tibetan | 23.2 | 113.2 | Han | N.A. | Wang_C_et_al_2021 | PASS |
| Han1432 | Han_Guangdong | sEA | China | modern | 0 | Sino-Tibetan | 23.2 | 113.2 | Han | N.A. | Wang_C_et_al_2021 | PASS |
| Han1434 | Han_Guangdong | sEA | China | modern | 0 | Sino-Tibetan | 23.2 | 113.2 | Han | N.A. | Wang_C_et_al_2021 | PASS |
| Han2087 | Han_Guangdong | sEA | China | modern | 0 | Sino-Tibetan | 23.2 | 113.2 | Han | N.A. | Wang_C_et_al_2021 | PASS |
| Han2088 | Han_Guangdong | sEA | China | modern | 0 | Sino-Tibetan | 23.2 | 113.2 | Han | N.A. | Wang_C_et_al_2021 | PASS |
| Han894 | Han_Henan | nEA | China | modern | 0 | Sino-Tibetan | 34.8 | 113.6 | Han | N.A. | Wang_C_et_al_2021 | PASS |
| Han1226 | Han_Henan | nEA | China | modern | 0 | Sino-Tibetan | 34.8 | 113.6 | Han | N.A. | Wang_C_et_al_2021 | PASS |
| Han1243 | Han_Henan | nEA | China | modern | 0 | Sino-Tibetan | 34.8 | 113.6 | Han | N.A. | Wang_C_et_al_2021 | PASS |
| Han1713 | Han_Henan | nEA | China | modern | 0 | Sino-Tibetan | 34.8 | 113.6 | Han | N.A. | Wang_C_et_al_2021 | PASS |

| | | | | | | | | | | | | |
|---|---|---|---|---|---|---|---|---|---|---|---|---|
| Han1714 | Han_Henan | nEA | China | modern | 0 | Sino-Tibetan | 34.8 | 113.6 | Han | N.A. | Wang_C_et_al_2021 | PASS |
| Han788 | Han_Hubei | sEA | China | modern | 0 | Sino-Tibetan | 30.5 | 114.3 | Han | N.A. | Wang_C_et_al_2021 | PASS |
| Han789 | Han_Hubei | sEA | China | modern | 0 | Sino-Tibetan | 30.5 | 114.3 | Han | N.A. | Wang_C_et_al_2021 | PASS |
| Han874 | Han_Hubei | sEA | China | modern | 0 | Sino-Tibetan | 30.5 | 114.3 | Han | N.A. | Wang_C_et_al_2021 | PASS |
| Han2161 | Han_Hubei | sEA | China | modern | 0 | Sino-Tibetan | 30.5 | 114.3 | Han | N.A. | Wang_C_et_al_2021 | PASS |
| Han2162 | Han_Hubei | sEA | China | modern | 0 | Sino-Tibetan | 30.5 | 114.3 | Han | N.A. | Wang_C_et_al_2021 | PASS |
| Han1452 | Han_Jiangsu | sEA | China | modern | 0 | Sino-Tibetan | 32.1 | 118.8 | Han | N.A. | Wang_C_et_al_2021 | PASS |
| Han1962 | Han_Jiangsu | sEA | China | modern | 0 | Sino-Tibetan | 32.1 | 118.8 | Han | N.A. | Wang_C_et_al_2021 | PASS |
| Han2054 | Han_Jiangsu | sEA | China | modern | 0 | Sino-Tibetan | 32.1 | 118.8 | Han | N.A. | Wang_C_et_al_2021 | PASS |
| Han2057 | Han_Jiangsu | sEA | China | modern | 0 | Sino-Tibetan | 32.1 | 118.8 | Han | N.A. | Wang_C_et_al_2021 | PASS |
| Han2058 | Han_Jiangsu | sEA | China | modern | 0 | Sino-Tibetan | 32.1 | 118.8 | Han | N.A. | Wang_C_et_al_2021 | PASS |
| Han2076 | Han_Jiangsu | sEA | China | modern | 0 | Sino-Tibetan | 32.1 | 118.8 | Han | N.A. | Wang_C_et_al_2021 | PASS |
| Han2090 | Han_Jiangsu | sEA | China | modern | 0 | Sino-Tibetan | 32.1 | 118.8 | Han | N.A. | Wang_C_et_al_2021 | PASS |
| Han4 | Han_Shandong | nEA | China | modern | 0 | Sino-Tibetan | 36.6 | 117 | Han | N.A. | Wang_C_et_al_2021 | PASS |
| Han25 | Han_Shandong | nEA | China | modern | 0 | Sino-Tibetan | 36.6 | 117 | Han | N.A. | Wang_C_et_al_2021 | PASS |
| Han853 | Han_Shandong | nEA | China | modern | 0 | Sino-Tibetan | 36.6 | 117 | Han | N.A. | Wang_C_et_al_2021 | PASS |
| Han1329 | Han_Shandong | nEA | China | modern | 0 | Sino-Tibetan | 36.6 | 117 | Han | N.A. | Wang_C_et_al_2021 | PASS |
| Han1616 | Han_Shandong | nEA | China | modern | 0 | Sino-Tibetan | 36.6 | 117 | Han | N.A. | Wang_C_et_al_2021 | PASS |
| Han1840 | Han_Shandong | nEA | China | modern | 0 | Sino-Tibetan | 36.6 | 117 | Han | N.A. | Wang_C_et_al_2021 | PASS |
| Han1896 | Han_Shandong | nEA | China | modern | 0 | Sino-Tibetan | 36.6 | 117 | Han | N.A. | Wang_C_et_al_2021 | PASS |
| Han1916 | Han_Shandong | nEA | China | modern | 0 | Sino-Tibetan | 36.6 | 117 | Han | N.A. | Wang_C_et_al_2021 | PASS |
| Han1978 | Han_Shandong | nEA | China | modern | 0 | Sino-Tibetan | 36.6 | 117 | Han | N.A. | Wang_C_et_al_2021 | PASS |
| Han220 | Han_Shandong | nEA | China | modern | 0 | Sino-Tibetan | 36.6 | 117 | Han | N.A. | Wang_C_et_al_2021 | PASS |
| Han24 | Han_Shanghai | sEA | China | modern | 0 | Sino-Tibetan | 31.2 | 121.5 | Han | N.A. | Wang_C_et_al_2021 | PASS |
| Han496 | Han_Shanghai | sEA | China | modern | 0 | Sino-Tibetan | 31.2 | 121.5 | Han | N.A. | Wang_C_et_al_2021 | PASS |
| Han115 | Han_Shanxi | nEA | China | modern | 0 | Sino-Tibetan | 37.9 | 112.5 | Han | N.A. | Wang_C_et_al_2021 | PASS |
| Han790 | Han_Shanxi | nEA | China | modern | 0 | Sino-Tibetan | 37.9 | 112.5 | Han | N.A. | Wang_C_et_al_2021 | PASS |
| Han855 | Han_Shanxi | nEA | China | modern | 0 | Sino-Tibetan | 37.9 | 112.5 | Han | N.A. | Wang_C_et_al_2021 | PASS |
| Han1603 | Han_Shanxi | nEA | China | modern | 0 | Sino-Tibetan | 37.9 | 112.5 | Han | N.A. | Wang_C_et_al_2021 | PASS |
| Han1843 | Han_Shanxi | nEA | China | modern | 0 | Sino-Tibetan | 37.9 | 112.5 | Han | N.A. | Wang_C_et_al_2021 | PASS |
| Han1897 | Han_Shanxi | nEA | China | modern | 0 | Sino-Tibetan | 37.9 | 112.5 | Han | N.A. | Wang_C_et_al_2021 | PASS |
| Han1899 | Han_Shanxi | nEA | China | modern | 0 | Sino-Tibetan | 37.9 | 112.5 | Han | N.A. | Wang_C_et_al_2021 | PASS |
| Han1970 | Han_Shanxi | nEA | China | modern | 0 | Sino-Tibetan | 37.9 | 112.5 | Han | N.A. | Wang_C_et_al_2021 | PASS |
| Han1709 | Han_Sichuan | sEA | China | modern | 0 | Sino-Tibetan | 30.7 | 104.1 | Han | N.A. | Wang_C_et_al_2021 | PASS |
| Han1917 | Han_Sichuan | sEA | China | modern | 0 | Sino-Tibetan | 30.7 | 104.1 | Han | N.A. | Wang_C_et_al_2021 | PASS |
| Han2044 | Han_Sichuan | sEA | China | modern | 0 | Sino-Tibetan | 30.7 | 104.1 | Han | N.A. | Wang_C_et_al_2021 | PASS |
| Han2145 | Han_Sichuan | sEA | China | modern | 0 | Sino-Tibetan | 30.7 | 104.1 | Han | N.A. | Wang_C_et_al_2021 | PASS |
| Han2147 | Han_Sichuan | sEA | China | modern | 0 | Sino-Tibetan | 30.7 | 104.1 | Han | N.A. | Wang_C_et_al_2021 | PASS |
| Han2151 | Han_Sichuan | sEA | China | modern | 0 | Sino-Tibetan | 30.7 | 104.1 | Han | N.A. | Wang_C_et_al_2021 | PASS |
| Han2155 | Han_Sichuan | sEA | China | modern | 0 | Sino-Tibetan | 30.7 | 104.1 | Han | N.A. | Wang_C_et_al_2021 | PASS |
| Han53 | Han_Zhejiang | sEA | China | modern | 0 | Sino-Tibetan | 30.3 | 120.2 | Han | N.A. | Wang_C_et_al_2021 | PASS |

| | | | | | | | | | | | | |
|---|---|---|---|---|---|---|---|---|---|---|---|---|
| Han192 | Han_Zhejiang | sEA | China | modern | 0 | Sino-Tibetan | 30.3 | 120.2 | Han | N.A. | Wang_C_et_al_2021 | PASS |
| Han278 | Han_Zhejiang | sEA | China | modern | 0 | Sino-Tibetan | 30.3 | 120.2 | Han | N.A. | Wang_C_et_al_2021 | PASS |
| Han605 | Han_Zhejiang | sEA | China | modern | 0 | Sino-Tibetan | 30.3 | 120.2 | Han | N.A. | Wang_C_et_al_2021 | PASS |
| Han993 | Han_Zhejiang | sEA | China | modern | 0 | Sino-Tibetan | 30.3 | 120.2 | Han | N.A. | Wang_C_et_al_2021 | PASS |
| DBB006 | Qiang_Danba | sEA | China | modern | 0 | Sino-Tibetan | 30.8 | 101.9 | Qiang | N.A. | Wang_C_et_al_2021 | PASS |
| DBA19 | Qiang_Danba | sEA | China | modern | 0 | Sino-Tibetan | 30.8 | 101.9 | Qiang | N.A. | Wang_C_et_al_2021 | PASS |
| DBA021 | Qiang_Danba | sEA | China | modern | 0 | Sino-Tibetan | 30.8 | 101.9 | Qiang | N.A. | Wang_C_et_al_2021 | PASS |
| DBB007 | Qiang_Danba | sEA | China | modern | 0 | Sino-Tibetan | 30.8 | 101.9 | Qiang | N.A. | Wang_C_et_al_2021 | PASS |
| DBA017 | Qiang_Danba | sEA | China | modern | 0 | Sino-Tibetan | 30.8 | 101.9 | Qiang | N.A. | Wang_C_et_al_2021 | PASS |
| DBA018 | Qiang_Danba | sEA | China | modern | 0 | Sino-Tibetan | 30.8 | 101.9 | Qiang | N.A. | Wang_C_et_al_2021 | PASS |
| DBB001 | Qiang_Danba | sEA | China | modern | 0 | Sino-Tibetan | 30.8 | 101.9 | Qiang | N.A. | Wang_C_et_al_2021 | PASS |
| DBA01 | Qiang_Danba | sEA | China | modern | 0 | Sino-Tibetan | 30.8 | 101.9 | Qiang | N.A. | Wang_C_et_al_2021 | PASS |
| DBB002 | Qiang_Danba | sEA | China | modern | 0 | Sino-Tibetan | 30.8 | 101.9 | Qiang | N.A. | Wang_C_et_al_2021 | PASS |
| BM32 | Qiang_Daofu | sEA | China | modern | 0 | Sino-Tibetan | 30.9 | 101.1 | Qiang | N.A. | Wang_C_et_al_2021 | PASS |
| BM10 | Qiang_Daofu | sEA | China | modern | 0 | Sino-Tibetan | 30.9 | 101.1 | Qiang | N.A. | Wang_C_et_al_2021 | PASS |
| BM43 | Qiang_Daofu | sEA | China | modern | 0 | Sino-Tibetan | 30.9 | 101.1 | Qiang | N.A. | Wang_C_et_al_2021 | PASS |
| BM24 | Qiang_Daofu | sEA | China | modern | 0 | Sino-Tibetan | 30.9 | 101.1 | Qiang | N.A. | Wang_C_et_al_2021 | PASS |
| BM26 | Qiang_Daofu | sEA | China | modern | 0 | Sino-Tibetan | 30.9 | 101.1 | Qiang | N.A. | Wang_C_et_al_2021 | PASS |
| BM21 | Qiang_Daofu | sEA | China | modern | 0 | Sino-Tibetan | 30.9 | 101.1 | Qiang | N.A. | Wang_C_et_al_2021 | PASS |
| BM28 | Qiang_Daofu | sEA | China | modern | 0 | Sino-Tibetan | 30.9 | 101.1 | Qiang | N.A. | Wang_C_et_al_2021 | PASS |
| BM37 | Qiang_Daofu | sEA | China | modern | 0 | Sino-Tibetan | 30.9 | 101.1 | Qiang | N.A. | Wang_C_et_al_2021 | PASS |
| BM27 | Qiang_Daofu | sEA | China | modern | 0 | Sino-Tibetan | 30.9 | 101.1 | Qiang | N.A. | Wang_C_et_al_2021 | PASS |
| BM02 | Qiang_Daofu | sEA | China | modern | 0 | Sino-Tibetan | 30.9 | 101.1 | Qiang | N.A. | Wang_C_et_al_2021 | PASS |
| BM18 | Qiang_Daofu | sEA | China | modern | 0 | Sino-Tibetan | 30.9 | 101.1 | Qiang | N.A. | Wang_C_et_al_2021 | PASS |
| T141 | Tibetan_Chamdo | Tibet | China | modern | 0 | Sino-Tibetan | 31.1 | 97.2 | Tibetan | N.A. | Wang_C_et_al_2021 | PASS |
| T173 | Tibetan_Chamdo | Tibet | China | modern | 0 | Sino-Tibetan | 31.1 | 97.2 | Tibetan | N.A. | Wang_C_et_al_2021 | PASS |
| T274 | Tibetan_Chamdo | Tibet | China | modern | 0 | Sino-Tibetan | 31.1 | 97.2 | Tibetan | N.A. | Wang_C_et_al_2021 | PASS |
| T363 | Tibetan_Chamdo | Tibet | China | modern | 0 | Sino-Tibetan | 31.1 | 97.2 | Tibetan | N.A. | Wang_C_et_al_2021 | PASS |
| T247 | Tibetan_Chamdo | Tibet | China | modern | 0 | Sino-Tibetan | 31.1 | 97.2 | Tibetan | N.A. | Wang_C_et_al_2021 | PASS |
| T74 | Tibetan_Chamdo | Tibet | China | modern | 0 | Sino-Tibetan | 31.1 | 97.2 | Tibetan | N.A. | Wang_C_et_al_2021 | PASS |
| T144 | Tibetan_Chamdo | Tibet | China | modern | 0 | Sino-Tibetan | 31.1 | 97.2 | Tibetan | N.A. | Wang_C_et_al_2021 | PASS |
| T194 | Tibetan_Chamdo | Tibet | China | modern | 0 | Sino-Tibetan | 31.1 | 97.2 | Tibetan | N.A. | Wang_C_et_al_2021 | PASS |
| T196 | Tibetan_Chamdo | Tibet | China | modern | 0 | Sino-Tibetan | 31.1 | 97.2 | Tibetan | N.A. | Wang_C_et_al_2021 | PASS |
| T273 | Tibetan_Chamdo | Tibet | China | modern | 0 | Sino-Tibetan | 31.1 | 97.2 | Tibetan | N.A. | Wang_C_et_al_2021 | PASS |
| T282 | Tibetan_Chamdo | Tibet | China | modern | 0 | Sino-Tibetan | 31.1 | 97.2 | Tibetan | N.A. | Wang_C_et_al_2021 | PASS |
| T283 | Tibetan_Chamdo | Tibet | China | modern | 0 | Sino-Tibetan | 31.1 | 97.2 | Tibetan | N.A. | Wang_C_et_al_2021 | PASS |
| QH1 | Tibetan_Gangcha | Tibet | China | modern | 0 | Sino-Tibetan | 37.3 | 100.2 | Tibetan | N.A. | Wang_C_et_al_2021 | PASS |
| QH5 | Tibetan_Gangcha | Tibet | China | modern | 0 | Sino-Tibetan | 37.3 | 100.2 | Tibetan | N.A. | Wang_C_et_al_2021 | PASS |
| QH7 | Tibetan_Gangcha | Tibet | China | modern | 0 | Sino-Tibetan | 37.3 | 100.2 | Tibetan | N.A. | Wang_C_et_al_2021 | PASS |
| QH18 | Tibetan_Gangcha | Tibet | China | modern | 0 | Sino-Tibetan | 37.3 | 100.2 | Tibetan | N.A. | Wang_C_et_al_2021 | PASS |
| QH10 | Tibetan_Gangcha | Tibet | China | modern | 0 | Sino-Tibetan | 37.3 | 100.2 | Tibetan | N.A. | Wang_C_et_al_2021 | PASS |

| | | | | | | | | | | | | |
|---|---|---|---|---|---|---|---|---|---|---|---|---|
| QH9 | Tibetan_Gangcha | Tibet | China | modern | 0 | Sino-Tibetan | 37.3 | 100.2 | Tibetan | N.A. | Wang_C_et_al_2021 | PASS |
| QH14 | Tibetan_Gangcha | Tibet | China | modern | 0 | Sino-Tibetan | 37.3 | 100.2 | Tibetan | N.A. | Wang_C_et_al_2021 | PASS |
| QH17 | Tibetan_Gangcha | Tibet | China | modern | 0 | Sino-Tibetan | 37.3 | 100.2 | Tibetan | N.A. | Wang_C_et_al_2021 | PASS |
| QH2 | Tibetan_Gangcha | Tibet | China | modern | 0 | Sino-Tibetan | 37.3 | 100.2 | Tibetan | N.A. | Wang_C_et_al_2021 | PASS |
| QH3 | Tibetan_Gangcha | Tibet | China | modern | 0 | Sino-Tibetan | 37.3 | 100.2 | Tibetan | N.A. | Wang_C_et_al_2021 | PASS |
| QH4 | Tibetan_Gangcha | Tibet | China | modern | 0 | Sino-Tibetan | 37.3 | 100.2 | Tibetan | N.A. | Wang_C_et_al_2021 | PASS |
| QH8 | Tibetan_Gangcha | Tibet | China | modern | 0 | Sino-Tibetan | 37.3 | 100.2 | Tibetan | N.A. | Wang_C_et_al_2021 | PASS |
| QH11 | Tibetan_Gangcha | Tibet | China | modern | 0 | Sino-Tibetan | 37.3 | 100.2 | Tibetan | N.A. | Wang_C_et_al_2021 | PASS |
| QH13 | Tibetan_Gangcha | Tibet | China | modern | 0 | Sino-Tibetan | 37.3 | 100.2 | Tibetan | N.A. | Wang_C_et_al_2021 | PASS |
| QH15 | Tibetan_Gangcha | Tibet | China | modern | 0 | Sino-Tibetan | 37.3 | 100.2 | Tibetan | N.A. | Wang_C_et_al_2021 | PASS |
| QH16 | Tibetan_Gangcha | Tibet | China | modern | 0 | Sino-Tibetan | 37.3 | 100.2 | Tibetan | N.A. | Wang_C_et_al_2021 | PASS |
| QH19 | Tibetan_Gangcha | Tibet | China | modern | 0 | Sino-Tibetan | 37.3 | 100.2 | Tibetan | N.A. | Wang_C_et_al_2021 | PASS |
| QH20 | Tibetan_Gangcha | Tibet | China | modern | 0 | Sino-Tibetan | 37.3 | 100.2 | Tibetan | N.A. | Wang_C_et_al_2021 | PASS |
| QH6 | Tibetan_Gangcha | Tibet | China | modern | 0 | Sino-Tibetan | 37.3 | 100.2 | Tibetan | N.A. | Wang_C_et_al_2021 | PASS |
| QH12 | Tibetan_Gangcha | Tibet | China | modern | 0 | Sino-Tibetan | 37.3 | 100.2 | Tibetan | N.A. | Wang_C_et_al_2021 | PASS |
| GN01 | Tibetan_Gannan | Tibet | China | modern | 0 | Sino-Tibetan | 35 | 102.9 | Tibetan | N.A. | Wang_C_et_al_2021 | PASS |
| GN03 | Tibetan_Gannan | Tibet | China | modern | 0 | Sino-Tibetan | 35 | 102.9 | Tibetan | N.A. | Wang_C_et_al_2021 | PASS |
| GN10 | Tibetan_Gannan | Tibet | China | modern | 0 | Sino-Tibetan | 35 | 102.9 | Tibetan | N.A. | Wang_C_et_al_2021 | PASS |
| GN17 | Tibetan_Gannan | Tibet | China | modern | 0 | Sino-Tibetan | 35 | 102.9 | Tibetan | N.A. | Wang_C_et_al_2021 | PASS |
| GN19 | Tibetan_Gannan | Tibet | China | modern | 0 | Sino-Tibetan | 35 | 102.9 | Tibetan | N.A. | Wang_C_et_al_2021 | PASS |
| T21 | Tibetan_Lhasa | Tibet | China | modern | 0 | Sino-Tibetan | 30 | 91.1 | Tibetan | N.A. | Wang_C_et_al_2021 | PASS |
| T165 | Tibetan_Lhasa | Tibet | China | modern | 0 | Sino-Tibetan | 30 | 91.1 | Tibetan | N.A. | Wang_C_et_al_2021 | PASS |
| T170 | Tibetan_Lhasa | Tibet | China | modern | 0 | Sino-Tibetan | 30 | 91.1 | Tibetan | N.A. | Wang_C_et_al_2021 | PASS |
| T181 | Tibetan_Lhasa | Tibet | China | modern | 0 | Sino-Tibetan | 30 | 91.1 | Tibetan | N.A. | Wang_C_et_al_2021 | PASS |
| T183 | Tibetan_Lhasa | Tibet | China | modern | 0 | Sino-Tibetan | 30 | 91.1 | Tibetan | N.A. | Wang_C_et_al_2021 | PASS |
| T76 | Tibetan_Lhasa | Tibet | China | modern | 0 | Sino-Tibetan | 30 | 91.1 | Tibetan | N.A. | Wang_C_et_al_2021 | PASS |
| T125 | Tibetan_Lhasa | Tibet | China | modern | 0 | Sino-Tibetan | 30 | 91.1 | Tibetan | N.A. | Wang_C_et_al_2021 | PASS |
| T185 | Tibetan_Lhasa | Tibet | China | modern | 0 | Sino-Tibetan | 30 | 91.1 | Tibetan | N.A. | Wang_C_et_al_2021 | PASS |
| T18 | Tibetan_Lhasa | Tibet | China | modern | 0 | Sino-Tibetan | 30 | 91.1 | Tibetan | N.A. | Wang_C_et_al_2021 | PASS |
| T67 | Tibetan_Nagqu | Tibet | China | modern | 0 | Sino-Tibetan | 31.5 | 92.1 | Tibetan | N.A. | Wang_C_et_al_2021 | PASS |
| T91 | Tibetan_Nagqu | Tibet | China | modern | 0 | Sino-Tibetan | 31.5 | 92.1 | Tibetan | N.A. | Wang_C_et_al_2021 | PASS |
| T121 | Tibetan_Nagqu | Tibet | China | modern | 0 | Sino-Tibetan | 31.5 | 92.1 | Tibetan | N.A. | Wang_C_et_al_2021 | PASS |
| T187 | Tibetan_Nagqu | Tibet | China | modern | 0 | Sino-Tibetan | 31.5 | 92.1 | Tibetan | N.A. | Wang_C_et_al_2021 | PASS |
| T238 | Tibetan_Nagqu | Tibet | China | modern | 0 | Sino-Tibetan | 31.5 | 92.1 | Tibetan | N.A. | Wang_C_et_al_2021 | PASS |
| T278 | Tibetan_Nagqu | Tibet | China | modern | 0 | Sino-Tibetan | 31.5 | 92.1 | Tibetan | N.A. | Wang_C_et_al_2021 | PASS |
| T312 | Tibetan_Nagqu | Tibet | China | modern | 0 | Sino-Tibetan | 31.5 | 92.1 | Tibetan | N.A. | Wang_C_et_al_2021 | PASS |
| T320 | Tibetan_Nagqu | Tibet | China | modern | 0 | Sino-Tibetan | 31.5 | 92.1 | Tibetan | N.A. | Wang_C_et_al_2021 | PASS |
| T188 | Tibetan_Shannan | Tibet | China | modern | 0 | Sino-Tibetan | 29.2 | 91.8 | Tibetan | N.A. | Wang_C_et_al_2021 | PASS |
| T190 | Tibetan_Shannan | Tibet | China | modern | 0 | Sino-Tibetan | 29.2 | 91.8 | Tibetan | N.A. | Wang_C_et_al_2021 | PASS |
| T242 | Tibetan_Shannan | Tibet | China | modern | 0 | Sino-Tibetan | 29.2 | 91.8 | Tibetan | N.A. | Wang_C_et_al_2021 | PASS |
| T248 | Tibetan_Shannan | Tibet | China | modern | 0 | Sino-Tibetan | 29.2 | 91.8 | Tibetan | N.A. | Wang_C_et_al_2021 | PASS |

| | | | | | | | | | | | | |
|---|---|---|---|---|---|---|---|---|---|---|---|---|
| T126 | Tibetan_Shannan | Tibet | China | modern | 0 | Sino-Tibetan | 29.2 | 91.8 | Tibetan | N.A. | Wang_C_et_al_2021 | PASS |
| T127 | Tibetan_Shannan | Tibet | China | modern | 0 | Sino-Tibetan | 29.2 | 91.8 | Tibetan | N.A. | Wang_C_et_al_2021 | PASS |
| T199 | Tibetan_Shannan | Tibet | China | modern | 0 | Sino-Tibetan | 29.2 | 91.8 | Tibetan | N.A. | Wang_C_et_al_2021 | PASS |
| T200 | Tibetan_Shannan | Tibet | China | modern | 0 | Sino-Tibetan | 29.2 | 91.8 | Tibetan | N.A. | Wang_C_et_al_2021 | PASS |
| T289 | Tibetan_Shannan | Tibet | China | modern | 0 | Sino-Tibetan | 29.2 | 91.8 | Tibetan | N.A. | Wang_C_et_al_2021 | PASS |
| T22 | Tibetan_Shigatse | Tibet | China | modern | 0 | Sino-Tibetan | 29.3 | 88.9 | Tibetan | N.A. | Wang_C_et_al_2021 | PASS |
| T41 | Tibetan_Shigatse | Tibet | China | modern | 0 | Sino-Tibetan | 29.3 | 88.9 | Tibetan | N.A. | Wang_C_et_al_2021 | PASS |
| T77 | Tibetan_Shigatse | Tibet | China | modern | 0 | Sino-Tibetan | 29.3 | 88.9 | Tibetan | N.A. | Wang_C_et_al_2021 | PASS |
| T78 | Tibetan_Shigatse | Tibet | China | modern | 0 | Sino-Tibetan | 29.3 | 88.9 | Tibetan | N.A. | Wang_C_et_al_2021 | PASS |
| T80 | Tibetan_Shigatse | Tibet | China | modern | 0 | Sino-Tibetan | 29.3 | 88.9 | Tibetan | N.A. | Wang_C_et_al_2021 | PASS |
| T189 | Tibetan_Shigatse | Tibet | China | modern | 0 | Sino-Tibetan | 29.3 | 88.9 | Tibetan | N.A. | Wang_C_et_al_2021 | PASS |
| T192 | Tibetan_Shigatse | Tibet | China | modern | 0 | Sino-Tibetan | 29.3 | 88.9 | Tibetan | N.A. | Wang_C_et_al_2021 | PASS |
| T291 | Tibetan_Shigatse | Tibet | China | modern | 0 | Sino-Tibetan | 29.3 | 88.9 | Tibetan | N.A. | Wang_C_et_al_2021 | PASS |
| T297 | Tibetan_Shigatse | Tibet | China | modern | 0 | Sino-Tibetan | 29.3 | 88.9 | Tibetan | N.A. | Wang_C_et_al_2021 | PASS |
| T215 | Tibetan_Shigatse | Tibet | China | modern | 0 | Sino-Tibetan | 29.3 | 88.9 | Tibetan | N.A. | Wang_C_et_al_2021 | PASS |
| XL120 | Tibetan_Xinlong | Tibet | China | modern | 0 | Sino-Tibetan | 31 | 100.3 | Tibetan | N.A. | Wang_C_et_al_2021 | PASS |
| XL61 | Tibetan_Xinlong | Tibet | China | modern | 0 | Sino-Tibetan | 31 | 100.3 | Tibetan | N.A. | Wang_C_et_al_2021 | PASS |
| XL20 | Tibetan_Xinlong | Tibet | China | modern | 0 | Sino-Tibetan | 31 | 100.3 | Tibetan | N.A. | Wang_C_et_al_2021 | PASS |
| XL92 | Tibetan_Xinlong | Tibet | China | modern | 0 | Sino-Tibetan | 31 | 100.3 | Tibetan | N.A. | Wang_C_et_al_2021 | PASS |
| XL60 | Tibetan_Xinlong | Tibet | China | modern | 0 | Sino-Tibetan | 31 | 100.3 | Tibetan | N.A. | Wang_C_et_al_2021 | PASS |
| XL47 | Tibetan_Xinlong | Tibet | China | modern | 0 | Sino-Tibetan | 31 | 100.3 | Tibetan | N.A. | Wang_C_et_al_2021 | PASS |
| XL76 | Tibetan_Xinlong | Tibet | China | modern | 0 | Sino-Tibetan | 31 | 100.3 | Tibetan | N.A. | Wang_C_et_al_2021 | PASS |
| XL22 | Tibetan_Xinlong | Tibet | China | modern | 0 | Sino-Tibetan | 31 | 100.3 | Tibetan | N.A. | Wang_C_et_al_2021 | PASS |
| XL37 | Tibetan_Xinlong | Tibet | China | modern | 0 | Sino-Tibetan | 31 | 100.3 | Tibetan | N.A. | Wang_C_et_al_2021 | PASS |
| XL05 | Tibetan_Xinlong | Tibet | China | modern | 0 | Sino-Tibetan | 31 | 100.3 | Tibetan | N.A. | Wang_C_et_al_2021 | PASS |
| XHTB7 | Tibetan_Xunhua | Tibet | China | modern | 0 | Sino-Tibetan | 35.8 | 102.5 | Tibetan | N.A. | Wang_C_et_al_2021 | PASS |
| XHTB17 | Tibetan_Xunhua | Tibet | China | modern | 0 | Sino-Tibetan | 35.8 | 102.5 | Tibetan | N.A. | Wang_C_et_al_2021 | PASS |
| XHTB20 | Tibetan_Xunhua | Tibet | China | modern | 0 | Sino-Tibetan | 35.8 | 102.5 | Tibetan | N.A. | Wang_C_et_al_2021 | PASS |
| XHTB5 | Tibetan_Xunhua | Tibet | China | modern | 0 | Sino-Tibetan | 35.8 | 102.5 | Tibetan | N.A. | Wang_C_et_al_2021 | PASS |
| YJ86 | Tibetan_Yajiang | Tibet | China | modern | 0 | Sino-Tibetan | 30 | 101 | Tibetan | N.A. | Wang_C_et_al_2021 | PASS |
| YJ14 | Tibetan_Yajiang | Tibet | China | modern | 0 | Sino-Tibetan | 30 | 101 | Tibetan | N.A. | Wang_C_et_al_2021 | PASS |
| YJ39 | Tibetan_Yajiang | Tibet | China | modern | 0 | Sino-Tibetan | 30 | 101 | Tibetan | N.A. | Wang_C_et_al_2021 | PASS |
| YJ01 | Tibetan_Yajiang | Tibet | China | modern | 0 | Sino-Tibetan | 30 | 101 | Tibetan | N.A. | Wang_C_et_al_2021 | PASS |
| YJ140 | Tibetan_Yajiang | Tibet | China | modern | 0 | Sino-Tibetan | 30 | 101 | Tibetan | N.A. | Wang_C_et_al_2021 | PASS |
| YJ37 | Tibetan_Yajiang | Tibet | China | modern | 0 | Sino-Tibetan | 30 | 101 | Tibetan | N.A. | Wang_C_et_al_2021 | PASS |
| YJ12 | Tibetan_Yajiang | Tibet | China | modern | 0 | Sino-Tibetan | 30 | 101 | Tibetan | N.A. | Wang_C_et_al_2021 | PASS |
| YJ111 | Tibetan_Yajiang | Tibet | China | modern | 0 | Sino-Tibetan | 30 | 101 | Tibetan | N.A. | Wang_C_et_al_2021 | PASS |
| YJ25 | Tibetan_Yajiang | Tibet | China | modern | 0 | Sino-Tibetan | 30 | 101 | Tibetan | N.A. | Wang_C_et_al_2021 | PASS |
| YJ104 | Tibetan_Yajiang | Tibet | China | modern | 0 | Sino-Tibetan | 30 | 101 | Tibetan | N.A. | Wang_C_et_al_2021 | PASS |
| Tibetan870 | Tibetan_Yunnan | Tibet | China | modern | 0 | Sino-Tibetan | 27.8 | 99.7 | Tibetan | N.A. | Wang_C_et_al_2021 | PASS |
| Tibetan871 | Tibetan_Yunnan | Tibet | China | modern | 0 | Sino-Tibetan | 27.8 | 99.7 | Tibetan | N.A. | Wang_C_et_al_2021 | PASS |

| | | | | | | | | | | | | |
|---|---|---|---|---|---|---|---|---|---|---|---|---|
| Tibetan872 | Tibetan_Yunnan | Tibet | China | modern | 0 | Sino-Tibetan | 27.8 | 99.7 | Tibetan | N.A. | Wang_C_et_al_2021 | PASS |
| Tibetan937 | Tibetan_Yunnan | Tibet | China | modern | 0 | Sino-Tibetan | 27.8 | 99.7 | Tibetan | N.A. | Wang_C_et_al_2021 | PASS |
| Dong02 | Dong_Guizhou | sEA | China | modern | 0 | Tai-Kadai | 26.7 | 106.6 | Dong | N.A. | Wang_C_et_al_2021 | PASS |
| Dong03 | Dong_Guizhou | sEA | China | modern | 0 | Tai-Kadai | 26.7 | 106.6 | Dong | N.A. | Wang_C_et_al_2021 | PASS |
| Dong05 | Dong_Guizhou | sEA | China | modern | 0 | Tai-Kadai | 26.7 | 106.6 | Dong | N.A. | Wang_C_et_al_2021 | PASS |
| Dong08 | Dong_Guizhou | sEA | China | modern | 0 | Tai-Kadai | 26.7 | 106.6 | Dong | N.A. | Wang_C_et_al_2021 | PASS |
| Dong09 | Dong_Guizhou | sEA | China | modern | 0 | Tai-Kadai | 26.7 | 106.6 | Dong | N.A. | Wang_C_et_al_2021 | PASS |
| Dong10 | Dong_Guizhou | sEA | China | modern | 0 | Tai-Kadai | 26.7 | 106.6 | Dong | N.A. | Wang_C_et_al_2021 | PASS |
| Dong11 | Dong_Guizhou | sEA | China | modern | 0 | Tai-Kadai | 26.7 | 106.6 | Dong | N.A. | Wang_C_et_al_2021 | PASS |
| Dong14 | Dong_Guizhou | sEA | China | modern | 0 | Tai-Kadai | 26.7 | 106.6 | Dong | N.A. | Wang_C_et_al_2021 | PASS |
| Dong13 | Dong_Guizhou | sEA | China | modern | 0 | Tai-Kadai | 26.7 | 106.6 | Dong | N.A. | Wang_C_et_al_2021 | PASS |
| Dong01 | Dong_Guizhou | sEA | China | modern | 0 | Tai-Kadai | 26.7 | 106.6 | Dong | N.A. | Wang_C_et_al_2021 | PASS |
| Dong007 | Dong_Guizhou | sEA | China | modern | 0 | Tai-Kadai | 26.7 | 106.6 | Dong | N.A. | Wang_C_et_al_2021 | PASS |
| HN1-243 | Dong_Hunan | sEA | China | modern | 0 | Tai-Kadai | 27.4 | 109.2 | Dong | N.A. | Wang_C_et_al_2021 | PASS |
| HN1-273 | Dong_Hunan | sEA | China | modern | 0 | Tai-Kadai | 27.4 | 109.2 | Dong | N.A. | Wang_C_et_al_2021 | PASS |
| HN1-232 | Dong_Hunan | sEA | China | modern | 0 | Tai-Kadai | 27.4 | 109.2 | Dong | N.A. | Wang_C_et_al_2021 | PASS |
| HN1-218 | Dong_Hunan | sEA | China | modern | 0 | Tai-Kadai | 27.4 | 109.2 | Dong | N.A. | Wang_C_et_al_2021 | PASS |
| HN1-206 | Dong_Hunan | sEA | China | modern | 0 | Tai-Kadai | 27.4 | 109.2 | Dong | N.A. | Wang_C_et_al_2021 | PASS |
| Dong04 | Dong_Hunan | sEA | China | modern | 0 | Tai-Kadai | 27.4 | 109.2 | Dong | N.A. | Wang_C_et_al_2021 | PASS |
| HN1-260 | Dong_Hunan | sEA | China | modern | 0 | Tai-Kadai | 27.4 | 109.2 | Dong | N.A. | Wang_C_et_al_2021 | PASS |
| Dong12 | Dong_Hunan | sEA | China | modern | 0 | Tai-Kadai | 27.4 | 109.2 | Dong | N.A. | Wang_C_et_al_2021 | PASS |
| HN1-106 | Dong_Hunan | sEA | China | modern | 0 | Tai-Kadai | 27.4 | 109.2 | Dong | N.A. | Wang_C_et_al_2021 | PASS |
| GL01 | Gelao | sEA | China | modern | 0 | Tai-Kadai | 24.8 | 105.3 | Gelao | N.A. | Wang_C_et_al_2021 | PASS |
| GL02 | Gelao | sEA | China | modern | 0 | Tai-Kadai | 24.8 | 105.3 | Gelao | N.A. | Wang_C_et_al_2021 | PASS |
| GL09 | Gelao | sEA | China | modern | 0 | Tai-Kadai | 24.8 | 105.3 | Gelao | N.A. | Wang_C_et_al_2021 | PASS |
| GL07 | Gelao | sEA | China | modern | 0 | Tai-Kadai | 24.8 | 105.3 | Gelao | N.A. | Wang_C_et_al_2021 | PASS |
| GL10 | Gelao | sEA | China | modern | 0 | Tai-Kadai | 24.8 | 105.3 | Gelao | N.A. | Wang_C_et_al_2021 | PASS |
| GL23 | Gelao | sEA | China | modern | 0 | Tai-Kadai | 24.8 | 105.3 | Gelao | N.A. | Wang_C_et_al_2021 | PASS |
| GL13 | Gelao | sEA | China | modern | 0 | Tai-Kadai | 24.8 | 105.3 | Gelao | N.A. | Wang_C_et_al_2021 | PASS |
| GL16 | Gelao | sEA | China | modern | 0 | Tai-Kadai | 24.8 | 105.3 | Gelao | N.A. | Wang_C_et_al_2021 | PASS |
| GL20 | Gelao | sEA | China | modern | 0 | Tai-Kadai | 24.8 | 105.3 | Gelao | N.A. | Wang_C_et_al_2021 | PASS |
| GL24 | Gelao | sEA | China | modern | 0 | Tai-Kadai | 24.8 | 105.3 | Gelao | N.A. | Wang_C_et_al_2021 | PASS |
| CS411 | Li | sEA | China | modern | 0 | Tai-Kadai | 18.5 | 110 | Li | N.A. | Wang_C_et_al_2021 | PASS |
| LS001 | Li | sEA | China | modern | 0 | Tai-Kadai | 18.5 | 110 | Li | N.A. | Wang_C_et_al_2021 | PASS |
| SY071 | Li | sEA | China | modern | 0 | Tai-Kadai | 18.5 | 110 | Li | N.A. | Wang_C_et_al_2021 | PASS |
| WZS058 | Li | sEA | China | modern | 0 | Tai-Kadai | 18.5 | 110 | Li | N.A. | Wang_C_et_al_2021 | PASS |
| W14 | Maonan | sEA | China | modern | 0 | Tai-Kadai | 24.8 | 108.3 | Maonan | N.A. | Wang_C_et_al_2021 | PASS |
| W10 | Maonan | sEA | China | modern | 0 | Tai-Kadai | 24.8 | 108.3 | Maonan | N.A. | Wang_C_et_al_2021 | PASS |
| J6 | Maonan | sEA | China | modern | 0 | Tai-Kadai | 24.8 | 108.3 | Maonan | N.A. | Wang_C_et_al_2021 | PASS |
| J58 | Maonan | sEA | China | modern | 0 | Tai-Kadai | 24.8 | 108.3 | Maonan | N.A. | Wang_C_et_al_2021 | PASS |
| J17 | Maonan | sEA | China | modern | 0 | Tai-Kadai | 24.8 | 108.3 | Maonan | N.A. | Wang_C_et_al_2021 | PASS |

| | | | | | | | | | | | | |
|---|---|---|---|---|---|---|---|---|---|---|---|---|
| J9 | Maonan | sEA | China | modern | 0 | Tai-Kadai | 24.8 | 108.3 | Maonan | N.A. | Wang_C_et_al_2021 | PASS |
| J29 | Maonan | sEA | China | modern | 0 | Tai-Kadai | 24.8 | 108.3 | Maonan | N.A. | Wang_C_et_al_2021 | PASS |
| W29 | Maonan | sEA | China | modern | 0 | Tai-Kadai | 24.8 | 108.3 | Maonan | N.A. | Wang_C_et_al_2021 | PASS |
| W3 | Maonan | sEA | China | modern | 0 | Tai-Kadai | 24.8 | 108.3 | Maonan | N.A. | Wang_C_et_al_2021 | PASS |
| W11 | Maonan | sEA | China | modern | 0 | Tai-Kadai | 24.8 | 108.3 | Maonan | N.A. | Wang_C_et_al_2021 | PASS |
| J32 | Maonan | sEA | China | modern | 0 | Tai-Kadai | 24.8 | 108.3 | Maonan | N.A. | Wang_C_et_al_2021 | PASS |
| J33 | Maonan | sEA | China | modern | 0 | Tai-Kadai | 24.8 | 108.3 | Maonan | N.A. | Wang_C_et_al_2021 | PASS |
| J36 | Maonan | sEA | China | modern | 0 | Tai-Kadai | 24.8 | 108.3 | Maonan | N.A. | Wang_C_et_al_2021 | PASS |
| W13 | Maonan | sEA | China | modern | 0 | Tai-Kadai | 24.8 | 108.3 | Maonan | N.A. | Wang_C_et_al_2021 | PASS |
| W26 | Maonan | sEA | China | modern | 0 | Tai-Kadai | 24.8 | 108.3 | Maonan | N.A. | Wang_C_et_al_2021 | PASS |
| W15 | Maonan | sEA | China | modern | 0 | Tai-Kadai | 24.8 | 108.3 | Maonan | N.A. | Wang_C_et_al_2021 | PASS |
| W41 | Maonan | sEA | China | modern | 0 | Tai-Kadai | 24.8 | 108.3 | Maonan | N.A. | Wang_C_et_al_2021 | PASS |
| B87 | Mulam | sEA | China | modern | 0 | Tai-Kadai | 24.8 | 108.9 | Mulam | N.A. | Wang_C_et_al_2021 | PASS |
| B26 | Mulam | sEA | China | modern | 0 | Tai-Kadai | 24.8 | 108.9 | Mulam | N.A. | Wang_C_et_al_2021 | PASS |
| A114 | Mulam | sEA | China | modern | 0 | Tai-Kadai | 24.8 | 108.9 | Mulam | N.A. | Wang_C_et_al_2021 | PASS |
| B3 | Mulam | sEA | China | modern | 0 | Tai-Kadai | 24.8 | 108.9 | Mulam | N.A. | Wang_C_et_al_2021 | PASS |
| B81 | Mulam | sEA | China | modern | 0 | Tai-Kadai | 24.8 | 108.9 | Mulam | N.A. | Wang_C_et_al_2021 | PASS |
| B48 | Mulam | sEA | China | modern | 0 | Tai-Kadai | 24.8 | 108.9 | Mulam | N.A. | Wang_C_et_al_2021 | PASS |
| B49 | Mulam | sEA | China | modern | 0 | Tai-Kadai | 24.8 | 108.9 | Mulam | N.A. | Wang_C_et_al_2021 | PASS |
| B8 | Mulam | sEA | China | modern | 0 | Tai-Kadai | 24.8 | 108.9 | Mulam | N.A. | Wang_C_et_al_2021 | PASS |
| B14 | Mulam | sEA | China | modern | 0 | Tai-Kadai | 24.8 | 108.9 | Mulam | N.A. | Wang_C_et_al_2021 | PASS |
| B43 | Mulam | sEA | China | modern | 0 | Tai-Kadai | 24.8 | 108.9 | Mulam | N.A. | Wang_C_et_al_2021 | PASS |
| B18 | Mulam | sEA | China | modern | 0 | Tai-Kadai | 24.8 | 108.9 | Mulam | N.A. | Wang_C_et_al_2021 | PASS |
| B32 | Mulam | sEA | China | modern | 0 | Tai-Kadai | 24.8 | 108.9 | Mulam | N.A. | Wang_C_et_al_2021 | PASS |
| A019 | Mulam | sEA | China | modern | 0 | Tai-Kadai | 24.8 | 108.9 | Mulam | N.A. | Wang_C_et_al_2021 | PASS |
| A015 | Mulam | sEA | China | modern | 0 | Tai-Kadai | 24.8 | 108.9 | Mulam | N.A. | Wang_C_et_al_2021 | PASS |
| A005 | Mulam | sEA | China | modern | 0 | Tai-Kadai | 24.8 | 108.9 | Mulam | N.A. | Wang_C_et_al_2021 | PASS |
| A016 | Mulam | sEA | China | modern | 0 | Tai-Kadai | 24.8 | 108.9 | Mulam | N.A. | Wang_C_et_al_2021 | PASS |
| A123 | Mulam | sEA | China | modern | 0 | Tai-Kadai | 24.8 | 108.9 | Mulam | N.A. | Wang_C_et_al_2021 | PASS |
| Zhuang01 | Zhuang | sEA | China | modern | 0 | Tai-Kadai | 22.8 | 108.4 | Zhuang | N.A. | Wang_C_et_al_2021 | PASS |
| Zhuang012 | Zhuang | sEA | China | modern | 0 | Tai-Kadai | 22.8 | 108.4 | Zhuang | N.A. | Wang_C_et_al_2021 | PASS |
| Zhuang019 | Zhuang | sEA | China | modern | 0 | Tai-Kadai | 22.8 | 108.4 | Zhuang | N.A. | Wang_C_et_al_2021 | PASS |
| Zhuang02 | Zhuang | sEA | China | modern | 0 | Tai-Kadai | 22.8 | 108.4 | Zhuang | N.A. | Wang_C_et_al_2021 | PASS |
| Zhuang04 | Zhuang | sEA | China | modern | 0 | Tai-Kadai | 22.8 | 108.4 | Zhuang | N.A. | Wang_C_et_al_2021 | PASS |
| Zhuang013 | Zhuang | sEA | China | modern | 0 | Tai-Kadai | 22.8 | 108.4 | Zhuang | N.A. | Wang_C_et_al_2021 | PASS |
| Zhuang09 | Zhuang | sEA | China | modern | 0 | Tai-Kadai | 22.8 | 108.4 | Zhuang | N.A. | Wang_C_et_al_2021 | PASS |
| Zhuang08 | Zhuang | sEA | China | modern | 0 | Tai-Kadai | 22.8 | 108.4 | Zhuang | N.A. | Wang_C_et_al_2021 | PASS |
| Zhuang03 | Zhuang | sEA | China | modern | 0 | Tai-Kadai | 22.8 | 108.4 | Zhuang | N.A. | Wang_C_et_al_2021 | PASS |
| Zhuang015 | Zhuang | sEA | China | modern | 0 | Tai-Kadai | 22.8 | 108.4 | Zhuang | N.A. | Wang_C_et_al_2021 | PASS |
| Zhuang018 | Zhuang | sEA | China | modern | 0 | Tai-Kadai | 22.8 | 108.4 | Zhuang | N.A. | Wang_C_et_al_2021 | PASS |
| Zhuang11 | Zhuang | sEA | China | modern | 0 | Tai-Kadai | 22.8 | 108.4 | Zhuang | N.A. | Wang_C_et_al_2021 | PASS |

| | | | | | | | | | | | | |
|---|---|---|---|---|---|---|---|---|---|---|---|---|
| Zhuang017 | Zhuang | sEA | China | modern | 0 | Tai-Kadai | 22.8 | 108.4 | Zhuang | N.A. | Wang_C_et_al_2021 | PASS |
| Zhuang020 | Zhuang | sEA | China | modern | 0 | Tai-Kadai | 22.8 | 108.4 | Zhuang | N.A. | Wang_C_et_al_2021 | PASS |
| Zhuang16 | Zhuang | sEA | China | modern | 0 | Tai-Kadai | 22.8 | 108.4 | Zhuang | N.A. | Wang_C_et_al_2021 | PASS |
| Zhuang10 | Zhuang | sEA | China | modern | 0 | Tai-Kadai | 22.8 | 108.4 | Zhuang | N.A. | Wang_C_et_al_2021 | PASS |
| Zhuang06 | Zhuang | sEA | China | modern | 0 | Tai-Kadai | 22.8 | 108.4 | Zhuang | N.A. | Wang_C_et_al_2021 | PASS |
| Zhuang05 | Zhuang | sEA | China | modern | 0 | Tai-Kadai | 22.8 | 108.4 | Zhuang | N.A. | Wang_C_et_al_2021 | PASS |
| Zhuang07 | Zhuang | sEA | China | modern | 0 | Tai-Kadai | 22.8 | 108.4 | Zhuang | N.A. | Wang_C_et_al_2021 | PASS |
| Z01 | Zhuang | sEA | China | modern | 0 | Tai-Kadai | 22.8 | 108.4 | Zhuang | N.A. | Wang_C_et_al_2021 | PASS |
| Z02 | Zhuang | sEA | China | modern | 0 | Tai-Kadai | 22.8 | 108.4 | Zhuang | N.A. | Wang_C_et_al_2021 | PASS |
| Z05 | Zhuang | sEA | China | modern | 0 | Tai-Kadai | 22.8 | 108.4 | Zhuang | N.A. | Wang_C_et_al_2021 | PASS |
| HK03 | Kazakh_China | nEA | China | modern | 0 | Turkic | 38.5 | 94.3 | N.A. | N.A. | Wang_C_et_al_2021 | PASS |
| HK4 | Kazakh_China | nEA | China | modern | 0 | Turkic | 38.5 | 94.3 | N.A. | N.A. | Wang_C_et_al_2021 | PASS |
| HK14 | Kazakh_China | nEA | China | modern | 0 | Turkic | 38.5 | 94.3 | N.A. | N.A. | Wang_C_et_al_2021 | PASS |
| HK15 | Kazakh_China | nEA | China | modern | 0 | Turkic | 38.5 | 94.3 | N.A. | N.A. | Wang_C_et_al_2021 | PASS |
| HKC90 | Kazakh_China | nEA | China | modern | 0 | Turkic | 38.5 | 94.3 | N.A. | N.A. | Wang_C_et_al_2021 | PASS |
| HKC29 | Kazakh_China | nEA | China | modern | 0 | Turkic | 38.5 | 94.3 | N.A. | N.A. | Wang_C_et_al_2021 | PASS |
| HKC109 | Kazakh_China | nEA | China | modern | 0 | Turkic | 38.5 | 94.3 | N.A. | N.A. | Wang_C_et_al_2021 | PASS |
| HKC140 | Kazakh_China | nEA | China | modern | 0 | Turkic | 38.5 | 94.3 | N.A. | N.A. | Wang_C_et_al_2021 | PASS |
| KZ78 | Kyrgyz_China | nEA | China | modern | 0 | Turkic | 43.8 | 87.7 | N.A. | N.A. | Wang_C_et_al_2021 | PASS |
| KZ35 | Kyrgyz_China | nEA | China | modern | 0 | Turkic | 43.8 | 87.7 | N.A. | N.A. | Wang_C_et_al_2021 | PASS |
| KZ15 | Kyrgyz_China | nEA | China | modern | 0 | Turkic | 43.8 | 87.7 | N.A. | N.A. | Wang_C_et_al_2021 | PASS |
| KZ77 | Kyrgyz_China | nEA | China | modern | 0 | Turkic | 43.8 | 87.7 | N.A. | N.A. | Wang_C_et_al_2021 | PASS |
| KZ52 | Kyrgyz_China | nEA | China | modern | 0 | Turkic | 43.8 | 87.7 | N.A. | N.A. | Wang_C_et_al_2021 | PASS |
| KZ12 | Kyrgyz_China | nEA | China | modern | 0 | Turkic | 43.8 | 87.7 | N.A. | N.A. | Wang_C_et_al_2021 | PASS |
| KZ83 | Kyrgyz_China | nEA | China | modern | 0 | Turkic | 43.8 | 87.7 | N.A. | N.A. | Wang_C_et_al_2021 | PASS |
| KZ46 | Kyrgyz_China | nEA | China | modern | 0 | Turkic | 43.8 | 87.7 | N.A. | N.A. | Wang_C_et_al_2021 | PASS |
| KZ81 | Kyrgyz_China | nEA | China | modern | 0 | Turkic | 43.8 | 87.7 | N.A. | N.A. | Wang_C_et_al_2021 | PASS |
| KZ87 | Kyrgyz_China | nEA | China | modern | 0 | Turkic | 43.8 | 87.7 | N.A. | N.A. | Wang_C_et_al_2021 | PASS |
| KZ65 | Kyrgyz_China | nEA | China | modern | 0 | Turkic | 43.8 | 87.7 | N.A. | N.A. | Wang_C_et_al_2021 | PASS |
| KZ22 | Kyrgyz_China | nEA | China | modern | 0 | Turkic | 43.8 | 87.7 | N.A. | N.A. | Wang_C_et_al_2021 | PASS |
| KZ73 | Kyrgyz_China | nEA | China | modern | 0 | Turkic | 43.8 | 87.7 | N.A. | N.A. | Wang_C_et_al_2021 | PASS |
| SL1 | Salar | nEA | China | modern | 0 | Turkic | 35.8 | 102.5 | N.A. | N.A. | Wang_C_et_al_2021 | PASS |
| SL2 | Salar | nEA | China | modern | 0 | Turkic | 35.8 | 102.5 | N.A. | N.A. | Wang_C_et_al_2021 | PASS |
| SL3 | Salar | nEA | China | modern | 0 | Turkic | 35.8 | 102.5 | N.A. | N.A. | Wang_C_et_al_2021 | PASS |
| SL6 | Salar | nEA | China | modern | 0 | Turkic | 35.8 | 102.5 | N.A. | N.A. | Wang_C_et_al_2021 | PASS |
| SL7 | Salar | nEA | China | modern | 0 | Turkic | 35.8 | 102.5 | N.A. | N.A. | Wang_C_et_al_2021 | PASS |
| SL9 | Salar | nEA | China | modern | 0 | Turkic | 35.8 | 102.5 | N.A. | N.A. | Wang_C_et_al_2021 | PASS |
| SL10 | Salar | nEA | China | modern | 0 | Turkic | 35.8 | 102.5 | N.A. | N.A. | Wang_C_et_al_2021 | PASS |
| SL15 | Salar | nEA | China | modern | 0 | Turkic | 35.8 | 102.5 | N.A. | N.A. | Wang_C_et_al_2021 | PASS |
| Ba37 | Bahun | SouthAsia | Nepal | modern | 0 | Indo-European | 27.4 | 85.3 | N.A. | N.A. | Wang_C_et_al_2021 | PASS |
| Ba48 | Bahun | SouthAsia | Nepal | modern | 0 | Indo-European | 27.4 | 85.3 | N.A. | N.A. | Wang_C_et_al_2021 | PASS |

| | | | | | | | | | | | | |
|---|---|---|---|---|---|---|---|---|---|---|---|---|
| Ba15 | Bahun | SouthAsia | Nepal | modern | 0 | Indo-European | 27.4 | 85.3 | N.A. | N.A. | Wang_C_et_al_2021 | PASS |
| Ba25 | Bahun | SouthAsia | Nepal | modern | 0 | Indo-European | 27.4 | 85.3 | N.A. | N.A. | Wang_C_et_al_2021 | PASS |
| Ba02 | Bahun | SouthAsia | Nepal | modern | 0 | Indo-European | 27.4 | 85.3 | N.A. | N.A. | Wang_C_et_al_2021 | PASS |
| Rai16 | Rai | SouthAsia | Nepal | modern | 0 | Indo-European | 27.4 | 85.3 | N.A. | N.A. | Wang_C_et_al_2021 | PASS |
| Rai12 | Rai | SouthAsia | Nepal | modern | 0 | Indo-European | 27.4 | 85.3 | N.A. | N.A. | Wang_C_et_al_2021 | PASS |
| Rai09 | Rai | SouthAsia | Nepal | modern | 0 | Indo-European | 27.4 | 85.3 | N.A. | N.A. | Wang_C_et_al_2021 | PASS |
| Rai14 | Rai | SouthAsia | Nepal | modern | 0 | Indo-European | 27.4 | 85.3 | N.A. | N.A. | Wang_C_et_al_2021 | PASS |
| Rai04 | Rai | SouthAsia | Nepal | modern | 0 | Indo-European | 27.4 | 85.3 | N.A. | N.A. | Wang_C_et_al_2021 | PASS |
| Te18 | Tharu | SouthAsia | Nepal | modern | 0 | Indo-European | 27.4 | 86.2 | N.A. | N.A. | Wang_C_et_al_2021 | PASS |
| Te16 | Tharu | SouthAsia | Nepal | modern | 0 | Indo-European | 27.4 | 86.2 | N.A. | N.A. | Wang_C_et_al_2021 | PASS |
| Te09 | Tharu | SouthAsia | Nepal | modern | 0 | Indo-European | 27.4 | 86.2 | N.A. | N.A. | Wang_C_et_al_2021 | PASS |
| Te15 | Tharu | SouthAsia | Nepal | modern | 0 | Indo-European | 27.4 | 86.2 | N.A. | N.A. | Wang_C_et_al_2021 | PASS |
| Te06 | Tharu | SouthAsia | Nepal | modern | 0 | Indo-European | 27.4 | 86.2 | N.A. | N.A. | Wang_C_et_al_2021 | PASS |
| Gu18 | Gurung | SouthAsia | Nepal | modern | 0 | Sino-Tibetan | 27.4 | 86.2 | Gurung | N.A. | Wang_C_et_al_2021 | PASS |
| Gu22 | Gurung | SouthAsia | Nepal | modern | 0 | Sino-Tibetan | 27.4 | 86.2 | Gurung | N.A. | Wang_C_et_al_2021 | PASS |
| Gu24 | Gurung | SouthAsia | Nepal | modern | 0 | Sino-Tibetan | 27.4 | 86.2 | Gurung | N.A. | Wang_C_et_al_2021 | PASS |
| Gu26 | Gurung | SouthAsia | Nepal | modern | 0 | Sino-Tibetan | 27.4 | 86.2 | Gurung | N.A. | Wang_C_et_al_2021 | PASS |
| Gu27 | Gurung | SouthAsia | Nepal | modern | 0 | Sino-Tibetan | 27.4 | 86.2 | Gurung | N.A. | Wang_C_et_al_2021 | PASS |
| Ma41 | Magar | SouthAsia | Nepal | modern | 0 | Sino-Tibetan | 27.4 | 86.2 | Magar | N.A. | Wang_C_et_al_2021 | PASS |
| Ma11 | Magar | SouthAsia | Nepal | modern | 0 | Sino-Tibetan | 27.4 | 86.2 | Magar | N.A. | Wang_C_et_al_2021 | PASS |
| Ma30 | Magar | SouthAsia | Nepal | modern | 0 | Sino-Tibetan | 27.4 | 86.2 | Magar | N.A. | Wang_C_et_al_2021 | PASS |
| Ma23 | Magar | SouthAsia | Nepal | modern | 0 | Sino-Tibetan | 27.4 | 86.2 | Magar | N.A. | Wang_C_et_al_2021 | PASS |
| Ma07 | Magar | SouthAsia | Nepal | modern | 0 | Sino-Tibetan | 27.4 | 86.2 | Magar | N.A. | Wang_C_et_al_2021 | PASS |
| Ma28 | Magar | SouthAsia | Nepal | modern | 0 | Sino-Tibetan | 27.4 | 86.2 | Magar | N.A. | Wang_C_et_al_2021 | PASS |
| Ne41 | Newar | SouthAsia | Nepal | modern | 0 | Sino-Tibetan | 27.4 | 85.3 | Newar | N.A. | Wang_C_et_al_2021 | PASS |
| Ne72 | Newar | SouthAsia | Nepal | modern | 0 | Sino-Tibetan | 27.4 | 85.3 | Newar | N.A. | Wang_C_et_al_2021 | PASS |
| Ne30 | Newar | SouthAsia | Nepal | modern | 0 | Sino-Tibetan | 27.4 | 85.3 | Newar | N.A. | Wang_C_et_al_2021 | PASS |
| Ne08 | Newar | SouthAsia | Nepal | modern | 0 | Sino-Tibetan | 27.4 | 85.3 | Newar | N.A. | Wang_C_et_al_2021 | PASS |
| Ne66 | Newar | SouthAsia | Nepal | modern | 0 | Sino-Tibetan | 27.4 | 85.3 | Newar | N.A. | Wang_C_et_al_2021 | PASS |
| Ne61 | Newar | SouthAsia | Nepal | modern | 0 | Sino-Tibetan | 27.4 | 85.3 | Newar | N.A. | Wang_C_et_al_2021 | PASS |
| Ne35 | Newar | SouthAsia | Nepal | modern | 0 | Sino-Tibetan | 27.4 | 85.3 | Newar | N.A. | Wang_C_et_al_2021 | PASS |
| Ne01 | Newar | SouthAsia | Nepal | modern | 0 | Sino-Tibetan | 27.4 | 85.3 | Newar | N.A. | Wang_C_et_al_2021 | PASS |
| Sherpa08 | Sherpa | SouthAsia | Nepal | modern | 0 | Sino-Tibetan | 27.4 | 85.3 | Sherpa | N.A. | Wang_C_et_al_2021 | PASS |
| Sherpa04 | Sherpa | SouthAsia | Nepal | modern | 0 | Sino-Tibetan | 27.4 | 85.3 | Sherpa | N.A. | Wang_C_et_al_2021 | PASS |
| Sherpa13 | Sherpa | SouthAsia | Nepal | modern | 0 | Sino-Tibetan | 27.4 | 85.3 | Sherpa | N.A. | Wang_C_et_al_2021 | PASS |
| Sherpa09 | Sherpa | SouthAsia | Nepal | modern | 0 | Sino-Tibetan | 27.4 | 85.3 | Sherpa | N.A. | Wang_C_et_al_2021 | PASS |
| Ta39 | Tamang | SouthAsia | Nepal | modern | 0 | Sino-Tibetan | 27.4 | 86.2 | Tamang | N.A. | Wang_C_et_al_2021 | PASS |
| Ta43 | Tamang | SouthAsia | Nepal | modern | 0 | Sino-Tibetan | 27.4 | 86.2 | Tamang | N.A. | Wang_C_et_al_2021 | PASS |
| Ta24 | Tamang | SouthAsia | Nepal | modern | 0 | Sino-Tibetan | 27.4 | 86.2 | Tamang | N.A. | Wang_C_et_al_2021 | PASS |
| Ta64 | Tamang | SouthAsia | Nepal | modern | 0 | Sino-Tibetan | 27.4 | 86.2 | Tamang | N.A. | Wang_C_et_al_2021 | PASS |
| Ta80 | Tamang | SouthAsia | Nepal | modern | 0 | Sino-Tibetan | 27.4 | 86.2 | Tamang | N.A. | Wang_C_et_al_2021 | PASS |

| | | | | | | | | | | | | |
|---|---|---|---|---|---|---|---|---|---|---|---|---|
| Ta52 | Tamang | SouthAsia | Nepal | modern | 0 | Sino-Tibetan | 27.4 | 86.2 | Tamang | N.A. | Wang_C_et_al_2021 | PASS |
| Ta05 | Tamang | SouthAsia | Nepal | modern | 0 | Sino-Tibetan | 27.4 | 86.2 | Tamang | N.A. | Wang_C_et_al_2021 | PASS |
| Ta03 | Tamang | SouthAsia | Nepal | modern | 0 | Sino-Tibetan | 27.4 | 86.2 | Tamang | N.A. | Wang_C_et_al_2021 | PASS |
| TY | Tianyuan_40000BP | nEA | China | ancient | 40000 | N.A. | 39.4 | 115.5 | N.A. | N.A. | Yang_et_al_2017 | PASS |
| Yumin | Yumin_8400BP | nEA | Mongolia | ancient | 8375 | N.A. | 42 | 114.2 | N.A. | N.A. | Yang_et_al_2020 | PASS |
| Bianbian | Bianbian_9500BP | nEA | China | ancient | 9513 | N.A. | 36.1 | 118.5 | N.A. | N.A. | Yang_et_al_2020 | PASS |
| BS | Boshan_8200BP | nEA | China | ancient | 8180 | N.A. | 36.5 | 117.9 | N.A. | N.A. | Yang_et_al_2020 | PASS |
| XJS1309_M7 | Xiaojingshan_7800BP | nEA | China | ancient | 7797 | N.A. | 36.5 | 117.9 | N.A. | N.A. | Yang_et_al_2020 | PASS |
| XJS1311_M16 | Xiaojingshan_7800BP | nEA | China | ancient | 7861 | N.A. | 36.5 | 117 | N.A. | N.A. | Yang_et_al_2020 | PASS |
| XJS1309_M4 | Xiaojingshan_7800BP | nEA | China | ancient | 7806 | N.A. | 36.5 | 117 | N.A. | N.A. | Yang_et_al_2020 | PASS |
| Xiaogao | Xiaogao_8700BP | nEA | China | ancient | 8684 | N.A. | 37.9 | 117.6 | N.A. | N.A. | Yang_et_al_2020 | PASS |
| Qihe2_d | Qihe_8400BP | sEA | China | ancient | 8394 | N.A. | 25.4 | 117.6 | N.A. | N.A. | Yang_et_al_2020 | PASS |
| LD1 | Liangdao_7750BP | sEA | Taiwan | ancient | 8190 | N.A. | 26.3 | 120.2 | N.A. | N.A. | Yang_et_al_2020 | PASS |
| LD2 | Liangdao_7750BP | sEA | Taiwan | ancient | 7575 | N.A. | 26.3 | 120.2 | N.A. | N.A. | Yang_et_al_2020 | PASS |
| SuogangB1_d | Suogang_4550BP | sEA | Taiwan | ancient | 4550 | N.A. | 23.5 | 119.6 | N.A. | N.A. | Yang_et_al_2020 | PASS |
| SuogangB3_d | Suogang_4550BP | sEA | Taiwan | ancient | 4550 | N.A. | 23.5 | 119.6 | N.A. | N.A. | Yang_et_al_2020 | PASS |
| L5705 | Xitoucun_4500BP | sEA | China | ancient | 4333 | N.A. | 26.2 | 119.1 | N.A. | N.A. | Yang_et_al_2020 | PASS |
| L5700 | Xitoucun_4500BP | sEA | China | ancient | 4474 | N.A. | 26.2 | 119.1 | N.A. | N.A. | Yang_et_al_2020 | PASS |
| L5692_d | Xitoucun_4500BP | sEA | China | ancient | 4474 | N.A. | 26.2 | 119.1 | N.A. | N.A. | Yang_et_al_2020 | imiss |
| L5706_d | Xitoucun_4500BP | sEA | China | ancient | 4467 | N.A. | 26.2 | 119.1 | N.A. | N.A. | Yang_et_al_2020 | PASS |
| L5704_d | Xitoucun_4500BP | sEA | China | ancient | 4502 | N.A. | 26.2 | 119.1 | N.A. | N.A. | Yang_et_al_2020 | PASS |
| L5703_d | Xitoucun_4500BP | sEA | China | ancient | 4572 | N.A. | 26.2 | 119.1 | N.A. | N.A. | Yang_et_al_2020 | imiss |
| L5701_d | Xitoucun_4500BP | sEA | China | ancient | 4329 | N.A. | 26.2 | 119.1 | N.A. | N.A. | Yang_et_al_2020 | PASS |
| L7415 | Tanshishan_4350BP | sEA | China | ancient | 4333 | N.A. | 26.1 | 119.2 | N.A. | N.A. | Yang_et_al_2020 | PASS |
| L7417_d | Tanshishan_4350BP | sEA | China | ancient | 4472 | N.A. | 26.1 | 119.2 | N.A. | N.A. | Yang_et_al_2020 | PASS |
| L5698_d | Tanshishan_4350BP | sEA | China | ancient | 4318 | N.A. | 26.2 | 119.1 | N.A. | N.A. | Yang_et_al_2020 | imiss |
| L5696_d | Tanshishan_4350BP | sEA | China | ancient | NA | N.A. | 26.2 | 119.1 | N.A. | N.A. | Yang_et_al_2020 | imiss |
| L5694 | Chuanyun_300BP | sEA | China | ancient | 308 | N.A. | 25.6 | 117.3 | N.A. | N.A. | Yang_et_al_2020 | PASS |
| I3354 | Boisman_6900BP | nEA | Russia | ancient | 7237 | N.A. | 42.79 | 131.28 | N.A. | N.A. | Wang_C_et_al_2021 | PASS |
| I1196 | Boisman_6900BP | nEA | Russia | ancient | 7058 | N.A. | 42.79 | 131.28 | N.A. | N.A. | Wang_C_et_al_2021 | PASS |
| I1198 | Boisman_6900BP | nEA | Russia | ancient | 6872 | N.A. | 42.79 | 131.28 | N.A. | N.A. | Wang_C_et_al_2021 | PASS |
| I1193 | Boisman_6900BP | nEA | Russia | ancient | 6832 | N.A. | 42.79 | 131.28 | N.A. | N.A. | Wang_C_et_al_2021 | PASS |
| I3355 | Boisman_6900BP | nEA | Russia | ancient | 6821 | N.A. | 42.79 | 131.28 | N.A. | N.A. | Wang_C_et_al_2021 | PASS |
| I1197 | Boisman_6900BP | nEA | Russia | ancient | 6804 | N.A. | 42.79 | 131.28 | N.A. | N.A. | Wang_C_et_al_2021 | PASS |
| I1190 | Boisman_6900BP | nEA | Russia | ancient | 6675 | N.A. | 42.79 | 131.28 | N.A. | N.A. | Wang_C_et_al_2021 | PASS |
| I1194 | Boisman_6900BP | nEA | Russia | ancient | 6652 | N.A. | 42.79 | 131.28 | N.A. | N.A. | Wang_C_et_al_2021 | PASS |
| I14819 | Boisman_5600BP | nEA | Russia | ancient | 5625 | N.A. | 42.79 | 131.28 | N.A. | N.A. | Wang_C_et_al_2021 | PASS |
| I3356 | Boisman_5600BP | nEA | Russia | ancient | 5605 | N.A. | 42.79 | 131.28 | N.A. | N.A. | Wang_C_et_al_2021 | PASS |
| I1206 | Boisman_6900BP | nEA | Russia | ancient | 6954 | N.A. | 42.79 | 131.28 | N.A. | N.A. | Wang_C_et_al_2021 | PASS |
| I13886 | Jomon_3500BP | nEA | Japan | ancient | 4003 | N.A. | 35.55 | 140.16 | N.A. | N.A. | Wang_C_et_al_2021 | PASS |
| I13884 | Jomon_3500BP | nEA | Japan | ancient | 4351 | N.A. | 35.55 | 140.16 | N.A. | N.A. | Wang_C_et_al_2021 | PASS |

| | | | | | | | | | | | | |
|---|---|---|---|---|---|---|---|---|---|---|---|---|
| I6341 | Jomon_3500BP | nEA | Japan | ancient | 4200 | N.A. | 45.37 | 141.03 | N.A. | N.A. | Wang_C_et_al_2021 | PASS |
| I13882 | Jomon_3500BP | nEA | Japan | ancient | 3234 | N.A. | 35.55 | 140.16 | N.A. | N.A. | Wang_C_et_al_2021 | PASS |
| I13885 | Jomon_3500BP | nEA | Japan | ancient | 3216 | N.A. | 35.55 | 140.16 | N.A. | N.A. | Wang_C_et_al_2021 | PASS |
| I13883 | Jomon_3500BP | nEA | Japan | ancient | 2859 | N.A. | 35.55 | 140.16 | N.A. | N.A. | Wang_C_et_al_2021 | PASS |
| I6358 | Mongolia_7300BP | nEA | Mongolia | ancient | 7167 | N.A. | 48.29 | 115.1 | N.A. | N.A. | Wang_C_et_al_2021 | PASS |
| I7021 | Mongolia_7300BP | nEA | Mongolia | ancient | 7050 | N.A. | 48 | 113.93 | N.A. | N.A. | Wang_C_et_al_2021 | PASS |
| I11696 | Mongolia_7300BP | nEA | Mongolia | ancient | 7528 | N.A. | 49.39 | 102.7 | N.A. | N.A. | Wang_C_et_al_2021 | PASS |
| I11698 | Mongolia_7300BP | nEA | Mongolia | ancient | 7520 | N.A. | 49.39 | 102.7 | N.A. | N.A. | Wang_C_et_al_2021 | PASS |
| I11697 | Mongolia_7300BP | nEA | Mongolia | ancient | 7500 | N.A. | 49.39 | 102.7 | N.A. | N.A. | Wang_C_et_al_2021 | PASS |
| I13179 | Mongolia_7300BP | nEA | Mongolia | ancient | 7515 | N.A. | 49.39 | 102.7 | N.A. | N.A. | Wang_C_et_al_2021 | PASS |
| I13698 | Mongolia_7300BP | nEA | Mongolia | ancient | 7504 | N.A. | 49.53 | 103.28 | N.A. | N.A. | Wang_C_et_al_2021 | PASS |
| I14000 | Mongolia_7300BP | nEA | Mongolia | ancient | 7480 | N.A. | 49.53 | 103.28 | N.A. | N.A. | Wang_C_et_al_2021 | PASS |
| S97.EC | Wuzhuangguoliang_5000BP | nEA | China | ancient | 5050 | N.A. | 37.82 | 109.05 | N.A. | N.A. | Wang_C_et_al_2021 | PASS |
| S123.EC | Wuzhuangguoliang_5000BP | nEA | China | ancient | 5050 | N.A. | 37.82 | 109.05 | N.A. | N.A. | Wang_C_et_al_2021 | PASS |
| 18R21262.EC | Wuzhuangguoliang_5000BP | nEA | China | ancient | 5050 | N.A. | 37.82 | 109.05 | N.A. | N.A. | Wang_C_et_al_2021 | PASS |
| S91.EC | Wuzhuangguoliang_5000BP | nEA | China | ancient | 5050 | N.A. | 37.82 | 109.05 | N.A. | N.A. | Wang_C_et_al_2021 | imiss |
| S118.EC | Wuzhuangguoliang_5000BP | nEA | China | ancient | 5050 | N.A. | 37.82 | 109.05 | N.A. | N.A. | Wang_C_et_al_2021 | imiss |
| S120.EC | Wuzhuangguoliang_5000BP | nEA | China | ancient | 5050 | N.A. | 37.82 | 109.05 | N.A. | N.A. | Wang_C_et_al_2021 | imiss |
| 18R21265.EC | Wuzhuangguoliang_5000BP | nEA | China | ancient | 5050 | N.A. | 37.82 | 109.05 | N.A. | N.A. | Wang_C_et_al_2021 | imiss |
| 18R21266.EC | Wuzhuangguoliang_5000BP | nEA | China | ancient | 4785 | N.A. | 37.82 | 109.05 | N.A. | N.A. | Wang_C_et_al_2021 | PASS |
| I6221 | Afanasievo_5000BP | nEA | Mongolia | ancient | 4996 | N.A. | 46.4 | 100.82 | N.A. | N.A. | Wang_C_et_al_2021 | PASS |
| I13957 | Afanasievo_4600BP | nEA | Mongolia | ancient | 4625 | N.A. | 49.34 | 88.71 | N.A. | N.A. | Wang_C_et_al_2021 | PASS |
| I6347 | CenterWest_3100BP | nEA | Mongolia | ancient | 3295 | N.A. | 49.31 | 95.45 | N.A. | N.A. | Wang_C_et_al_2021 | PASS |
| I6264 | CenterWest_3100BP | nEA | Mongolia | ancient | 3275 | N.A. | 45.92 | 100.83 | N.A. | N.A. | Wang_C_et_al_2021 | PASS |
| I6262 | CenterWest_3100BP | nEA | Mongolia | ancient | 3270 | N.A. | 45.92 | 100.83 | N.A. | N.A. | Wang_C_et_al_2021 | PASS |
| I13766 | CenterWest_3100BP | nEA | Mongolia | ancient | 3208 | N.A. | 46.43 | 100.82 | N.A. | N.A. | Wang_C_et_al_2021 | PASS |
| I13767 | CenterWest_3100BP | nEA | Mongolia | ancient | 3198 | N.A. | 48.11 | 102.55 | N.A. | N.A. | Wang_C_et_al_2021 | PASS |
| I12975 | CenterWest_3100BP | nEA | Mongolia | ancient | 3110 | N.A. | 51.5 | 100.67 | N.A. | N.A. | Wang_C_et_al_2021 | PASS |
| I7039 | CenterWest_3100BP | nEA | Mongolia | ancient | 3066 | N.A. | 47.42 | 92.23 | N.A. | N.A. | Wang_C_et_al_2021 | PASS |
| I13505 | CenterWest_3100BP | nEA | Mongolia | ancient | 3001 | N.A. | 46.43 | 100.82 | N.A. | N.A. | Wang_C_et_al_2021 | PASS |
| I6362 | CenterWest_3100BP | nEA | Mongolia | ancient | 2921 | N.A. | 46.06 | 92.03 | N.A. | N.A. | Wang_C_et_al_2021 | PASS |
| I6351 | CenterWest_2900BP | nEA | Mongolia | ancient | 2885 | N.A. | 49.7 | 93.8 | N.A. | N.A. | Wang_C_et_al_2021 | PASS |
| I12978 | Chemurchek_4450BP | nEA | Mongolia | ancient | 4472 | N.A. | 46.12 | 91.57 | N.A. | N.A. | Wang_C_et_al_2021 | PASS |
| I12957 | Chemurchek_4450BP | nEA | Mongolia | ancient | 4462 | N.A. | 46.12 | 91.57 | N.A. | N.A. | Wang_C_et_al_2021 | PASS |
| I3620 | Hanben_1550BP | sEA | Taiwan | ancient | 1857 | N.A. | 24.33 | 121.77 | N.A. | N.A. | Wang_C_et_al_2021 | PASS |
| I3615 | Hanben_1550BP | sEA | Taiwan | ancient | 1833 | N.A. | 24.33 | 121.77 | N.A. | N.A. | Wang_C_et_al_2021 | PASS |
| I3617 | Hanben_1550BP | sEA | Taiwan | ancient | 1644 | N.A. | 24.33 | 121.77 | N.A. | N.A. | Wang_C_et_al_2021 | PASS |
| I3616 | Hanben_1550BP | sEA | Taiwan | ancient | 1641 | N.A. | 24.33 | 121.77 | N.A. | N.A. | Wang_C_et_al_2021 | PASS |
| I3621 | Hanben_1550BP | sEA | Taiwan | ancient | 1595 | N.A. | 24.33 | 121.77 | N.A. | N.A. | Wang_C_et_al_2021 | PASS |
| I3618 | Hanben_1550BP | sEA | Taiwan | ancient | 1550 | N.A. | 24.33 | 121.77 | N.A. | N.A. | Wang_C_et_al_2021 | PASS |
| I3731 | Hanben_1550BP | sEA | Taiwan | ancient | 1550 | N.A. | 24.33 | 121.77 | N.A. | N.A. | Wang_C_et_al_2021 | PASS |

| | | | | | | | | | | | | |
|---|---|---|---|---|---|---|---|---|---|---|---|---|
| I3727 | Hanben_1550BP | sEA | Taiwan | ancient | 1550 | N.A. | 24.33 | 121.77 | N.A. | N.A. | Wang_C_et_al_2021 | PASS |
| I3736 | Hanben_1550BP | sEA | Taiwan | ancient | 1550 | N.A. | 24.33 | 121.77 | N.A. | N.A. | Wang_C_et_al_2021 | PASS |
| I13692 | Hanben_1550BP | sEA | Taiwan | ancient | 1550 | N.A. | 24.33 | 121.77 | N.A. | N.A. | Wang_C_et_al_2021 | PASS |
| I8074 | Hanben_1550BP | sEA | Taiwan | ancient | 1550 | N.A. | 24.33 | 121.77 | N.A. | N.A. | Wang_C_et_al_2021 | PASS |
| I14929 | Hanben_1550BP | sEA | Taiwan | ancient | 1550 | N.A. | 24.33 | 121.77 | N.A. | N.A. | Wang_C_et_al_2021 | PASS |
| I14931 | Hanben_1550BP | sEA | Taiwan | ancient | 1550 | N.A. | 24.33 | 121.77 | N.A. | N.A. | Wang_C_et_al_2021 | PASS |
| I8080 | Hanben_1550BP | sEA | Taiwan | ancient | 1550 | N.A. | 24.33 | 121.77 | N.A. | N.A. | Wang_C_et_al_2021 | PASS |
| I14933 | Hanben_1550BP | sEA | Taiwan | ancient | 1550 | N.A. | 24.33 | 121.77 | N.A. | N.A. | Wang_C_et_al_2021 | PASS |
| I14934 | Hanben_1550BP | sEA | Taiwan | ancient | 1550 | N.A. | 24.33 | 121.77 | N.A. | N.A. | Wang_C_et_al_2021 | PASS |
| I8076 | Hanben_1550BP | sEA | Taiwan | ancient | 1550 | N.A. | 24.33 | 121.77 | N.A. | N.A. | Wang_C_et_al_2021 | PASS |
| I14925 | Hanben_1550BP | sEA | Taiwan | ancient | 1550 | N.A. | 24.33 | 121.77 | N.A. | N.A. | Wang_C_et_al_2021 | imiss |
| I3614 | Hanben_1550BP | sEA | Taiwan | ancient | 1550 | N.A. | 24.33 | 121.77 | N.A. | N.A. | Wang_C_et_al_2021 | PASS |
| I3728 | Hanben_1550BP | sEA | Taiwan | ancient | 1496 | N.A. | 24.33 | 121.77 | N.A. | N.A. | Wang_C_et_al_2021 | PASS |
| I3619 | Hanben_1550BP | sEA | Taiwan | ancient | 1473 | N.A. | 24.33 | 121.77 | N.A. | N.A. | Wang_C_et_al_2021 | PASS |
| I8075 | Hanben_1550BP | sEA | Taiwan | ancient | 1459 | N.A. | 24.33 | 121.77 | N.A. | N.A. | Wang_C_et_al_2021 | PASS |
| I13695 | Hanben_1550BP | sEA | Taiwan | ancient | 1459 | N.A. | 24.33 | 121.77 | N.A. | N.A. | Wang_C_et_al_2021 | PASS |
| I8081 | Hanben_1550BP | sEA | Taiwan | ancient | 1409 | N.A. | 24.33 | 121.77 | N.A. | N.A. | Wang_C_et_al_2021 | PASS |
| I3732 | Hanben_1550BP | sEA | Taiwan | ancient | 1235 | N.A. | 24.33 | 121.77 | N.A. | N.A. | Wang_C_et_al_2021 | PASS |
| I8072 | Hanben_1550BP | sEA | Taiwan | ancient | 1550 | N.A. | 24.33 | 121.77 | N.A. | N.A. | Wang_C_et_al_2021 | PASS |
| I3734 | Hanben_1550BP | sEA | Taiwan | ancient | 1575 | N.A. | 24.33 | 121.77 | N.A. | N.A. | Wang_C_et_al_2021 | PASS |
| I3735 | Hanben_1550BP | sEA | Taiwan | ancient | 1532 | N.A. | 24.33 | 121.77 | N.A. | N.A. | Wang_C_et_al_2021 | PASS |
| I3612 | Hanben_1550BP | sEA | Taiwan | ancient | 1800 | N.A. | 24.33 | 121.77 | N.A. | N.A. | Wang_C_et_al_2021 | PASS |
| I3611 | Hanben_1550BP | sEA | Taiwan | ancient | 1715 | N.A. | 24.33 | 121.77 | N.A. | N.A. | Wang_C_et_al_2021 | PASS |
| I12973 | Mongolia_2500BP | nEA | Mongolia | ancient | 3226 | N.A. | 49.41 | 102.69 | N.A. | N.A. | Wang_C_et_al_2021 | PASS |
| I13174 | MongunTaiga_3350BP | nEA | Mongolia | ancient | 3410 | N.A. | 47.07 | 91.83 | N.A. | N.A. | Wang_C_et_al_2021 | PASS |
| I12976 | MongunTaiga_3350BP | nEA | Mongolia | ancient | 3332 | N.A. | 49.36 | 88.71 | N.A. | N.A. | Wang_C_et_al_2021 | PASS |
| I6363 | MongunTaiga_3050BP | nEA | Mongolia | ancient | 3095 | N.A. | 45.35 | 90.85 | N.A. | N.A. | Wang_C_et_al_2021 | PASS |
| I7033 | MongunTaiga_3050BP | nEA | Mongolia | ancient | 3066 | N.A. | 49.96 | 92.05 | N.A. | N.A. | Wang_C_et_al_2021 | PASS |
| I12955 | Munkhkhairkhan_3700BP | nEA | Mongolia | ancient | 3824 | N.A. | 49.7 | 96.8 | N.A. | N.A. | Wang_C_et_al_2021 | PASS |
| I13173 | Munkhkhairkhan_3700BP | nEA | Mongolia | ancient | 3722 | N.A. | 47.07 | 91.83 | N.A. | N.A. | Wang_C_et_al_2021 | PASS |
| I6348 | Munkhkhairkhan_3700BP | nEA | Mongolia | ancient | 3632 | N.A. | 49.39 | 102.7 | N.A. | N.A. | Wang_C_et_al_2021 | PASS |
| I12958 | Munkhkhairkhan_3500BP | nEA | Mongolia | ancient | 3483 | N.A. | 47.07 | 91.83 | N.A. | N.A. | Wang_C_et_al_2021 | PASS |
| I13964 | Ulaanzukh_3350BP | nEA | Mongolia | ancient | 3350 | N.A. | 46.77 | 111.6 | N.A. | N.A. | Wang_C_et_al_2021 | PASS |
| I12972 | Ulaanzukh_3350BP | nEA | Mongolia | ancient | 3347 | N.A. | 46.77 | 111.6 | N.A. | N.A. | Wang_C_et_al_2021 | PASS |
| I14037 | Ulaanzukh_3350BP | nEA | Mongolia | ancient | 3306 | N.A. | 46.77 | 111.6 | N.A. | N.A. | Wang_C_et_al_2021 | PASS |
| I13180 | Ulgii_4700BP | nEA | Mongolia | ancient | 4862 | N.A. | 49.3 | 88.83 | N.A. | N.A. | Wang_C_et_al_2021 | PASS |
| I12977 | Ulgii_4700BP | nEA | Mongolia | ancient | 4822 | N.A. | 49.36 | 88.71 | N.A. | N.A. | Wang_C_et_al_2021 | PASS |
| I6361 | Ulgii_4700BP | nEA | Mongolia | ancient | 4483 | N.A. | 49.34 | 88.71 | N.A. | N.A. | Wang_C_et_al_2021 | PASS |
| I6357 | SlabGrave_2850BP | nEA | Mongolia | ancient | 2200 | N.A. | 47.7 | 106.4 | N.A. | N.A. | Wang_C_et_al_2021 | PASS |
| I6364 | Mongolia_2500BP | nEA | Mongolia | ancient | 2927 | N.A. | 45.39 | 90.8 | N.A. | N.A. | Wang_C_et_al_2021 | PASS |
| I14194 | Mongolia_2500BP | nEA | Mongolia | ancient | 2834 | N.A. | 46.77 | 111.6 | N.A. | N.A. | Wang_C_et_al_2021 | PASS |

| | | | | | | | | | | | | |
|---|---|---|---|---|---|---|---|---|---|---|---|---|
| I13504 | Mongolia_2500BP | nEA | Mongolia | ancient | 2368 | N.A. | 50.7 | 99.2 | N.A. | N.A. | Wang_C_et_al_2021 | PASS |
| I13965 | Mongolia_2500BP | nEA | Mongolia | ancient | 2270 | N.A. | 48.68 | 88.38 | N.A. | N.A. | Wang_C_et_al_2021 | PASS |
| I6263 | Pazyryk_2200BP | nEA | Mongolia | ancient | 2221 | N.A. | 48.68 | 88.38 | N.A. | N.A. | Wang_C_et_al_2021 | PASS |
| I6224 | Sagly_2250BP | nEA | Mongolia | ancient | 2225 | N.A. | 49.96 | 92.05 | N.A. | N.A. | Wang_C_et_al_2021 | PASS |
| I6232 | Sagly_2250BP | nEA | Mongolia | ancient | 2239 | N.A. | 49.96 | 92.05 | N.A. | N.A. | Wang_C_et_al_2021 | PASS |
| I12970 | Sagly_2250BP | nEA | Mongolia | ancient | 2286 | N.A. | 49.96 | 92.05 | N.A. | N.A. | Wang_C_et_al_2021 | PASS |
| I7029 | Sagly_2250BP | nEA | Mongolia | ancient | 2225 | N.A. | 49.96 | 92.05 | N.A. | N.A. | Wang_C_et_al_2021 | PASS |
| I7027 | Sagly_2250BP | nEA | Mongolia | ancient | 2278 | N.A. | 49.96 | 92.05 | N.A. | N.A. | Wang_C_et_al_2021 | PASS |
| I6356 | Sagly_2250BP | nEA | Mongolia | ancient | 2250 | N.A. | 49.96 | 92.05 | N.A. | N.A. | Wang_C_et_al_2021 | PASS |
| I7030 | Sagly_2250BP | nEA | Mongolia | ancient | 2244 | N.A. | 49.96 | 92.05 | N.A. | N.A. | Wang_C_et_al_2021 | PASS |
| I7024 | Sagly_2250BP | nEA | Mongolia | ancient | 2244 | N.A. | 49.96 | 92.05 | N.A. | N.A. | Wang_C_et_al_2021 | PASS |
| I7022 | Sagly_2250BP | nEA | Mongolia | ancient | 2244 | N.A. | 49.96 | 92.05 | N.A. | N.A. | Wang_C_et_al_2021 | PASS |
| I7023 | Sagly_2250BP | nEA | Mongolia | ancient | 2226 | N.A. | 49.96 | 92.05 | N.A. | N.A. | Wang_C_et_al_2021 | PASS |
| I6233 | Sagly_2250BP | nEA | Mongolia | ancient | 2225 | N.A. | 49.96 | 92.05 | N.A. | N.A. | Wang_C_et_al_2021 | PASS |
| I6226 | Sagly_2250BP | nEA | Mongolia | ancient | 2220 | N.A. | 49.96 | 92.05 | N.A. | N.A. | Wang_C_et_al_2021 | PASS |
| I12960 | SlabGrave_2850BP | nEA | Mongolia | ancient | 3332 | N.A. | 46.92 | 102.76 | N.A. | N.A. | Wang_C_et_al_2021 | PASS |
| I12969 | SlabGrave_2850BP | nEA | Mongolia | ancient | 3001 | N.A. | 47.18 | 109.19 | N.A. | N.A. | Wang_C_et_al_2021 | PASS |
| I6352 | SlabGrave_2850BP | nEA | Mongolia | ancient | 2949 | N.A. | 46.9 | 102.77 | N.A. | N.A. | Wang_C_et_al_2021 | PASS |
| I7032 | SlabGrave_2850BP | nEA | Mongolia | ancient | 2849 | N.A. | 48.69 | 110.19 | N.A. | N.A. | Wang_C_et_al_2021 | PASS |
| I12971 | SlabGrave_2850BP | nEA | Mongolia | ancient | 2849 | N.A. | 49.15 | 114.87 | N.A. | N.A. | Wang_C_et_al_2021 | PASS |
| I13963 | SlabGrave_2850BP | nEA | Mongolia | ancient | 2829 | N.A. | 47.19 | 109.19 | N.A. | N.A. | Wang_C_et_al_2021 | PASS |
| I6349 | SlabGrave_2850BP | nEA | Mongolia | ancient | 2781 | N.A. | 45.3 | 113.85 | N.A. | N.A. | Wang_C_et_al_2021 | PASS |
| I6365 | SlabGrave_2850BP | nEA | Mongolia | ancient | 2744 | N.A. | 49.66 | 99.93 | N.A. | N.A. | Wang_C_et_al_2021 | PASS |
| I6359 | SlabGrave_2850BP | nEA | Mongolia | ancient | 2700 | N.A. | 50.12 | 100.05 | N.A. | N.A. | Wang_C_et_al_2021 | PASS |
| I6369 | SlabGrave_2850BP | nEA | Mongolia | ancient | 2554 | N.A. | 47.38 | 110.32 | N.A. | N.A. | Wang_C_et_al_2021 | PASS |
| I13175 | Xianbei_1500BP | nEA | Mongolia | ancient | 1483 | N.A. | 45.39 | 90.8 | N.A. | N.A. | Wang_C_et_al_2021 | PASS |
| I6228 | Xiongnu_1900BP | nEA | Mongolia | ancient | 1919 | N.A. | 49.96 | 92.05 | N.A. | N.A. | Wang_C_et_al_2021 | PASS |
| I1202 | Yankovsky_2850BP | nEA | Russia | ancient | 2850 | N.A. | 43.06 | 131.89 | N.A. | N.A. | Wang_C_et_al_2021 | PASS |
| I3358 | Heishui_Mohe_750BP | nEA | Russia | ancient | 769 | N.A. | 45.91 | 134.88 | N.A. | N.A. | Wang_C_et_al_2021 | PASS |
| I12974 | Mongolia_650BP | nEA | Mongolia | ancient | 682 | N.A. | 47.38 | 110.32 | N.A. | N.A. | Wang_C_et_al_2021 | PASS |
| I13961 | Mongolia_650BP | nEA | Mongolia | ancient | 609 | N.A. | 46.15 | 114.87 | N.A. | N.A. | Wang_C_et_al_2021 | PASS |
| I13176 | Mongolia_1000BP | nEA | Mongolia | ancient | 1011 | N.A. | 45.35 | 90.85 | N.A. | N.A. | Wang_C_et_al_2021 | PASS |
| Qihe3 | Qihe_11550BP | sEA | China | ancient | 11552 | N.A. | 25.4 | 117.6 | N.A. | N.A. | Wang_T_et_al_2021 | PASS |
| Longlin_1 | Longlin_10550BP | sEA | China | ancient | 10563 | N.A. | 24.64 | 105.17 | N.A. | N.A. | Wang_T_et_al_2021 | PASS |
| Dushan4_1 | Dushan_8800BP | sEA | China | ancient | 8784 | N.A. | 23.6 | 107.13 | N.A. | N.A. | Wang_T_et_al_2021 | PASS |
| Baojianshan5_M2 | Baojianshan_7350BP | sEA | China | ancient | 7368 | N.A. | 22.34 | 106.85 | N.A. | N.A. | Wang_T_et_al_2021 | PASS |
| Baojianshan5_M1 | Baojianshan_7350BP | sEA | China | ancient | 7368 | N.A. | 22.34 | 106.85 | N.A. | N.A. | Wang_T_et_al_2021 | PASS |
| BalongKD07 | BaBanQinCen_1550BP | sEA | China | ancient | 1552 | N.A. | 24.15 | 107.51 | N.A. | N.A. | Wang_T_et_al_2021 | PASS |
| BalongKD10 | BaBanQinCen_1500BP | sEA | China | ancient | 1489 | N.A. | 24.15 | 107.51 | N.A. | N.A. | Wang_T_et_al_2021 | PASS |
| QinchangKD14 | BaBanQinCen_1500BP | sEA | China | ancient | 1476 | N.A. | 24.1 | 107.51 | N.A. | N.A. | Wang_T_et_al_2021 | PASS |
| QinchangKD13 | BaBanQinCen_1500BP | sEA | China | ancient | 1442 | N.A. | 24.1 | 107.51 | N.A. | N.A. | Wang_T_et_al_2021 | PASS |

| | | | | | | | | | | | | |
|---|---|---|---|---|---|---|---|---|---|---|---|---|
| BandaKD15 | BaBanQinCen_1500BP | sEA | China | ancient | 1435 | N.A. | 24.05 | 107.5 | N.A. | N.A. | Wang_T_et_al_2021 | PASS |
| CenxunKP05 | BaBanQinCen_1500BP | sEA | China | ancient | 1387 | N.A. | 23.4 | 107.4 | N.A. | N.A. | Wang_T_et_al_2021 | PASS |
| BandaKD11 | BaBanQinCen_1500BP | sEA | China | ancient | 1387 | N.A. | 24.05 | 107.5 | N.A. | N.A. | Wang_T_et_al_2021 | PASS |
| BalongKD06 | BaBanQinCen_1500BP | sEA | China | ancient | 1500 | N.A. | 24.15 | 107.51 | N.A. | N.A. | Wang_T_et_al_2021 | imiss |
| BalongKD08 | BaBanQinCen_1500BP | sEA | China | ancient | 1500 | N.A. | 24.15 | 107.51 | N.A. | N.A. | Wang_T_et_al_2021 | imiss |
| CenxunKP13 | LaCen_1350BP | sEA | China | ancient | 1411 | N.A. | 23.4 | 107.4 | N.A. | N.A. | Wang_T_et_al_2021 | PASS |
| LadaKH01 | LaCen_1350BP | sEA | China | ancient | 1388 | N.A. | 24 | 107.6 | N.A. | N.A. | Wang_T_et_al_2021 | PASS |
| CenxunKP07 | LaCen_1350BP | sEA | China | ancient | 1330 | N.A. | 23.4 | 107.4 | N.A. | N.A. | Wang_T_et_al_2021 | PASS |
| LayiKD01 | Layi_1450BP | sEA | China | ancient | 1468 | N.A. | 24.08 | 107.62 | N.A. | N.A. | Wang_T_et_al_2021 | PASS |
| ShenxianKP09 | Shenxian_1300BP | sEA | China | ancient | 1315 | N.A. | 23.31 | 107.59 | N.A. | N.A. | Wang_T_et_al_2021 | PASS |
| YiyangKP17 | Yiyang_1350BP | sEA | China | ancient | 1387 | N.A. | 23.33 | 107.5 | N.A. | N.A. | Wang_T_et_al_2021 | PASS |
| HuatuyanNL06 | GaoHuaHua_450BP | sEA | China | ancient | 500 | N.A. | 25.1 | 107.66 | N.A. | N.A. | Wang_T_et_al_2021 | PASS |
| HuatuyanNL18 | GaoHuaHua_450BP | sEA | China | ancient | 500 | N.A. | 25.1 | 107.66 | N.A. | N.A. | Wang_T_et_al_2021 | PASS |
| HuatuyanNL04 | GaoHuaHua_450BP | sEA | China | ancient | 500 | N.A. | 25.1 | 107.66 | N.A. | N.A. | Wang_T_et_al_2021 | imiss |
| Yinwang | GaoHuaHua_450BP | sEA | China | ancient | 500 | N.A. | 23.82 | 107.52 | N.A. | N.A. | Wang_T_et_al_2021 | imiss |
| HuatuyanNL11 | GaoHuaHua_450BP | sEA | China | ancient | 446 | N.A. | 25.1 | 107.66 | N.A. | N.A. | Wang_T_et_al_2021 | PASS |
| HuatuyanNL17 | GaoHuaHua_450BP | sEA | China | ancient | 434 | N.A. | 25.1 | 107.66 | N.A. | N.A. | Wang_T_et_al_2021 | PASS |
| GaofengNL23 | GaoHuaHua_450BP | sEA | China | ancient | 421 | N.A. | 25.11 | 107.7 | N.A. | N.A. | Wang_T_et_al_2021 | imiss |
| HuaqiaoNL26 | GaoHuaHua_450BP | sEA | China | ancient | 419 | N.A. | 25.11 | 107.66 | N.A. | N.A. | Wang_T_et_al_2021 | PASS |
| HuatuyanNL21 | GaoHuaHua_450BP | sEA | China | ancient | 405 | N.A. | 25.1 | 107.66 | N.A. | N.A. | Wang_T_et_al_2021 | PASS |
| HuatuyanNL02 | GaoHuaHua_450BP | sEA | China | ancient | 386 | N.A. | 25.1 | 107.66 | N.A. | N.A. | Wang_T_et_al_2021 | PASS |
| HuatuyanNL19 | GaoHuaHua_450BP | sEA | China | ancient | 375 | N.A. | 25.1 | 107.66 | N.A. | N.A. | Wang_T_et_al_2021 | PASS |
| La368 | PhaFaen_7000BP | MSEA | Laos | ancient | 7040 | N.A. | 18.55 | 103.52 | N.A. | N.A. | McColl_et_al_2018 | PASS |
| Ma911 | GuaChaCave_3900BP | MSEA | Malaysia | ancient | 3872 | N.A. | 5.15 | 101.69 | N.A. | N.A. | McColl_et_al_2018 | PASS |
| Ma912 | GuaChaCave_2400BP | MSEA | Malaysia | ancient | 2409 | N.A. | 5.15 | 101.69 | N.A. | N.A. | McColl_et_al_2018 | PASS |
| I0627 | ManBac_3800BP | MSEA | Vietnam | ancient | 3963 | N.A. | 20.54 | 106.41 | N.A. | N.A. | Lipson_et_al_2018 | PASS |
| I1137 | ManBac_3800BP | MSEA | Vietnam | ancient | 3763 | N.A. | 20.54 | 106.41 | N.A. | N.A. | Lipson_et_al_2018 | PASS |
| I1859 | ManBac_3800BP | MSEA | Vietnam | ancient | 3765 | N.A. | 20.54 | 106.41 | N.A. | N.A. | Lipson_et_al_2018 | PASS |
| I2731 | ManBac_3800BP | MSEA | Vietnam | ancient | 3718 | N.A. | 20.54 | 106.41 | N.A. | N.A. | Lipson_et_al_2018 | PASS |
| I2947_new | ManBac_3800BP | MSEA | Vietnam | ancient | 3750 | N.A. | 20.54 | 106.41 | N.A. | N.A. | Lipson_et_al_2018 | PASS |
| Vt778 | NamTun_2550BP | MSEA | Vietnam | ancient | 2549 | N.A. | 22.42 | 102.32 | N.A. | N.A. | McColl_et_al_2018 | PASS |
| Vt777 | MaiDaDieu_2300BP | MSEA | Vietnam | ancient | 2275 | N.A. | 20.13 | 105.32 | N.A. | N.A. | McColl_et_al_2018 | PASS |
| Vt833 | MaiDaDieu_3800BP | MSEA | Vietnam | ancient | 3788 | N.A. | 20.13 | 105.32 | N.A. | N.A. | McColl_et_al_2018 | PASS |
| Vt880 | HonHaiCoTien_4000BP | MSEA | Vietnam | ancient | 4000 | N.A. | 21.4 | 107.47 | N.A. | N.A. | McColl_et_al_2018 | PASS |
| La364 | TamPaLing_2850BP | MSEA | Laos | ancient | 2865 | N.A. | 20.77 | 104.01 | N.A. | N.A. | McColl_et_al_2018 | PASS |
| La727 | TamHang_2200BP | MSEA | Laos | ancient | 2320 | N.A. | 20.77 | 104.01 | N.A. | N.A. | McColl_et_al_2018 | PASS |
| La898 | TamHang_2200BP | MSEA | Laos | ancient | 2000 | N.A. | 20.77 | 104.01 | N.A. | N.A. | McColl_et_al_2018 | PASS |
| I4011_new | Oakaie_2950BP | MSEA | Myanmar | ancient | 2950 | N.A. | 22.76 | 95.39 | N.A. | N.A. | Lipson_et_al_2018 | PASS |
| In661 | LoyangUjungCave_2000BP | MSEA | Indonesia | ancient | 1917 | N.A. | 3.85 | 96.47 | N.A. | N.A. | McColl_et_al_2018 | PASS |
| In662 | LoyangUjungCave_2000BP | MSEA | Indonesia | ancient | 2152 | N.A. | 3.85 | 96.47 | N.A. | N.A. | McColl_et_al_2018 | PASS |
| Vt779 | NuiNap_2200BP | MSEA | Vietnam | ancient | 2242 | N.A. | 19.98 | 105.69 | N.A. | N.A. | McColl_et_al_2018 | PASS |

| | | | | | | | | | | | | |
|---|---|---|---|---|---|---|---|---|---|---|---|---|
| Vt781 | NuiNap_2200BP | MSEA | Vietnam | ancient | 2248 | N.A. | 19.98 | 105.69 | N.A. | N.A. | McColl_et_al_2018 | PASS |
| Vt796 | NuiNap_2200BP | MSEA | Vietnam | ancient | 2143 | N.A. | 19.98 | 105.69 | N.A. | N.A. | McColl_et_al_2018 | PASS |
| Vt808 | NuiNap_2200BP | MSEA | Vietnam | ancient | 2255 | N.A. | 19.98 | 105.69 | N.A. | N.A. | McColl_et_al_2018 | PASS |
| I2497 | NuiNap_2200BP | MSEA | Vietnam | ancient | 2000 | N.A. | 19.98 | 105.69 | N.A. | N.A. | Lipson_et_al_2018 | PASS |
| I2948_new | NuiNap_1950BP | MSEA | Vietnam | ancient | 1948 | N.A. | 19.98 | 105.69 | N.A. | N.A. | Lipson_et_al_2018 | PASS |
| I1680_new | VatKomnou_1800BP | MSEA | Cambodia | ancient | 1810 | N.A. | 10.98 | 104.97 | N.A. | N.A. | Lipson_et_al_2018 | PASS |
| Th519 | LongLongRak_1750BP | MSEA | Thailand | ancient | 1792 | N.A. | 19.55 | 98.27 | N.A. | N.A. | McColl_et_al_2018 | PASS |
| Th521 | LongLongRak_1750BP | MSEA | Thailand | ancient | 1785 | N.A. | 19.55 | 98.27 | N.A. | N.A. | McColl_et_al_2018 | PASS |
| Th530 | LongLongRak_1750BP | MSEA | Thailand | ancient | 1756 | N.A. | 19.55 | 98.27 | N.A. | N.A. | McColl_et_al_2018 | PASS |
| Th531 | LongLongRak_1750BP | MSEA | Thailand | ancient | 1687 | N.A. | 19.55 | 98.27 | N.A. | N.A. | McColl_et_al_2018 | PASS |
| Th703 | LongLongRak_1750BP | MSEA | Thailand | ancient | 1758 | N.A. | 19.55 | 98.27 | N.A. | N.A. | McColl_et_al_2018 | PASS |
| Vt719 | HonHaiCoTien_200BP | MSEA | Vietnam | ancient | 223 | N.A. | 21.4 | 107.47 | N.A. | N.A. | McColl_et_al_2018 | PASS |
| Ma554 | SupuHujung_400BP | MSEA | Malaysia | ancient | 383 | N.A. | 6.7 | 116.8 | N.A. | N.A. | McColl_et_al_2018 | PASS |
| Ma555 | Kinabatagan_300BP | MSEA | Malaysia | ancient | 299 | N.A. | 5.08 | 117.87 | N.A. | N.A. | McColl_et_al_2018 | PASS |
| LIA001002.TF1.1 | Liang_Bua_2600BP | ISEA | Indonesia | ancient | 2600 | N.A. | -8.53 | 120.44 | N.A. | N.A. | Oliveira_et_al_2021 | PASS |
| TanjungPinang1.TF | Tanjung_Pinang_2100BP | ISEA | Indonesia | ancient | 2100 | N.A. | 2 | 128.42 | N.A. | N.A. | Oliveira_et_al_2021 | PASS |
| TanjungPinang2.TF | Tanjung_Pinang_2100BP | ISEA | Indonesia | ancient | 2100 | N.A. | 2 | 128.42 | N.A. | N.A. | Oliveira_et_al_2021 | PASS |
| TanjungPinang4.TF | Tanjung_Pinang_2100BP | ISEA | Indonesia | ancient | 2100 | N.A. | 2 | 128.42 | N.A. | N.A. | Oliveira_et_al_2021 | PASS |
| TanjungPinang6.TF | Tanjung_Pinang_2100BP | ISEA | Indonesia | ancient | 2100 | N.A. | 2 | 128.42 | N.A. | N.A. | Oliveira_et_al_2021 | PASS |
| Uattamdi1.TF | Uattamdi_1900BP | ISEA | Indonesia | ancient | 1900 | N.A. | 0.05 | 127.41 | N.A. | N.A. | Oliveira_et_al_2021 | PASS |
| AMA001.B0101.TF1.1 | Aru_Manara_2150BP | ISEA | Indonesia | ancient | 2150 | N.A. | 2.3 | 128.8 | N.A. | N.A. | Oliveira_et_al_2021 | PASS |
| AMA0038.A0101.TF1.1 | Aru_Manara_2150BP | ISEA | Indonesia | ancient | 2150 | N.A. | 2.3 | 128.8 | N.A. | N.A. | Oliveira_et_al_2021 | PASS |
| AMA004.AB0101.TF1.1 | Aru_Manara_2150BP | ISEA | Indonesia | ancient | 2150 | N.A. | 2.3 | 128.8 | N.A. | N.A. | Oliveira_et_al_2021 | PASS |
| AMA005.A0101.TF1.1 | Aru_Manara_2150BP | ISEA | Indonesia | ancient | 2150 | N.A. | 2.3 | 128.8 | N.A. | N.A. | Oliveira_et_al_2021 | PASS |
| AMA009.AB0101.TF1.1 | Aru_Manara_950BP | ISEA | Indonesia | ancient | 950 | N.A. | 2.3 | 128.8 | N.A. | N.A. | Oliveira_et_al_2021 | PASS |
| TOP002.A0101.TF1.1 | Topogaro_250BP | ISEA | Indonesia | ancient | 250 | N.A. | -2.2 | 121.66 | N.A. | N.A. | Oliveira_et_al_2021 | PASS |
| TOP004.A0101.TF1.1 | Topogaro_250BP | ISEA | Indonesia | ancient | 250 | N.A. | -2.2 | 121.66 | N.A. | N.A. | Oliveira_et_al_2021 | PASS |
| JAB001.A0101.TF1.1 | Jareng_Bori_450BP | ISEA | Indonesia | ancient | 450 | N.A. | -8.42 | 124.12 | N.A. | N.A. | Oliveira_et_al_2021 | PASS |
| KMO001.A0101.TF1.1 | Komodo_750BP | ISEA | Indonesia | ancient | 750 | N.A. | -8.6 | 119.44 | N.A. | N.A. | Oliveira_et_al_2021 | PASS |
| LIT001.A0101.TF1.1 | Liang_Toge_850BP | ISEA | Indonesia | ancient | 850 | N.A. | -8.65 | 120.99 | N.A. | N.A. | Oliveira_et_al_2021 | PASS |
| Lapita_I1368 | Vanuatu_2900BP | Oceania | Vanuatu | ancient | 2983 | N.A. | -17.79 | 168.37 | N.A. | N.A. | Skoglund_et_al_2016 | PASS |
| Lapita_I1369 | Vanuatu_2900BP | Oceania | Vanuatu | ancient | 3045 | N.A. | -17.79 | 168.37 | N.A. | N.A. | Skoglund_et_al_2016 | PASS |
| Lapita_I1370 | Vanuatu_2900BP | Oceania | Vanuatu | ancient | 3083 | N.A. | -17.79 | 168.37 | N.A. | N.A. | Skoglund_et_al_2016 | PASS |
| Lapita_Sk10 | Tonga_2600BP | Oceania | Tonga | ancient | 2594 | N.A. | -21.18 | -175.12 | N.A. | N.A. | Skoglund_et_al_2016 | PASS |
| TON001 | Tonga_2600BP | Oceania | Tonga | ancient | 2625 | N.A. | -21.18 | 175.11 | N.A. | N.A. | Posth_et_al_2018 | PASS |
| TON002 | Tonga_2600BP | Oceania | Tonga | ancient | 2625 | N.A. | -21.18 | 175.11 | N.A. | N.A. | Posth_et_al_2018 | PASS |
| MAL006 | Vanuatu_2400BP | Oceania | Vanuatu | ancient | 2665 | N.A. | -16.07 | 167.45 | N.A. | N.A. | Posth_et_al_2018 | imiss |
| MAL004 | Vanuatu_2400BP | Oceania | Vanuatu | ancient | 2567 | N.A. | -16.07 | 167.45 | N.A. | N.A. | Posth_et_al_2018 | PASS |
| MAL002 | Vanuatu_2400BP | Oceania | Vanuatu | ancient | 2525 | N.A. | -16.07 | 167.45 | N.A. | N.A. | Posth_et_al_2018 | PASS |
| TAN002 | Vanuatu_2400BP | Oceania | Vanuatu | ancient | 2490 | N.A. | -19.33 | 169.34 | N.A. | N.A. | Posth_et_al_2018 | PASS |
| MAL001 | Vanuatu_2400BP | Oceania | Vanuatu | ancient | 2240 | N.A. | -15.9 | 167.3 | N.A. | N.A. | Posth_et_al_2018 | PASS |

| | | | | | | | | | | | | |
|---|---|---|---|---|---|---|---|---|---|---|---|---|
| MAL008 | Vanuatu_2400BP | Oceania | Vanuatu | ancient | 2238 | N.A. | -16.07 | 167.45 | N.A. | N.A. | Posth_et_al_2018 | imiss |
| MAL007 | Vanuatu_2400BP | Oceania | Vanuatu | ancient | 2038 | N.A. | -16.07 | 167.45 | N.A. | N.A. | Posth_et_al_2018 | PASS |
| FUT002 | Vanuatu_1200BP | Oceania | Vanuatu | ancient | 1238 | N.A. | -19.52 | 170.23 | N.A. | N.A. | Posth_et_al_2018 | PASS |
| FUT006 | Vanuatu_1200BP | Oceania | Vanuatu | ancient | 1209 | N.A. | -19.52 | 170.23 | N.A. | N.A. | Posth_et_al_2018 | PASS |
| FUT007 | Vanuatu_1200BP | Oceania | Vanuatu | ancient | 1204 | N.A. | -19.52 | 170.23 | N.A. | N.A. | Posth_et_al_2018 | PASS |
| FUT001 | Vanuatu_1200BP | Oceania | Vanuatu | ancient | 1159 | N.A. | -19.52 | 170.23 | N.A. | N.A. | Posth_et_al_2018 | PASS |
| LHA001 | Tonga_800BP | Oceania | Tonga | ancient | 844 | N.A. | -21.18 | -175.12 | N.A. | N.A. | Posth_et_al_2018 | PASS |
| MAI002 | Solomon_Islands_500BP | Oceania | Solomon_Islands | ancient | 467 | N.A. | -9.25 | 161.22 | N.A. | N.A. | Posth_et_al_2018 | PASS |
| TAP003 | French_Polynesia_400BP | Oceania | French_Polynesia | ancient | 366 | N.A. | -16.84 | -151.36 | N.A. | N.A. | Posth_et_al_2018 | PASS |
| TAP004 | French_Polynesia_200BP | Oceania | French_Polynesia | ancient | 228 | N.A. | -16.84 | -151.36 | N.A. | N.A. | Posth_et_al_2018 | PASS |
| TAP002 | French_Polynesia_200BP | Oceania | French_Polynesia | ancient | 203 | N.A. | -16.84 | -151.36 | N.A. | N.A. | Posth_et_al_2018 | PASS |
| TAN001 | Vanuatu_300BP | Oceania | Vanuatu | ancient | 202 | N.A. | -19.56 | 169.28 | N.A. | N.A. | Posth_et_al_2018 | PASS |
| I1370 | Vanuatu_2900BP | Oceania | Vanuatu | ancient | 2945 | N.A. | -17.79 | 168.37 | N.A. | N.A. | Lipson_et_al_2018_b | imiss |
| I1369 | Vanuatu_2900BP | Oceania | Vanuatu | ancient | 2885 | N.A. | -17.79 | 168.37 | N.A. | N.A. | Lipson_et_al_2018_b | imiss |
| I1368 | Vanuatu_2900BP | Oceania | Vanuatu | ancient | 2870 | N.A. | -17.79 | 168.37 | N.A. | N.A. | Lipson_et_al_2018_b | imiss |
| I5951 | Vanuatu_2900BP | Oceania | Vanuatu | ancient | 2820 | N.A. | -17.79 | 168.37 | N.A. | N.A. | Lipson_et_al_2018_b | PASS |
| I4451 | Vanuatu_2400BP | Oceania | Vanuatu | ancient | 2260 | N.A. | -17.69 | 168.29 | N.A. | N.A. | Lipson_et_al_2018_b | PASS |
| I3921 | Vanuatu_1200BP | Oceania | Vanuatu | ancient | 1260 | N.A. | -16.71 | 168.13 | N.A. | N.A. | Lipson_et_al_2018_b | PASS |
| I5259_published | Vanuatu_300BP | Oceania | Vanuatu | ancient | 480 | N.A. | -17.64 | 168.21 | N.A. | N.A. | Lipson_et_al_2018_b | PASS |
| I4450 | Vanuatu_300BP | Oceania | Vanuatu | ancient | 215 | N.A. | -17.68 | 168.53 | N.A. | N.A. | Lipson_et_al_2018_b | PASS |
| I4105 | Vanuatu_300BP | Oceania | Vanuatu | ancient | 150 | N.A. | -16.89 | 168.3 | N.A. | N.A. | Lipson_et_al_2018_b | PASS |
| I4419 | Vanuatu_300BP | Oceania | Vanuatu | ancient | 140 | N.A. | -17.8 | 168.52 | N.A. | N.A. | Lipson_et_al_2018_b | PASS |
| I4424 | Vanuatu_300BP | Oceania | Vanuatu | ancient | 140 | N.A. | -17.77 | 168.29 | N.A. | N.A. | Lipson_et_al_2018_b | PASS |
| I4106 | Vanuatu_300BP | Oceania | Vanuatu | ancient | 140 | N.A. | -16.89 | 168.3 | N.A. | N.A. | Lipson_et_al_2018_b | PASS |
| I4425 | Vanuatu_300BP | Oceania | Vanuatu | ancient | 135 | N.A. | -17.65 | 168.4 | N.A. | N.A. | Lipson_et_al_2018_b | PASS |
| I5265 | Vanuatu_2900BP | Oceania | Vanuatu | ancient | 2875 | N.A. | -17.79 | 168.39 | N.A. | N.A. | Lipson_et_al_2020 | imiss |
| I5266 | Vanuatu_2900BP | Oceania | Vanuatu | ancient | 2875 | N.A. | -17.79 | 168.39 | N.A. | N.A. | Lipson_et_al_2020 | PASS |
| I5268 | Vanuatu_2900BP | Oceania | Vanuatu | ancient | 2995 | N.A. | -17.79 | 168.39 | N.A. | N.A. | Lipson_et_al_2020 | imiss |
| I5267 | Vanuatu_2900BP | Oceania | Vanuatu | ancient | 3050 | N.A. | -17.79 | 168.39 | N.A. | N.A. | Lipson_et_al_2020 | imiss |
| I6188 | Vanuatu_2400BP | Oceania | Vanuatu | ancient | 2400 | N.A. | -17.7 | 168.27 | N.A. | N.A. | Lipson_et_al_2020 | imiss |
| I14493 | Vanuatu_300BP | Oceania | Vanuatu | ancient | 350 | N.A. | -17.64 | 168.15 | N.A. | N.A. | Lipson_et_al_2020 | PASS |
| I10969 | Vanuatu_300BP | Oceania | Vanuatu | ancient | 350 | N.A. | -17.64 | 168.15 | N.A. | N.A. | Lipson_et_al_2020 | PASS |
| I10968 | Vanuatu_300BP | Oceania | Vanuatu | ancient | 350 | N.A. | -17.64 | 168.15 | N.A. | N.A. | Lipson_et_al_2020 | PASS |
| I10967 | Vanuatu_300BP | Oceania | Vanuatu | ancient | 180 | N.A. | -17.64 | 168.2 | N.A. | N.A. | Lipson_et_al_2020 | PASS |
| I10966 | Vanuatu_300BP | Oceania | Vanuatu | ancient | 350 | N.A. | -17.64 | 168.2 | N.A. | N.A. | Lipson_et_al_2020 | PASS |
| EFE005 | Vanuatu_300BP | Oceania | Vanuatu | ancient | 234 | N.A. | -17.81 | 168.52 | N.A. | N.A. | Lipson_et_al_2020 | PASS |
| SP4210.all | Guam_2200BP | Oceania | US | ancient | 2200 | N.A. | 13.62 | 144.87 | N.A. | N.A. | Pugach_et_al_2021 | PASS |
| SP4211.all | Guam_2200BP | Oceania | US | ancient | 2200 | N.A. | 13.62 | 144.87 | N.A. | N.A. | Pugach_et_al_2021 | PASS |
| mota.SG | Mota_4470BP | Africa | Ethiopia | ancient | 4470 | N.A. | 6.797495 | 38.207852 | N.A. | N.A. | Llorente_et_al_2015 | PASS (qpAdm) |
| Ust_Ishim_published.DG | Ust_Ishim_44350BP | nEA | Russia | ancient | 44366 | N.A. | 57.7 | 71.1 | N.A. | N.A. | Fu_et_al_2014 | PASS (qpAdm) |
| Kostenki14.SG | Kostenki_38050BP | nEA | Russia | ancient | 38052 | N.A. | 51.23 | 39.3 | N.A. | N.A. | Seguin-Orlando_et_al_2014 | PASS (qpAdm) |

| | | | | | | | | | | | | |
|---|---|---|---|---|---|---|---|---|---|---|---|---|
| I1290 | Iran_10000BP | WestAsia | Iran | ancient | 9806 | N.A. | 34.45 | 48.116 | N.A. | N.A. | Lazaridis_et_al_2016 | PASS (qpAdm) |
| I1944 | Iran_10000BP | WestAsia | Iran | ancient | 9800 | N.A. | 34.45 | 48.116 | N.A. | N.A. | Narasimhan_et_al_2019 | PASS (qpAdm) |
| I1945 | Iran_10000BP | WestAsia | Iran | ancient | 9800 | N.A. | 34.45 | 48.116 | N.A. | N.A. | Narasimhan_et_al_2019 | PASS (qpAdm) |
| I1947 | Iran_10000BP | WestAsia | Iran | ancient | 9992 | N.A. | 34.45 | 48.116 | N.A. | N.A. | Narasimhan_et_al_2019 | PASS (qpAdm) |
| I1949 | Iran_10000BP | WestAsia | Iran | ancient | 10042 | N.A. | 34.45 | 48.116 | N.A. | N.A. | Narasimhan_et_al_2019 | PASS (qpAdm) |
| I1954 | Iran_10000BP | WestAsia | Iran | ancient | 10162 | N.A. | 34.45 | 48.116 | N.A. | N.A. | Narasimhan_et_al_2019 | PASS (qpAdm) |
| I1951 | Iran_10000BP | WestAsia | Iran | ancient | 9848 | N.A. | 34.45 | 48.116 | N.A. | N.A. | Narasimhan_et_al_2019 | PASS (qpAdm) |
| I7527 | Iran_10000BP | WestAsia | Iran | ancient | 9900 | N.A. | 34.45 | 48.116 | N.A. | N.A. | Narasimhan_et_al_2019 | PASS (qpAdm) |
| Yana_old.SG | Yana_31850BP | nEA | Russia | ancient | 31850 | N.A. | 70.72 | 135.42 | N.A. | N.A. | Sikora_et_al_2019 | PASS (qpAdm) |
| I0061.SG | Karelia_8450BP | nEA | Russia | ancient | 8450 | N.A. | 61.65 | 35.65 | N.A. | N.A. | Fu_et_al_2016 | PASS (qpAdm) |
| I2123 | Indus_Periphery_4500BP | WestAsia | Turkmenistan | ancient | 4211 | N.A. | 38.21228 | 62.03443 | N.A. | N.A. | Narasimhan_et_al_2019 | PASS (qpAdm) |
| I11456_published | Indus_Periphery_4500BP | WestAsia | Iran | ancient | 4500 | N.A. | 30.649857 | 61.400311 | N.A. | N.A. | Narasimhan_et_al_2019 | PASS (qpAdm) |
| I11459_published | Indus_Periphery_4500BP | WestAsia | Iran | ancient | 4699 | N.A. | 30.649857 | 61.400311 | N.A. | N.A. | Narasimhan_et_al_2019 | PASS (qpAdm) |
| I11466_published | Indus_Periphery_4500BP | WestAsia | Iran | ancient | 4200 | N.A. | 30.649857 | 61.400311 | N.A. | N.A. | Narasimhan_et_al_2019 | PASS (qpAdm) |
| I8726 | Indus_Periphery_4500BP | WestAsia | Iran | ancient | 5000 | N.A. | 30.649857 | 61.400311 | N.A. | N.A. | Narasimhan_et_al_2019 | PASS (qpAdm) |
| I8728_published | Indus_Periphery_4500BP | WestAsia | Iran | ancient | 4500 | N.A. | 30.649857 | 61.400311 | N.A. | N.A. | Narasimhan_et_al_2019 | PASS (qpAdm) |
| I11458_published | Indus_Periphery_4500BP | WestAsia | Iran | ancient | 4600 | N.A. | 30.649857 | 61.400311 | N.A. | N.A. | Narasimhan_et_al_2019 | imiss |
| USR1.SG | Upward_Sun_River_11400BP | America | US | ancient | 11425 | N.A. | 64.22 | -145.7 | N.A. | N.A. | Moreno-Mayar_et_al_2017 | PASS (qpAdm) |
| Kolyma_River.SG | Kolyma_9750BP | nEA | Russia | ancient | 9775 | N.A. | 68.6 | 159.1 | N.A. | N.A. | Sikora_et_al_2019 | PASS (qpAdm) |
| IK002.SG | Jomon_2800BP | nEA | Japan | ancient | 2782 | N.A. | 35.0183 | 137.294 | N.A. | N.A. | McColl_et_al_2018 | PASS (qpAdm) |
| RISE515.SG | Okunevo_4300BP | nEA | Russia | ancient | 4196 | N.A. | 53.156486 | 90.207811 | N.A. | N.A. | Damgaard_et_al_2018 | PASS (qpAdm) |
| RISE667.SG | Okunevo_4300BP | nEA | Russia | ancient | 4300 | N.A. | 53.156486 | 90.207811 | N.A. | N.A. | Damgaard_et_al_2018 | PASS (qpAdm) |
| RISE670.SG | Okunevo_4300BP | nEA | Russia | ancient | 3959 | N.A. | 53.156486 | 90.207811 | N.A. | N.A. | Damgaard_et_al_2018 | PASS (qpAdm) |
| RISE671.SG | Okunevo_4300BP | nEA | Russia | ancient | 4300 | N.A. | 53.156486 | 90.207811 | N.A. | N.A. | Damgaard_et_al_2018 | PASS (qpAdm) |
| RISE672.SG | Okunevo_4300BP | nEA | Russia | ancient | 4300 | N.A. | 53.156486 | 90.207811 | N.A. | N.A. | Damgaard_et_al_2018 | PASS (qpAdm) |
| RISE674.SG | Okunevo_4300BP | nEA | Russia | ancient | 4064 | N.A. | 53.156486 | 90.207811 | N.A. | N.A. | Damgaard_et_al_2018 | PASS (qpAdm) |
| RISE675.SG | Okunevo_4300BP | nEA | Russia | ancient | 4517 | N.A. | 53.708561 | 90.359808 | N.A. | N.A. | Damgaard_et_al_2018 | PASS (qpAdm) |
| RISE677.SG | Okunevo_4300BP | nEA | Russia | ancient | 4409 | N.A. | 53.708561 | 90.359808 | N.A. | N.A. | Damgaard_et_al_2018 | PASS (qpAdm) |
| RISE680.SG | Okunevo_4300BP | nEA | Russia | ancient | 4300 | N.A. | 53.708561 | 90.359808 | N.A. | N.A. | Damgaard_et_al_2018 | PASS (qpAdm) |
| RISE681.SG | Okunevo_4300BP | nEA | Russia | ancient | 4300 | N.A. | 53.708561 | 90.359808 | N.A. | N.A. | Damgaard_et_al_2018 | PASS (qpAdm) |
| RISE683.SG | Okunevo_4300BP | nEA | Russia | ancient | 3973 | N.A. | 53.708561 | 90.359808 | N.A. | N.A. | Damgaard_et_al_2018 | PASS (qpAdm) |
| RISE684.SG | Okunevo_4300BP | nEA | Russia | ancient | 4239 | N.A. | 53.708561 | 90.359808 | N.A. | N.A. | Damgaard_et_al_2018 | PASS (qpAdm) |
| RISE685.SG | Okunevo_4300BP | nEA | Russia | ancient | 4300 | N.A. | 53.708561 | 90.359808 | N.A. | N.A. | Damgaard_et_al_2018 | PASS (qpAdm) |
| RISE719.SG | Okunevo_4300BP | nEA | Russia | ancient | 4300 | N.A. | 54.371767 | 91.506856 | N.A. | N.A. | Damgaard_et_al_2018 | PASS (qpAdm) |
